# Trading acyls and swapping sugars: metabolic innovations in *Solanum* trichomes

**DOI:** 10.1101/2023.06.05.542877

**Authors:** Paul D. Fiesel, Rachel E. Kerwin, A. Daniel Jones, Robert L. Last

## Abstract

Solanaceae (nightshade family) species synthesize a remarkable array of clade- and tissue-specific specialized metabolites. Protective acylsugars, one such class of structurally diverse metabolites, are produced by AcylSugar AcylTransferases from sugars and acyl-coenzyme A esters. Published research revealed trichome acylsugars composed of glucose and sucrose cores in species across the family. In addition, acylsugars were analyzed across a small fraction of the >1200 species in the phenotypically megadiverse *Solanum* genus, with a handful containing inositol and glycosylated inositol cores. The current study sampled several dozen species across subclades of the *Solanum* to get a more detailed view of acylsugar chemodiversity. In depth characterization of acylsugars from the Clade II species *Solanum melongena* (brinjal eggplant) led to the identification of eight unusual structures with inositol or inositol glycoside cores, and hydroxyacyl chains. Liquid chromatography-mass spectrometry analysis of 31 additional species in the *Solanum* genus revealed striking acylsugar diversity with some traits restricted to specific clades and species. Acylinositols and inositol-based acyldisaccharides were detected throughout much of the genus. In contrast, acylglucoses and acylsucroses were more restricted in distribution. Analysis of tissue-specific transcriptomes and interspecific acylsugar acetylation differences led to the identification of the *S. melongena* AcylSugar AcylTransferase 3-Like 1 (SmASAT3-L1; SMEL4.1_12g015780) enzyme. This enzyme is distinct from previously characterized acylsugar acetyltransferases, which are in the ASAT4 clade, and appears to be a functionally divergent ASAT3. This study provides a foundation for investigating the evolution and function of diverse *Solanum* acylsugar structures and harnessing this diversity in breeding and synthetic biology.

## Introduction

Plants are remarkable synthetic chemists, producing a multitude of structurally complex specialized metabolites that differ from the products of general, or primary, metabolism in their lineage-specific distribution and tissue or cell type-specific biosynthesis. In contrast to the negative selection against changes in primary metabolism (for example, amino acids, energy metabolism intermediates and vitamin cofactors), less constrained evolution of specialized metabolism led to accumulation of hundreds of thousands of taxonomically restricted metabolites in broad classes (Pichersky and Gang, 2000; Zähner, 1979). Specialized metabolites play many critical roles, including abiotic and biotic stress protection (Agati and Tattini, 2010; De Moraes et al., 2001; Landry et al., 1995), pollinator attraction (Kretschmar and Baumann, 1999) and mediation of beneficial and pathogenic microbe interactions (Yu et al., 2021). These diverse and bioactive small molecules have historical and modern uses in human medicine, including the chemotherapeutics vinblastine and paclitaxel, antimalarial artemisinin, and painkillers such as morphine (Anand et al., 2019; Wink, 2015). Their distinct phylogenetic distributions and relatively rapid diversification make specialized metabolic enzymes excellent models to study evolution (Firn and Jones, 2009; Fraenkel, 1959; Pichersky and Lewinsohn, 2011).

Acylsugars are specialized metabolites produced across the Solanaceae (nightshade) family, aiding in defense against herbivores, fungi, and bacteria (Goffreda et al., 1989; Leckie et al., 2016; Luu et al., 2017; Weinhold and Baldwin, 2011). In types I- and IV-glandular trichomes, BAHD-type <u>A</u>cyl<u>S</u>ugar <u>A</u>cyl<u>T</u>ransferase (ASAT) enzymes assemble acylsugars from the basic building blocks of sugars, often sucrose, and short- to medium-length acyl chains derived from acyl-coenzyme A (acyl-CoA) esters (Fan et al., 201; Lou et al., 2021; Moghe et al., 2017; Schilmiller et al., 2015; Schilmiller et al., 2012). Despite their simple components, acylsugars exhibit remarkable chemical diversity arising from variations in sugar core composition and acyl chain length, branching pattern, position, and number (Fan et al., 2019; Fiesel et al., 2022; Ghosh et al., 2014; Hurney, 2018; Lou et al., 2021; Moghe et al., 2017; Schenck et al., 2022). For example, acylsucroses, composed of a sucrose disaccharide core, accumulate in cultivated tomato (*Solanum lycopersicum*) trichomes, while acylsucroses and acylglucoses have been observed in the trichomes of the wild tomato species, *Solanum pennellii*. Acylsugar structural variation also impacts biological activity; for example, differential oviposition deterrence was demonstrated from naturally derived acylsugar mixtures (Leckie et al., 2016).

Solanaceae acylsugars have become an exemplary model to study evolution of a diverse, biologically relevant trait in a plant family with genomic and phylogenetic resources. Utilization of this model revealed gene duplication, neofunctionalization, co-option, and loss involved in acylsugar evolution (Fan et al., 2020, 2017; Leong et al., 2019; Moghe et al., 2017). For example, neofunctionalization of an invertase-like enzyme, AcylSucrose FructoFuranosidase 1 (ASFF1) and functional divergence of core ASAT enzymes is responsible for differences in sugar core type and acyl chain positions between cultivated tomato *S. lycopersicum* and wild tomato *S. pennellii* acylsugars (Fan et al., 2017; Leong et al., 2019). Identifying mechanisms of acylsugar evolution across the broader *Solanum* genus requires a detailed understanding of acylsugar diversity and biosynthesis, which is lacking for many species.

Nearly half of the Solanaceae falls into the large (>1200 species) and phenotypically diverse *Solanum* genus (Gagnon et al., 2022). This genus is split into several major clades, including Potato, Regmandra, Dulcamaroid-Morelloid (DulMo), Valdiviense-Archaesolanum-Normania non-spiny (VANAns), and Clade II (Gagnon et al., 2022). The well-studied Potato clade contains cultivated tomato and potato and their wild relatives. The Regmandra, DulMo, and VANAns clades contain mostly non-spiny species, including the black nightshade, *S*. *nigrum.* The large Clade II contains nearly 900 chiefly spiny species, including the cultivated brinjal eggplant, *S*. *melongena* (Bohs, 2005; Bohs and Olmstead, 1997; Gagnon et al., 2022; Levin et al., 2006; PBI Solanum Project, 2022; Särkinen et al., 2013; Stern et al., 2011; Tepe et al., 2016; Weese and Bohs, 2007). To date, documentation of *Solanum* acylsugar diversity largely focused on a handful of species within the Potato clade. These efforts identified at least 38 nuclear magnetic resonance (NMR) spectroscopy-resolved acylsugar structures and liquid chromatography mass spectrometry (LC-MS) supported annotations of many more (Fan et al., 2017; Ghosh et al., 2014; Lybrand et al., 2020; Schilmiller et al., 2016). Though acylsugar reports outside of the Potato clade are limited, they reveal novel structural variants not observed among cultivated tomato relatives. For example, acylsugars with *myo*-inositol sugar cores (i.e., acylinositols) were characterized in three species: *S. nigrum*, from the DulMo clade, and *S. lanceolatum* and *S. quitoense,* from Clade II (Herrera-Salgado et al., 2005; Hurney, 2018; Leong et al., 2020; Lou et al., 2021). These discoveries highlight the benefits of a comprehensive description of *Solanum* acylsugars within the well-developed phylogenetic framework.

Here we report analysis of *Solanum* acylsugar chemical diversity in species from the relatively unexplored *Solanum* clades DulMo, VANAns, and Clade II, which together comprise ∼1000 *Solanum* species. We first established the Clade II brinjal eggplant, *S. melongena*, as a reference species. Eggplant is a major worldwide fruit crop with extensive genomic, transcriptomic, and germplasm resources (Barchi et al., 2021; Li et al., 2021; Mennella et al., 2010). We characterized eggplant trichome acylinositols, acylinositol glycosides, and acylsugars with unusual hydroxylated acyl chains using electrospray ionization LC-quadrupole time of flight-MS (ESI LC-QToF-MS) and NMR. These atypical structures likely reflect participation of enzymes that are distinct from the cultivated tomato acylsucrose pathway. Moving beyond this model organism framework, LC-MS phylogenetic screening of 31 additional Clade II, DulMo, and VANAns species led to the identification of remarkable acylsugar structural variation. LC-MS features with characteristics of acylinositols were found in 25 of the 26 acylsugar-producing species, suggesting one or a small number of evolutionary origins. In contrast, acylglucoses were detected in DulMo and VANAns species, but not in any tested Clade II species.

As a first step towards revealing the molecular basis of this extensive acylsugar diversity, we leveraged interspecific acylsugar variation and RNA sequencing of multiple eggplant tissues to identify an acylinositol biosynthetic enzyme, *S. melongena* Acylsugar AcylTransferase 3-Like 1 (SmASAT3-L1; SMEL4.1_12g015780), responsible for acetylating a triacylinositol glycoside acyl acceptor. SmASAT3-L1 exhibits different acyl-CoA substrate specificity than previously characterized ASAT3 homologs, demonstrating how gene duplication and functional divergence created acylsugar metabolic novelty in this part of the *Solanum* clade.

## Results and Discussion

### Eggplant glandular trichomes accumulate acylsugar-like metabolites

We began our investigation of *Solanum* acylsugar diversity with the brinjal eggplant *S. melongena* due to its economic importance, availability of genomic resources, and phylogenetic position within the monophyletic Eastern Hemisphere Spiny clade of Clade II (formerly known as the ‘Old World spiny clade’) (Gagnon et al., 2022). Across eight eggplant accessions (listed in Table S1), we observed glandular trichomes resembling the acylsugar-producing structures found in other *Solanum* species (Leong et al., 2020; Lou et al., 2021; Schilmiller et al., 2012) on hypocotyls, cotyledons, and the first three true leaves of young plants (Figure 1 and Figure S1). In contrast, we observed only non-glandular stellate trichomes on leaves and stems of mature eggplants (Figure S1), and these are types that typically do not accumulate acylsugars or other specialized metabolites (Levin, 1973; Wagner, 1991).

**Figure 1.**
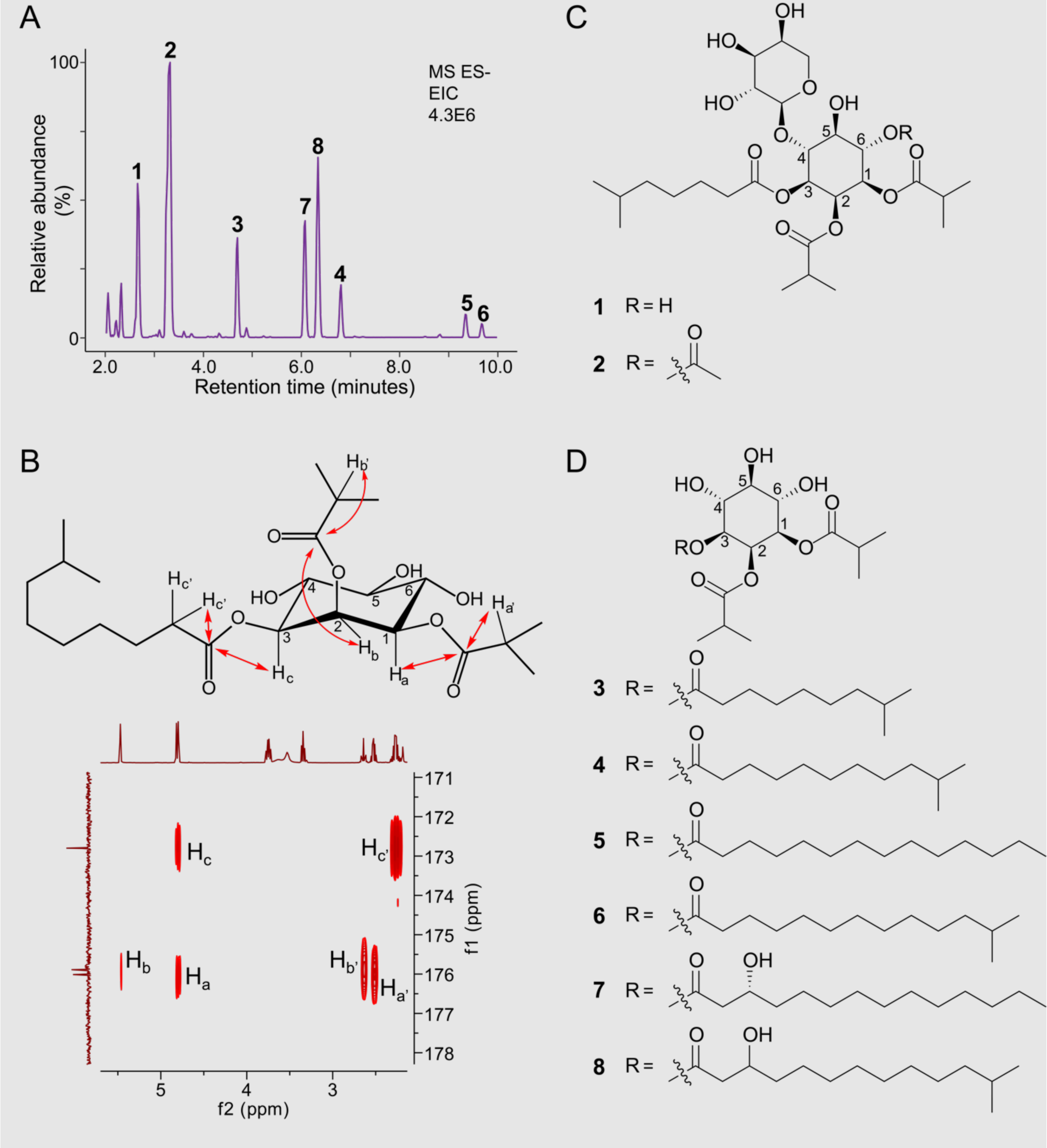
Profiling of *S. melongena* acylsugars. **(A)** LC-MS base peak intensity (BPI) chromatogram of *S. melongena* acylsugars. Peaks characterized by NMR are indicated with their compound numbers from Panels C and D. Metabolites were collected from seedling tissue with glandular trichomes similar to the ones displayed in the closeup photo of a *S. melongena* hypocotyl. **(B)** Annotation of HMBC experiments used to identify acylation positions of specific acyl chains. At the top, I3:18(4,4,10) is displayed in the chair conformation along with hydrogens important for determining acylation position. An HMBC NMR spectrum showing couplings between hydrogens and carbonyl carbons is shown. **(C)** NMR-characterized acyldisaccharide structures 1 and 2, with inositol ring carbons numbered. **(D)** NMR-characterized acylinositol monosaccharide structures 3-8.

We analyzed surface metabolite extracts from young and mature eggplant tissues using LC-QToF-MS coupled with collision-induced dissociation (CID) in negative and positive ion mode. We used these data to annotate acylsugars based on relative mass defect, molecular adduct ion masses, retention times, and ions present in CID mass spectra. Briefly, acylsugars were annotated from masses of fatty acid carboxylate fragment ions, as well as fragment ions corresponding to stepwise losses of acyl chains from the pseudomolecular ion to a sugar core fragment ion. The acylsugar annotation methods and confidence criteria are explained in detail in the Methods. This analysis revealed abundant acylsugars in extracts from young, glandular trichome-producing eggplant tissues, but not from mature, non-glandular trichome producing tissues.

We annotated 38 acylsugars in leaf surface extracts of young eggplant from eight of nine accessions (Table S1), consisting of 16 acylhexoses and 22 acyldisaccharides (Table 1). LC-MS based acylsugar annotations are described using a shorthand nomenclature modified from previous work (Leong et al., 2020) as follows: UX:Y:Z(A,B,C,D), where U – as a single or multi-letter designation – represents the sugar core, X represents the number of acyl chains, Y is the sum of acyl chain carbons, Z indicates the number of unsaturated bonds in the acyl chains (when present), and A-D represent the number of carbon atoms in the individual acyl chains. For example, the eggplant acylhexose I3:18(4,4,10) consists of a *myo*-inositol core with three acyl chains with a total of 18 carbon atoms.

**Table 1.**
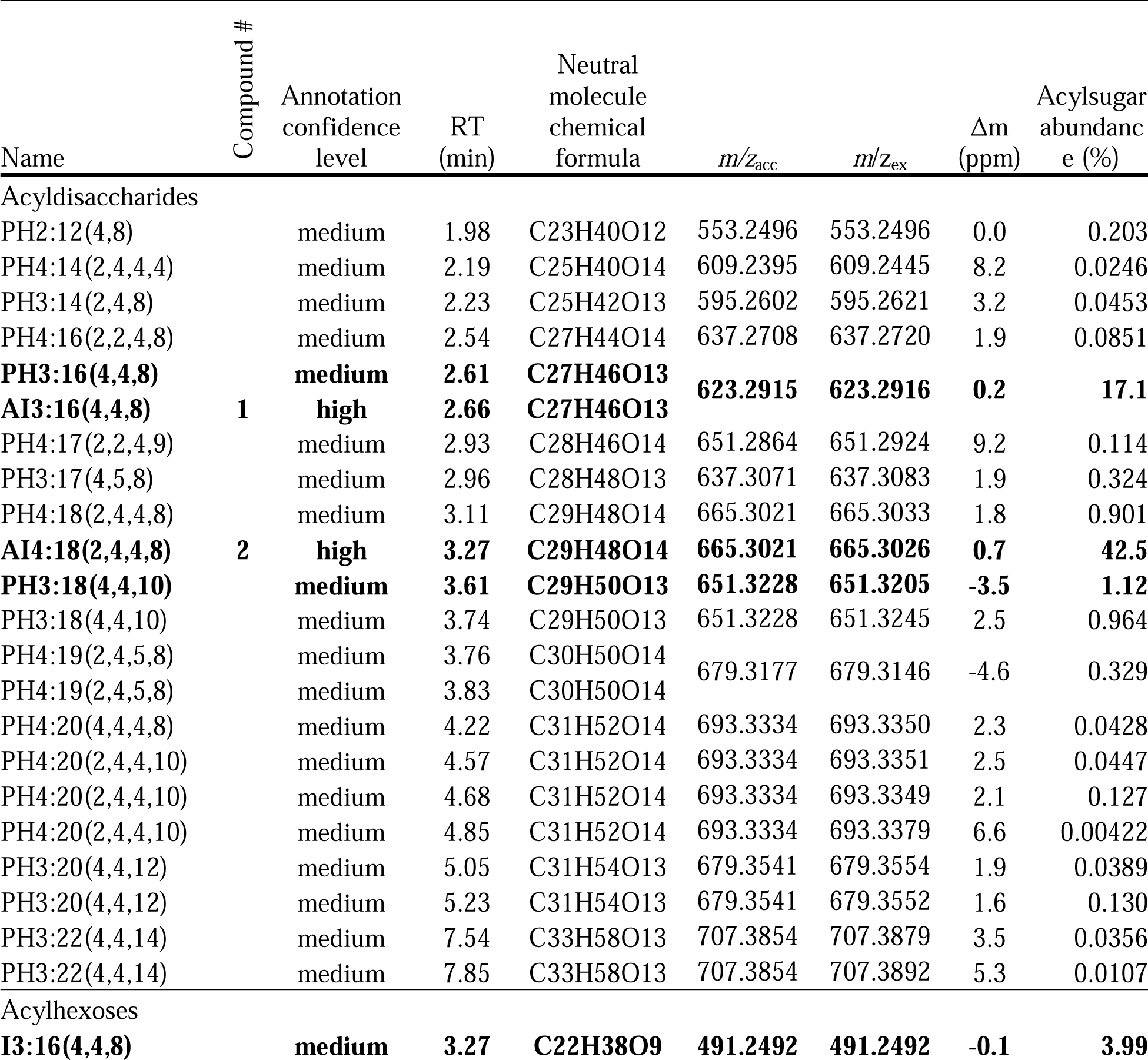
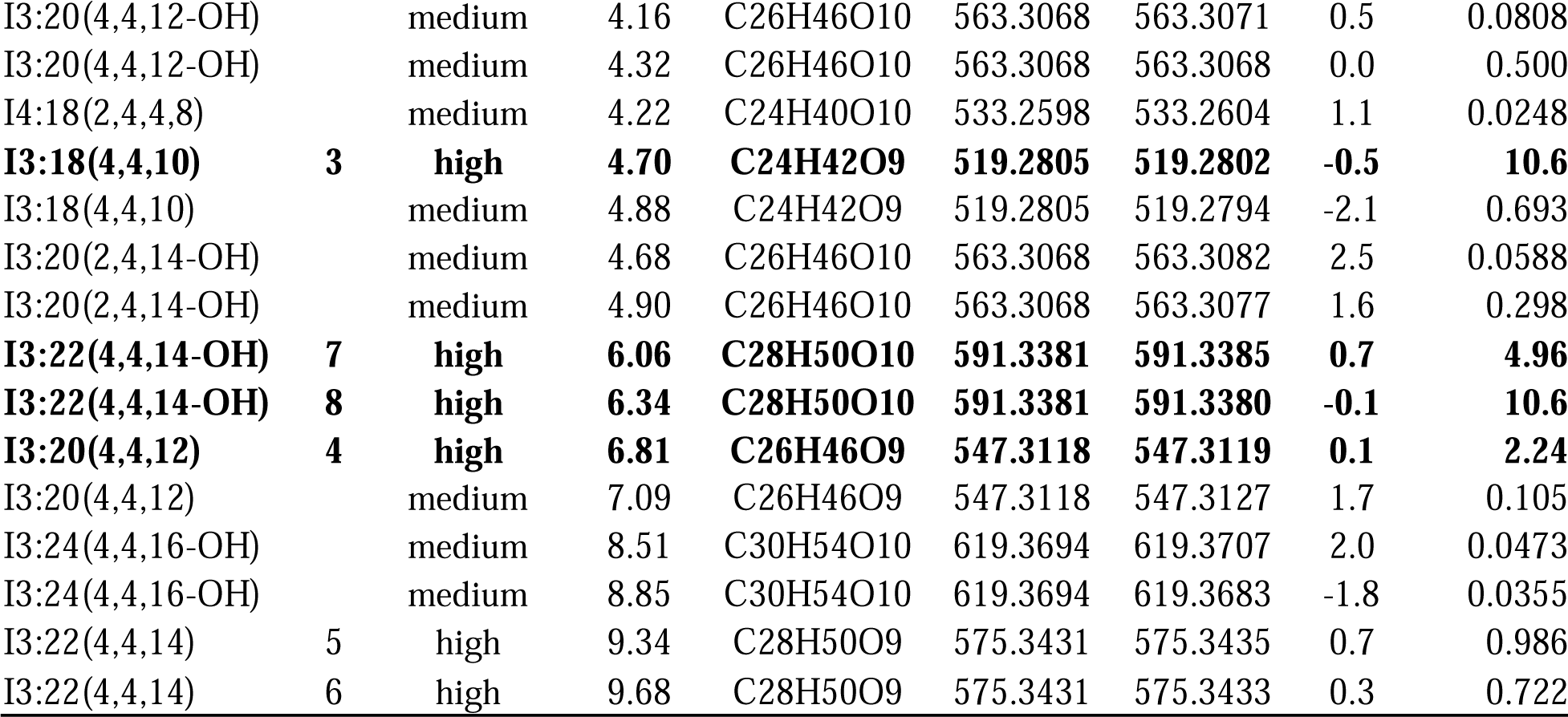
Summary of annotated acylsugars detected in *S. melongena* leaf surface extracts. PH = pentose-hexose; AI = arabinose-inositol, I = inositol. RT = retention time; *m*/*z*_acc_ = theoretical monoisotopic formate adduct mass; *m*/*z*_ex_ = experimental formate adduct mass; Δm (ppm) = mass measurement error in parts per million. Compound numbers are listed for NMR characterized compounds (Figure 1).. Acylsugars composing ≥1% of total acylsugar abundance area are in boldface text. Acylsugar abundance was calculated by the software Progenesis QI v3.0 using the sum of each compound’s adduct ion intensities (see Methods for parameters). Percent acylsugar abundance was calculated by dividing average acylsugar abundance for each compound by the sum of all acylsugar abundances, as reported in the right most column. These values were averaged over 15 samples. The annotation method is described in the Acylsugar Annotation subsection of the Methods. Acylsugars are sorted by number of sugar moieties and then by elution order. Some acylsugar isomers were unable to be deconvoluted by Progensis QI and were reported with shared m/*z*, Δm, and percent abundance values as indicated by the merged table cells.

Curiously, eggplant acyldisaccharides contain an atypical pentose-hexose core, as evidenced by a fragment ion mass of *m/z* 293.09 in negative-ion mode, corresponding to a fully deacylated sugar core minus a hydroxyl group and a proton. Further evidence was provided by positive mode CID, which promotes glycosidic bond cleavage, yielding a fragment ion corresponding to the neutral loss of an unacylated pentose ring. All acylhexoses annotated with medium to high confidence formed abundant fragment ions in positive mode but few fragment ions in negative mode as illustrated in Figure S2; this pattern is characteristic of acylinositols found in *S. quitoense* and *S. nigrum,* but has not been observed for acylglucoses, suggesting that the detected *S. melongena* acylhexoses are acylinositols (Hurney, 2018; Leong et al., 2020; Lou et al., 2021).

The eggplant acylsugars each contained three to four acylations, all on the hexose core, including one medium eight-carbon (C8) to C14 acyl chain and two to three shorter C4 or C5 acyl chains. The medium acyl chains (C8, C10, C12, and C14) were either iso-branched or straight as revealed by GC-MS acyl chain analysis (Figure S3). Additionally, we identified hydroxylated C12, C14, and C16 acyl chains not previously reported in Solanaceae acylsugars (Table 1). Although we did not observe large differences between the eight eggplant accessions, eggplant acylsugars differ in chain length, functional groups, and acyl chain composition from the reported *Solanum* acylinositols (Herrera-Salgado et al., 2005; Hurney, 2018; Lou et al., 2021).

Although LC-MS provided information about sugar core mass, it did not reveal the sugar core structure, prompting analysis using gas chromatography-mass spectrometry (GC-MS) of derivatized *S. melongena* acylsugar cores. When free sugar cores, produced by metabolite extract saponification, were derivatized to form alditol acetates, GC-MS peaks corresponding to derivatized *myo*-inositol and glucose were detected, supporting the presence of acylinositols (Figure S4A). Because the detection of glucose might have resulted from other compounds in the leaf surface metabolite extracts, the disaccharide sugar core composition was determined by hydrolyzing the saponified sugar cores with formic acid to cleave the glycosidic linkage.

Comparison of the hydrolyzed and unhydrolyzed samples revealed a peak corresponding to arabinose only in the hydrolyzed plant samples (Figure S4B). Taken together, the results of saponification with and without hydrolysis, followed by derivatization, confirmed identification of *myo*-inositol sugar cores and identified the pentose moiety of the hexose-pentose disaccharide core as arabinose.

### NMR analysis of eight eggplant acylsugars

While MS analysis provided valuable information about acyl chain number and length, sugar core mass, as well as acyl chain number, complementary information about acyl chain branch structure, sugar core stereochemistry, and acyl chain positions was obtained using NMR. We purified and resolved the structures of eight abundant eggplant acylsugars from accession PI 555598 using a combination of 1D and 2D NMR experiments (Figure 1). All eight structures are newly described, and because atomic connections were determined by NMR and MS data, the proposed structures meet Metabolomics Standards Initiative level 1 criteria for metabolite identification (Sumner et al., 2007).

A series of NMR experiments confirmed that the acylhexose and acyldisaccharide sugar cores are *myo*-inositol and 4-*O*-β-arabinopyranosyl *myo*-inositol, respectively (Figure 1C,D; Supplemental Data File S1). We assigned all sugar ring proton signals of each sugar core’s spin system with total correlation spectroscopy (TOCSY). Inferences from correlation spectroscopy (COSY) data then identified the order of ring protons and revealed pyranose and cyclitol ring structures for the pentose and hexose rings, respectively. Relative stereochemistry at ring positions was subsequently determined through comparison of spin-spin splitting (multiplicities and coupling constants) referenced to expected patterns inferred from chemical principles and previously reported acylinositols (Hurney, 2018; Leong et al., 2020). The disaccharide glycosidic linkage position was determined by heteronuclear multiple bond correlation (HMBC) correlations to be at position 4 and position 1 of *myo*-inositol and arabinose, respectively. The arabinose β anomeric configuration was inferred from the anomeric carbon ^1^*J*_CH_ (162 Hz) as revealed by a coupled-heteronuclear single quantum coherence (coupled-HSQC) experiment of the free disaccharide sugar core after saponification. Our sugar core assignments identifying this unusual disaccharide agreed with the previous GC-MS sugar core results, supporting the efficacy of the sugar core GC-MS identification method (Figure S4) This disaccharide core differs in the pentose moiety identity from a recently reported acylated 4-*O*-*ß-*xylopyranosyl *myo*-inositol in the *Solanum* Clade II species *S. quitoense* (Hurney, 2018). To our knowledge, this is the first report of acylated 4-*O*-*ß*-arabinosyl *myo*-inositol sugars.

Acyl chain positions, branching patterns, and hydroxyl positions were also resolved through integration of different NMR experiments. We found that all acyl chains were confined to *myo*-inositol, consistent with the LC-MS results. All eight acylsugars are decorated with two short iso-branched iC4 acyl chain esters at positions 1 and 2, and one medium C8 to C14 acyl chain ester at position 3 (Figure 1). Compound 2 (Figure 1C), the only tetraacylated acylsugar identified in eggplant, additionally carried an acetylation at position 6 (Figure 1C). The medium-length acyl chains at position 3 were resolved as straight (nC14) or terminally iso-branched (iC8, iC10, iC12, iC14) based on signals characteristic of protons near the branched carbons.

Strikingly, we identified unusual hydroxylated straight and iso-branched 3-hydroxytetradecanoate acyl chains, 3-OH-nC14 and 3-OH-iC14, in compounds 7 and 8, respectively (Figure 1D). We assigned hydroxylation positions of 3-OH-nC14 and 3-OH-iC14 to the third acyl carbon based on a downfield shifted signal in the ^1^H NMR spectrum at 3.92 ppm, corresponding to one hydrogen at that position. In contrast, the non-hydroxylated medium acyl chains observed in compounds 1-6 (Figure 1C,D), carry two hydrogen atoms at the third acyl carbon, and these have a characteristic signal near 1.50 ppm. We are not aware of previous reports of 3-OH-nC14 and 3-OH-iC14 chains in Solanaceae acylsugars. While 3-OH-nC14 chains are observed in the acylsugar-like bacterial Lipid A glycolipids, the hydroxylation position differentiates these eggplant chains from hydroxyacyl chains in castor bean (*Ricinus communis*) seed oil, *Silene gallica* gallicasides, *Ibicella lutea* fatty acid glycosides, and Convolvulaceae resin glycosides (Asai et al., 2010; Asai and Fujimoto, 2010; Bah and Pereda-Miranda, 1996; Smith, 1971).

### The OH-tetradecanoate acyl chain has *R* stereochemistry

To gain insight into the biosynthetic origin of the hydroxylated acyl chains, we tested chirality of the 3-OH-nC14 acyl chain of compound 7 as this chain has available commercial standards. Two-step chiral derivatization and GC-MS analysis was performed on compound 7 and two commercial standards, (3*R*)-OH-nC14 and the corresponding racemate (3*R*/*S*)-OH-nC14. Fatty acids were converted to ethyl esters to aid volatilization, followed by esterification of the acyl chain hydroxyl group with the Mosher acid (*R*)-(−)-α-methoxy-α-(trifluoromethyl)phenylacetate (*R*-MPTA) to distinguish stereoisomers by GC separation of the diastereomeric derivatives. GC-MS analysis of the derivatized (3*R*/*S*)-OH-nC14 standard compounds yielded two peaks at retention times of 42.21 and 42.40 minutes, while one peak at 42.40 minutes was obtained for (3*R*)-OH-nC14 (Figure 2). We observed expected fragment ions corresponding to the MPTA fragment ion, fatty acid ethyl ester, and fatty acid which supported peak assignment (Figure S5). Thus (3*S*)-OH-nC14 and (3*R*)-OH-nC14 elute at 42.21 and 42.40 minutes, respectively. The compound 7 acyl chain was assigned as (3*R*)-OH-nC14 as its derivatized 3-OH-tetradecanoate acyl chain eluted at 42.40 minutes (Figure 2).

**Figure 2.**
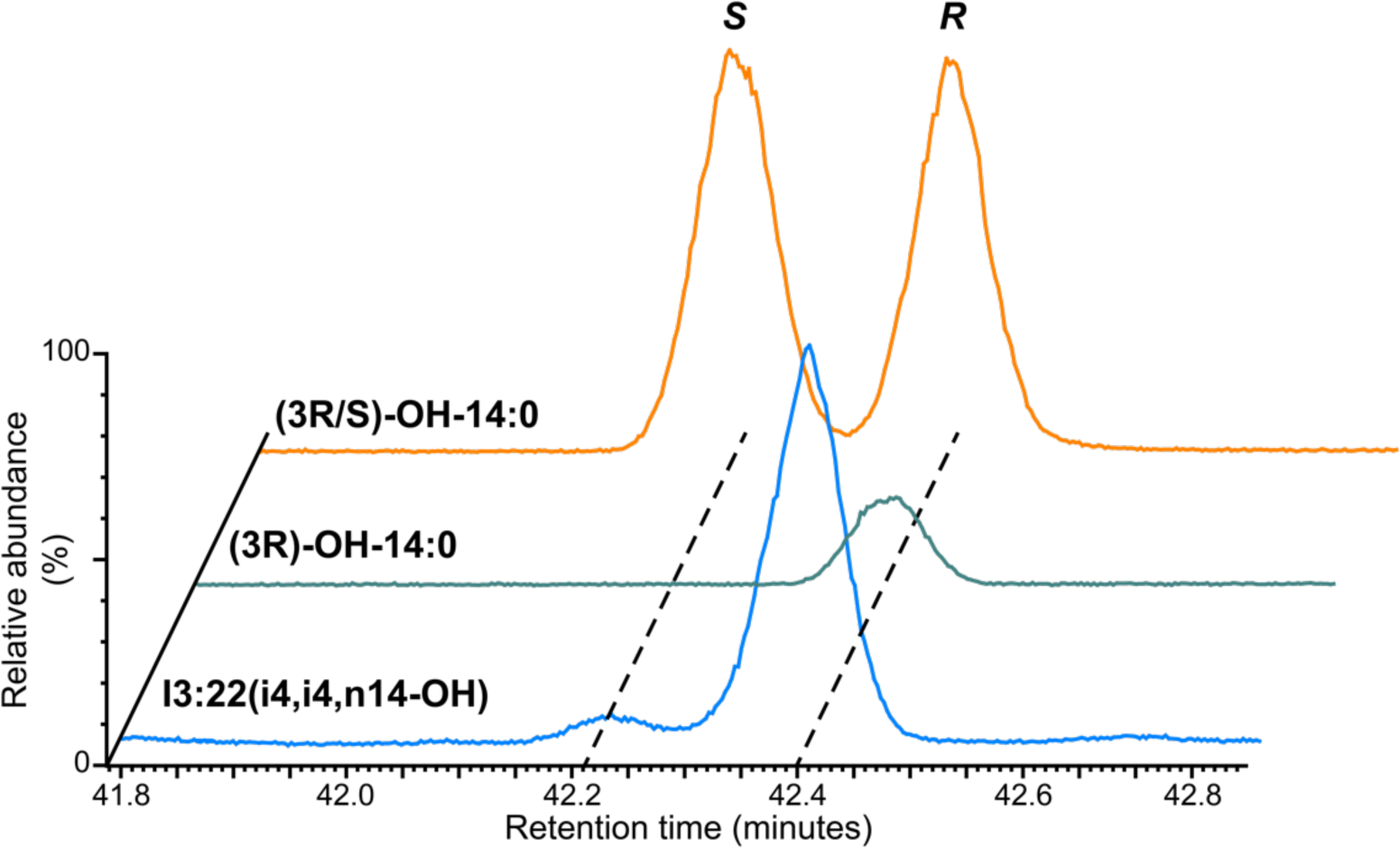
Evidence that the 3-OH-n14 acyl chain on Compound 7, I3:22(i4,i4,n14-OH) is in the *R* configuration. Hydroxyacyl chains analyzed by GC-selected ion monitoring-MS (GC-SIM-MS) after ethyl transesterification and derivatization to their diastereomeric (*S*)-MPTA form. Fragment ions with *m/z* of 189, 209, and 255 were monitored, and relative abundance was normalized to the highest peak. Yellow and green traces represent derivatives from commercial (3*R*/*S*)- and (3*R*)-OH-14:0 standards. The main derivatized 3-OH-14:0 acyl chain from Compound 7 (Figure 1), shown with the blue trace, comigrates with the peak corresponding to the *R* enantiomer. The minor peak from I3:22 is either from a small amount of endogenous *S* isomer or instrument noise.

The (3*R*)-OH-nC14 chain hydroxyl position and stereochemistry implicate possible metabolic pathways leading to its biosynthesis. As fatty acid hydroxylases typically act on the acyl chain terminal region, they are unlikely to catalyze C3 hydroxylation of a C14 fatty acid (Pinot and Beisson, 2011). Of the other well-characterized pathways, only mitochondrial and plastidial fatty acid metabolism mechanisms produce (3*R*)-OH-acyl-thioester intermediates. Further, previous work identified two trichome-expressed, mitochondrial enzymes, Acylsugar Enoyl-CoA Hydratase 1 (AECH1) and Acylsugar Acyl-CoA Synthetase 1 (AACS1), and one trichome-expressed, plastidial enzyme, a beta-ketoacyl-(acyl-carrier-protein) reductase (SpKAR1), involved in medium C10 and C12 acyl chain production in cultivated tomato and its wild relative, *S. pennellii* (Fan et al., 2020; Ji et al., 2023). We hypothesize that acylsugar hydroxyacyl chains evolved through changes in acyl-CoA substrate availability, possibly through the actions of thioesterases or acyl-CoA synthetases. Future genetic and biochemical studies should reveal the mechanisms behind (3*R*)-OH acyl chain production, potentially providing new approaches to engineer unique acylsugars with hydroxyacyl chains and hydroxylated fatty acids for polymeric building blocks.

### Enormous *Solanum* acylsugar diversity revealed by LC-MS metabolite screening

To date, most acylsugar screening across the *Solanum* has focused on cultivated tomato and its Potato clade relatives (Fan et al., 2017; Ghosh et al., 2014; Lybrand et al., 2020; Schilmiller et al., 2016), which represent a small fraction of the species diversity in this genus (PBI Solanum Project, 2022). Though limited in number, studies profiling non-Potato clade species, including *S. nigrum* (Lou et al., 2021), *S. lanceolatum* (Herrera-Salgado et al., 2005), *S. quitoense* (Hurney, 2018; Leong et al., 2020), and *S. melongena* (this work) demonstrate acylsugar diversity across the *Solanum* is vast, and largely uncharacterized. To address this knowledge gap, we analyzed tissue surface extracts from 31 additional *Solanum* species, including 25 Clade II, five DulMo clade, and one VANAns clade members (listed in Table S1). While NMR is the gold standard for structural elucidation, it is prohibitively time consuming for a largescale metabolite diversity survey. LC-MS coupled with CID provides substantial structural information and is compatible with high-throughput screening. To strike a balance between data quality and quantity, we performed LC-MS-CID screening on all 31 species and annotated acylsugars as described for *S. melongena*. Twenty-four of the 31 additional analyzed species produced detectable acylsugars from visible surface glandular trichomes, which were typically found only on seedlings, a pattern observed with eggplant. Species lacking detectable acylsugars had no or few observable glandular trichomes in the tissues analyzed, though we cannot rule out the possibility that acylsugar-producing glandular trichomes may be present on unobserved and untested tissue types and developmental stages. Targeted analysis of leaf surface extracts from 32 species, including *S. melongena*, and fruit surface extracts from two species, uncovered previously unreported acylsugars and identified the distributions of unusual acylsugar chemical traits in the *Solanum* genus (Figure 3; Tables 1 and S2-27), revealing substantial acylsugar variation between species (Figure 4; Tables 1 and S2-27).

**Figure 3.**
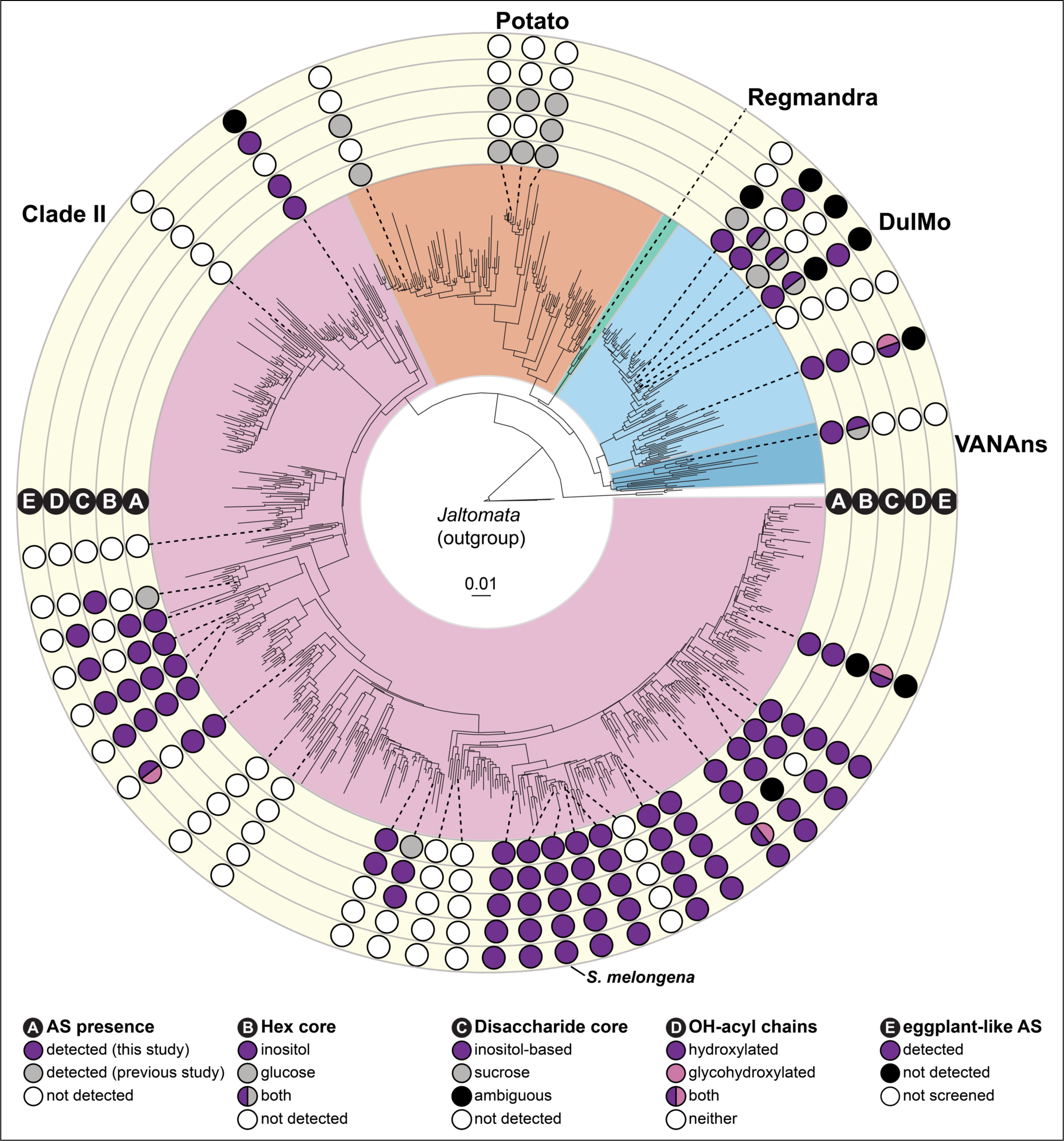
Distribution of acylsugar traits across the *Solanum* genus. Acylsugar phenotypes were mapped onto a previously published Supermatrix maximum likelihood phylogeny (Gagnon et al., 2022). Five rings (labeled **A-E**) surrounding the phylogeny represent different acylsugar traits that vary across *Solanum* species (summarized in Table S2). **A) AS presence:** presence of detectable acylsugars in aerial surface (i.e., trichome) extracts. **B) Hex core:** identity of hexose sugar core in detected acylhexoses. **C) Disaccharide core:** identity of disaccharide sugar core in detected acyldisaccharides. If acyldisaccharides were detected but the sugar core identity could not be resolved, it was categorized as ambiguous. **D) OH-acyl chains:** identity of unusual acyl chains bearing hydroxyl or glycosylated hydroxyl groups in detected acylsugars. **E) eggplant-like AS:** presence of acylsugars with the same acyl chain and sugar core composition that also coelute with eggplant acylsugars (and are therefore likely identical). Acylsugar annotations from this study are based upon LC-MS and NMR data. Annotations from previous studies were limited to those with NMR structural data.

**Figure 4.**
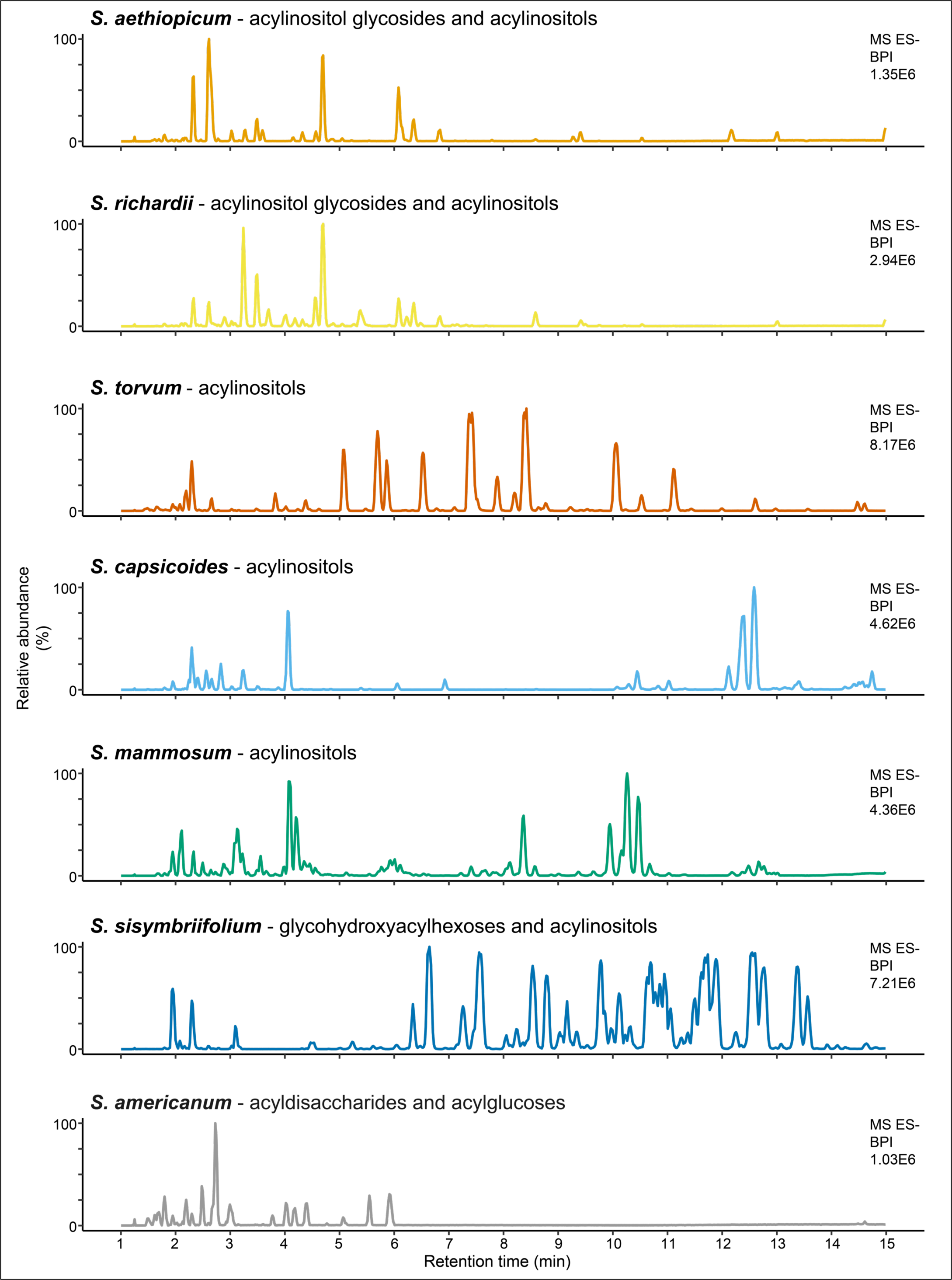
*Solanum* species produce dramatically different leaf surface metabolite profiles. The seven representative species display morphological differences, as demonstrated by the flower and fruit images, and they also exhibit diverse metabolite profiles as demonstrated by base peak intensity LC-MS chromatograms from their leaf surface metabolites. Most of the peaks shown consist of acylsugars with varied sugar cores and acyl chain types and numbers.Acylsugar classes are listed for each species. Acylsugar profiles for these species are detailed in Tables S5, S8, S18, S21, and S25-27.

### Identification of diverse acyl chain types in *Solanum* acylsugars

Our survey of *Solanum* species uncovered acylsugars with a surprising diversity of acyl chain lengths, functional groups, and combinations of chain types and positions. We identified C2 to C18 acyl chains, including unsaturated and hydroxylated forms, based on analysis of fatty acid fragment carboxylate ions in negative mode and neutral losses in both negative and positive mode (Table S2). Acyl chain compositions differed between acylsugars within and across species: acylinositols with primarily medium C8 to C14 acyl chains were observed in three species (Figure 3, Tables S14, S15, S25), while acylsugars with two short C4 to C6 acyl chains and one medium C8 to C14 acyl chain were observed in 17 species, an acylation pattern also observed in eggplant and wild tomato (Figure 1; Table 1) (Lybrand et al., 2020). Acylsugars with unsaturated acyl chains, including C5:1, C18:1, C18:2, and C18:3, were detected only in *S. torvum* (Table S18). To our knowledge, this is the first report of unsaturated C18 chains in Solanaceae acylsugars. In contrast, hydroxylated medium-length acyl chains were surprisingly common: we annotated acylsugars bearing hydroxylated acyl chains in three DulMo clade species and 18 Clade II species (Figure 3). These hydroxylated acyl chains are ‘chemical handles’ susceptible to further modifications, including glycosylations and acylations, which would result in even greater acylsugar diversity.

Indeed, we obtained evidence for glycosylated acyl chains in three Clade II species, *S. prinophyllum*, *S. sisymbriifolium*, *S. lasiophyllum*, and one DulMo species, *S. dulcamara* (Tables S12, S13, S24 and S27). Several lines of evidence led us to conclude that these species accumulate acylsugars with glycosyl groups attached via ether linkages to medium hydroxyacyl chains esterified to the hexose core. First, the compounds fragmented abnormally under negative mode CID: the glycosyl group was observed as a neutral loss from the [M-H]^-^ ion, producing a [sugar – H]^-^ fragment ion (Figure S6). This contrasts with extensive observations that CID spectra of other characterized acyldisaccharides do not exhibit negative mode disaccharide glycosidic linkage cleavage (Figure S2 and Figure S7) (Ghosh et al., 2014; Hurney, 2018; Lou et al., 2021). Further evidence that the glycosylation is not on the primary acylated hexose ring was obtained by subjecting acylsugar extracts to saponification, which induces ester linkage breakage while retaining ether linkages. Upon analyzing saponified acylsugar extracts by LC-MS, we detected pentose and hexose sugars with C14 or C16 acyl chains rather than glycosylated hexoses, indicating that the glycosyl group is connected to a medium hydroxyacyl chain by an ether linkage that is not cleaved during alkaline saponification (Figure S8). We named these compounds glycohydroxyacylhexoses and annotated them by modifying the conventional acylsugar nomenclature where the molecule in Figure S6 is named H3:24(4,4,16-O-p) with 16-O-p representing the pentosylated (p) C16 hydroxyacyl chain (16-O). To our knowledge, this is the first report of acylhexoses with glycosylated hydroxyacyl chains in Solanaceae acylsugars. Resin glycosides produced by Convolvulaceae family species contain similar hydroxyacyl chain glycosidic linkages. However, in resin glycosides, hydroxyacyl chains are connected to the oligosaccharide core at two points – by an ester and ether linkage, respectively – forming a macrocyclic structure not observed in the *Solanum* glycohydroxyacylhexoses (Bah and Pereda-Miranda, 1996; Kruse et al., 2022; Pereda-Miranda et al., 1993).

The unusual glycohydroxyacylhexoses raises questions regarding their biosynthetic and evolutionary origins. As these compounds are found in different lineages (Eastern Hemisphere Spiny, Dulcamaroid, and Sisymbriifolium sections) – and closely related species lack detectable glycohydroxyacylhexoses – this trait appears to be highly labile (Figure 3). Because we observe the cognate, non-glycosylated acylinositol in each of these species, we hypothesize that glycohydroxyacylhexoses consist of a *myo*-inositol hexose core. Further analysis of acylsugars from additional species may identify other types of modified acylsugar hydroxyacyl chains in *Solanum* acylsugars.

Sixteen species accumulate acylsugars with the same molecular masses, sugar core masses, and acyl chain complements as the NMR-characterized eggplant acylinositols, suggesting they may be identical. To test this hypothesis, we mixed purified eggplant NMR-resolved standards with metabolite extracts from each species and performed liquid co-chromatography. Coelution provided one line of evidence that the eggplant-like acylsugars detected in 11 Clade II species are identical to characterized eggplant acylinositols (Tables S5-14, S27). In contrast, the eggplant-like acylsugars from five other species, including three DulMo clade members and two Clade II members, eluted separately from characterized eggplant acylinositols, indicating they are acylation positional isomers or acyl chain branching isomers (Figure S9). Examples of acylinositol positional isomers were described between the DulMo clade species *S. nigrum* and the Clade II species *S. quitoense* and *S. melongena*, providing precedent that the non-coeluting compounds might be positional isomers of the eggplant acylsugars shown in Figure 1 (Leong et al., 2020; Lou et al., 2021). Our data suggest a phylogenetic pattern of acylinositol distribution in which Clade II species generally accumulate eggplant-like acylinositol isomers, whereas DulMo clade species accumulate non-eggplant-like acylinositol isomers (Figure 3). While the eggplant-like isomers coelute with eggplant acylsugars, further analysis with complementary analytical methods is needed to confirm whether these are structurally identical across species.

### Acylhexoses and acyldisaccharides accumulate throughout Clade II, DulMo, and VANAns *Solanum* species

Acylhexoses were detected in all analyzed acylsugar-producing species. They were annotated as acylglucoses or acylinositols based upon differences in characteristic fragmentation behavior under negative ion mode MS conditions (Leong et al., 2020; Lou et al., 2021). Identification of acylglucoses was based on observation of a sugar core fragment ion of *m*/*z* 143.03 (C H O ^-^) and stepwise acyl chain losses at the lowest collision energy level (0 V) (Figure S10). In contrast, acylinositol annotation was based on an absence of acyl chain fragment ions at 0 V and MS^e^ CID negative mode functions (Figure S2). Using these rules, we identified acylinositols in all acylsugar-producing species tested except for *S. americanum* (Tables 1, S2-27). In contrast, acylglucoses were not detected in any Clade II species but were commonly detected among the DulMo and VANAns species, found in all analyzed acylsugar-producing species except *S. dulcamara* (Tables S2, S19, S20, S23, S24, S26; Figure 3; Lou et al., 2021).

Differing from the widespread acylhexose distribution, acyldisaccharides were detected sporadically in Clade II and DulMo clade species, being observed in 15 of 21 Clade II and 2 of 4 DulMo acylsugar-producing analyzed species (Figure 3 and Table S2). Based on negative [M-H]^-^ fragment masses and positive mode fragmentation of the glycosidic linkage, the acyldisaccharides were composed of hexose-hexose, pentose-hexose, or deoxyhexose-hexose sugar cores (Table S2, Figure S2 and Figure S7). Though complete structural information cannot be obtained from MS fragmentation alone, two lines of evidence lead us to hypothesize that these acyldisaccharides are composed of glycosylated inositol cores. First, most of the species with detectable acyldisaccharides also accumulate cognate acylinositols with the same acyl chain lengths (leaf and fruit surface extracts from 9 of 15 and two of two different species, respectively), suggesting a shared inositol-containing core structure (Tables 1, S3-9, S12-17, S20, S22). Second, acylinositol glycosides with varying sugar core sizes and stereochemistries were described in *S. lanceolatum* (Herrera-Salgado et al., 2005), *S. quitoense* (Hurney, 2018), and *S. melongena* (Figure 1), suggesting that the species in this study may accumulate similar compounds. The ratio of acylhexose to acyldisaccharide peak number and abundance differs across the surveyed species; for example, *S. mammosum* only accumulated detectable acylinositols while *S. anguivi* primarily accumulated acyldisaccharides that comprised 80% of the total acylsugar peak area (Tables S21 and S4). This is reminiscent of varied acylhexose accumulation observed in trichomes of evolutionarily distant *Solanum* species: *S. nigrum* only contains detectable acylinositol and acylglucose monosaccharides (Lou et al., 2021), while *S. pennellii* accessions have mixtures of acylglucoses and acylsucroses ranging from 41-95% of total acylsugars as acylglucoses. (Lybrand et al., 2020; Shapiro et al., 1994).

### Sticky fruits accumulate acylinositols

We investigated the chemical basis of sticky fruit surface substances previously reported in botanical species descriptions of the Clade II species *S. acerifolium* and *S. atropurpureum* (Nee, 2022a, 2022b, 1991). LC-MS analysis of fruit surface extracts revealed similar acylsugar profiles between the two species, with each extract containing what appeared to be the same 128 acylsugars. These fruit acylsugars were distinct from the 22 and 19 trichome acylsugars identified in *S. acerifolium* and *S. atropurpureum*, respectively, as evidenced by their LC elution and MS characteristics (Tables S14-17; Figure 5). Seventy-seven fruit acylsugars were identified as acylinositols with two or three medium C8 to C14 acyl chains, including one to three hydroxyacyl chains. The remaining 51 fruit acylsugars are acyldisaccharides containing an unusual deoxyhexose-hexose core not previously reported in Solanaceae acylsugars. In contrast, *S. acerifolium* and *S. atropurpureum* trichome acylsugars, composed of a pentose-hexose disaccharide core decorated with short C5 or C6 acyl chains, were not detected in fruit surface extracts.

**Figure 5.**
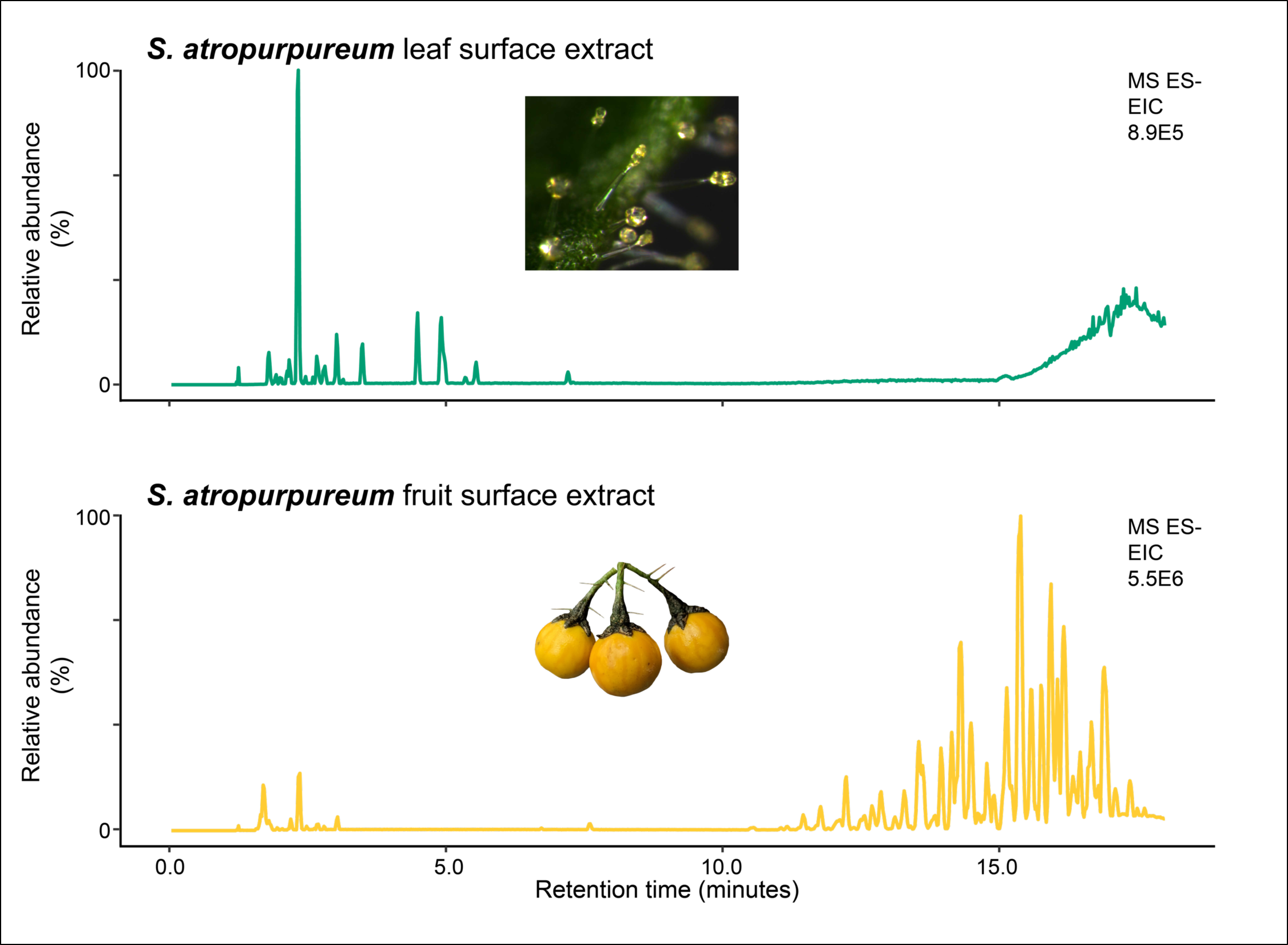
*S. atropurpureum* trichomes and fruit exhibit dramatically different metabolite profiles. The top LC-MS base peak intensity (BPI) chromatogram displays metabolites from a leaf surface extract, while the bottom LC-MS base peak intensity chromatogram displays metabolites from a fruit surface extract. The leaf surface extract acylsugars elute from 2.2 to 7.6 min (Table S17), while fruit surface extract acylsugars elute from 4.6 to 18.4 min (Table S15).

*S. mammosum* and *S. capsicoides,* which are closely related to *S. acerifolium* and *S. atropurpureum*, do not produce noticeably sticky fruit surfaces, suggesting this trait evolved in the common ancestor of *S. acerifolium* and *S. atropurpureum,* perhaps as recently as 2 Mya (Särkinen et al., 2013). Fruit surface acylsugars were previously described within the *Physalis* genus and likely represent an independent evolutionary origin from the *S. acerifolium* and *S. atropurpureum* fruit acylsugars (Bernal et al., 2018; Cao et al., 2015; Cicchetti et al., 2018; Maldonado et al., 2006). Considering that, like eggplant, *S. acerifolium* and *S. atropurpureum* fruits lack trichomes (Nee, 2022a, 2022b), some other organ or cell type presumably synthesizes the fruit surface acylsugars. Structures similar to *Solanum fernadesii* petiolar resin glands (Sampaio et al., 2021), *Hypericum androsaemum* microscopic fruit glands (Caprioli et al., 2016), and fennel and chamomile fruit secretory ducts (vittae) (Zizovic et al., 2007) may be involved in *S. acerifolium* and *S. atropurpureum* fruit acylsugar secretion. Future work will be needed to understand the cellular, biosynthetic, and evolutionary relationships between trichome and fruit acylsugars.

### Sugar core evolution across the Solanaceae

Our screening revealed acylinositols to be broadly distributed across Clade II, DulMo, and VANAns (Figure 3, Table S1). Outside of this study, acylinositols were only reported in a small number of *Solanum* species, including two Clade II members, *S. quitoense* and *S. lanceolatum*, and one DulMo clade member, *S. nigrum*. The acylinositol distribution suggests that acylinositol biosynthesis arose one or more times within the *Solanum* genus. However, our ability to elucidate the likely number of acylinositol origins is impeded by the lack of resolution of the phylogenetic relationships between the major *Solanum* clades, including Clade II, DulMo, VANAns, Potato, and Regmandra (Gagnon et al., 2022). Further biochemical and genetic analyses of acylinositol biosynthesis may reveal how this trait evolved, and whether the pathway is conserved between Clade II, DulMo, and VANAns species.

Acylglucoses were reported to be sporadically present across the Solanaceae, including in *Salpiglossis sinuata*, *Petunia*, *Nicotiana* spp., *Datura*, the DulMo clade *S. nigrum*, and Potato clade *S. pennellii* (Castillo et al., 1989; Chortyk et al., 1997; Fiesel et al., 2022; Fobes et al., 1985; Hurney, 2018; King and Calhoun, 1988; Lou et al., 2021; Matsuzaki et al., 1989; Schenck et al., 2022; Van Dam and Hare, 1998). Recent work in *S. pennellii* and *S. nigrum* revealed that acylglucoses are synthesized from acylsucroses by a neofunctionalized invertase-like enzyme, AcylSucrose FructoFuranosidase 1 (ASFF1) (Leong et al., 2019; Lou et al., 2021). These results show that conversion of acylsucroses to acylglucoses arose at least twice in *Solanum* because *S. nigrum* and *S. pennellii* employ non-orthologous ASFF1 enzymes. Our current study extends documented cases of *Solanum* acylglucose distribution to the VANAns clade, as well as documenting it in additional DulMo species. In contrast, we found no acylglucose-accumulating species among the 26 *Solanum* Clade II species analyzed in this study (Figure 3 and Table S2). Elucidation of acylglucose biosynthesis outside of the *Solanum* clade will be needed to resolve the number of evolutionary origins and biochemical mechanisms leading to synthesis of these metabolites across the Solanaceae.

Varied acyldisaccharides were observed across DulMo and Clade II, and we predict they are biosynthesized from acylinositols. Within the Potato clade as well as outside the *Solanum* genus, acylsucroses are the dominant acyldisaccharide in the reported species and tissues (Fiesel et al., 2022). In contrast, no acylsucroses were detected in any species screened in this study (Figure 3 and Table S2). However, acylsucrose biosynthesis might be found in other DulMo and VANAns species or accessions because acylglucoses were detected in all but one DulMo and VANAns species and that acylsucroses can act as intermediates in acylglucose biosynthesis (Leong et al., 2019; Lou et al., 2021). The lack of both acylglucoses and acylsucroses in the analyzed Clade II species (Figure 3 and Table S2) suggests a loss of acylsucrose biosynthesis in Clade II. Rather, our LC-MS data are consistent with the hypothesis that the acyldisaccharides observed in this study consist of glycosylated inositol cores. Recent *in vitro* biochemistry and *in vivo* genetic evidence in *S. quitoense* suggests that acylinositols are the precursors to acylinositol disaccharides in Clade II *Solanum* species trichomes (Leong et al., 2022, 2019) and tomato roots (Kerwin et al., 2024). In this scenario, acylinositols would be converted to their cognate glycosides by one or more glycosyltransferases, with different sugars added either by a conserved promiscuous glycosyltransferase or multiple glycosyltransferases. *S. quitoense* and *S. melongena*, with their acylinositol glycosides bearing different glycosyl moieties, are promising systems for addressing the biochemical origins of these unusual *Solanum* trichome acyldisaccharides.

### *S. aethiopicum* is defective in expression of a trichome acylsugar acyltransferase enzyme

We began analyzing the biochemical mechanisms underlying the observed *Solanum* acylsugar diversity in seven of 23 members from the Eggplant clade and related Anguivi grade, two small subclades within Clade II that includes brinjal eggplant and its close relatives (Figure S11) (Aubriot et al., 2018). In contrast to the other six analyzed eggplant species, *S. aethiopicum* (scarlet or Ethiopian eggplant) is unique in that it had unacetylated AI3:16(4,4,8) (Compound 1) but did not accumulate detectable levels of the acetylated form, AI4:18(2,4,4,8) – the highest abundance *S. melongena* acylsugar (Compound 2; Figure 6A, Figure S11, Tables 1, S2, S5). We hypothesized that the absence of AI4:18(2,4,4,8) in *S. aethiopicum* is either due to loss-of-function or altered expression of an acetylating enzyme in this species. The availability of *S. melongena* and *S. aethiopicum* genomic sequences (Barchi et al., 2021; Song et al., 2019) provided the opportunity to seek a trichome expressed acetyltransferase responsible for producing AI4:18 and the mechanism behind the lack of detectable AI4:18 in *S. aethiopicum*.

**Figure 6.**
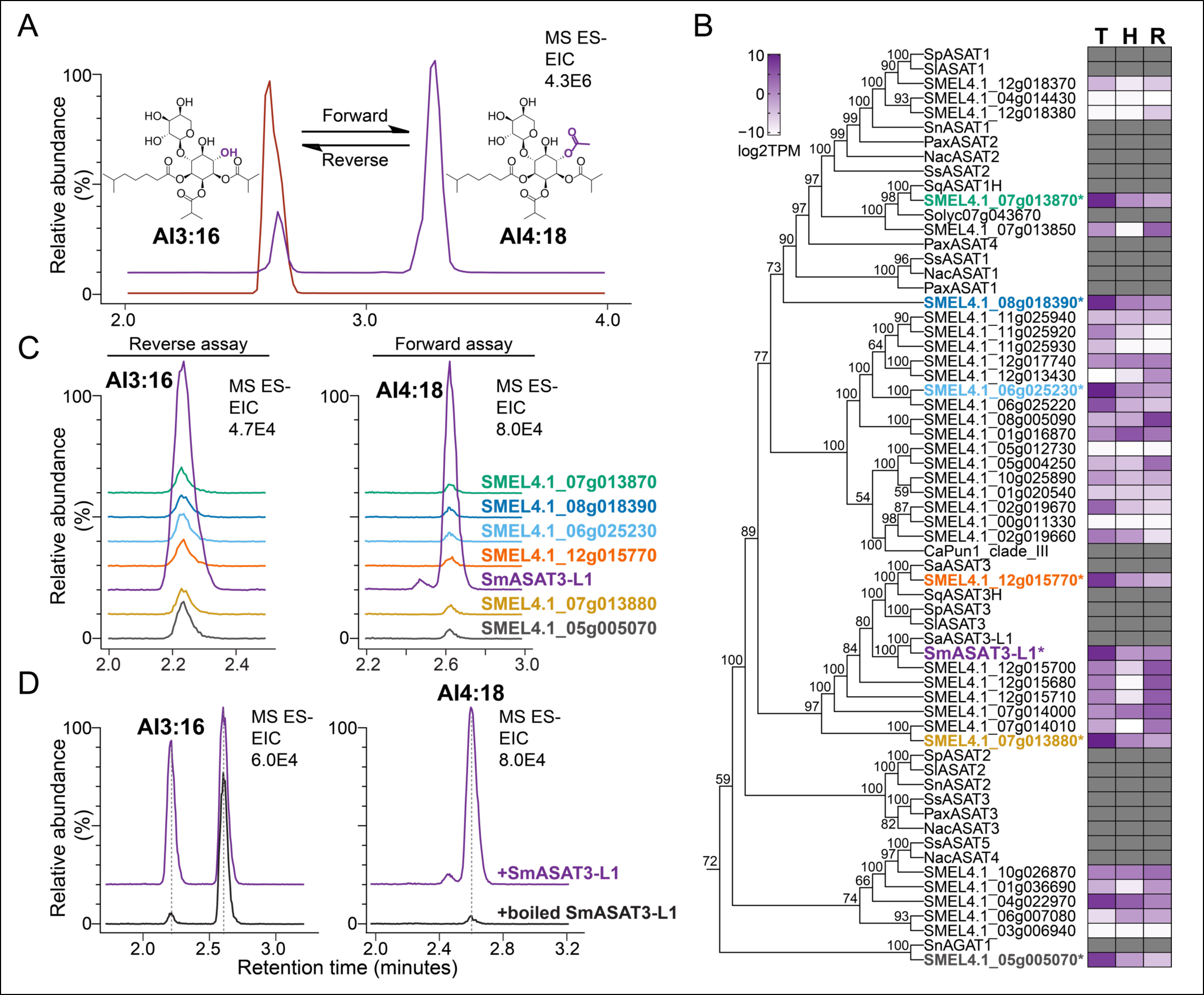
SmASAT3-L1 acetylates AI3:16 to produce AI4:18. Trichome expressed ASAT homologs were identified and tested for AI3:16 acetyl transferase activity. (A) LC-MS analysis of AI3:16(4,4,8) and AI4(2,4,4,8) in *S. aethiopicum* (red) and *S. melongena* (purple) leaf surface extracts. *S. aethiopicum* does not produce detectable levels of AI4:18. (B) Shown is a clade III BAHD tree subset from a phylogeny including 106 predicted BAHDs (PF002458) in the eggplant Smel_V4.1 reference genome, published reference BAHD sequences for clades I-VII, characterized ASAT sequences from other Solanaceae species, and the SaASAT3 and SaASAT3-L1 candidates from *S. aethiopicum* (see Figure S12 for full phylogeny). The maximum likelihood tree was inferred from amino acid sequences using the Jones-Taylor-Thornton algorithm with seven rate categories in IQ-TREE v2.1.3. Values at nodes indicate bootstrap support calculated from 100,000 ultrafast bootstrap replicates. The heatmap shows absolute transcript abundance (log2 TPM) across trichomes, trichomeless hypocotyls, and roots from 7-day-old eggplant seedlings. Expression data for non-eggplant sequences are not included in the heatmap. Color gradient provides a visual marker to rank the transcript abundance from high (purple) to low (white) or absent (grey). ASAT, acylsugar acyltransferase. log2TPM, log2 transformed transcripts per million. Genes denoted with “*” were selected for biochemical characterization based on expression in trichomes and sequence similarity to characterized ASAT enzymes in other species. (C) Of the seven enzymes tested *in vitro*, only SmASAT3-L1 successfully acetylated AI3:16(4,4,8) to form detectable AI4:18(2,4,4,8). The extracted ion chromatograms on the left display products from reverse enzyme assays and the formate adduct of AI3:16, *m/z* 623.29. The extracted ion chromatograms on the right display products from forward enzyme assays and the formate adduct of AI4:18, *m/z* 665.30. (D) Acetylation of AI3:16 *in vitro* to produce AI4:18 is abolished upon boiling of SmASAT3-L1 enzyme. Forward assay extracted ion chromatograms display the formate adduct of AI4:18, *m/z* 665.30, and reverse assay extracted ion chromatograms display the formate adduct of AI3:16, *m/z* 623.29. The reverse assay chromatograms display peaks for both AI3:16 and AI4:18 due to in source fragmentation of remaining AI4:18.

To identify enzyme candidates, we sequenced transcriptomes of isolated trichomes, trichome-depleted shaved hypocotyls, and whole roots collected from 7-day-old *S. melongena* seedlings and performed differential gene expression analysis (see Methods). Of the 23,251 eggplant genes expressed in at least one of the three tissues, 745 were significantly enriched (log2 fold-change >2, *p*-value < 0.05) in trichomes compared to hypocotyls and roots, including 20 BAHDs (Tables S37 and S38). As shown in Table S39, we selected for further testing seven BAHDs that were abundantly expressed in trichomes (TPM > 200) and homologous to characterized *Solanum* ASATs (sequences in colored text and denoted with “*” in Figure 6B) (D’Auria, 2006; Fan et al., 2016; Leong et al., 2022, 2020; Lou et al., 2021; Schenck et al., 2022). We expressed each of the seven candidates in *Escherichia coli* and tested for acetylation of Compound 1, AI3:16(4,4,8). An SlASAT3 outparalog (Koonin, 2005) SMEL4.1_12g015780, ACYLSUGAR ACYLTRANSFERASE3-LIKE1 (SmASAT3-L1) was the only enzyme to exhibit detectable forward activity acetylating Compound 1, AI3:16(4,4,8), to Compound 2, AI4:18(2,4,4,8) (Figure 6C). As shown in Figure S13, LC-MS analysis of the SmASAT3-L1 *in vitro* assay product revealed that it had identical molecular masses and elution times to plant-derived AI4:18(2,i4,i4,i8) (Compound 1) providing evidence that SmASAT3-L1 acetylates AI3:16(i4,i4,i8). Characterization of reverse activities of the seven BAHD acyltransferases (Leong et al., 2020; Lou et al., 2021), in which an acyl chain can be removed from an acylsugar by incubating it with the enzyme and free Coenzyme A, confirmed that SmASAT3-L1 was the only tested enzyme that removed an acetyl chain from AI4:18(2,4,4,8) (Figure 6D and Figure S13). Taken together, these results indicate that trichome-expressed *SmASAT3-L1* encodes an acylinositol acetyltransferase responsible for AI4:18(2,4,4,8) biosynthesis.

We tested the hypothesis that *S. aethiopicum* trichome extracts lack detectable acetylated AI4:18(2,4,4,8) due to a defect in *SmASAT3-L1* ortholog expression. Indeed, reverse transcription (RT)-PCR comparing isolated trichome and trichomeless shaved hypocotyl cDNA revealed that *SaASAT3-L1* (GenBank ID: OQ547782) transcript was undetectable in *S. aethiopicum* trichomes (Figure S14). This result implicates the defect in ASAT3-L1 enzyme gene expression as responsible for lack of AI4:18 in *S. aethiopicum*. The interspecific variation in *ASAT3-L1* expression is reminiscent of the ASAT4 expression differences among accessions of the wild tomato *Solanum habrochaites*. In that case, a subgroup of *S. habrochaites* accessions possessed a functional ASAT4 copy, but its low levels of gene expression correlated with reduced accumulation of acetylated acylsugars in these ‘subgroup E’ accessions (Kim et al., 2012).

### Natural variation potentiates novel activities and uses

Until recently, it appeared that acylsucroses dominate acylsugar profiles of Solanaceae species with a small number of documented examples of acylglucose- and acylinositol-producing *Solanum* species (Fiesel et al., 2022; Fobes et al., 1985; Herrera-Salgado et al., 2005; Hurney, 2018; Kerwin et al., 2024; King and Calhoun, 1988; Leong et al., 2020; Liu et al., 2017; Lou et al., 2021; Moghe et al., 2017; Schenck et al., 2022). Our chemotaxonomic survey of nearly three dozen *Solanum* species from the sparsely sampled Clade II, DulMo, and VANAns clades revealed widespread occurrence of glucose-, inositol-, and non-sucrose disaccharide-based acylsugars decorated with unusual acyl chains, including medium-length (C12-C16) hydroxyacyl chains, glycosylated hydroxyacyl chains, and unsaturated chains. Adding to interspecific variation we observe here, the recent identification of acylsugars in cultivated tomato root exudates (Korenblum et al., 2020) and root tissues (Kerwin et al., 2024) supports the value of metabolite screening in additional tissue types as well as species.

The characteristics of ASAT3-L1, the enzyme absent from the *S. aethiopicum* trichome transcriptome reveals the dynamic nature of plant specialized metabolism. First, SmASAT3-L1 is unique among characterized ASATs in its acyl acceptor specificity, acetylating an acylinositol glycoside rather than an acylated sucrose, glucose, or *myo*-inositol. Second, acetyl-CoA donor activity was previously only described for a subset of Clade III BAHDs, including ASAT4 and ASAT5 (Figure 7). This theme was also observed in trichomes of two other *Solanum* plants, more closely related to the species screened in this study: the Clade II *S. quitoense* triacylinositol acetyl transferase SqTAIAT (Leong et al., 2020) and DulMo *S. nigrum* acylglucose acetyl transferase SnAGAT1 (Lou et al., 2021). Both enzymes reside outside of the ASAT3 clade, and thus are phylogenetically distinct from SmASAT3-L1. Intriguingly, the characteristics of trichome-expressed SmASAT3-L1 and *S. quitoense* SqASAT1H, (Leong et al., 2022, 2020), as well as cultivated tomato root-expressed SlASAT1-L (Kerwin et al., 2024) also support the hypothesis that acylinositols are synthesized through one or more pathway(s) distinct from that of acylsucroses.

**Figure 7.**
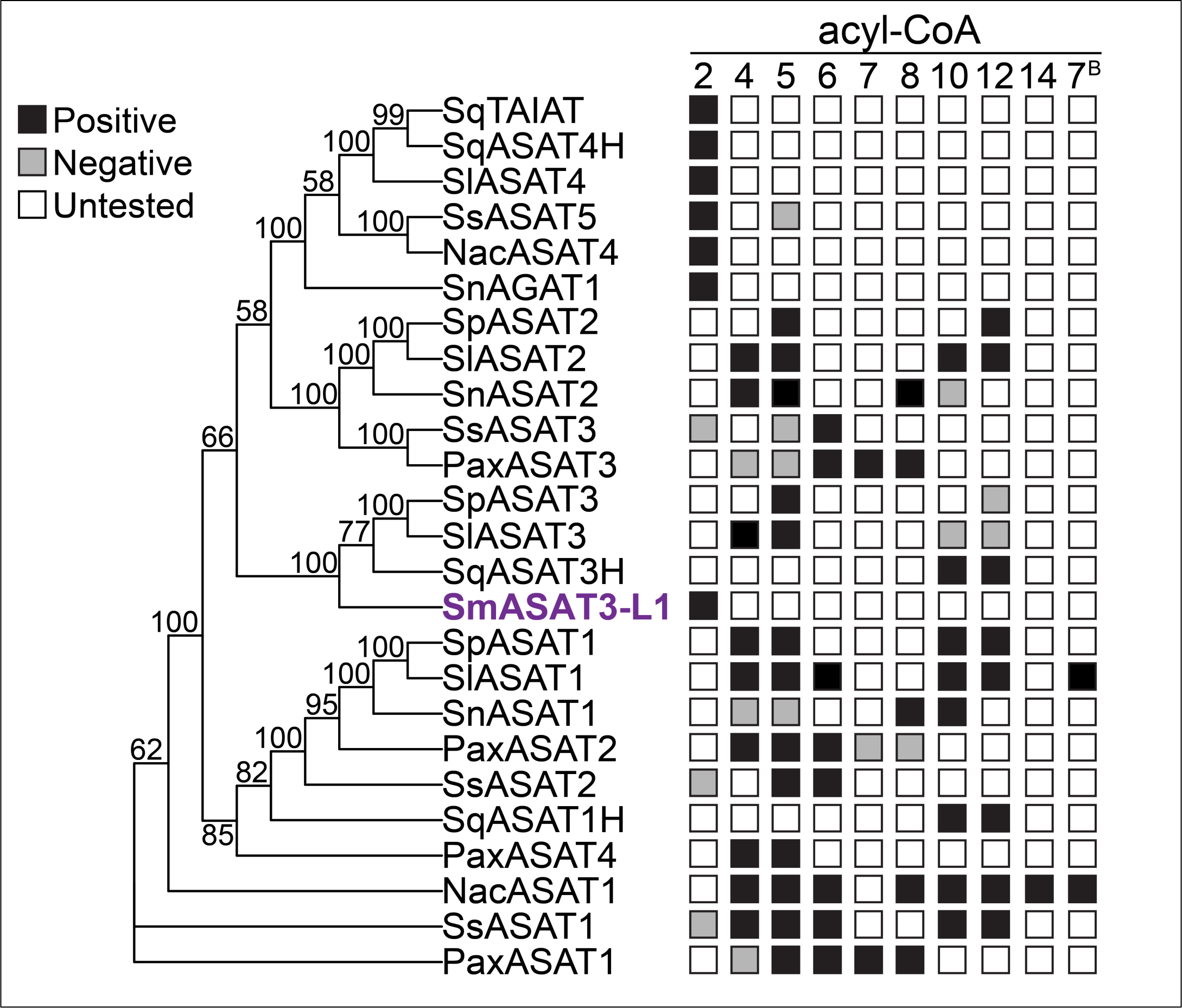
Characterized *in vitro* acyl-CoA usage by ASATs. The maximum likelihood phylogeny was inferred from amino acid sequences of characterized ASAT enzymes using a time-reversible algorithm specifying a plant-specific empirical substitution matrix, invariant sites, and four rate categories (Q.plant+I+G4) in IQ-TREE v2.1.3. Branch support was calculated from 100,000 ultrafast bootstrap replicates. Other than SmASAT3-L1, enzyme activities were described previously (Fan et al., 2017, 2015; Leong et al., 2022, 2020; Lou et al., 2021; Moghe et al., 2017; Nadakuduti et al., 2017; Schenck et al., 2022; Schilmiller et al., 2015, 2012). Acyl-CoA lengths are represented by their number of carbons. Acyl chain 7^B^ corresponds to a benzoyl-CoA. Reported positive activities are indicated by black squares; reported negative activities are indicated by gray squares; untested activities are indicated by white squares.

The remarkably varied acylsugar structures reported here likely have distinct bioactivities. For example, hydrophobic acylsugars with longer acyl chains and few free hydroxyl groups may disrupt membranes, reminiscent of less polar triterpene saponin variants (Augustin et al., 2011; Baumann et al., 2000). Variation in acyl chain linkage chemistries also likely influence acylsugar mode of action. For example, ester linked short chain acylsucroses are digested by *Manduca sexta* (hawkmoth) larvae and the volatile organic acids released into the environment, attracting ant predators (Weinhold and Baldwin, 2011). In contrast, longer chain hydroxylated and ether-linked glycohydroxyacyl chains observed in the study are likely to have quite different metabolic fates, perhaps persisting in the digestive systems of herbivores, and alternative metabolic fates in microbes. Recent work showed that free medium-chain 3-hydroxy fatty acids elicit a strong immune response in *Arabidopsis* and treating plants with 3-OH-C10 fatty acid increased resistance to *Pseudomonas syringae* (Kutschera et al., 2019). If hydroxylated acyl chains are released from acylsugars upon tissue damage, this could trigger a similar plant immune response, thereby conferring resistance to pathogen infection. Future work is needed to determine the function of hydroxylated acyl chain-containing acylsugars. Hydroxyacyl chains also present promising opportunities to expand upon natural acylsugar diversity through synthetic biology and/or synthetic chemistry. The hydroxyl acts as a reactive chemical handle, which was exploited in a subset of *Solanum* species trichomes to create glycohydroxyacylhexoses, and synthetic biology approaches could be used to add unusual sugars, acyl chains, and aromatic groups through promiscuous enzymes or synthetic chemistry. This handle can be exploited with synthetic chemistry using these chains as new feedstocks for creating polymers and may be useful to create synthetic acylsugars for use as food-grade emulsifiers. These structural changes would likely impact biological activities and may be useful for developing more pest resistant plants.

## Methods

### Plant material and growth conditions

*Solanum* spp. seeds were obtained from the sources described in Table S1. Seeds were treated with 10% (v/v) bleach for 10 min while being rocked at 24 rpm with a GyroMini nutating mixer (Labnet, Edison, NJ, USA), and then rinsed 5-6 times with distilled water. Unless otherwise noted, seeds were germinated on Whatman filter paper (MilliporeSigma, Burlington, MA, USA) at 28°C and in the dark. Germinated seedlings were transferred to peat pots (Jiffy, Zwijndrecht, Netherlands), and grown at 25°C, 16/8-h day/night light cycle, and ∼70 μmol m^-2^s^-1^ photosynthetic photon flux density with cool white fluorescent bulbs. Mature plants were grown in controlled environment growth chambers or in a greenhouse. The growth chamber conditions consisted of 25°/18°C day/night temperatures, 16/8-h day/night light cycle, and ∼100 μmol m^-2^s^-1^ photosynthetic photon flux density under LED bulbs. The greenhouse conditions consisted of a 16/8-h day/night light cycle achieved with supplemental sodium iodide lighting. Plants were fertilized weekly with 0.5X Hoagland’s solution.

### Surface metabolite extractions

Surface metabolites were extracted as described previously (Leong et al., 2019; Lou and Leong, 2019). We collected tissue primarily from seedlings as many of these *Solanum* species do not produce glandular trichomes on mature tissues. The only exceptions were *S. mammosum* and *S. sisymbriifolium* which produce glandular trichomes throughout their reproductive stages and from which we sampled mature leaf tissue. Briefly, 0.1-1 g of leaf, stem, hypocotyl, or cotyledon tissue was collected into a 1.5 mL screw-cap tube (Dot Scientific, Inc., Burton, MI, USA), 1 mL extraction solvent (3:3:2 acetonitrile:isopropanol:water, 0.1% formic acid, 10 µM propyl 4-hydroxybenzoate (internal standard)) was added and the sample was rocked at 24 rpm by a GyroMini nutating mixer for two min. After extraction, the supernatant was transferred into a glass 2 mL autosampler vial (Restek, Bellefonte, PA, USA) and sealed with a 9 mm cap containing a PTFE/silicone septum (J.G. Finneran, Vineland, NJ, USA). For fruit surface metabolite extractions, we modified the protocol by placing 1-2 fruit into a 15 mL polypropylene conical tube (Corning Inc., Corning, NY, USA) containing 5 mL extraction solvent then proceeded to the nutation step above.

### Bulk eggplant acylsugar extraction and purification for NMR analysis

Surface metabolites were bulk extracted from approximately 1000 *S. melongena* PI 555598 seedlings. Seeds were treated with 10% (v/v) bleach (Clorox, Oakland, CA, USA) for 10 min while being gently rocked at 24 rpm with a GyroMini nutating mixer and subsequently rinsed 5-6 times with distilled water. Seeds were sown in moist SUREMIX soil (Michigan Grower Products, Galesburg, MI, USA) in Perma-Next plant trays, 22 x 11 x 2.5 inches (Growers Supply Company, Dexter, MI, USA), covered with a humidity dome, 22 x 11 x 3 inches (Growers Supply Company), then immediately transferred to the growth chamber. Seedlings were harvested when 2-3 true leaves were observed, approximately one week after their germination, by cutting them at the base and placing them into two 2L beakers each containing 1L 100% acetonitrile. Surface metabolites were extracted with gentle agitation with a metal spatula for five min at room temperature. Plant material and sediment was removed by vacuum filtration through a Büchner funnel lined with Whatman filter paper (MilliporeSigma), Solvent was removed *in vacuo* by rotary evaporation and dried residue was dissolved in 20 mL of acetonitrile with 0.1% formic acid. Solvent was removed again using a vacuum centrifuge and the dried residue was dissolved in 1 mL of acetonitrile:water:formic acid (80:20:0.001). The semi-preparative LC method is described in detail in Table S28 and is described in brief here. *S. melongena* acylsugars were separated with a Waters 2795 HPLC (Waters Corporation, Milford, MA, USA) equipped with an Acclaim 120 C18 HPLC column (4.6 x 150 mm, 5 μm; Thermo Fisher Scientific, Waltham, MA, USA). Solvent A was water with 0.1% formic acid and Solvent B was acetonitrile. The reverse-phase linear gradient was as follows: 5% B at 0 min, 60% B at 2 min, 80% B at 40 min, 100% B at 42 min, 5% B at 42.01 min, held at 5% B until 44 min. Flow rate was 1.5 mL/min, injection volume was 100 µL, and the column temperature was 40L. Fractions were collected automatically at 0.25 min intervals by a 2211 Superrac fraction collector (LKB Bromma, Stockholm, Sweden) and assessed for acylsugar presence and purity by LC-MS analysis, as described below. Column fractions were collected in the same tubes for each method run which worked to pool the fractions between each run.

### LC-MS acylsugar analysis

Acylsugars were analyzed by LC-MS with each method described below and in Tables S29-33. For each LC method, the mobile phase consisted of aqueous 10 mM ammonium formate, adjusted to pH 2.8 with formic acid (Solvent A) and 100% acetonitrile (Solvent B). The flow rate was maintained at 0.3 mL/min.

Acylsugar extracts were analyzed with a 22 min LC gradient using a Waters Acquity UPLC coupled to a Waters Xevo G2-XS QToF mass spectrometer (Waters Corporation, Milford, MA) equipped with electrospray ionization (ESI) operating in positive (ESI^+^) and negative (ESI^-^) modes. Ten microliter acylsugar extracts were separated on an Acquity UPLC BEH C18 column (10 cm x 2.1 mm, 130 L, 1.7 µm; Waters), kept at 40L, using a binary solvent gradient. The 22 min linear gradient was as follows: 5% B at 0 min, 60% B at 2 min, 100% B at 16 min, held at 100% B until 20 min, 5% B at 20.01 min, held at 5% B until 22 min. Acylsugar extracts were analyzed with both ESI^-^, and ESI^+^. For ESI^-^, the following parameters were used: capillary voltage, 2 kV; sampling cone voltage, 35 V; source temperature, 100°C; desolvation temperature 350°C; cone gas flow, 50 L/Hr; desolvation gas flow, 600 L/Hr. For ESI^+^, the following parameters were used: capillary voltage, 3 kV; sampling cone voltage, 35 V; source temperature, 100°C; desolvation temperature 300°C; cone gas flow, 50 L/Hr; desolvation gas flow, 600 L/Hr. Acylsugars were fragmented in either MS^E^ or data-dependent acquisition (DDA) MS/MS modes as described previously (Lou et al., 2021; Lybrand et al., 2020). DDA survey and MS/MS functions acquired over *m/z* 50 to 1500 with scan times of 0.2 s. Ions selected for MS/MS were fragmented with a ramped collision energy where voltage varies based on ion mass. Collision energies followed a ramp with the voltage changing linearly for ions between the low mass setting (*m/z* 50, 15 to 30 V) and high mass setting (*m/z* 1500, 30 to 60 V). To increase mass accuracy, lock mass correction was performed during data collection, with leucine enkephalin as the reference.

Semi-preparative LC fractions were analyzed with direct infusion and a 14-min LC gradient using a LC-20ADvp ternary pump (Shimadzu, Kyoto, Japan) coupled to a Waters Xevo G2-XS QToF mass spectrometer (Waters Corporation). The direct infusion method quickly screened fractions for acylsugar presence while the 14-min LC method tested fraction purity. Ten microliters of acylsugar fractions were injected into an Ascentis Express C18 HPLC column (10 cm x 2.1 mm, 2.7 µm; Supelco, Bellefonte, PA, USA), kept at 40L. The 14-min gradient was as follows: 5% B at 0 min, 60% B at 2 min, 100% B at 10 min held until 12 min, 5% B at 12.01 min, and held at 5% until 14 min. Fractions were analyzed under ESI^+^ with the following parameters: capillary voltage, 3 kV; sampling cone voltage, 40 V; source temperature, 100°C; desolvation temperature 350°C; cone gas flow, 20 L/hr; desolvation gas flow, 500 L/hr. For direct infusion analysis, ions were acquired from *m/z* 50 to 1500 with a scan time of 0.1 s and one acquisition function with 0 V collision potential. For 14-min LC analysis, ions were acquired from *m/z* 50 to 1200 with a scan time of 0.1 s and three acquisition functions with different collision potentials (0, 25, 60 V). Lock mass calibration referenced to the leucine enkephalin [M+H]^+^ ion was applied during data acquisition.

Enzyme assays products were analyzed with a 7-min gradient using Waters Acquity UPLC coupled to a Waters Xevo G2-XS QToF mass spectrometer (Waters Corporation, Milford, MA) equipped with electrospray ionization (ESI) operating in negative (ESI^-^) mode. Reaction products were separated with an Ascentis Express C18 HPLC column (10 cm x 2.1 mm, 2.7 µm; Supelco), kept at 40L, using a binary solvent gradient. The 7 min linear gradient was as follows: 5% B at 0 min, 60% B at 1 min, 100% B at 5 min, held at 100% B until 6 min, 5% B at 6.01 min, held at 5% B until 7 min. The ESI^-^ parameters described for acylsugar extract analysis were used. Ions were acquired from *m/z* 50 to 1200 with a scan time of 0.1 s and three acquisition functions with different collision potentials (0, 25, 60 V). Lock mass calibration referenced to the leucine enkephalin [M-H]^-^ ion was applied during data acquisition.

For coelution analysis between enzymatically- and plant-produced AI3:16 and AI4:18, a 24-min linear gradient was used with an Acquity UPLC BEH C18 column (10 cm x 2.1 mm, 130 L, 1.7 µm; Waters), kept at 40LC, on the same instrument used for enzyme assay analysis. The binary solvent, linear gradient was as follows: 5% B at 0 min, 60% B at 18 min, 100% B at 18.01 min, held at 100% B until 22 min, 5% B at 22.01 min, held at 5% B until 24 min. ESI^-^ parameters and MS method from the above enzyme assay analysis were used.

Saponified acylsugars were analyzed using a Waters Acquity UPLC coupled to a Waters Xevo G2-XS QToF mass spectrometer (Waters Corporation, Milford, MA). Ten microliters of saponified acylsugars were injected into either an Acquity UPLC BEH C18 column (10 cm x 2.1 mm, 130 L, 1.7 µm; Waters) or an Acquity UPLC BEH Amide column (10 cm x 2.1 mm, 130 L, 1.7 µm; Waters), kept at 40L. Samples were analyzed on the C18 column with the 22-min method described above. To detect free sugars resulting from saponification, samples were analyzed on the BEH amide column with a 9 min method which was as follows: 95% B at 0 min held until 1 min, 60% B at 6 min, 5% B at 7 min, 95% B at 7.01 min, held at 95% B until 9 min. Both methods used the ESI-parameters and MS acquisition parameters described for enzyme assays.

### Acylsugar annotation

We inferred acylsugar structures utilizing negative and positive mode MS and MS/MS collision-induced dissociation (CID) as previously described (Ghosh et al., 2014; Hurney, 2018; Leong et al., 2019; Lou et al., 2021; Lybrand et al., 2020). We developed modified stepwise confidence assessments based on those stepwise criteria described by Lou and coworkers in their *Supplementary Text* ‘Acylsugar annotation’ section (Lou et al., 2021). Our criteria A-F were adapted directly from Lou et al: (A) Exact mass; (B) Presence of co-eluting sugar core fragments; (C) Presence of co-eluting fatty acid carboxylate fragments in high CID potential (30V); (D) Presence of co-eluting fragments that correspond to neutral losses of acyl chains in positive ion mode for acylinositols and negative ion mode for acyldissacharides and acylglucoses; (E.) Confirmed by MS/MS; (F) Consistency across samples. Putative acylsugars only meeting criterion A were annotated as low confidence, while those meeting all four criteria described in A-D were annotated as medium confidence. We added two additional criteria:(G) **NMR structural determination.** Acylsugars from *S. melongena* were further characterized by NMR experiments in which sugar core relative stereochemistry, acyl chain positions, and acyl chain branching were resolved. (H) **Coelution with NMR characterized compounds.** Acylsugars with the same acyl chains and sugar core size were analyzed by coelution with *S. melongena* acylsugars. If acylsugars coeluted with an NMR characterized peak and shared the same acyl chains and fragmentation patterns, they met this criterion and were annotated with high confidence. This criterion was employed for acylsugars from species other than *S. melongena*.

Acylsugars meeting all criteria in A-F and either G or H were annotated with highest confidence. Confidence levels for each reported acylsugar are specified in their species-specific annotation table (Tables S2-27).

### Acylsugar metabolomic analysis

To collect accurate formate adduct masses and relative abundances from putative acylsugars, negative mode LC-MS raw data from metabolite extracts were analyzed with Progenesis QI software v3.0 (Waters, Milford, MA, USA) using retention time alignment, peak detection, adduct grouping, and deconvolution. Lock mass correction was performed during data collection. We used the following analysis parameters: peak picking sensitivity, default; retention time range, 0-22 min; adducts detected, M+FA-H, M+Cl, M-H, 2M-H, and 2M+FA-H. Average percent peak abundance was calculated for each putative acylsugar by dividing its raw abundance by the sum of the total acylsugar raw abundance and then averaging that value across all samples for a species.

### NMR analysis of acylsugars

Purified acylsugars were dissolved in acetonitrile-*d*_3_ (99.96 atom % D; MilliporeSigma, Burlington, MA, USA) and transferred to Kontes NMR tubes (MilliporeSigma, Burlington, MA, USA). Samples were analyzed at the Michigan State University Max T. Rogers NMR Core with a Varian Inova 600 MHz spectrometer (Agilent, Santa Clara, CA, USA) equipped with a Nalorac 5 mm PFG switchable probe, a DirectDrive2 500 MHz spectrometer (Agilent, Santa Clara, CA, USA) equipped with a OneNMR probe, Bruker Avance NEO 800 MHz spectrometer (Bruker, Billerica, MA, USA) equipped with a 5 mm helium cryogenic HCN probe, or a Bruker Avance NEO 600 MHz spectrometer equipped with a 5 mm nitrogen cryogenic HCN Prodigy probe. All acylsugars were analyzed with a series of 1D (^1^H, and ^13^C) and 2D (gCOSY, TOCSY, gHSQC, gHMBC, gH2BC, *J*-resolved) experiments. Resulting spectra were referenced to acetonitrile-*d*_3_ (^O^H = 1.94 and ^O^C = 118.70 ppm). NMR spectra were processed and analyzed with MestReNova 12.0 software (MestreLab, A Coruña, Spain). For full NMR metadata, see Supplementary Datafile 1.

### Sugar core composition analysis

*S. melongena* PI 555598 acylsugar extracts were dried down *in vacuo* with centrifugation (Savant, ThermoFisher Scientific). Dried acylsugar extracts were dissolved in 1 mL of methanol. One mL of 3 M ammonium hydroxide was added, and the solution was mixed vigorously. This initial saponification reaction proceeded at room temperature for 48 hours. Solvent was removed by vacuum centrifugation. Saponified sugar cores were reduced and derivatized with acetate groups as previously described (Sassaki et al., 2008). Sugars were dissolved in 50 µL of 1 M ammonium hydroxide for 15 min at room temperature. Addition of 50 µL 20 mg/mL NaBH_4_ and incubation at 100°C for 10 min converted aldoses and ketoses to polyols. Excess sodium borohydride was quenched with 100 µL 1M trifluoroacetic acid. Two volumes of methanol were added, and the sample was dried down by a stream of nitrogen gas. Dried down reaction products were redissolved in 200 µL of methanolic 0.5 M HCl and heated at 100°C for 15 min in a closed ½ dram vial, 12 x 35 mm (Kimble Chase, Vineland, NJ, USA). Solvent was removed by a stream of nitrogen gas. To acetylate the sugar cores, residue was dissolved in 200 µL of pyridine:acetic anhydride (1:1) and incubated at 100°C for 30 min. Acetylated polyols were then dried down *in vacuo* in a vacuum centrifuge and redissolved in 100 µL hexane for GC-MS analysis.

To determine the composition of disaccharide sugar cores, saponified sugars were hydrolyzed prior to derivatization. Dried down sugars were redissolved in 98 µL water in a ½ dram vial, 12 x 35 mm (Kimble Chase). After addition of 102 µL of 88% formic acid, the vial was sealed and heated at 100L for 15 hours in a heat block. Hydrolyzed sugars were dried down *in vacuo* with a vacuum centrifuge and derivatized to alditol acetates following the method described above.

### Hydroxyl acyl chain stereochemistry analysis

We derivatized hydroxyacyl chains and extracted the derivatives as previously described (Jenske and Vetter, 2007). The commercial standards (3*R*)-OH-tetradecanoate (Cayman Chemical, Ann Arbor, MI, USA) and (3*R*/*S*)-OH-tetradecanoate (Cayman Chemical, Ann Arbor, MI, USA) were first derivatized to their respective ethyl ester derivatives. 600 µg of each fatty acid was dissolved in 0.5 mL of 0.5 M ethanolic KOH and incubated at 80°C for 5 min. After the solution was cooled on ice, 1 mL of ethanolic BF_3_ was added, and reactions were incubated at 80°C for five min. Two-phase partitioning with hexane and saturated sodium chloride yielded ethyl ester derivatives in the hexane partition.

The 3-OH-nC14 acyl chain from I3:22(iC4,iC4,3-OH-nC14) was derivatized to an ethyl ester following a previously published method (Fan et al., 2020; Ning et al., 2015). Half of the purified I3:22(iC4,iC4,3-OH-nC14) was dissolved in acetonitrile with 0.1% formic acid and transferred to a 1.5 mL microcentrifuge tube. The purified acylsugar was dried down *in vacuo* with vacuum centrifugation. The resulting dry, purified I3:22(iC4,iC4,3-OH-nC14) was dissolved in 300 µL of 21% (w/w) sodium ethoxide in ethanol (MilliporeSigma, Burlington, MA, USA). The reaction proceeded at room temperature for 30 min with constant rocking at 24 rpm by a GyroMini nutating mixer (Labnet, Edison, NJ, USA) and occasional vortexing. 400 µL hexane with 55 ng/µL tetradecane (MilliporeSigma, Burlington, MA, USA) as an internal standard was then added followed by vigorous vortexing. 500 µL of aqueous saturated sodium chloride was added to the hexane-ethanol mixture, and vortexed. The hexane layer was pipetted to another tube and two more two-phase partitions with saturated sodium chloride were completed. The final hexane layer containing the acyl chain ethyl esters was extracted and dried down by a stream of nitrogen gas.

The resulting fatty acid ethyl esters from the commercial standards and purified compound were derivatized with (*R*)-(-)-L-methoxy-L-trifluoromethylphenylacetyl chloride ((*R*)-(-)-MTPA-Cl; MilliporeSigma, Burlington, MA, USA) following a previously published procedure (Jenske and Vetter, 2007). Dried fatty acid ethyl esters were redissolved in 400 µL pyridine and 15 µL of (*R*)-(-)-MPTA-Cl. The reaction proceeded at room temperature for two hours, after which, 5 mL of water and tert-butyl methyl ether (TBME) were added along with solid K_2_CO_3_ (one spatula tip). The TBME phase was collected after three successive phase separations with 5 mL of water. TBME was evaporated to 1 mL and subjected to GC-MS analysis.

### GC-MS analysis

All GC-MS analyses employed an Agilent 5890 GC and an Agilent 5975 single quadrupole MS equipped with a FactorFour VF-5ms column (30 m x 0.25 mm, 0.25 um; Agilent) and 10 m EZ-Guard column (Agilent, Santa Clara, CA, USA). Helium was used as the carrier gas with a constant flow rate of 1 mL/min. Electron energy was set at 70 eV. MS source and quadrupole were maintained at 230°C and 150°C, respectively. Parameters specific to each analysis are listed below.

For analysis of sugar core alditol acetate derivatives, we followed previously published GC-MS parameters (Sassaki et al., 2008). Inlet temperature was maintained at 275°C. GC oven temperature was held at 60°C for one min and then was ramped at a rate of 40°C/min to 180°C. Oven temperature was then ramped to 240°C at a rate of 5°C/min and held for three min. Total run time was 19 min. The MS detector transfer line was maintained at 280°C. Split ratio was 10:1 with a split flow rate of 10 mL/min. A three min solvent delay was used. MS data was collected in full scan mode, *m/z* 50-600. Analysis parameters are also described in Table S34.

For analysis of MPTA fatty acid ethyl ester derivatives, we followed previously published GC-MS parameters (Jenske and Vetter, 2007). GC oven temperature was held at 60°C for 1.5 min and then was ramped at a rate of 40°C/min to 180°C. 180°C was held for two min after which the oven temperature was ramped to 230°C at a rate of 2°C/min. After 230°C was held for nine min, the oven temperature was ramped to 300°C at 10°C/min and held for 7.5 min. Total run time was 55 min. A 5-min solvent delay was applied. The MS detector transfer line was maintained at 280°C. Selective ion monitoring detected the ions *m*/*z* 189, 209, and 255 with a dwell time of 50 ms. Analysis parameters are also described in Table S35.

### Sample collection and transcriptome sequencing

We sequenced 18 transcriptomes: six biological replicates each of trichomes isolated from hypocotyls (trichomes), hypocotyls stripped of their trichomes (stripped hypocotyls), and whole roots (Supplemental data file 1). We sampled trichomes and stripped hypocotyls from 7- day-old *S. melongena* 67/3 seedlings following methods developed for root hair cell isolation (Bucher et al., 1997) with modifications. We grew lawns of eggplant seedlings in soil flats as described above for bulk leaf surface metabolite extraction. At 7 days post germination, we removed roots and cotyledons from seedlings, then transferred hypocotyls to liquid nitrogen in a plastic 2 L Dewar flask. Frozen hypocotyls were gently stirred with a glass rod for 20 minutes to physically shear trichomes from hypocotyls. After confirming under a dissecting scope that a sample of three hypocotyls had been stripped bare, we filtered trichomes into a 2 L glass beaker by slowly pouring the contents of the Dewar flask through a 500 μm wire mesh sieve (MilliporeSigma, Burlington, MA, USA). To maximize trichome recovery, stripped hypocotyls were returned to the Dewar, rinsed with liquid nitrogen, and filtered through the 500 μm sieve six more times. Stripped hypocotyls were divided into six pre-weighed 50 mL conical tubes, quickly weighed, then transferred to −80LC for storage. Trichomes were subsequently filtered through a 150 μm sieve (MilliporeSigma, Burlington, MA, USA) into a 500 mL beaker to increase sample purity then transferred to a 50 mL conical tube to allow the excess liquid nitrogen to evaporate. Finally, filtered trichomes were divided into six pre-weighed 2 mL screw-cap tubes (Dot Scientific, Inc., Burton, MI, USA), quickly weighed, then transferred to −80LC for storage.

We extracted total RNA using the RNeasy Plant Mini Kit (Qiagen, Hilden, Germany) following the manufacturer’s directions, measured RNA concentration using a Qubit 1.0 instrument (Thermo Fisher Scientific, Waltham, MA, USA) with the RNA HS assay, and the samples were processed by the Michigan State University Research Technology Support Facility Genomics Core (East Lansing, MI, USA) for library preparation and high-throughput sequencing. The core checked RNA quality using a Bioanalyzer 2100 (Agilent Technologies, Santa Clara, CA, USA), constructed sequencing libraries using an Illumina Stranded mRNA Prep kit (Illumina, Inc. San Diego, CA, USA), and sequenced all 18 libraries on a NovaSeq 6000 (Illumina, Inc. San Diego, CA) using a single S4 flow cell lane, producing 150-base pair paired-end reads (Supplemental datafile 1). The raw fastq files are available to download from the National Center for Biotechnology Information Sequence Read Archive database under BioProject PRJNA935765.

### Transcriptome alignment and differential gene expression analysis

To prepare the 150-bp paired-end sequences for alignment, we employed Trimmomatic (Bolger et al., 2014) to trim adapters and low-quality bases, then filter reads shorter than 75 bp, removing an average of 3.7% of sequences (range: 3.3% - 4.2%). We mapped the resulting 75– 150-bp (average length: 149 bp) paired-end RNAseq reads separately to the eggplant V4.1 and HQ reference genomes using STAR in two-pass mode, which enhances splice junction discovery and mapping sensitivity (Dobin et al., 2013; Dobin and Gingeras, 2015). Using this approach, an average of 79.2% (range: 60.1 - 87.1%) and 82.2% (range: 62.0 - 90.7%) of reads mapped to a unique genomic location in V4.1 and HQ, respectively (Table S36). We filtered the resulting transcriptome alignments according to best practices as defined by the Genome Analysis Toolkit (GATK) (DePristo et al., 2011; Van der Auwera et al., 2013). Briefly, we removed optical and PCR duplicates with MarkDuplicates from the Picard toolkit (http://broadinstitute.github.io/picard), parsed reads into exon segments and removed intron-spanning bases using SplitNCigarReads from GATK (McKenna et al., 2010). Finally, we selected unique alignments by eliminating reads with a mapping quality score below Q60 with the view command from SAMtools (Li et al., 2009). To generate raw read counts, we used the HTSeq command, htseq-count with the nonunique parameter set to all (Anders et al., 2015).

Raw read counts generated by HTSeq-count were used to perform differential gene expression analysis in edgeR (Robinson et al., 2010). To restrict comparisons to expressed genes, only transcripts with at least one read count-per-million (CPM) in at least one sample were retained for further analysis. This filtering step removed 11,665 (33.4%) and 14,544 (39.8%) annotated genes in the V4.1 and HQ eggplant genomes, respectively, yielding 23,251 and 22,024 expressed transcripts for differential gene expression analysis. Next, we normalized transcript abundances across the 18 transcriptomes in our eggplant V4.1 and HQ alignments using the default trimmed mean of M-values (TMM) method in the calcNormFactors function, then performed multidimensional scaling (MDS) with the plotMDS function to compare global gene expression profiles (Figure S15). This showed that our samples cluster tightly by tissue (i.e., trichome, shaved hypocotyl, or root), with no obvious differences between V4.1 and HQ alignments. To test differences in gene expression across tissues, we implemented a generalized linear model (GLM) using a quasi-likelihood (QL) approach: we generated an experimental design matrix specifying the three tissues (i.e., trichomes, trichomeless hypocotyls, and roots) with the model.matrix function, then used the glmQLFit function to fit our data to a QL-GLM model. To identify genes with a log2 fold-change (FC) > 2 between tissues, we used the glmTreat function, which performs threshold hypothesis testing, a rigorous statistical approach that evaluates variance and magnitude to detect expression differences greater than the specified value (e.g., log2 FC > 2), then applies false discovery rate (FDR) *p*-value corrections. Genes were classified as significantly differentially expressed between two tissues if log2 FC > 2 and FDR-corrected *p*-value < 0.05. We calculated absolute transcript abundance for all expressed genes as transcripts per million (TPM) with the calculateTPM function in scateR (McCarthy et al., 2017). Transcriptome expression values are in Tables S37 and S38.

### Phylogenetic analyses

Characterized ASAT enzymes fall into clade III of the BAHD family (Fan et al., 2017, 2015; Leong et al., 2022, 2020; Lou et al., 2021; Moghe et al., 2017; Nadakuduti et al., 2017; Schenck et al., 2022; Schilmiller et al., 2015, 2012). To identify ASAT candidates in eggplant, we searched for sequences containing the Pfam Transferase domain (PF02458) (Finn et al., 2010), associated with all characterized catalytically active BAHD proteins. The PF02458 HMM profile was obtained from the Pfam website (http://pfam.sanger.ac.uk/) and queried against the V4.1 and HQ eggplant proteomes using the hmmsearch tool from HMMER v3.2.1 (hmmer.org), revealing 106 and 108 putative BAHD sequences, respectively. Using MAFFT v7.471 in E-INS-i mode, we built multiple sequence alignments (MSA) of amino acid sequences from four sources: 1) V4.1 or HQ eggplant PF02458 hits, 2) published reference sequences for clades I-VII (Moghe et al., 2023), 3) characterized ASAT sequences from other Solanaceae species, and 4) the SaASAT3 and SaASAT3-L1 candidates from *S. aethiopicum* (Katoh and Standley, 2013). The E-INS-i algorithm implements local alignment with a generalized affine gap cost (Altschul, 1998), which aligns conserved regions (e.g., the BAHD transferase domain) and essentially ignores nonconserved regions. Phylogenetic reconstruction was performed using IQ-TREE v2.1.3 (Minh et al., 2020). The ModelFinder tool was implemented to identify the best maximum likelihood model for estimating evolutionary relationships (Kalyaanamoorthy et al., 2017), leading to selection of Jones-Taylor-Thornton (JTT)+F with seven rate categories and Q.plant+F with seven rate categories for the V4.1 and HQ MSAs, respectively. Phylogenetic trees were inferred by maximum likelihood using the chosen model, and branch support was obtained from 100,000 ultrafast bootstrap iterations (Hoang et al., 2018). The resulting BAHD phylogenies were visualized using the ggtree package in R (Yu et al., 2017). To generate the clade III BAHD heatmap-tree, we used the viewClade function in ggtree to subset the phylogeny and used gheatmap to visualize transcript abundance (log2 TPM) for the eggplant BAHDs.

### BAHD acyltransferase cloning, expression, and purification

Candidate ASATs were cloned into pET28b(+) (MilliporeSigma, Burlington, MA, USA). Open reading frames from genes were either synthesized by Twist Biosciences (South San Francisco, CA, USA) with or without codon-optimization for *E. coli* expression (Table S39) or amplified from genomic DNA or cDNA with primers listed in Table S40. Q5 2X Hotstart master mix (New England Biolabs, Ipswich, MA, USA) was used for cloning PCRs. The source of cloned DNA and whether a gene was codon optimized is described in Table S39. Amplified genes were purified by agarose gel electrophoresis and extraction with the Monarch DNA Gel Extraction Kit (New England Biolabs, Ipswich, MA, USA). Both the synthesized genes and PCR amplified genes were then inserted into a doubly digested BamHI/XhoI pET28b(+) through Gibson assembly using the 2X Gibson Assembly Master Mix (New England Biolabs, Ipswich, MA, USA) according to the manufacturer’s instructions. The constructs using synthesized genes were transformed into BL21(DE3) (MilliporeSigma, Burlington, MA, USA) and constructs with PCR amplified genes were transformed into BL21 Rosetta(DE3) cells (MilliporeSigma, Burlington, MA, USA). Constructs were verified with colony PCR and Sanger sequencing using T7 terminator and promoter primers (Table S40). Sanger sequencing was completed by the Michigan State University Research Technology Support Facility Genomics Core (East Lansing, MI, USA).

Protein expression occurred as previously described (Leong et al., 2022; Lou et al., 2021). Briefly, 50 mL cultures of picked transformation colonies were grown overnight at 37°C, shaking at 225 rpm in Luria-Bertani (LB) media (Neogen, Lansing, MI, USA) supplemented with 1% glucose (w/v). Fifteen mL of the overnight cultures were inoculated into 1 L of fresh LB medium, which was incubated at 37°C shaking at 225 rpm until an OD600 of 0.5 was reached. The cultures were incubated on ice for 25 min, after which, isopropylthio-β-galactoside was added to a final concentration of 50-500 µM. Then, cultures were incubated at 16°C shaking at 180 rpm overnight for 16 hours before cells were harvested by centrifugation at 4,000 rpm for 10 minutes at 4°C.

*S. melongena* BAHDs were purified as previously described (Leong et al., 2020) with the following modifications. The extraction buffer contained 10 mM imidazole, the wash buffer contained 20 mM imidazole, and the elution buffer contained 500 mM imidazole. Protein eluent was concentrated with 30-kD Amicon Ultra centrifugal filter units (MilliporeSigma).

### Enzyme assays

Enzyme assays were conducted in 100 mM sodium phosphate buffer at pH 6. For forward assays, acetyl-CoA (MilliporeSigma, Burlington, MA, USA) was added to a final concentration of 0.1 mM, and for reverse assays free CoA (MilliporeSigma, Burlington, MA, USA) was added to a final concentration of 1 mM. Purified AI3:16 and AI4:18 substrates were dried down using a vacuum centrifuge and redissolved in ethanol:water:formic acid (1:1:0.001). One microliter of the prepared acylsugars were used as acyl acceptors. Six microliters of enzyme were added to a final volume of 60 µL. For negative controls, 6 µL of enzyme that was heat inactivated at 95°C for 10 minutes was substituted in place of untreated enzyme. Assays were incubated at 30°C for 30 minutes after which 120 µL of acetonitrile:isopropanol:formic acid (1:1:0.001) with 1.5 µM telmisartan (MilliporeSigma, Burlington, MA, USA) stop solution was added. Reactions were then spun at 17,000 x *g* for 10 minutes to remove precipitate. Supernatant was placed in autosampler vials and analyzed by LC-MS.

### RT-PCR of *S. aethiopicum* BAHDs

We employed semi-quantitative RT-PCR to test *S. aethiopicum* BAHD expression in glandular trichomes with cDNA (Figure S14) and genomic DNA (gDNA) (Figure S16). Total RNA was isolated with the RNeasy Plant Mini Kit (Qiagen, Hilden, Germany), including an on-column DNase digestion (Qiagen, Hilden, Germany), from shaved hypocotyls from accession *S. aethiopicum* PI 666075 and glandular trichomes from accessions PI 666075 and Grif 14165 isolated as described above for *S. melongena* glandular trichomes. RNA was quantified with a Nanodrop 2000c instrument (Thermo Fisher Scientific, Waltham, MA, USA). We synthesized cDNA using 10 ng of RNA and SuperScript III Reverse Transcriptase (Invitrogen, Waltham, MA, USA). gDNA was isolated with the DNeasy Plant Mini Kit (Qiagen, Hilden, Germany) from young leaf tissue collected from mature *S. aethiopicum* PI 666075 plants. PCR reactions (25 µL) were set up with GoTaq Green Master Mix (Promega, Madison, WI, USA), 200 nM of forward and reverse primers (Table S40), and 1 µL of cDNA or gDNA. PCR was performed under these conditions: 2 min at 95°C followed by 22, 30, and 35 cycles of 30s at 95°C, 30s at 58°C, and 1 min at 72°C.

We identified the putative *S. aethiopicum* orthologs of the *S. melongena* candidate BAHDs by querying *S. aethiopicum* annotated transcripts (Song et al., 2019) with *S. melongena* candidate ASAT DNA sequences using BLASTn. SaASAT3 and SaASAT3-L1 (GenBank accession: OQ547782) were not annotated in the *S. aethiopicum* genome and were identified by querying *S. aethiopicum* scaffolds with BLASTn and then identifying open reading frames with Geneious software v9.1.8 (Dotmatics, Boston, MA, USA). The *S. aethiopicum* ef1α gene was identified by querying a putative *S. melongena* ef1α, SMEL4.1_06g005890, against the *S. aethiopicum* scaffolds.

### *S. melongena* acyl chain composition analysis

*S. melongena* acyl chain composition was assessed with a previously developed esterification and GC-MS methods (Fan et al., 2020; Ning et al., 2015; Schenck et al., 2022). Acyl chain identities were determined through authentic reference standards for nC8, nC10, nC12, and nC14 fatty acid ethyl esters (MilliporeSigma). The QuanLynx function of MassLynx v4.1 (Waters Corporation) integrated peaks from extracted ion chromatograms. Extracted ion chromatograms for the medium acyl chains iC8, nC8, iC10, nC10, iC12, nC12, iC14, and nC14 were generated for the *m/z* of 88 with a mass window of *m/z* of 0.50. Extracted ion chromatograms for the short chains iC4, aiC5, and iC5 were generated for with the *m/*z values of 71, 102, and 101, respectively, with a mass window of *m/z* of 0.50.

### Accession numbers

Raw RNA sequence data can be found in the National Center for Biotechnology Information (NCBI) Sequence Read Archive database under BioProject PRJNA935765. Sequence data for SaASAT3-L1 is available from the NCBI GenBank database under accession number OQ547782.

The NMR data for compounds 1-8 in Figure 1D have been deposited in the Natural Products Magnetic Resonance Database (np-mrd.org) under accession numbers NP0332515, NP0332562, NP0332563, NP0332564, NP0332571, NP0332572, NP0332573, NP0332574, and NP0332575 as documented in Table S41.

The LC-MS data used to annotate compounds in Tables 1, S3-27 have been deposited in the Metabolights database (https://www.ebi.ac.uk/metabolights) under the study ID MTBLS9459.

## Supporting information

Supplementarty NMR data and metadata

## Acknowledgements

We thank Dr. Christopher T. Martine for providing Australian *Solanum* ssp. seeds, Dr. Joyce van Eck for providing *Solanum prinophyllum* seeds, Dr. Guiseppe Leonardo Rotino and the Consiglio per la ricercar in agricoltura e l’analisi dell’economia agraria for providing *S. melongena* 67/3 and 305E40 seeds, and USDA-GRIN for providing *Solanum* ssp. seeds. We acknowledge the Michigan State University RTSF Mass Spectrometry and Metabolomics Core Facilities for LC-MS analysis support. We also acknowledge Dr. Daniel Holmes and Dr. Li Xie at the Michigan State University Max T. Rogers NMR Facility for experimental design and data analysis support. We are grateful to current and past members of the Last lab for feedback on the research and especially Drs. Jaynee Hart and Xingxing Li for comments on the manuscript.

## Author Contributions

PDF: Designed and performed research, analyzed data, wrote the paper

REK: Designed and performed research, analyzed data, wrote the paper

ADJ: Designed research, analyzed data, wrote the paper

RLL: Designed research, reviewed data, wrote the paper

