## Supplementary Figures S1-16 for "Trading acyls and swapping sugars: metabolic innovations in *Solanum* trichomes"

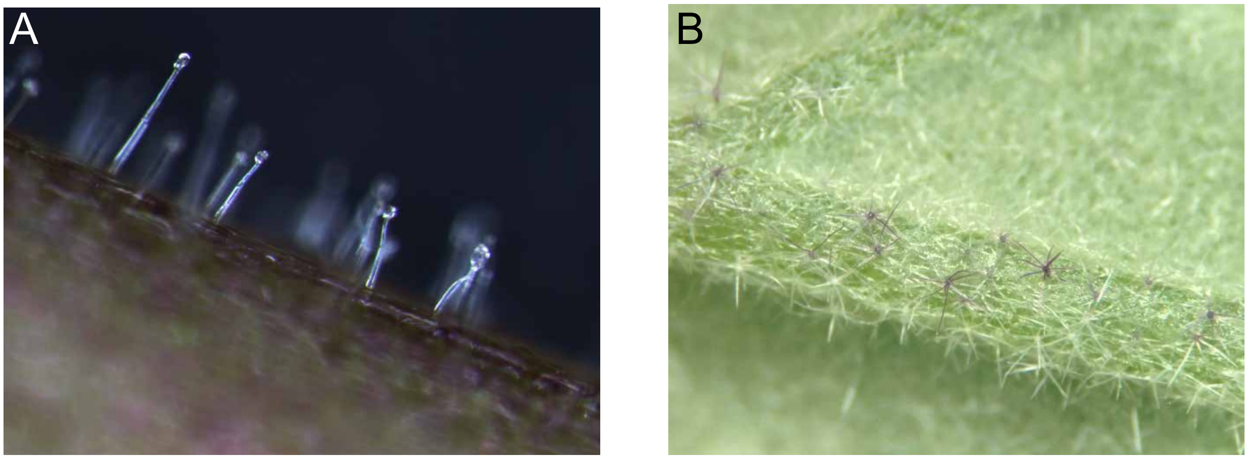


**Figure S1.** ***S. melongena*** **produces single stalked glandular trichomes on seedling above ground tissue but stellate nonglandular trichomes on non-seedling above ground tissue. (A)** Close up photo of a *S. melongena* hypocotyl displaying glandular trichomes with similar morphology to the acylsugar-producing *S. lycopersicum* Type I/IV trichomes (Luckwill, 1943; Schilmiller et al., 2012). **(B)** Close up photo of a *S. melongena* leaf from a reproductive stage plant displaying non-glandular stellate trichomes. Leaf surface metabolite extracts from this tissue do not have detectable acylsugars by LC-MS.

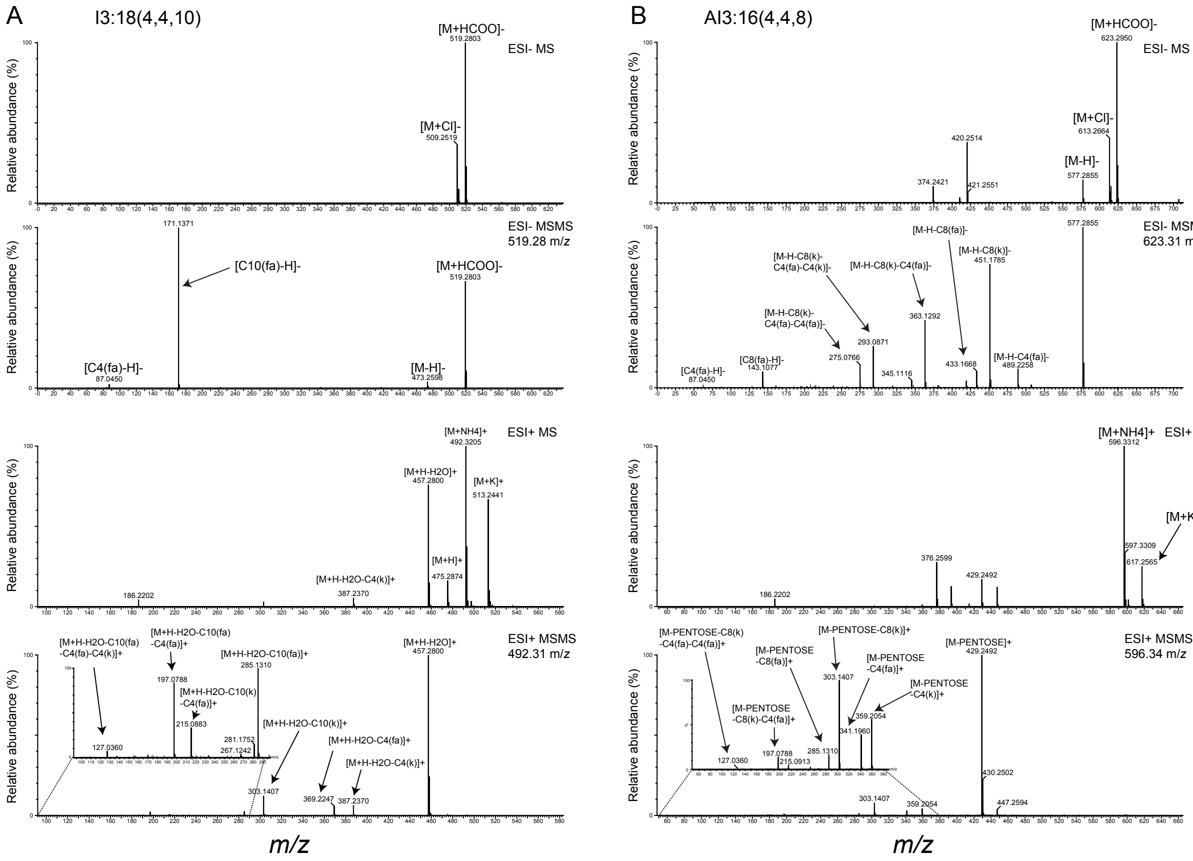


**Figure S2. Annotation of I3:18(4,4,10) and AI3:16(4,4,8) from *S. melongena* 67/3 using negative and positive mode MS and CID MS/MS fragmentation. (A)** I3:18(4,4,10) negative and positive mode MS and MS/MS fragmentation. **(B)** AI3:16(4,4,8) negative and positive MS and MS/MS fragmentation. For each compound, ESI- MS (top panels) display the formate adduct accurate mass which was used to determine chemical formulas. ESI- MS/MS (second from the top panels) exhibit acyl chain carboxylate fragment ions for both compounds and stepwise loss of acyl chains for only AI3:16. ESI+ MS/MS exhibit stepwise losses of acyl chains for both compounds. Positive mode CID of AI3:16 also produces a fragment ion (*m/z* 429.2492) corresponding to the neutral loss of the pentose moiety. This indicates that all acyl chains reside on the hexose ring. The acylsugar LC-MS annotation method is described in the Materials and Methods. fa = fatty acid; k = ketene.

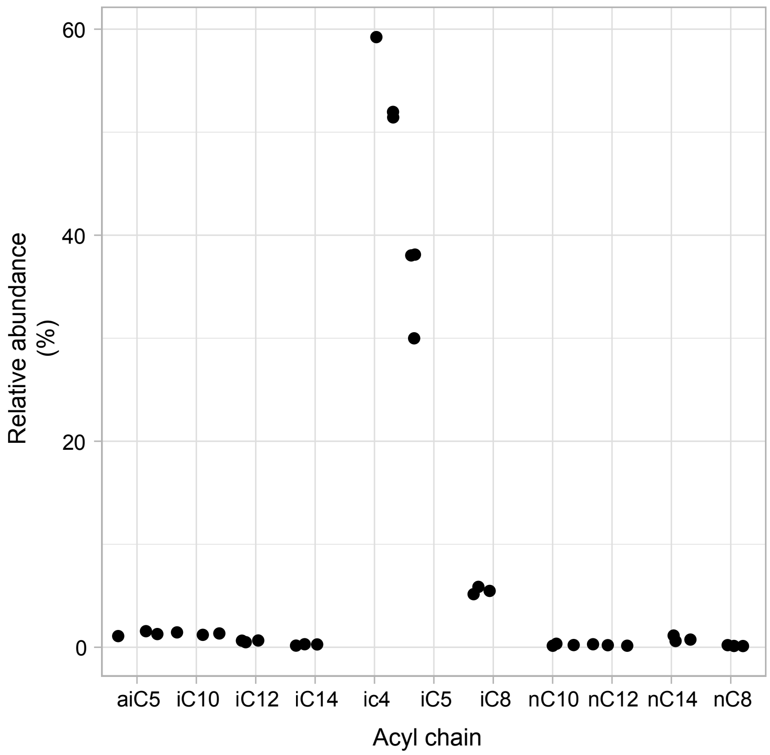


**Figure S3. Acyl chain composition of *S. melongena* acylsugars.** Relative abundance of acyl chains shown between three *S. melongena* PI 555598 leaf surface extracts from different seedlings. Straight acyl chains were identified with authentic reference standards and iso-branched chains were identified with NIST mass spectral library searches. Hydroxylated acyl chains were not included in this analysis and only detected acyl chains are included.

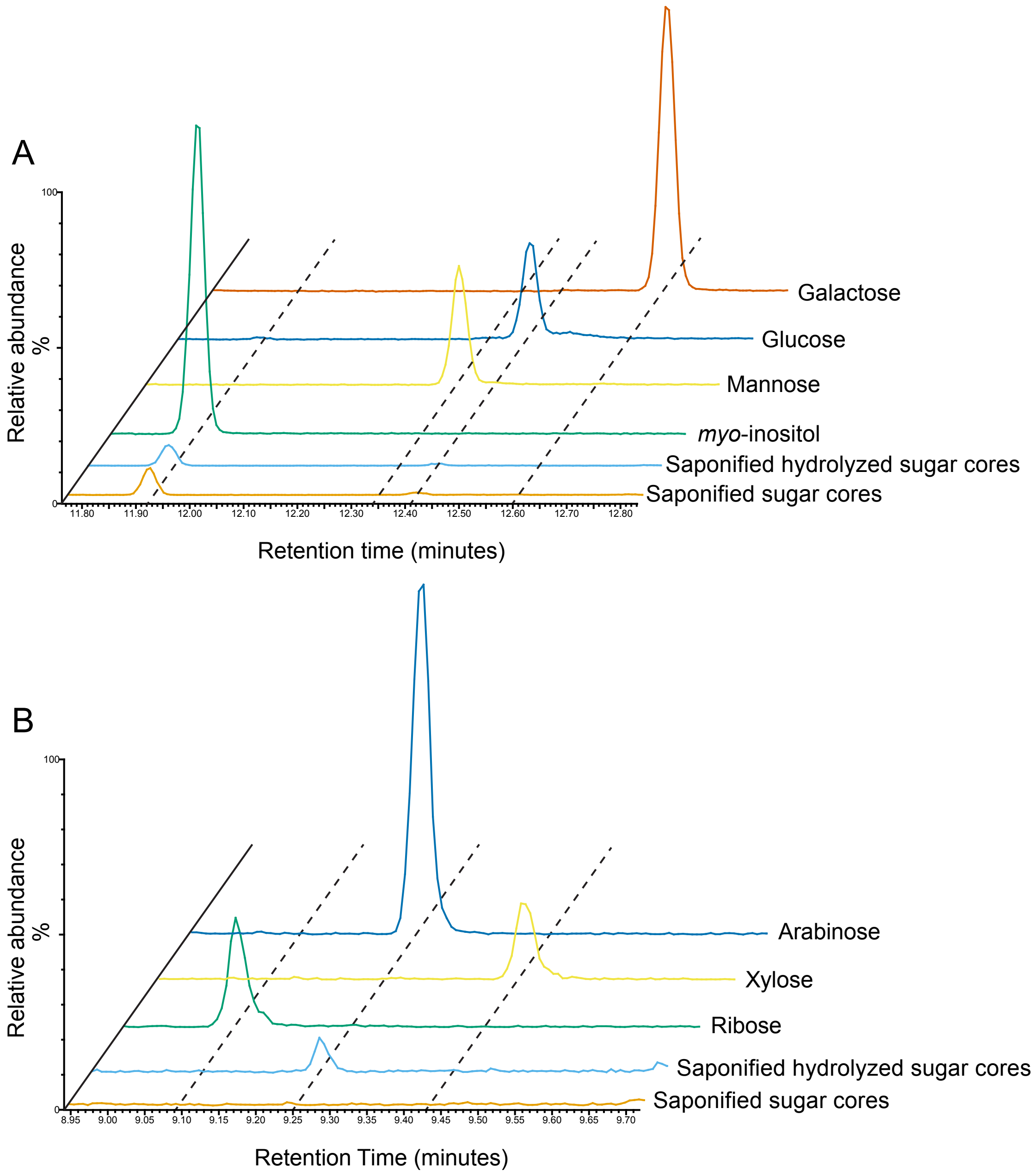


**Figure S4: Identification of *S. melongena* acylsugar core composition through GC-MS analysis of alditol acetate sugar derivatives.** *S. melongena* acylsugars collected from surface extracts were first saponified to remove acyl chains, and then, with or without acid hydrolysis to break the glycosidic linkages, sugar cores were derivatized to alditol acetates. **(A)** Alditol acetate derivatization of saponified *S. melongena* acylsugars yield a peak that comigrated with that of a *myo*-inositol standard. The traces displayed are GC-MS total ion chromatograms (TICs). **(B)** Alditol acetate pentose derivatives of saponified and hydrolyzed acylsugar cores comigrate with that of a derivatized arabinose standard. The traces displayed are GC-MS TICs. Vertical scales for both panels are normalized to the largest signal within the displayed region.

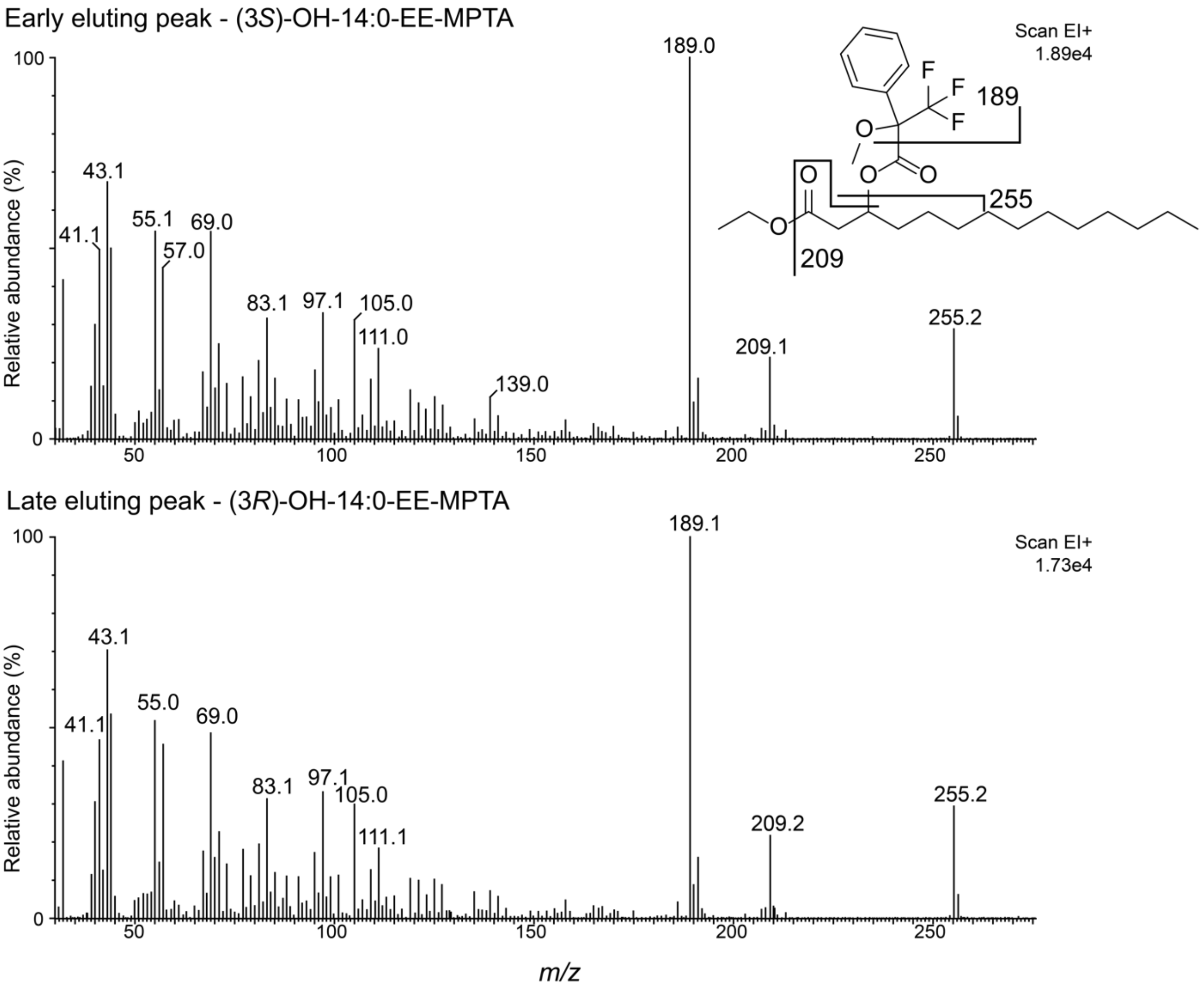


**Figure S5. GC-MS full scan mass spectra of 3-OH-14:0 ethyl ester stereoisomers as their MPTA derivatives.** The mass spectra contain ions at *m/z* 189, 209, and 255 corresponding to the expected fragmentation of the derivatized fatty acid. EE = ethyl ester. Early and late eluting peaks correspond to the peaks displayed in Figure 2.

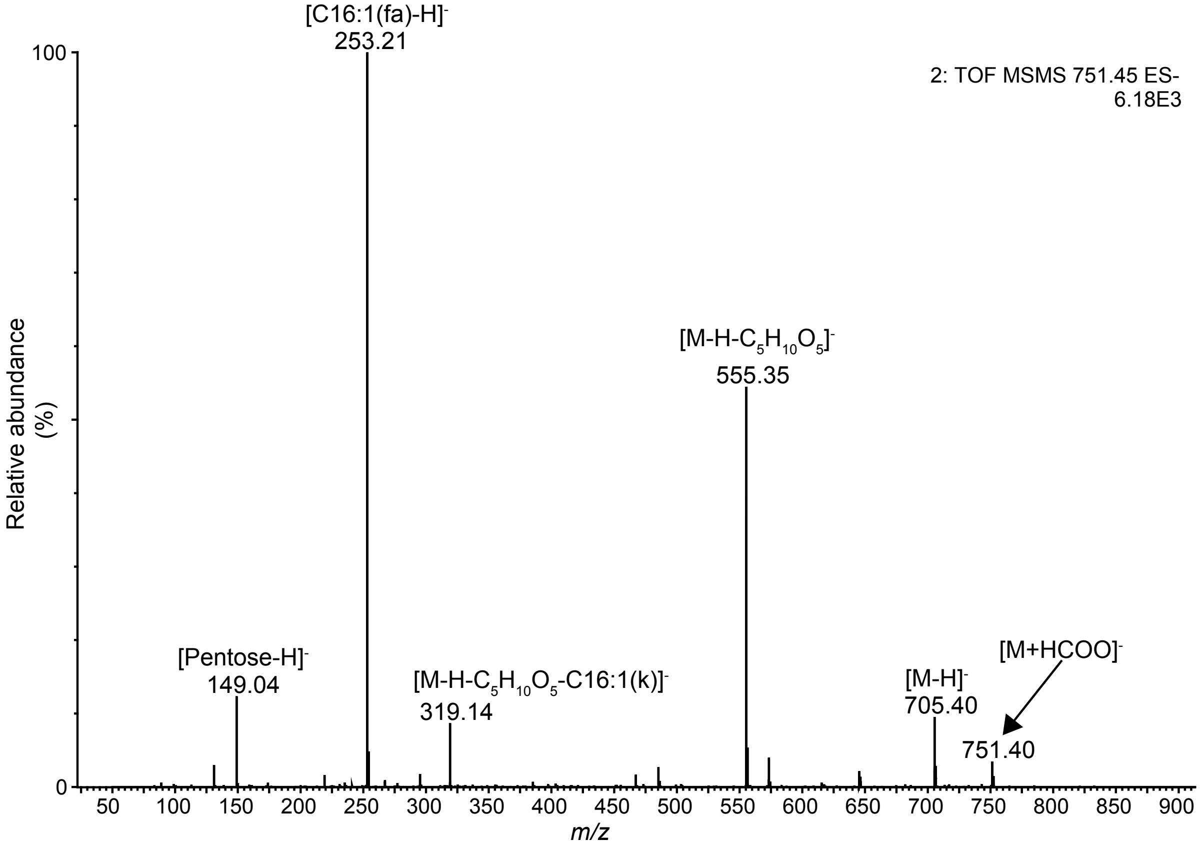


**Figure S6. Negative mode MS/MS fragmentation of H3:24(4,4,16-O-p) from *S. sisymbriifolium* results in neutral loss of a pentose moiety.** This neutral loss of a sugar group ($C_5H_{10}O_5$ plus formic acid) from the [M+formate]$^-$ ion *m/z* 751.4025 to form *m/z* 555.3522 is not observed with negative CID of other acyldisaccharides (Figures S2 and S7) suggesting an unusual glycosidic linkage on the acyl chain. fa = fatty acid; k = ketene; 16-O-p = pentosylated 16 carbon acyl chain.

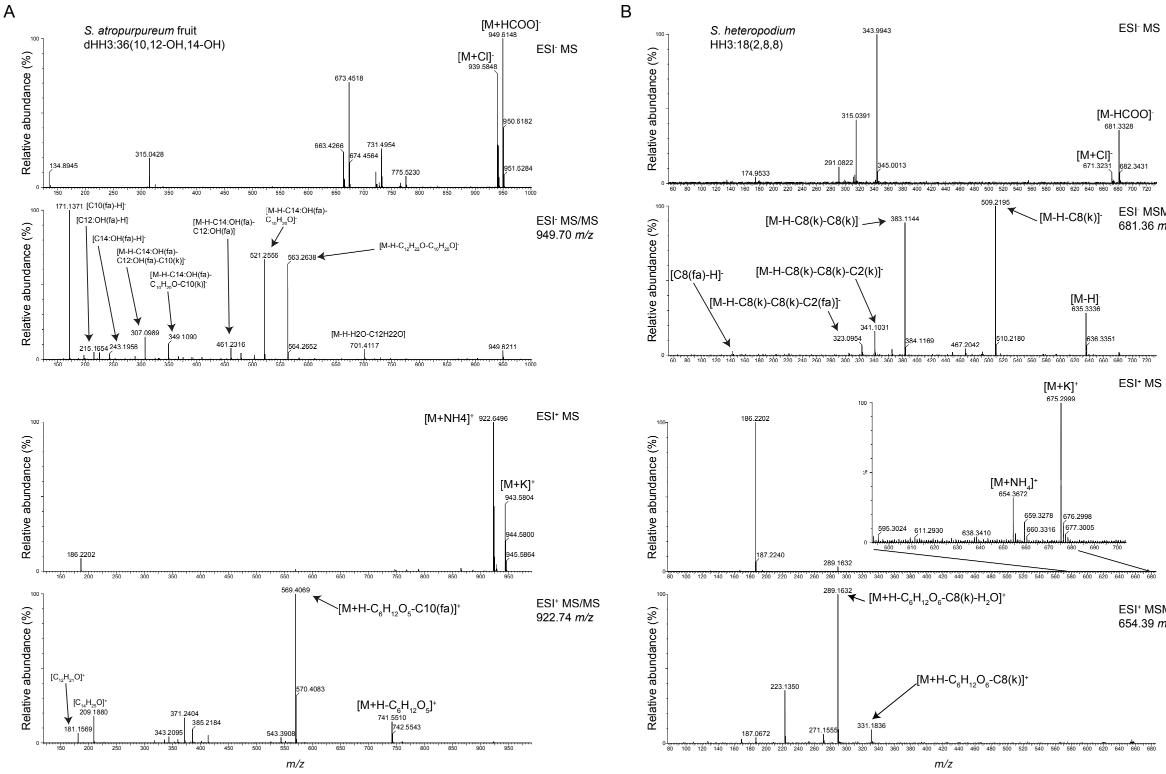


**Figure S7. Negative and positive mode MS and CID MS/MS fragmentation of dHH3:36(10,12-OH,14-OH) from a *S. atropurpureum* fruit surface extract and HH3:18(2,8,8) from a *S. heteropodium* leaf surface extract.** Formate and ammonium adduct ions were selected by data-dependent acquisition software and fragmented with a ramped collision energy detailed in the Methods. Under negative mode CID MS/MS, each compound produces fragment ions corresponding to stepwise acyl chain loss down to a free sugar core fragment ion. In contrast, under positive mode CID MS/MS each compound produces fragment ions corresponding to neutral loss of a sugar moiety**. (A)** dHH3:36(10,12-OH,14-OH) MS and MS/MS fragmentation. Positive mode CID produces a fragment ion (*m/z* 569.4069) corresponding to a neutral loss of a deoxyhexose acylated with a ten carbon acyl chain. dHH = deoxyhexose-hexose **(B)** HH3:18(2,8,8) MS and MS/MS fragmentation. Positive mode CID produces a fragment ion (*m/z* 331.1836) corresponding to a neutral loss of a hexose acylated with an eight carbon acyl chain. HH = hexose-hexose; fa = fatty acid; k = ketene.

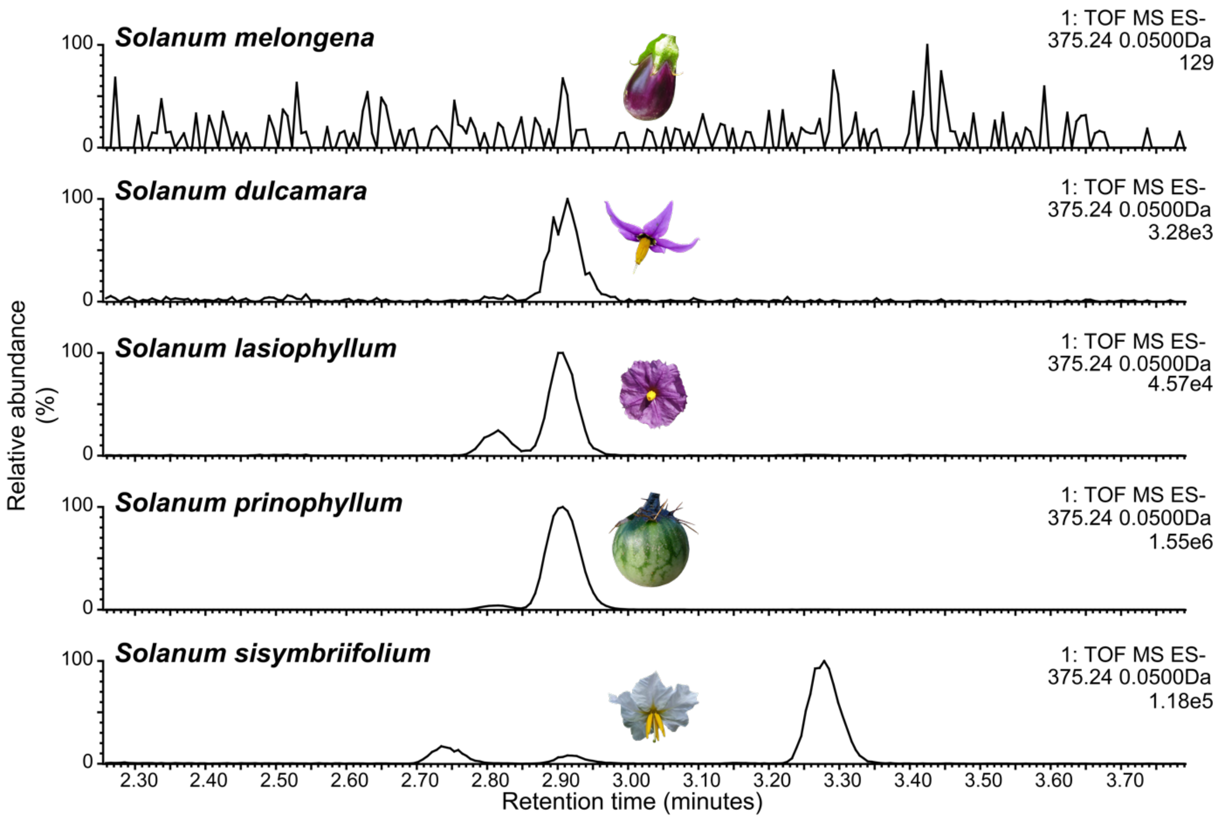


**Figure S8. Acylsugar saponification produces free glycosylated fatty acids in four *Solanum* species supporting the identification of glycohydroxyacyl hexoses.** All traces are extracted ion LC/MS chromatograms for pentosylated hydroxytetradecanoic acid, *m/z* 375.24±0.05 (retention time 2.90 min) from saponified leaf surface extracts. *S. melongena* does not contain detectable glycosylated fatty acids and acts as a negative control. Vertical scale is normalized to the largest signal within the displayed region (values in the upper right of each chromatogram).

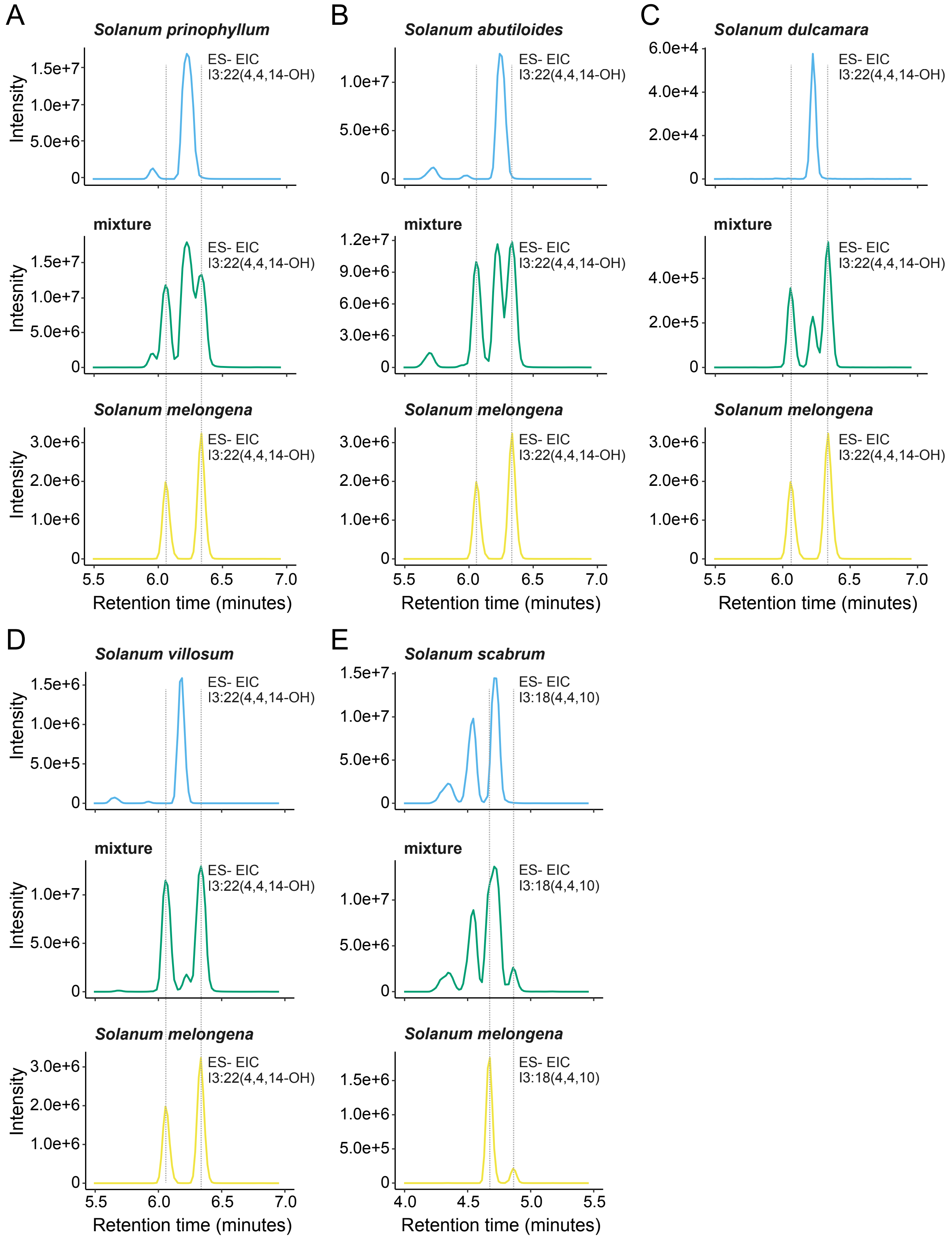
A
Solanum prinophyllum
mixture
Solanum melongena
B
Solanum abutiloides
C
Solanum dulcamara
D
Solanum villosum
E
Solanum scabrum
ES- EIC
I3:22(4,4,14-OH)
I3:18(4,4,10)
Intensity
Intesnity
Retention time (minutes)

**Figure S9. Coelution analysis reveals that five *Solanum* species accumulate acylsugar positional or branching isomers that differ from *S. melongena* acylsugars.** Extracted ion chromatograms were generated for combined signals of *m/z* 591.34 and 519.28 corresponding to I3:22(4,4,14-OH) and I3:18(4,4,10) $[M+formate]^-$ adducts, respectively. The yellow traces (bottom row) represent *S. melongena*, the blue traces represent the five other *Solanum* species, **(A)** *S. prinophylum,* **(B)** *S. abutiloides,* **(C)** *S. dulcamara,* **(D)** *S. villosum*, and **(E)** *S. scabrum*, and the middle row green traces represent acylsugar mixtures between *S. melongena* and the species in the trace below. The mixed samples allow for corrections of retention time drift that may occur between samples.

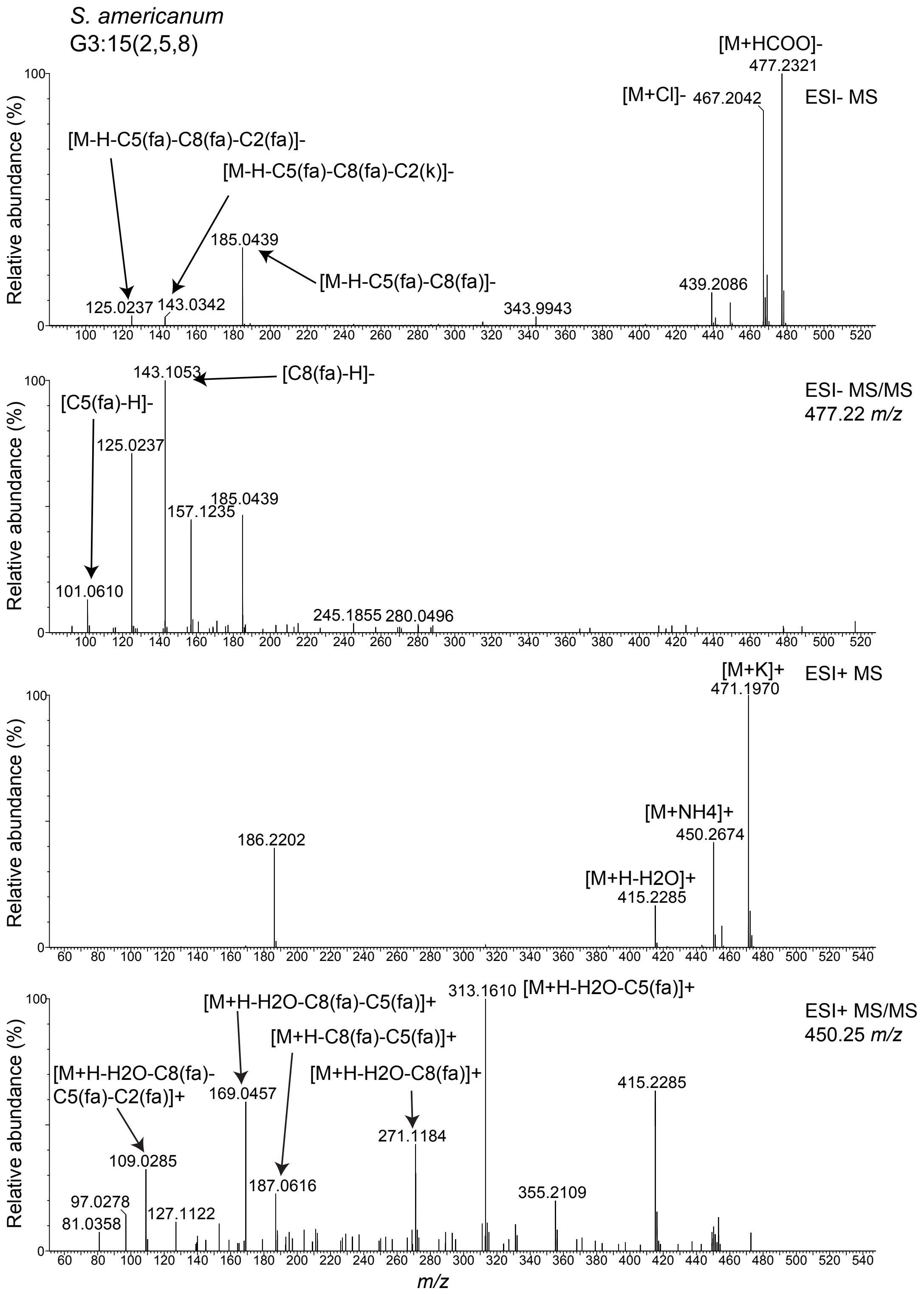

S. americanum
G3:15(2,5,8)
[M+HCOO]-
477.2321
[M+Cl]-
467.2042
ESI- MS
[M-H-C5(fa)-C8(fa)-C2(fa)]-
[M-H-C5(fa)-C8(fa)-C2(k)]-
185.0439
[M-H-C5(fa)-C8(fa)]-
125.0237
143.0342
343.9943
439.2086
143.1053
[C8(fa)-H]-
ESI- MS/MS
477.22 m/z
[C5(fa)-H]-
125.0237
185.0439
157.1235
101.0610
245.1855
280.0496
[M+K]+
471.1970
ESI+ MS
[M+NH4]+
450.2674
186.2202
[M+H-H2O]+
415.2285
[M+H-H2O-C8(fa)-C5(fa)]+
313.1610
[M+H-H2O-C5(fa)]+
ESI+ MS/MS
450.25 m/z
[M+H-C8(fa)-C5(fa)]+
[M+H-H2O-C8(fa)-
C5(fa)-C2(fa)]+
169.0457
[M+H-H2O-C8(fa)]+
415.2285
271.1184
109.0285
187.0616
355.2109
97.0278
81.0358
127.1122
Relative abundance (%)
m/z

**Figure S10. Negative and positive mode MS and CID MS/MS fragmentation of G3:15(2,5,8) from *S. americanum*.** Acylglucoses fragment characteristically in negative mode MS and MS/MS functions producing fragment ions corresponding to stepwise acyl chain loss. This is in stark contrast to acylinositol negative mode fragmentation which does not produce major fragment ions corresponding to stepwise acyl chain loss (Figure S2). This fragmentation difference distinguishes the two sugar cores and enables their annotation by LC-MS. G3:15 formate and ammonium adduct ions were selected by data-dependent acquisition software and fragmented with a ramped collision energy detailed in the Methods. fa = fatty acid; k = ketene.

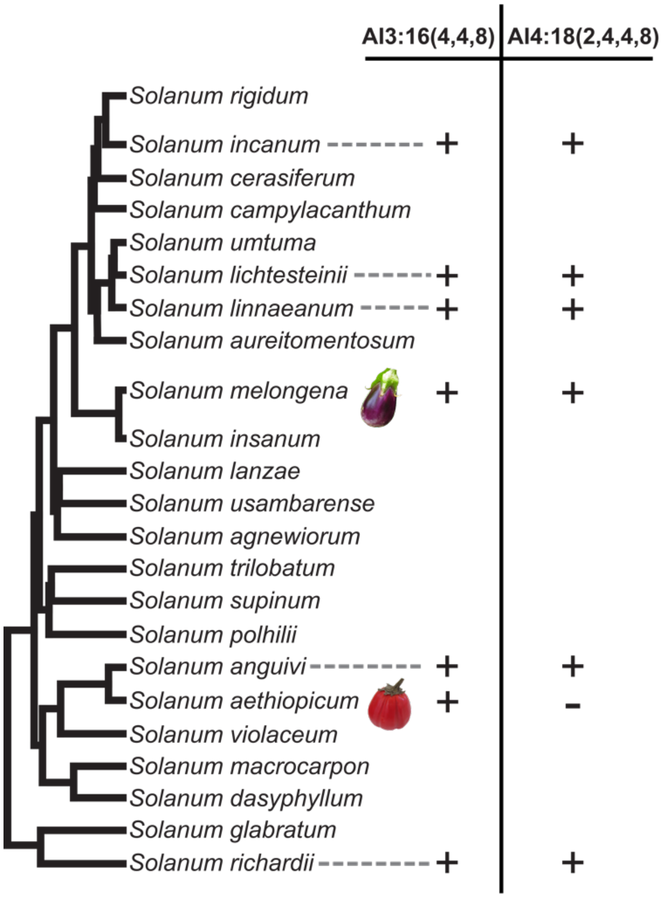


**Figure S11. Phylogenetic analysis of Eggplant Clade and Anguivi Grade species reveals *S. aethiopicum* is the only analyzed acylsugar-producing species to not accumulate detectable AI4:18(2,4,4,8).** AI3:16(4,4,8) and AI4:18(2,4,4,8) presence and absence in surface extracts as detected by LC-MS are plotted on a phylogeny of the Eggplant Clade and Anguivi Grade. Species that either do not accumulate detectable acylsugars (*S. macrocarpon* and *S. virginianum)* or were not analyzed did not have AI3:16 and AI4:18 presence plotted. The phylogeny was modified from a previously published version (Aubriot et al., 2018). Full acylsugar profiles are detailed Tables 1, S3-8.

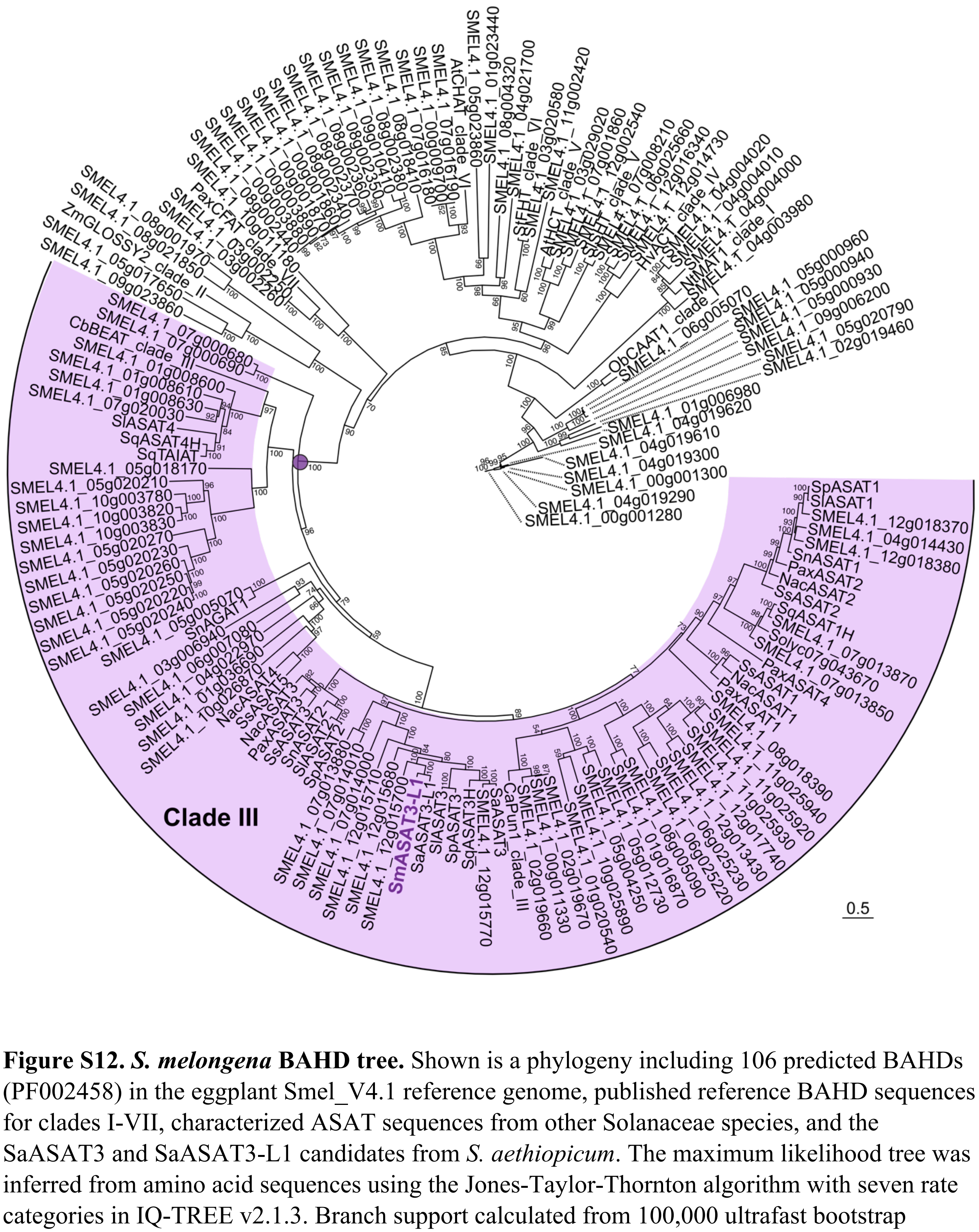


**Figure S12.** ***S. melongena*** **BAHD tree.** Shown is a phylogeny including 106 predicted BAHDs (PF002458) in the eggplant Smel_V4.1 reference genome, published reference BAHD sequences for clades I-VII, characterized ASAT sequences from other Solanaceae species, and the SaASAT3 and SaASAT3-L1 candidates from *S. aethiopicum*. The maximum likelihood tree was inferred from amino acid sequences using the Jones-Taylor-Thornton algorithm with seven rate categories in IQ-TREE v2.1.3. Branch support calculated from 100,000 ultrafast bootstrap replicates. Clade III BAHDs are highlighted in purple, with a purple dot marking the node. Branch lengths represent substitution rates (substitutions per site) as denoted by scale bar.

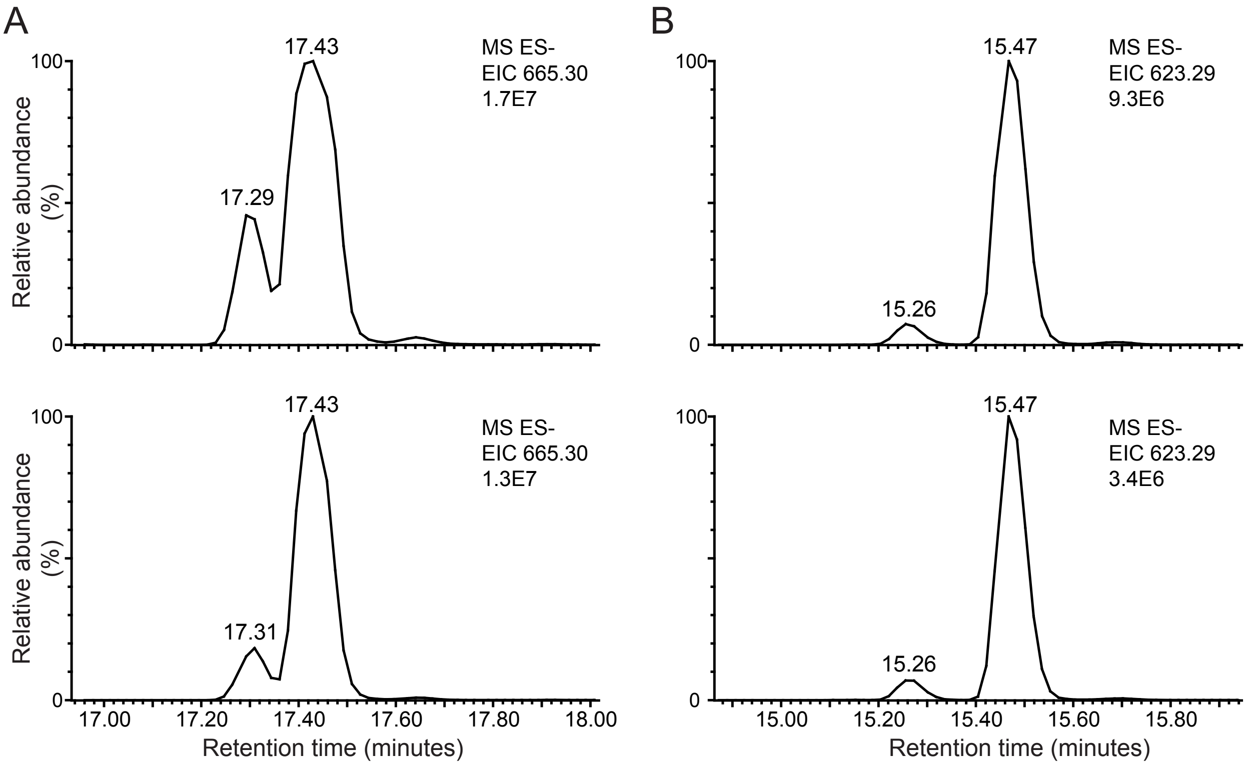


**Figure S13. SmASAT3-L1 forward and reverse assay produced AI4:18 and AI3:16, respectively, coelute with plant produced AI4:18 and AI3:16. (A)** Forward assay produced AI4:18 (top) coelutes with AI4:18 from a S. melongena 555598 leaf surface extract (bottom). The extracted ion chromatograms display the formate adduct of AI4:18, *m/z* 665.30. **(B)** Reverse assay produced AI3:16 (top) coelutes with AI3:16 from a S. melongena 555598 leaf surface extract (bottom). The extracted ion chromatograms display the formate adduct of AI3:16, *m/z* 623.20.

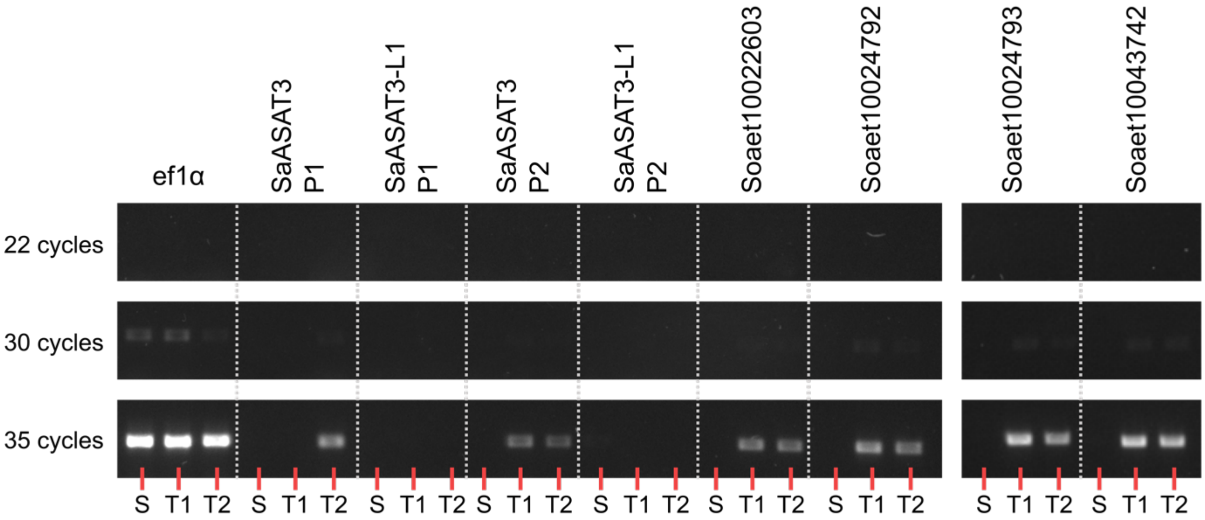


**Figure S14. Semi-quantitative RT-PCR analysis of *S. aethiopicum* BAHD expression in glandular trichomes.** *Elongation factor1α* (EF1α) was used as a positive control. P1 = primer pair 1; P2 = primer pair 2; S = shaved hypocotyl; T1 = PI 666075 glandular trichomes; T2 = Grif 14165 glandular trichomes.

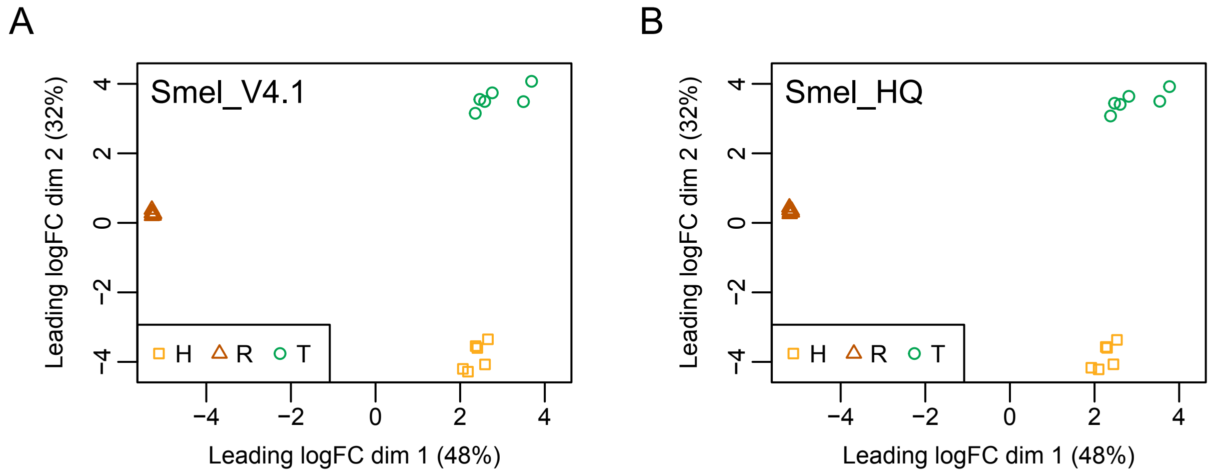


**Figure S15. MDS plot of *S. melongena* RNAseq data.** Genome-wide expression patterns across eggplant trichomes, trichomeless hypocotyls, and roots. Depicted are multidimensional scaling (MDS) plot demonstrating that our 18 eggplant RNAseq samples cluster tightly by tissue identity when reads were mapped against the **(A)** Smel_V4.1 or **(B)** Smel_HQ reference genome. Distance between points illustrates expression differences between pairs of samples, calculated as leading (i.e., largest absolute) log2 fold-change (FC) in two dimensions, dim 1 and dim 2. Along the y-axis (dim 2), samples are separated into three tissue-specific groups. Along the x-axis (dim 1), samples are separated into two groups representing aerial (trichomes and trichomeless hypocotyls) and subterranean (roots) tissues, respectively. Colors and shapes represent the three tissues. T, trichomes (green circles); H, trichomeless hypocotyls (yellow squares); R, roots (brown triangles).

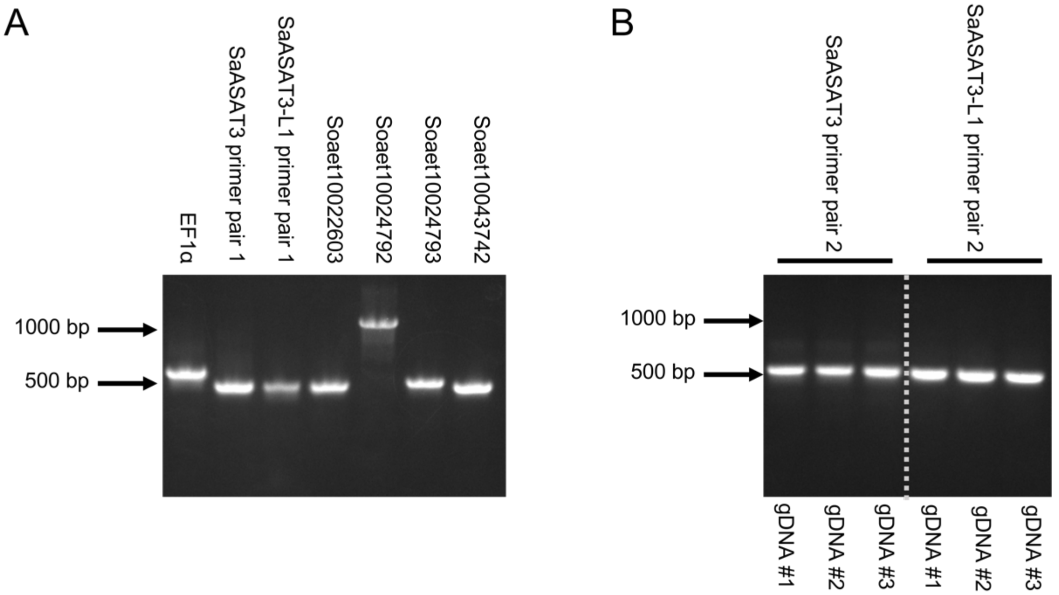


**Figure S16.** ***S. aethiopicum*** **RT-PCR primers validated with gDNA controls.** Soaet10024792 primers amplified a region containing an intron resulting in the larger amplicon length.
