## Supplementarty NMR data and metadata for "Trading acyls and swapping sugars: metabolic innovations in *Solanum* trichomes"

**Supplementary Data File S1**
**AI3:16(i4,i4,i8) Chemical shifts and coupling constants.**

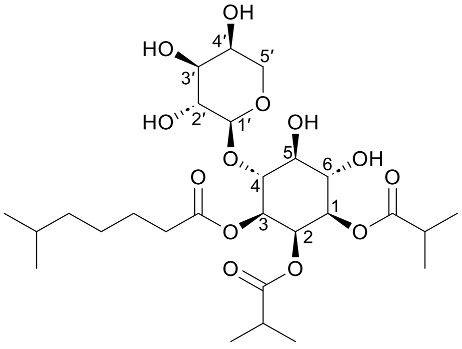


AI3:16(i4,i4,i8)

Molecular Formula: $C_{27}H_{46}O_{13}$

Instrument: Agilent 500 MHz DDR2

NMR solvent: $CD_3CN$
Fractions: 32-37

InChI Key: LLNCXHBBKCPKDQ-ZPDJPJEFSA-N

SMILES:
CC(C)CCCCC(=O)O[C@H]1[C@H](O[C@@H]2OC[C@H](O)[C@H](O)[C@H]2O)[C@@H](O)[C@H](O)[C@@H](OC(=O)C(C)C)[C@H]1OC(=O)C(C)C

| Carbon # (Group) | $^1$H ($\delta$, ppm) | $^{13}$C ($\delta$, ppm) |
|---|---|---|
| 1(CH) | 4.86 (dd, *J* = 2.91, 10.24 Hz) | 70.92 |
| -1(CO) | | 176.00 |
| -2(CH) | 2.61 (hept, *J* = 6.97 Hz) | 33.80 |
| -3,4($CH_3$) | 1.08 (d, *J* = 6.97 Hz) | 18.02 |
| | 1.06 (d, *J* = 6.97 Hz) | 18.24 |
| 2(CH) | 5.50 (t, *J* = 2.93 Hz) | 68.10 or 68.08[a] |
| -1(CO) | | 175.88 |
| -2(CH) | 2.47 (hept, *J* = 6.98 Hz) | 33.69 or 33.64[a] |
| -3,4($CH_3$) | 1.20 (d, *J* = 6.97 Hz) | 18.41, 18.40 |
| 3(CH) | 4.99 (dd, *J* = 3.04, 10.22 Hz) | 70.16 |
| -1(CO) | | 172.71 |
| -2($CH_2$) | 2.26 (m) | 33.69 or 33.64[a] |
| -3($CH_2$) | 1.53 (m) | 24.53 |
| -4($CH_2$) | 1.29 (m) | 26.57 |
| -5($CH_2$) | 1.19 (m) | 38.34 |
| -6(CH) | 1.55 (m) | 27.57 |
| -7,8($CH_3$) | 0.84 (d, *J* = 6.60 Hz) | 21.89 |
| 4(CH) | 3.85 (t, *J* = 10.00 Hz) | 80.60 |
| 5(CH) | 3.47 (t, *J* = 9.30 Hz) | 72.49 |
| 6(CH) | 3.78 (t, *J* = 9.85 Hz) | 70.66 |
| 1′(CH) | 4.26 (d, *J* = 7.01 Hz) | 104.17 |
| 2′(CH) | 3.41 (dd, *J* = 7.0, 9.36 Hz) | 71.10 |
| 3′(CH) | 3.47 (dd, *J* = 3.4, 9.3 Hz) | 72.49 |
| 4′(CH) | 3.73 (m) | 68.10 or 68.08[a] |
| 5′($CH_2$) | 3.88 (dd, *J* = 2.41, 12.58 Hz) | 66.13 |
| | 3.54 (dd, *J* = 1.54, 12.63 Hz) | |

a – $^{13}$C signals not resolved in 2D spectra.

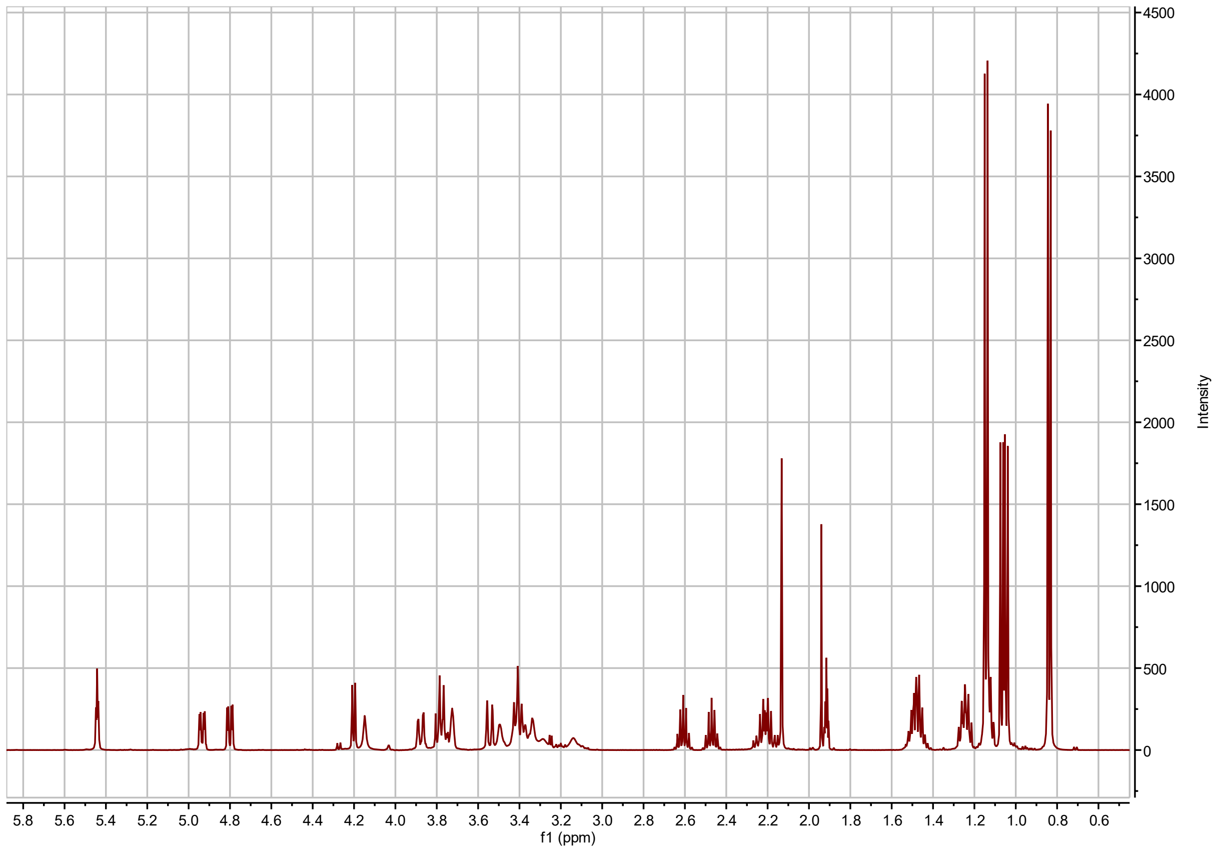


**AI3:16(4,4,8) $^{1}$H NMR**

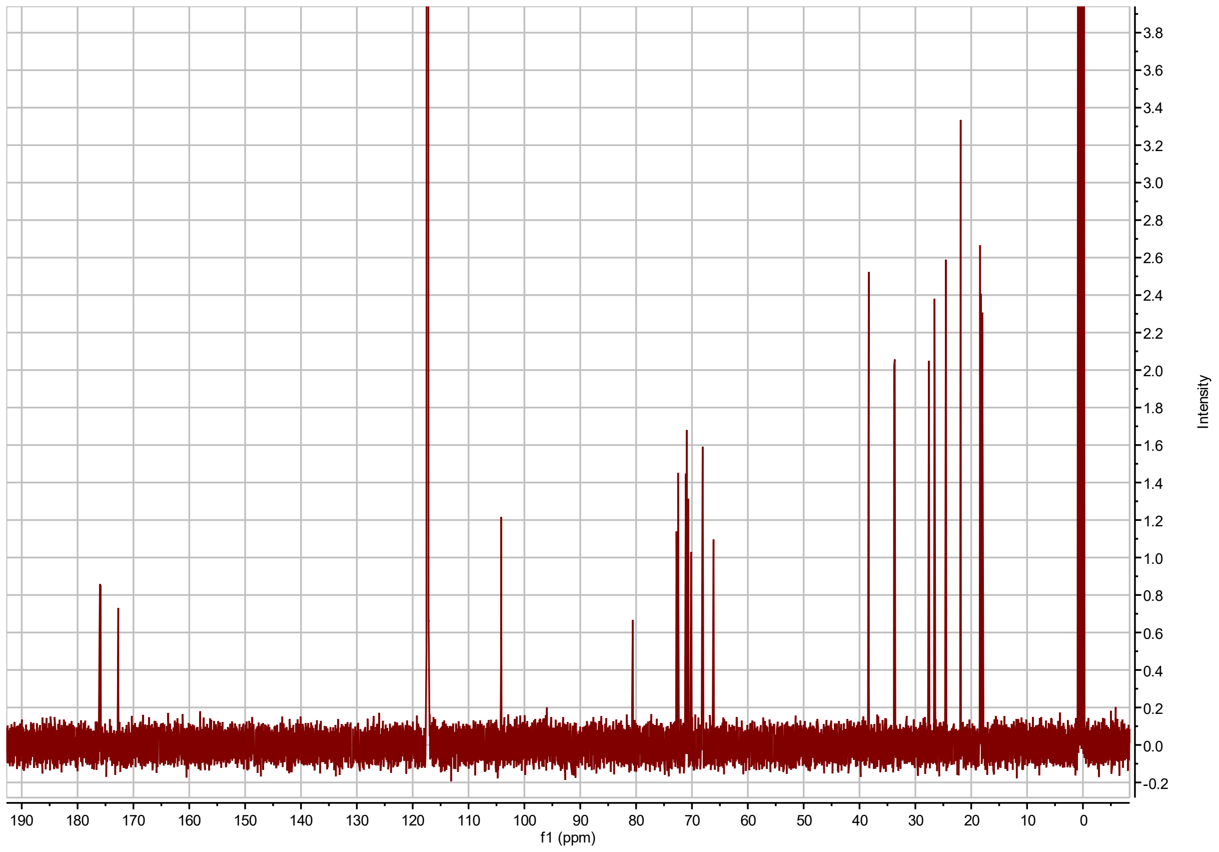


**AI3:16(4,4,8) $^{13}$C NMR**

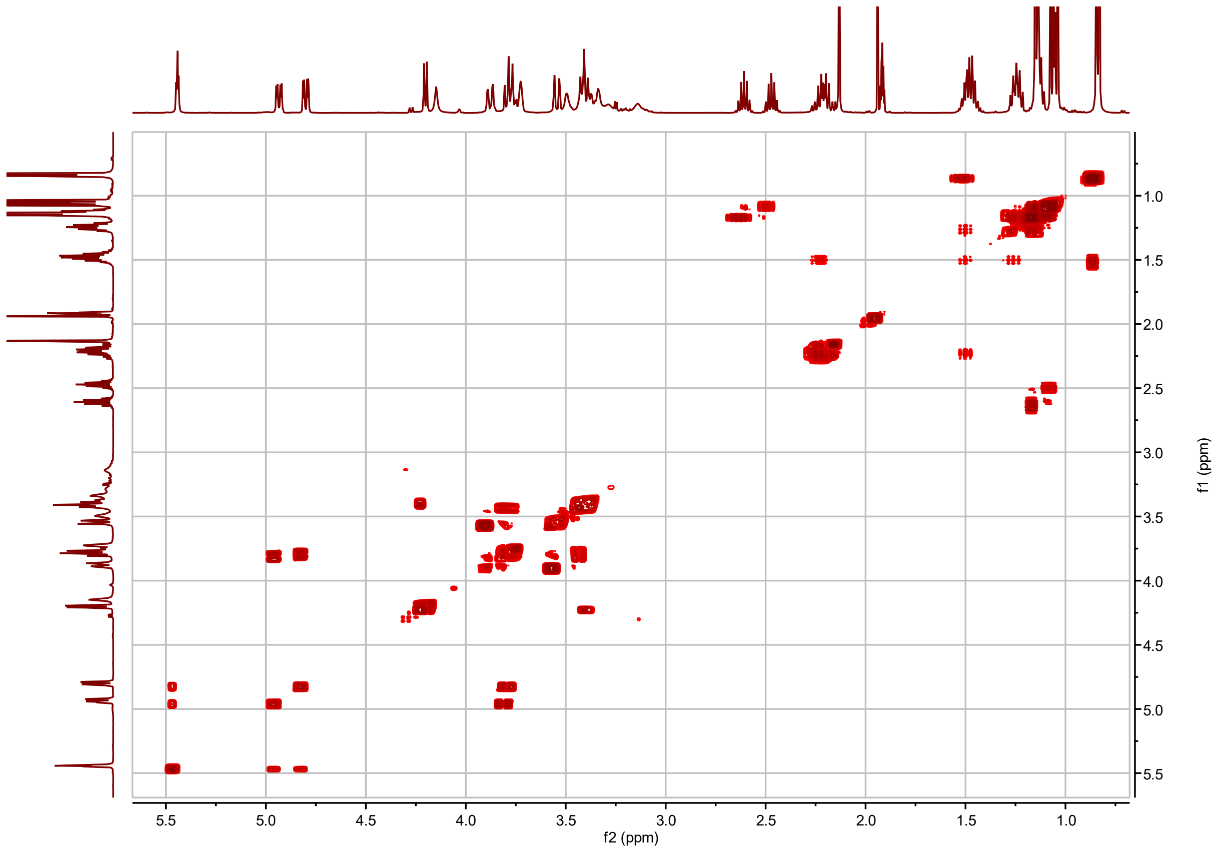


**AI3:16(4,4,8) $^{1}H$-$^{1}H$ COSY**

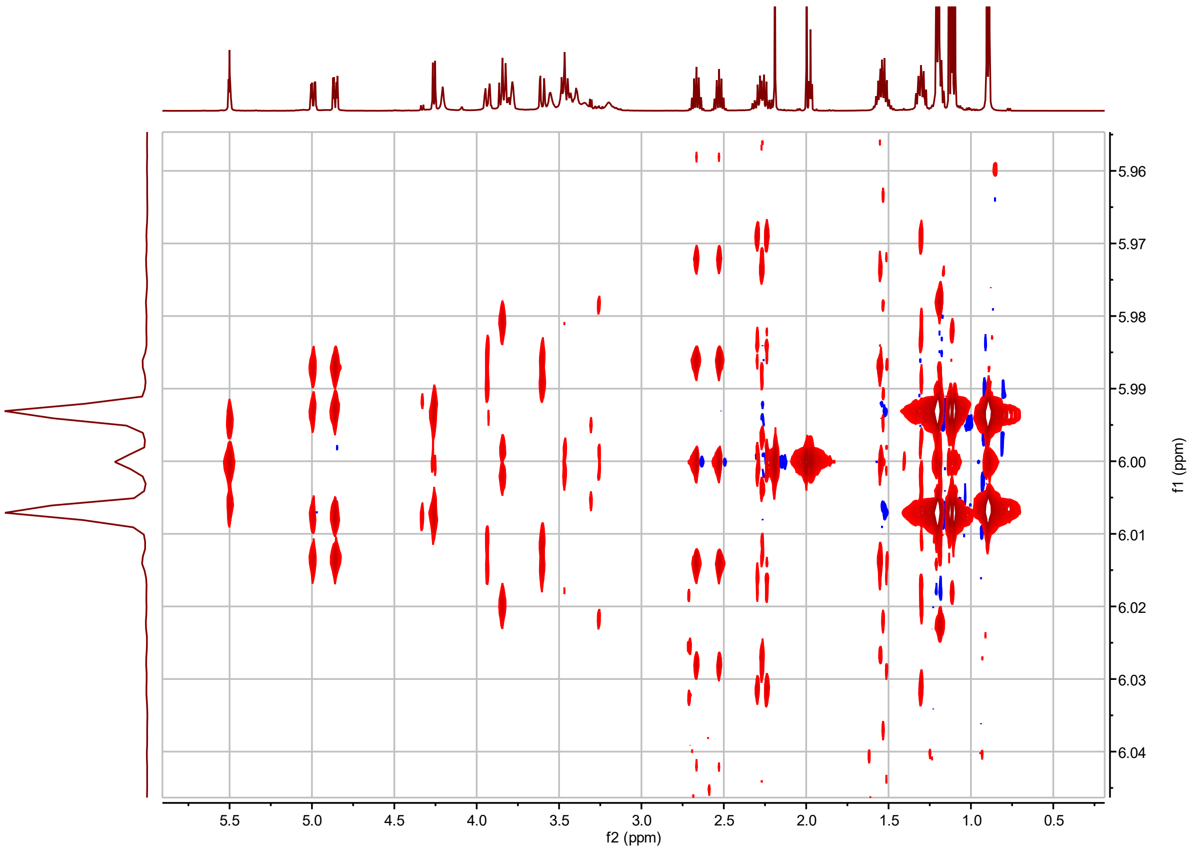


**AI3:16(4,4,8) *J*-resolved**

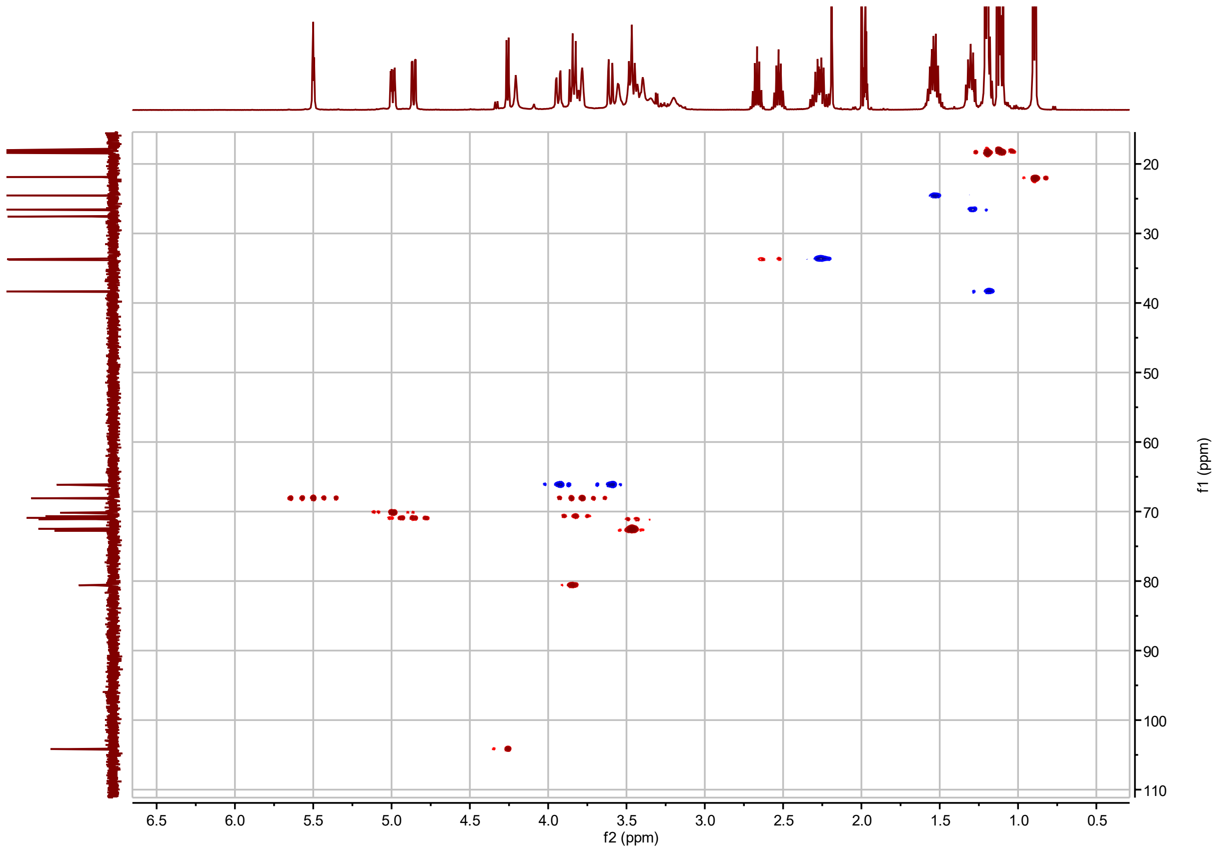


**AI3:16(4,4,8) $^{1}H$-$^{13}C$ HSQC**

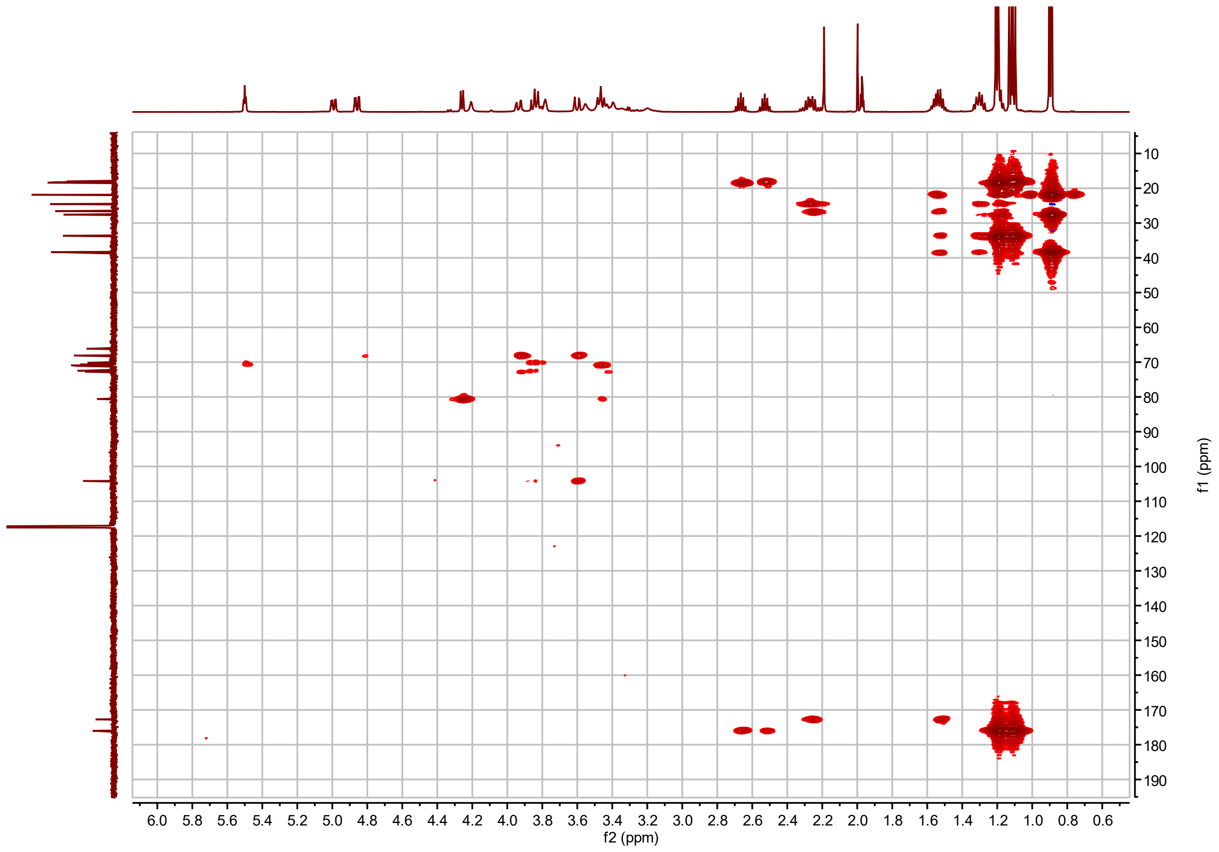


**AI3:16(4,4,8) $^{1}$H-$^{13}$C HMBC with apodization optimized for correlations between acyl chain carbonyl carbon and C2 proton(s)**

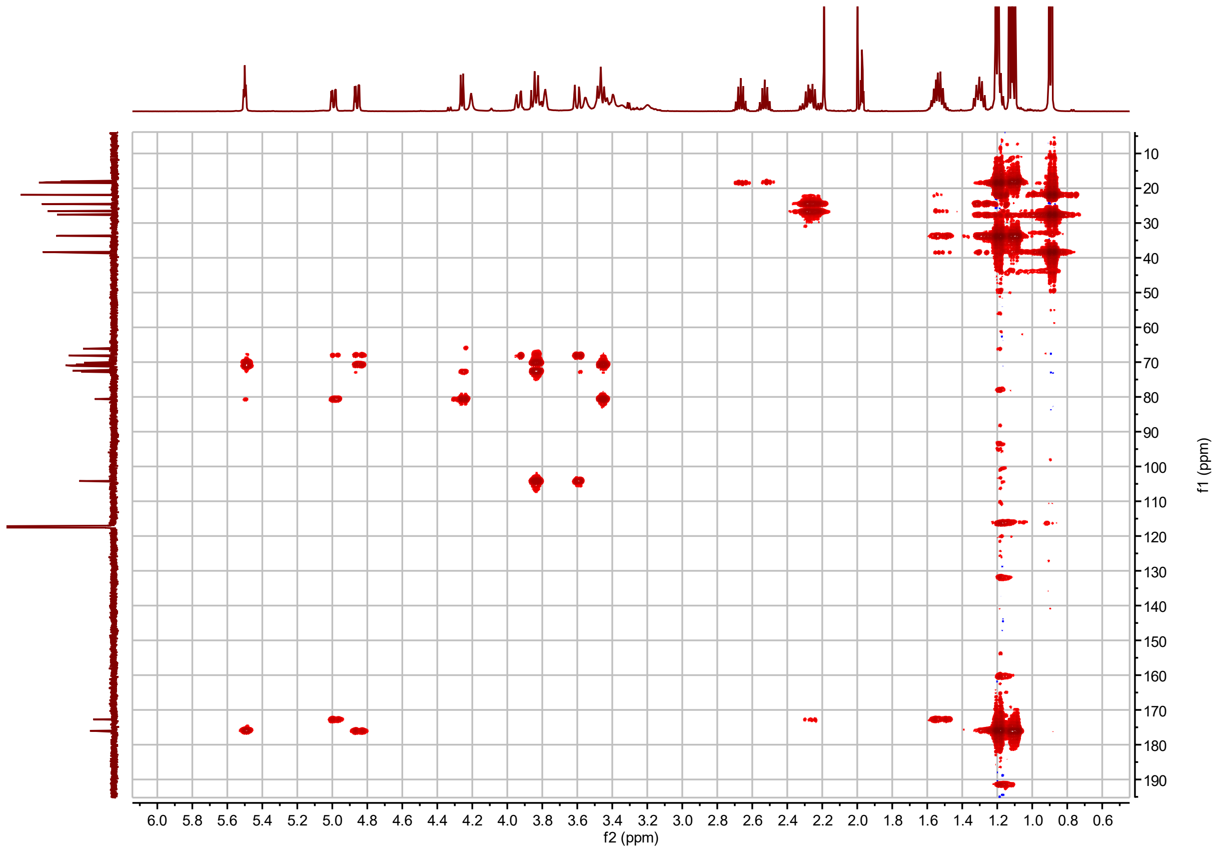


**AI3:16(4,4,8) $^{1}H$-$^{13}C$ HMBC with apodization optimized for correlations between acyl chain carbonyl carbon and sugar ring protons**

**AI4:18(2,i4,i4,i8) Chemical shifts and coupling constants.**

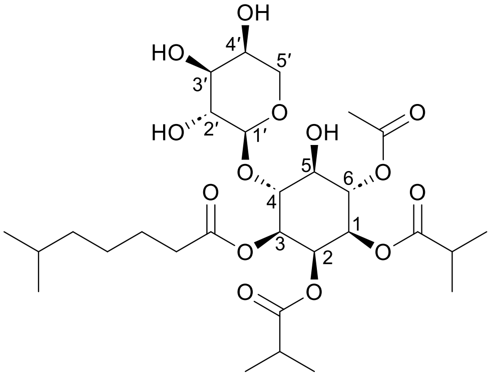


AI4:18(2,i4,i4,i8)

Molecular Formula: $C_{29}H_{48}O_{14}$

Instrument: Agilent 500 MHz DDR2 and Varian 600 MHz Inova

NMR solvent: $CD_3CN$

Fractions: 44-52

InChI Key: JIKVNSLWQPFKKW-XJPFAFDOSA-N

SMILES:
CC(C)CCCCC(=O)O[C@H]1[C@H](O[C@@H]2OC[C@H](O)[C@H](O)[C@H]2O)[C@@H](O)[C@H](OC(C)=O)[C@@H](OC(=O)C(C)C)[C@H]1OC(=O)C(C)C

| Carbon # (Group) | $^{1}$H($\delta$, ppm) | $^{13}$C ($\delta$, ppm) |
|---|---|---|
| 1(CH) | 5.03 (dd, $J$ = 2.9, 10.5 Hz) | 68.88 |
| -1(CO) | | 175.51 |
| -2(CH) | 2.45 (hept, $J$ = 7.0 Hz) | 33.67 |
| -3,4($CH_3$) | 1.07 (d, $J$ = 7.0 Hz) | 17.85 |
| 2(CH) | 5.54 (t, $J$ = 2.97 Hz) | 67.87 |
| -1(CO) | | 175.79 |
| -2(CH) | 2.69 (hept, $J$ = 6.94 Hz) | 33.78 |
| -3,4($CH_3$) | 1.21 (t, $J$ = 6.80 Hz) | 18.46 |
| 3(CH) | 5.05 (dd, $J$ = 2.9, 10.5 Hz) | 69.72 |
| -1(CO) | | 172.68 |
| -2($CH_2$) | 2.29 (m) | 33.60 |
| -3($CH_2$) | 1.53 (m) | 24.52 |
| -4($CH_2$) | 1.30 (m) | 26.54 |
| -5($CH_2$) | 1.19 (m) | 38.32 |
| -6(CH) | 1.55 (m) | 27.56 |
| -7,8($CH_3$) | 0.89 (d, $J$ = 6.6 Hz) | 21.88 |
| 4(CH) | 3.94 (t, $J$ = 9.7 Hz) | 80.93 |
| 5(CH) | 3.67 (t, $J$ = 9.48 Hz) | 70.22 |
| 6(CH) | 5.35 (t, $J$ = 10.10 Hz) | 71.07 |
| -1(CO) | | 169.75 |
| -2($CH_3$) | 2.02 (s) | 20.08 |
| 1′(CH) | 4.25 (d, $J$ = 6.98 Hz) | 104.22 |
| 2′(CH) | 3.43 (dd, $J$ = 7.00, 9.36 Hz) | 71.07 |
| 3′(CH) | 3.47 (dd, $J$ = 3.4, 9.3 Hz) | 72.80 |
| 4′(CH) | 3.78 (m) | 68.12 |
| 5′($CH_2$) | 3.93 (dd, $J$ = 2.5, 12.5 Hz)<br>3.61 (dd, $J$ = 1.5, 12.7 Hz) | 66.24 |

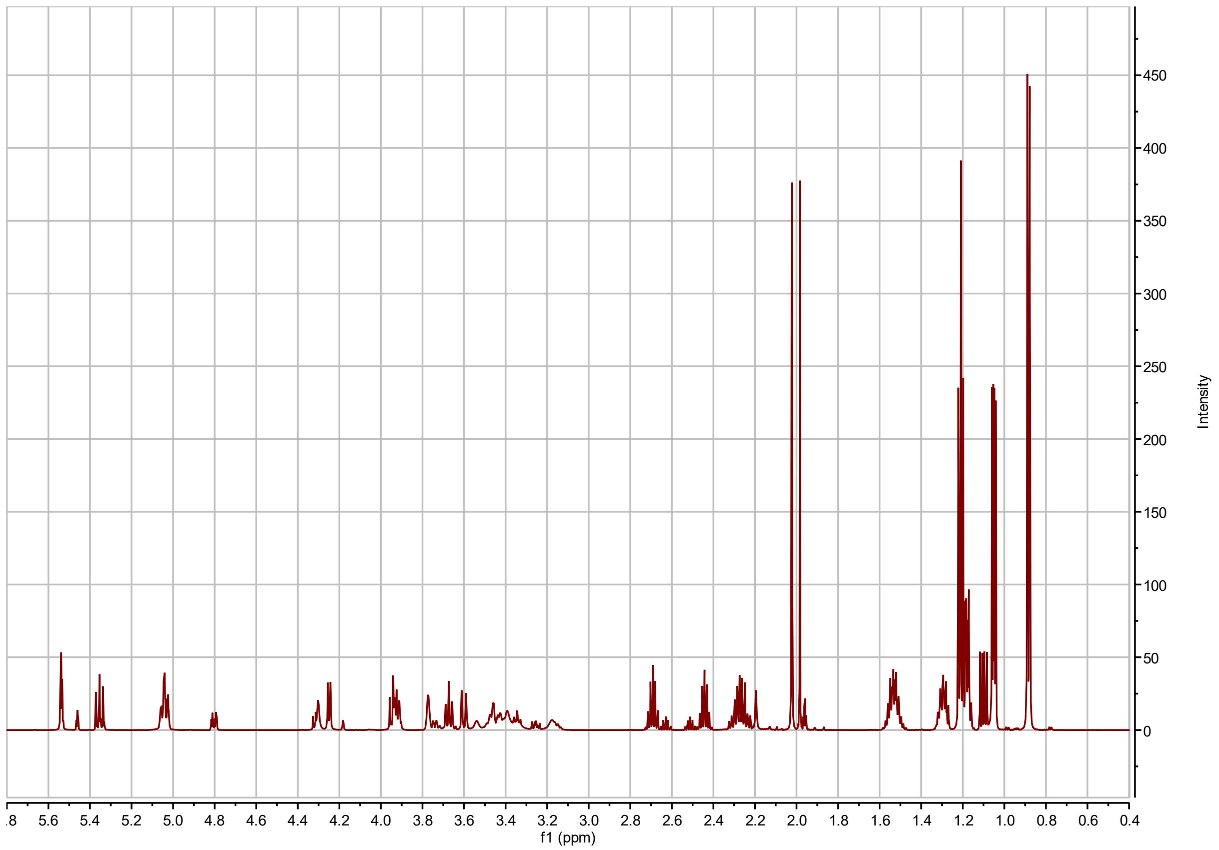


**AI4:18(2,4,4,8) $^{1}$H NMR**

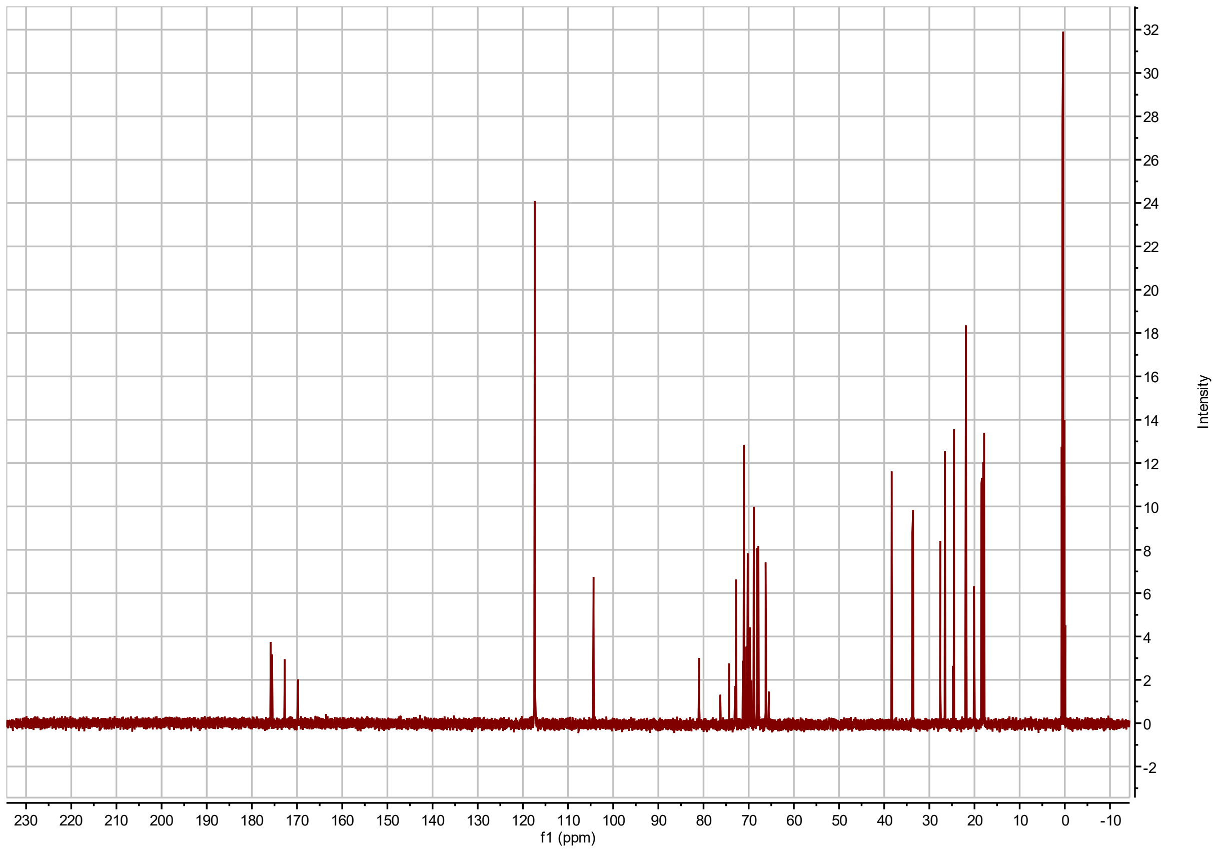


**AI4:18(2,4,4,8) $^{13}$C NMR**

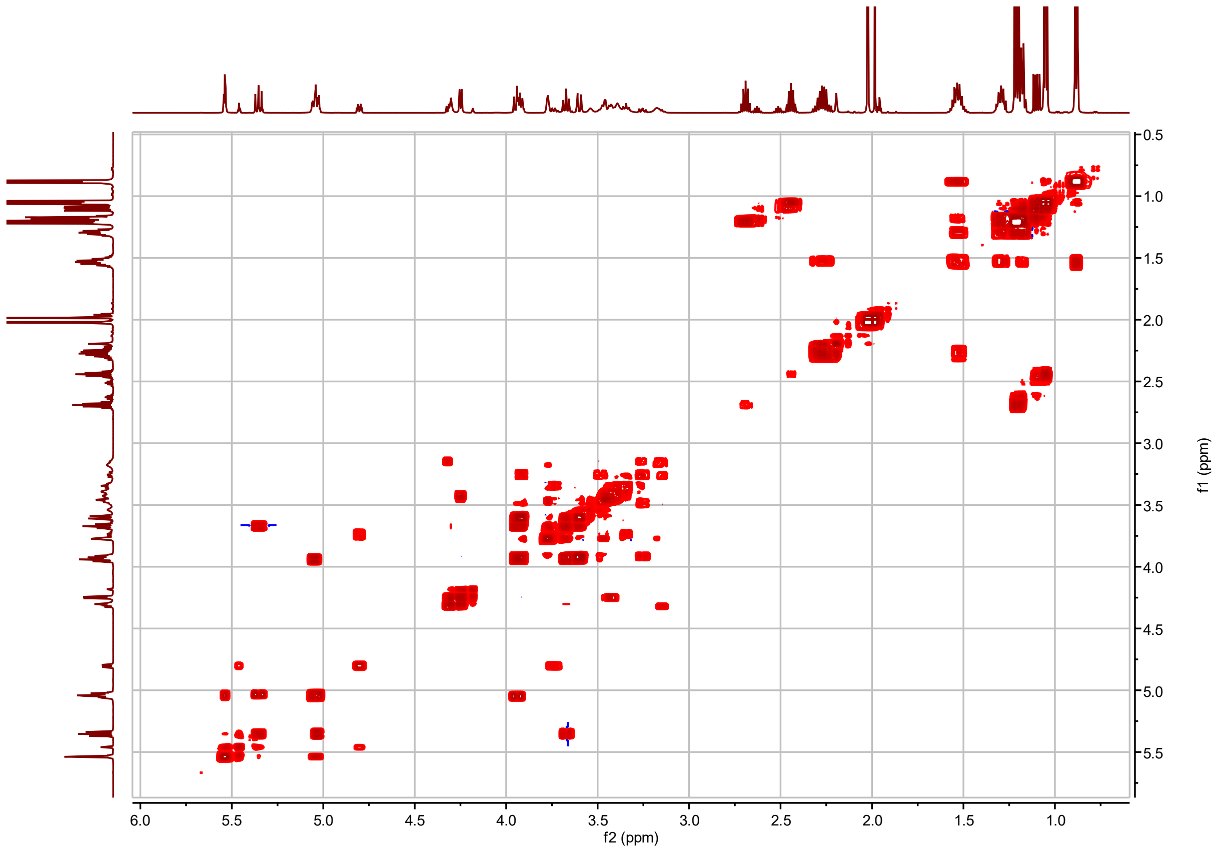


**AI4:18(2,4,4,8) $^{1}$H-$^{1}$H COSY**

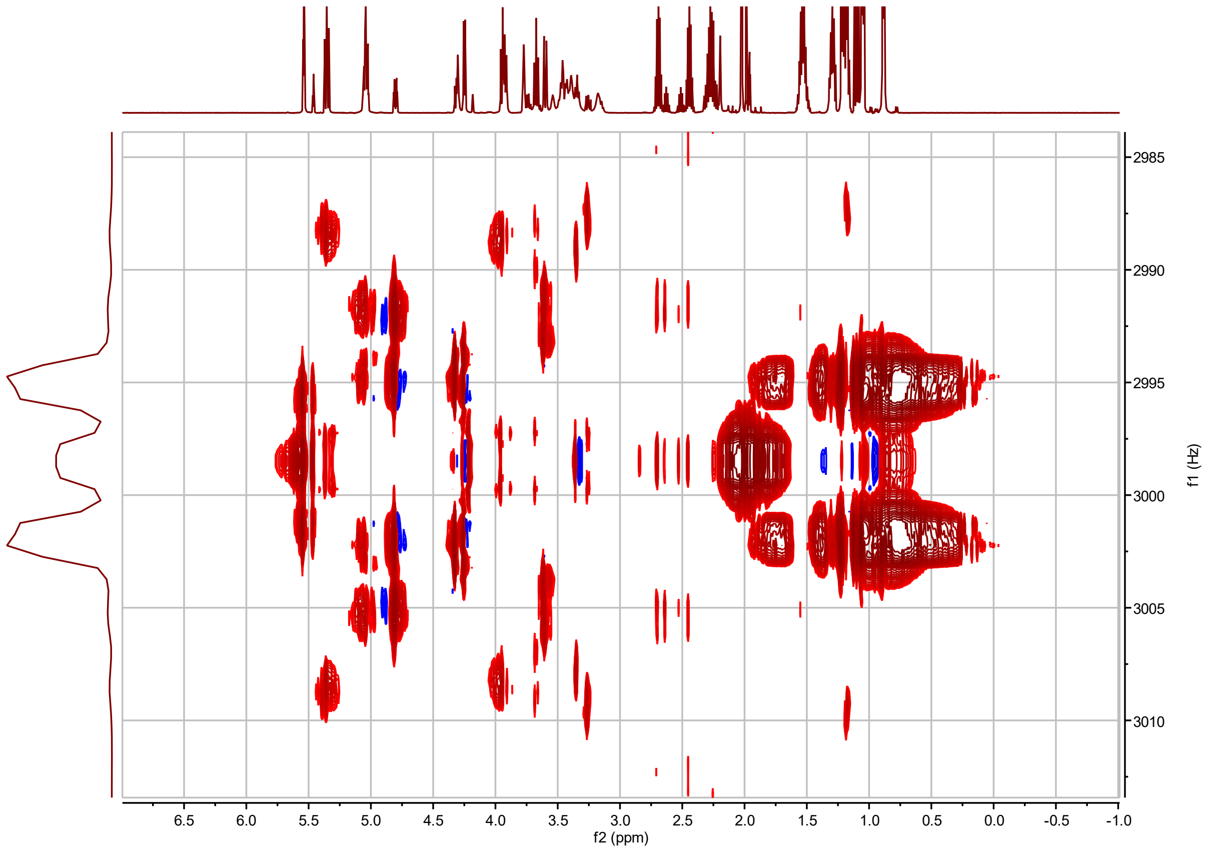


**AI4:18(2,4,4,8) *J*-resolved**

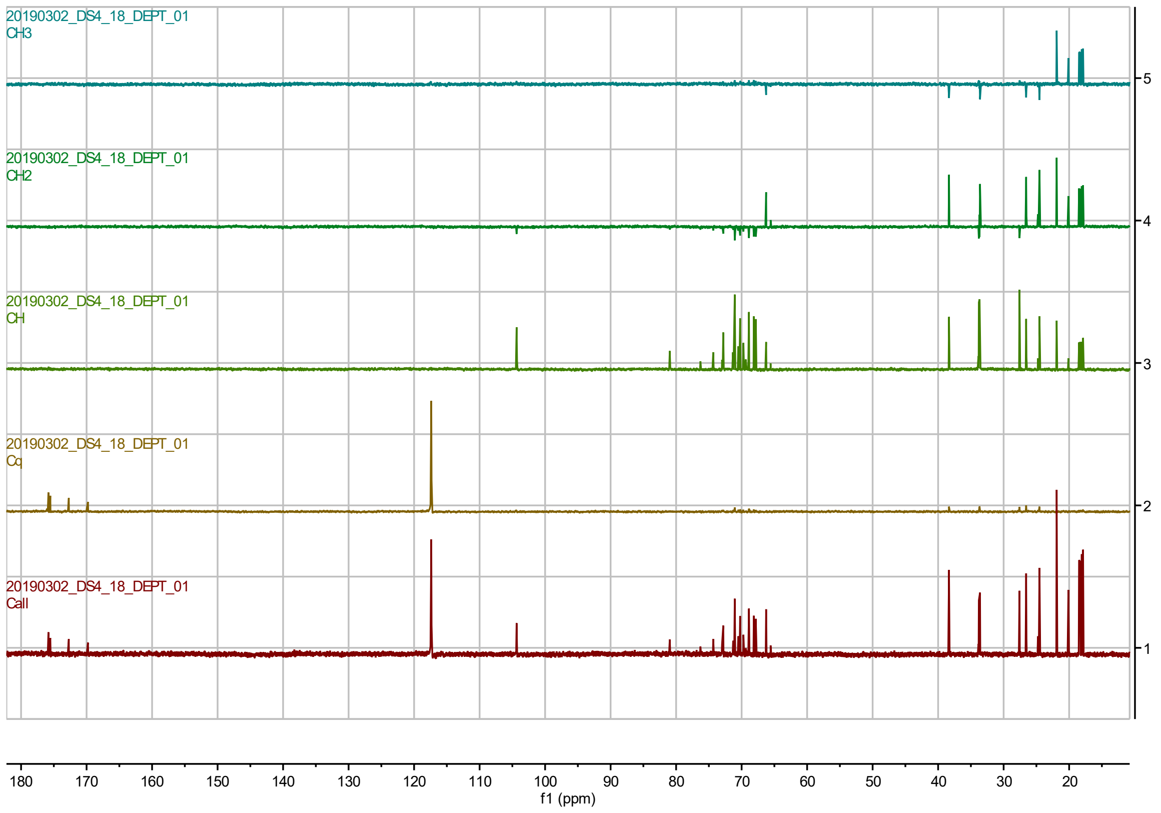


**AI4:18(2,4,4,8) $^{13}$C DEPT**

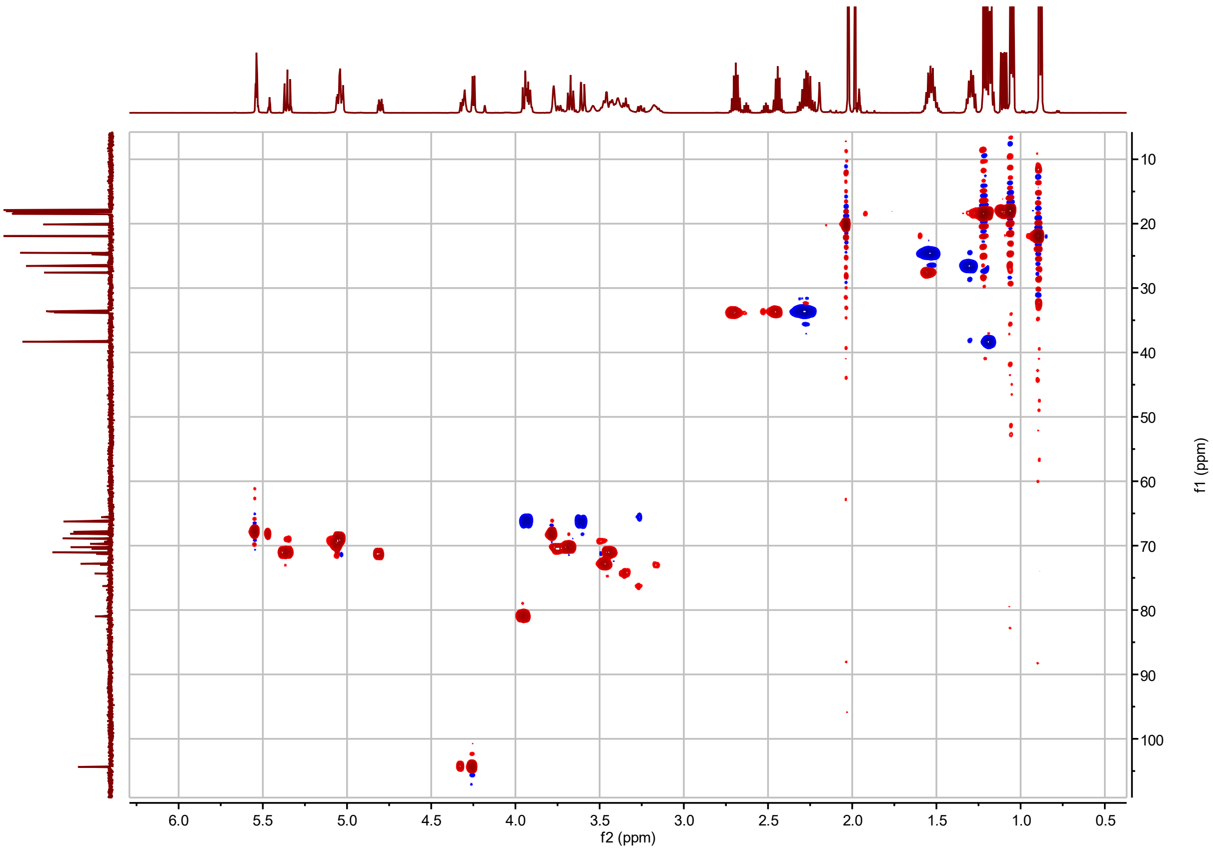


**AI4:18(2,4,4,8) $^{1}H$-$^{13}C$ HSQC**

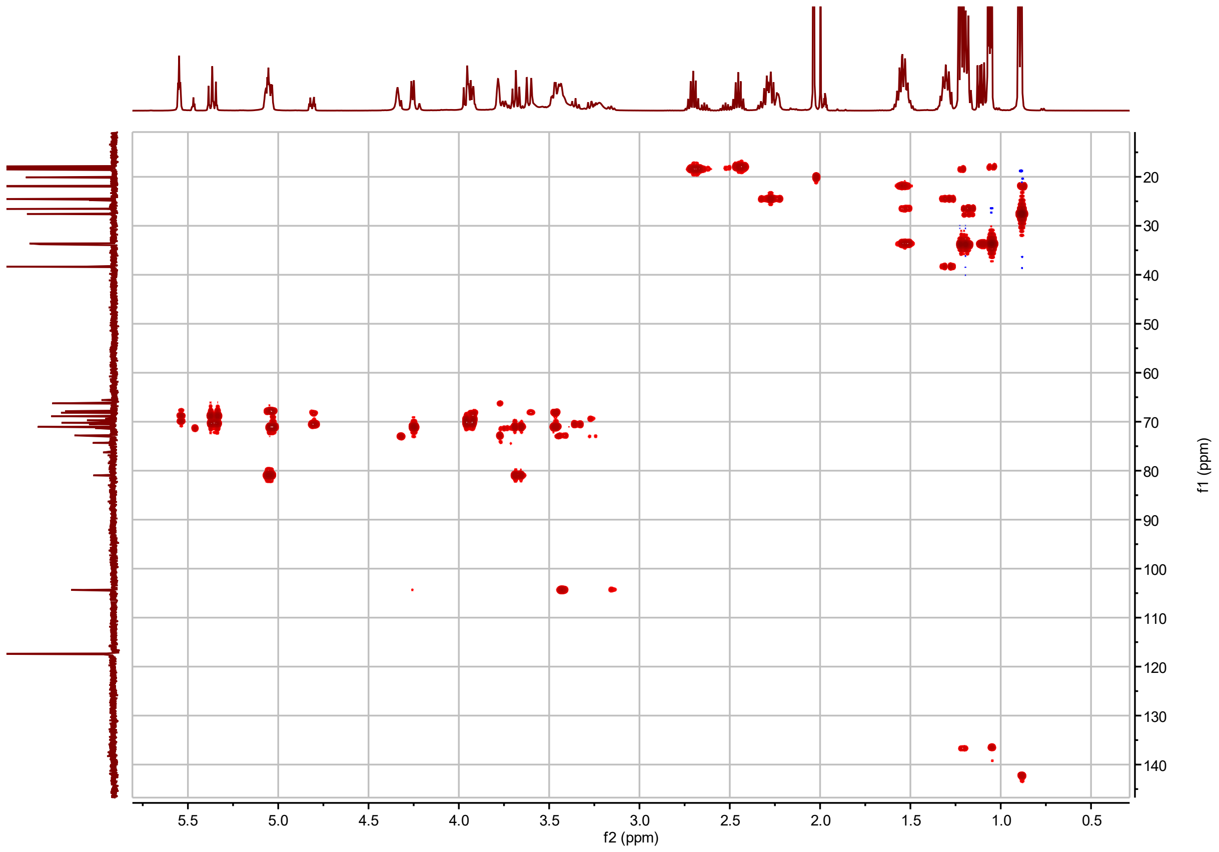


**AI4:18(2,4,4,8) $^{1}$H-$^{13}$C H2BC**

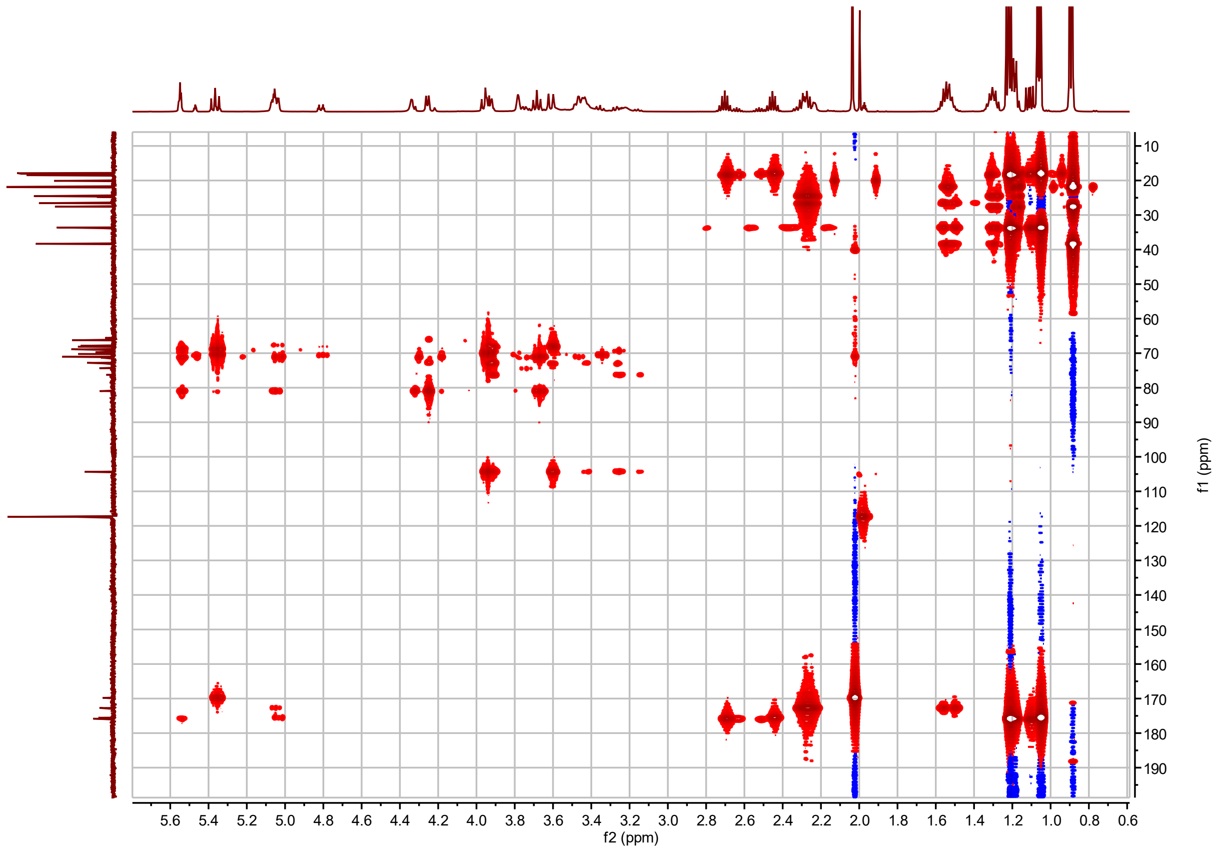


**AI4:18(2,4,4,8) $^{1}H$-$^{13}C$ HMBC**

**I3:18(i4,i4,i10) Chemical shifts and coupling constants.**

| | |
|---|---|
| 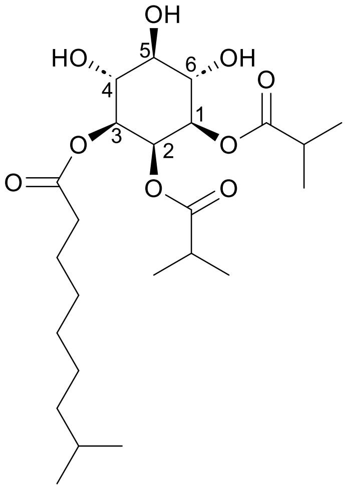 | I3:18(i4,i4,i10)<br><br>Molecular Formula: $C_{24}H_{42}O_9$<br><br>Instrument: Agilent 500 MHz DDR2<br><br>NMR solvent: $CD_3CN$<br><br>Fractions: 71-74<br><br>InChI Key: IRUZHFMTUQKAJJ-OGVIVNNGSA-N<br><br>SMILES:<br>CC(C)CCCCCCC(=O)O[C@H]1[C@H](O)[C@@H](O)[C@H](O)[C@@H](OC(=O)C(C)C)[C@H]1OC(=O)C(C)C |

| Carbon # (Group) | $^{1}$H ($\delta$, ppm) | $^{13}$C ($\delta$, ppm) |
|---|---|---|
| 1(CH) | 4.80 (dd, *J* = 2.95, 7.15 Hz) | 71.31 or 71.33[a] |
| -1(CO) | | 176.01 |
| -2(CH) | 2.52 (hept, *J* = 6.98 Hz) | 33.68 or 33.70[a] |
| -3,4($CH_3$) | 1.09, 1.11 (d, *J* = 7.01 Hz) | 18.01, 18.24 |
| 2(CH) | 5.47 (t, *J* = 2.98 Hz) | 68.16 |
| -1(CO) | | 179.89 |
| -2(CH) | 2.64 (hept, *J* = 6.97 Hz) | 33.85 |
| -3,4($CH_3$) | 1.18 (d, *J* = 6.95 Hz) | 18.35, 18.42 |
| 3(CH) | 4.80 (dd, *J* = 2.95, 7.15 Hz) | 71.31 or 71.33[a] |
| -1(CO) | | 172.80 |
| -2($CH_2$) | 2.26 (m) | 33.68 or 33.70[a] |
| -3($CH_2$) | 1.56 (m) | 24.55 |
| -4-6($CH_2$) | 1.36-1.25 (m) | 29.27(5), 28.73(4), 26.96(6) |
| -7($CH_2$) | 1.18 (m) | 38.70 |
| -8(CH) | 1.54 (hept, *J* = 6.7 Hz) | 27.74 |
| -9,10($CH_3$) | 0.89 (d, *J* = 6.6 Hz) | 21.93 |
| 4(CH) | 3.75 (t, *J* = 9.67 Hz) | 70.49 or 70.51[a] |
| 5(CH) | 3.34 (t, *J* = 9.29 Hz) | 70.49 or 70.51[a] |
| 6(CH) | 3.73 (t, *J* = 9.67 Hz) | 74.33 |

a – $^{13}$C signals not resolved in 2D spectra.

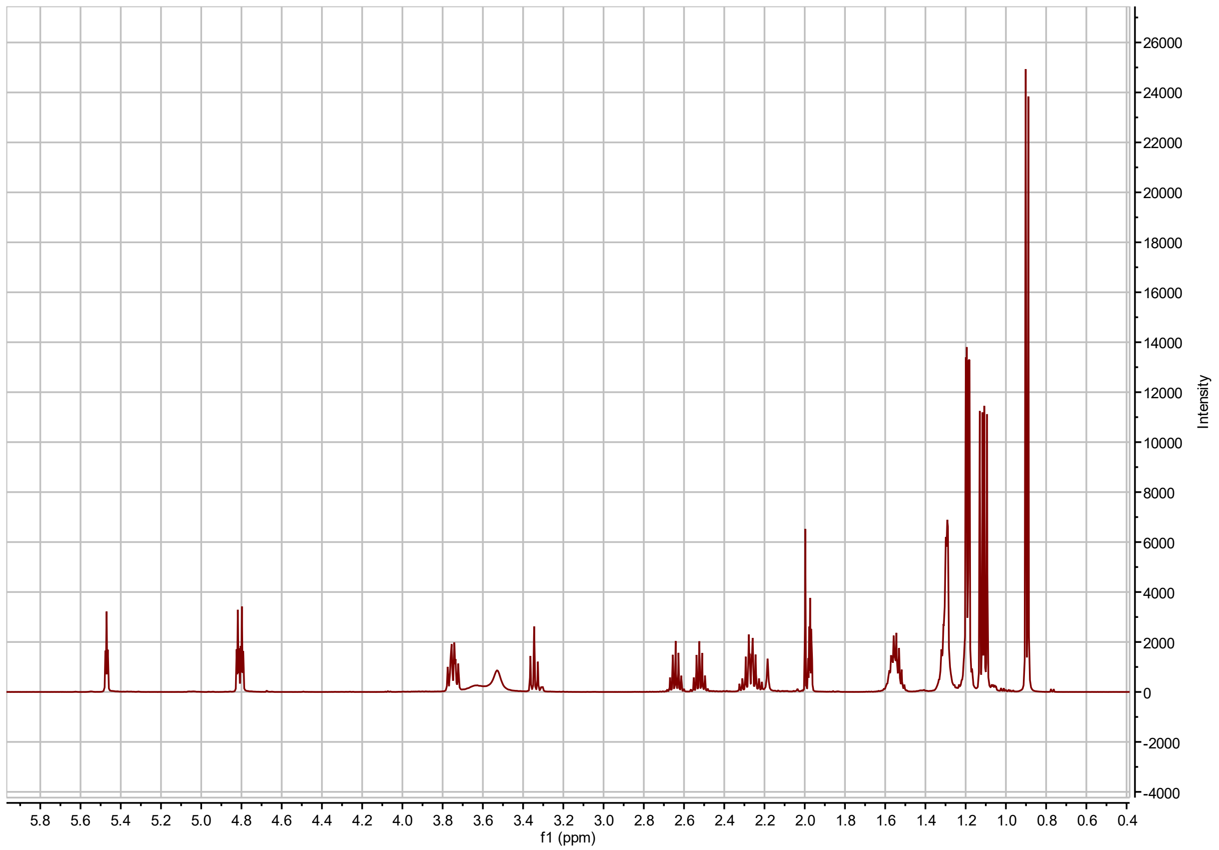


**I3:18(i4,i4,i10) $^{1}$H NMR**

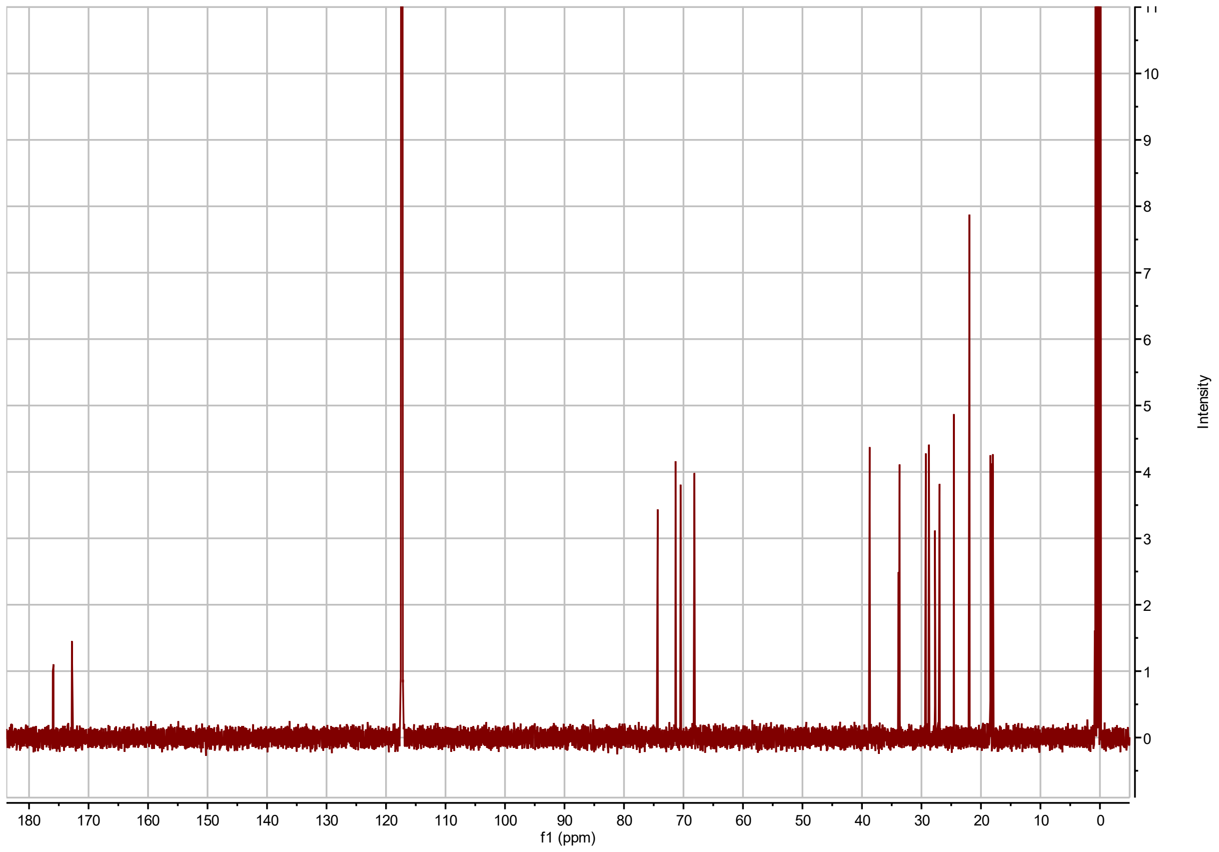


**I3:18(i4,i4,i10) $^{13}$C NMR**

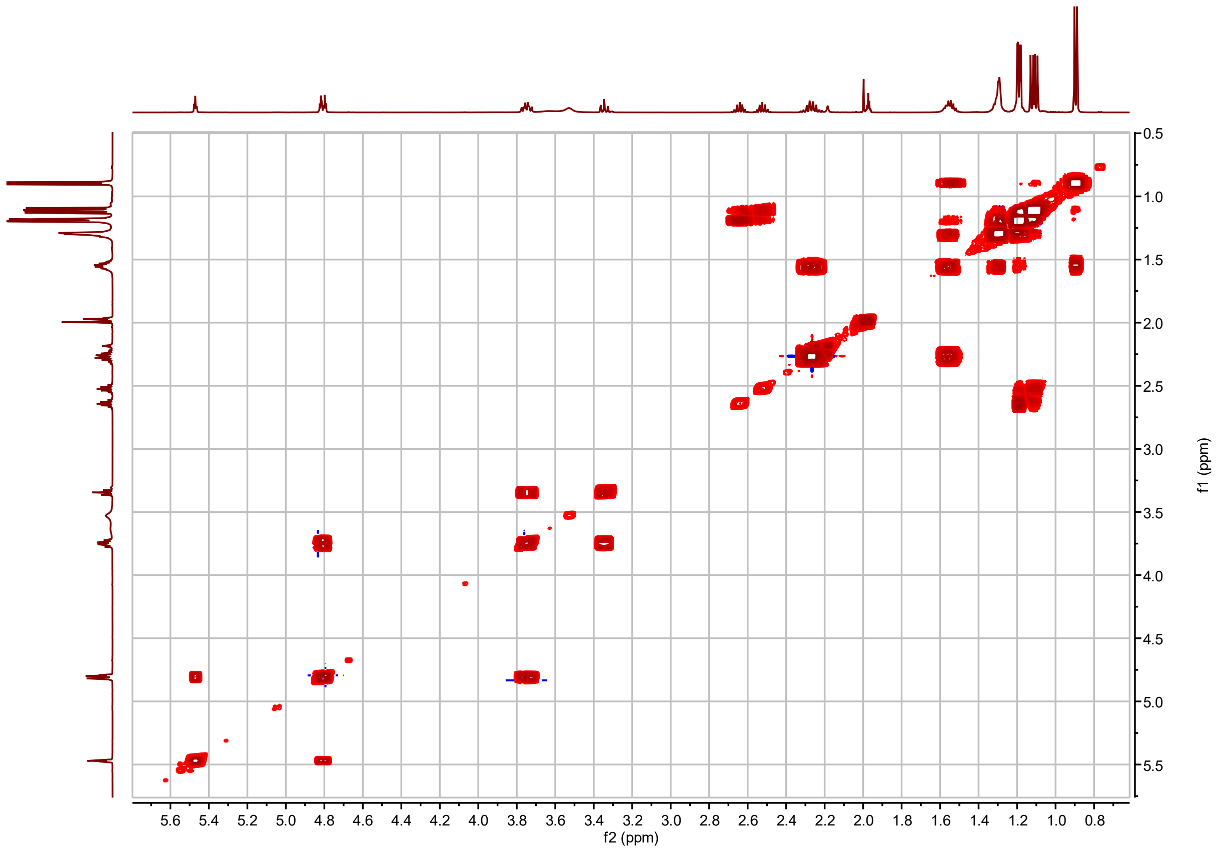


**I3:18(i4,i4,i10) $^{1}$H-$^{1}$H COSY**

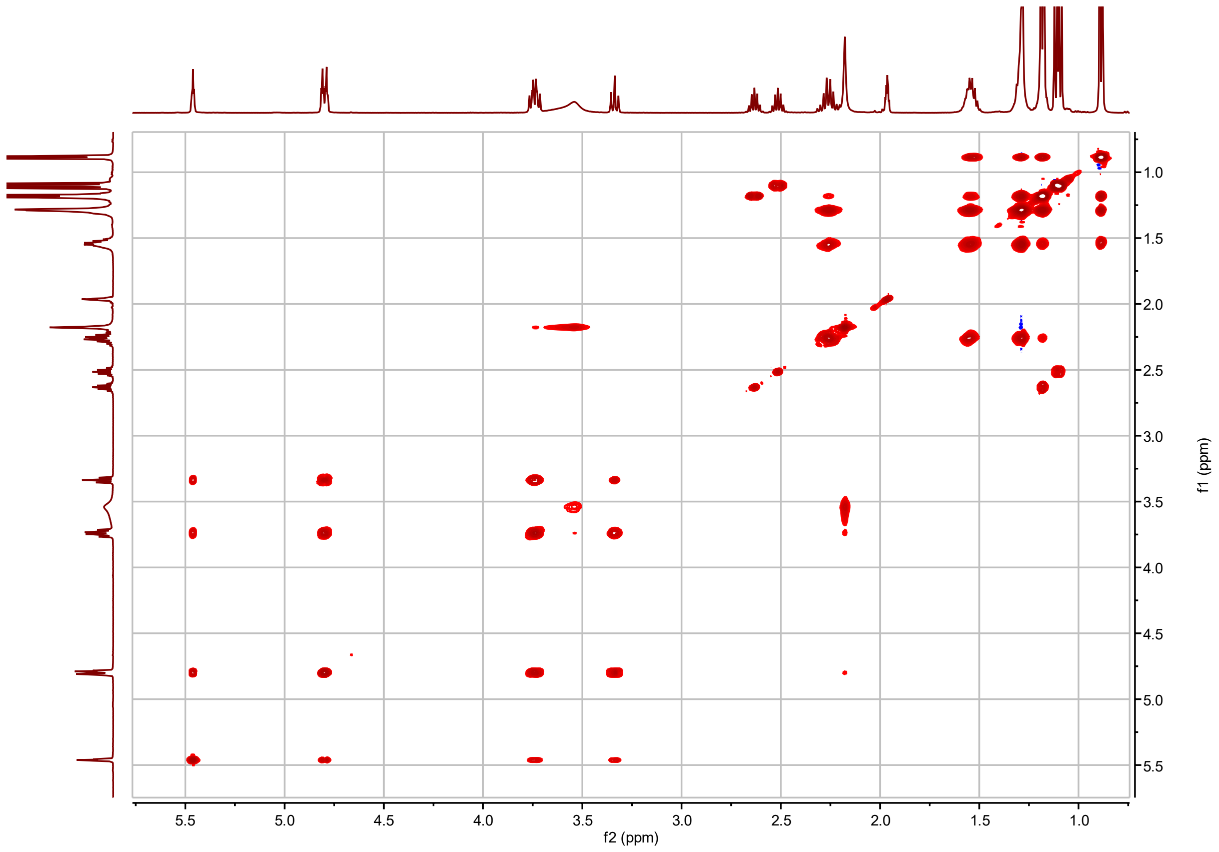


**I3:18(i4,i4,i10) $^{1}$H-$^{1}$H TOCSY**

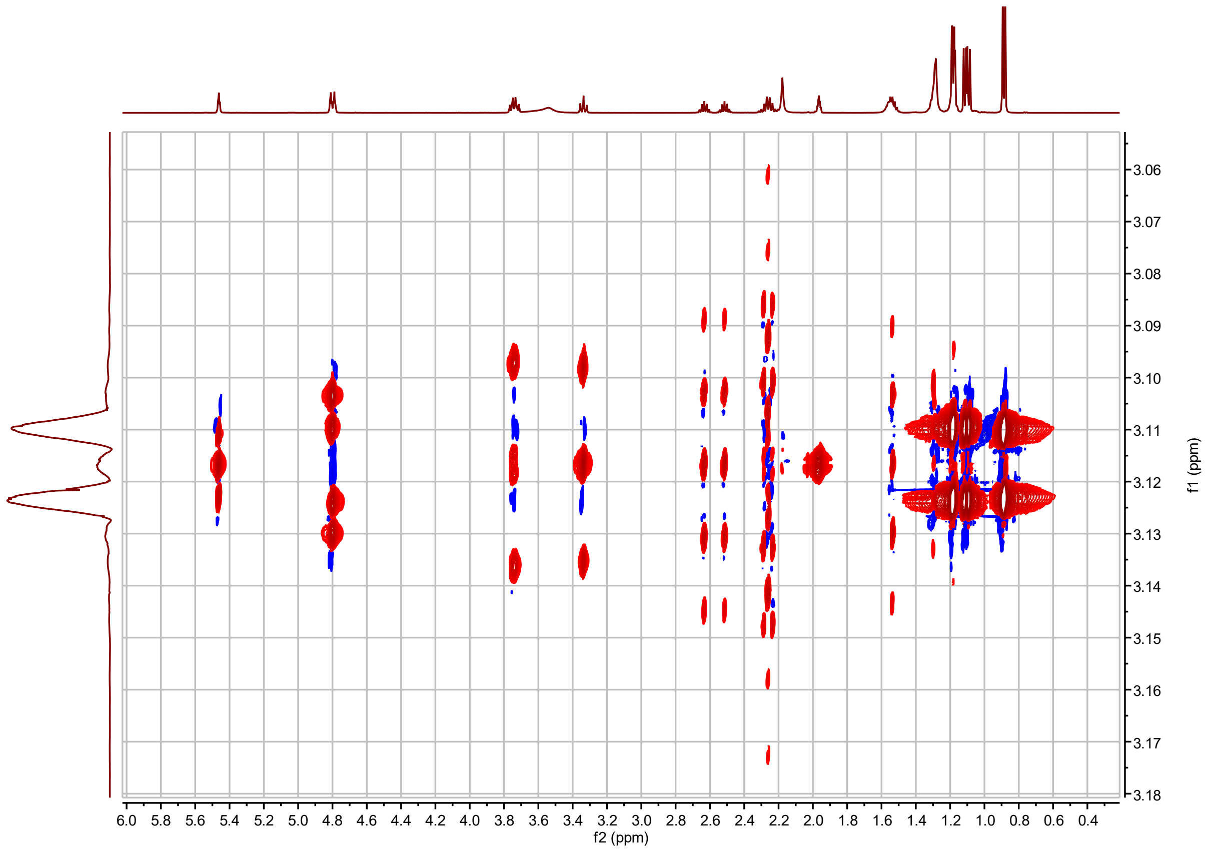


**I3:18(i4,i4,i10) *J*-resolved**

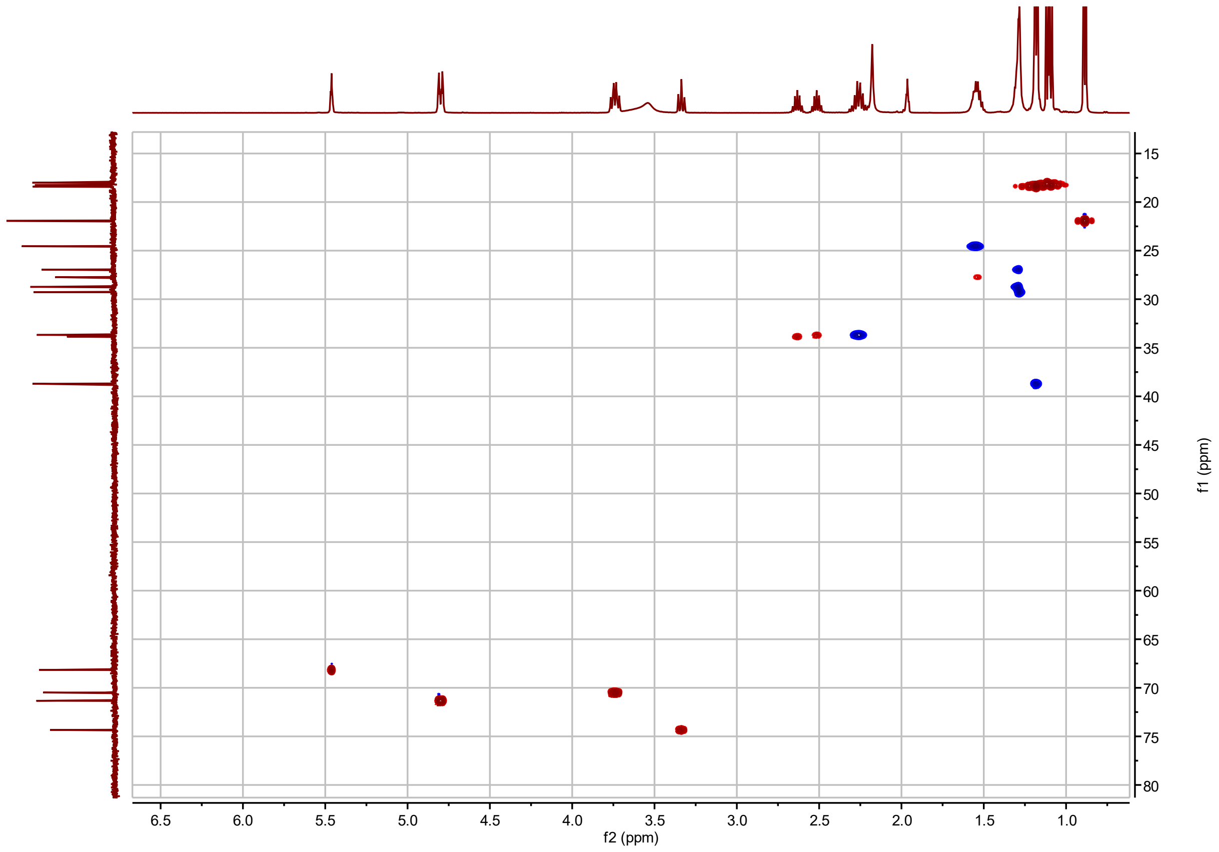


**I3:18(i4,i4,i10) $^{1}H$-$^{13}C$ HSQC**

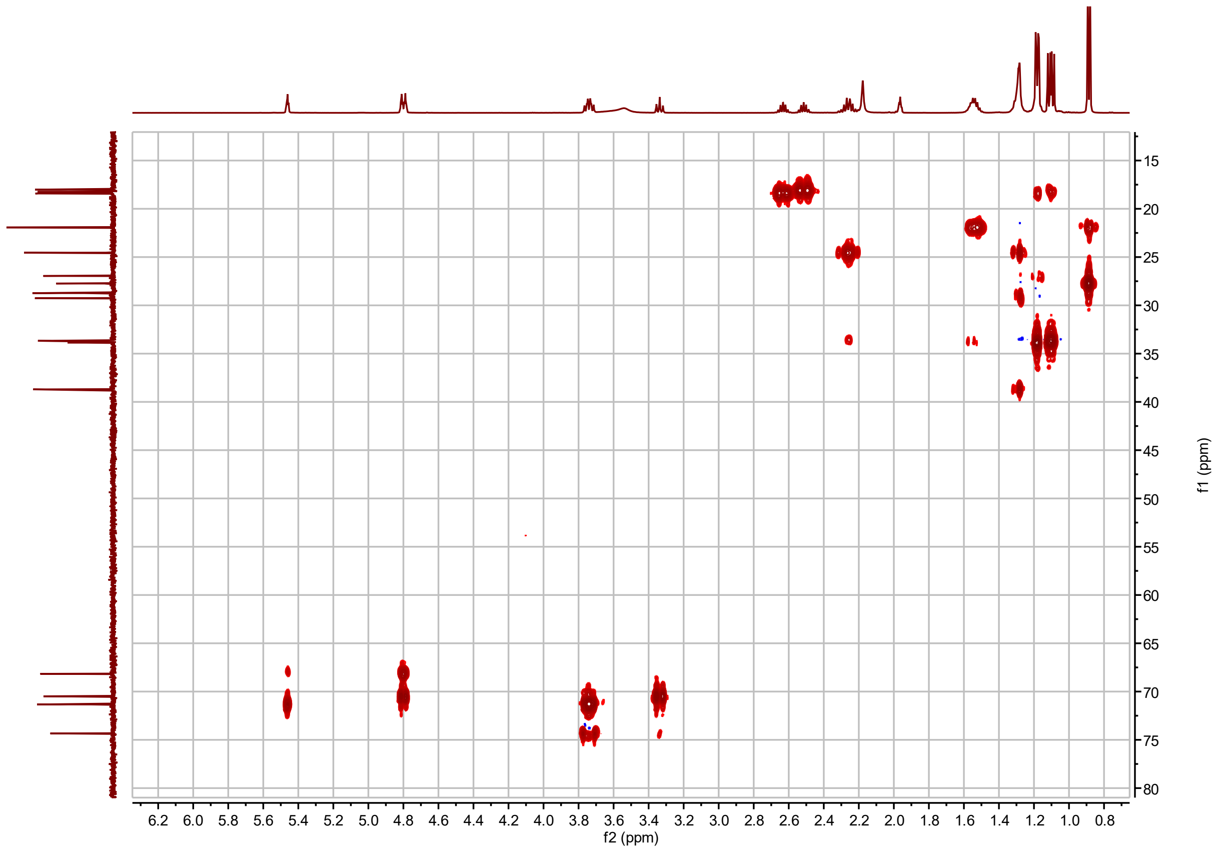


**I3:18(i4,i4,i10) $^{1}$H-$^{13}$C H2BC**

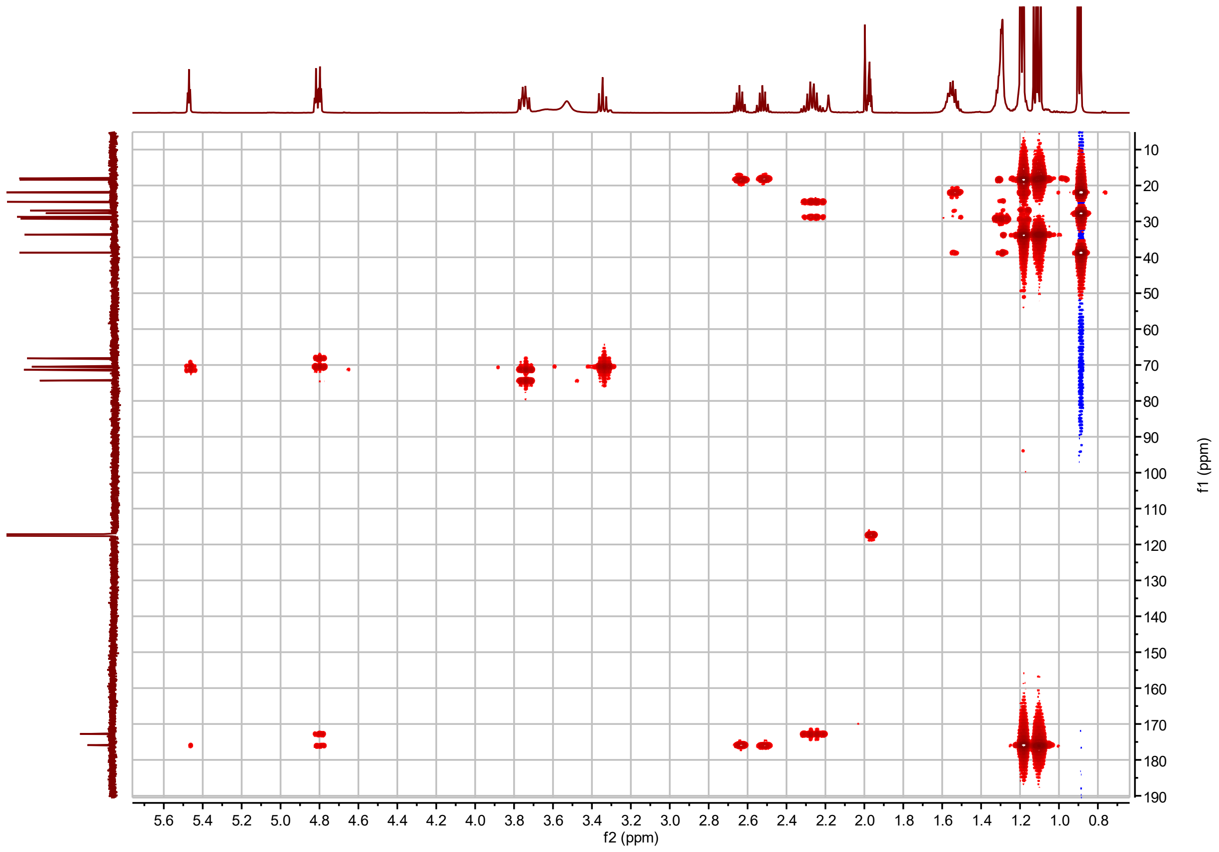


**I3:18(i4,i4,i10) $^{1}$H-$^{13}$C HMBC**

**I3:20(i4,i4,i12) Chemical shifts and coupling constants.**

<table>
<tr><td>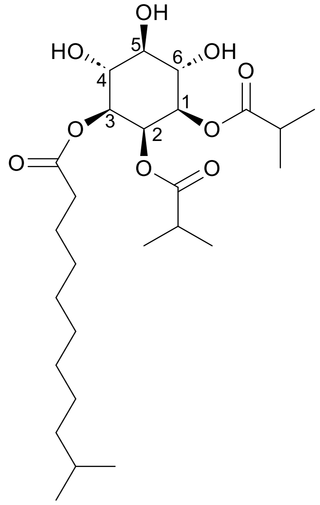</td><td colspan="2">I3:20(i4,i4,i12)<br><br>Molecular Formula: $C_{26}H_{46}O_9$<br><br>Instrument: Agilent 500 MHz DDR2<br><br>NMR solvent: $CD_3CN$<br><br>Fractions: 105-107<br><br>InChI Key: WFZVKWMBYUFKOS-VXAMWORUSA-N<br><br>SMILES:<br>CC(C)CCCCCCCCC(=O)O[C@H]1[C@H](O)[C@@H](O)[C@H](O)[C@@H](OC(=O)C(C)C)[C@H]1OC(=O)C(C)C</td></tr>
<tr><td>Carbon # (Group)</td><td>$^{1}$H ($\delta$, ppm)</td><td>$^{13}$C ($\delta$, ppm)</td></tr>
<tr><td>1(CH)<br>-1(CO)<br>-2(CH)<br>-3,4($CH_3$)</td><td>4.80 or 4.79 (dd, $J$ = 3.1, 10.1 Hz)<br><br>2.51 (hept, $J$ = 6.92 Hz)<br>1.11, 1.09 (d, $J$ = 7.0 Hz)</td><td>71.33 or 71.31[a]<br>157.99<br>33.69 or 33.67[a]<br>18.21, 17.99</td></tr>
<tr><td>2(CH)<br>-1(CO)<br>-2(CH)<br>-3,4($CH_3$)</td><td>5.46 (t, $J$ = 3.01 Hz)<br><br>2.62 (hept, $J$ = 6.95 Hz)<br>1.18 (d, $J$ = 7.0 Hz)</td><td>68.18<br>175.87<br>33.84<br>18.39, 18.32</td></tr>
<tr><td>3(CH)<br>-1(CO)<br>-2($CH_2$)<br>-3($CH_2$)<br>-4-8($CH_2$)<br><br>-9($CH_2$)<br>-10(CH)<br>-11,12($CH_3$)</td><td>4.80 or 4.79 (dd, $J$ = 3.1, 10.1 Hz)<br><br>2.26 (m)<br>1.54 (m)<br>1.33-1.27 (m)<br><br>1.19 (m)<br>1.54 (m)<br>0.88 (d, $J$ = 6.71 Hz)</td><td>71.33 or 71.31[a]<br>172.77<br>33.69 or 33.67[a]<br>24.53<br>28.68(4), 28.97(5), 29.18(6), 29.57 (7), 27.10(8)<br>38.76<br>27.73<br>21.91</td></tr>
<tr><td>4(CH)</td><td>3.75 or 3.73 (t, $J$ = 9.9 Hz)</td><td>70.53 or 70.51[a]</td></tr>
<tr><td>5(CH)</td><td>3.33 (t, $J$ = 9.30 Hz)</td><td>74.36</td></tr>
<tr><td>6(CH)</td><td>3.75 or 3.73 (t, $J$ = 9.9 Hz)</td><td>70.53 or 70.51[a]</td></tr>
</table>

a – $^{13}$C signals not resolved in 2D spectra.

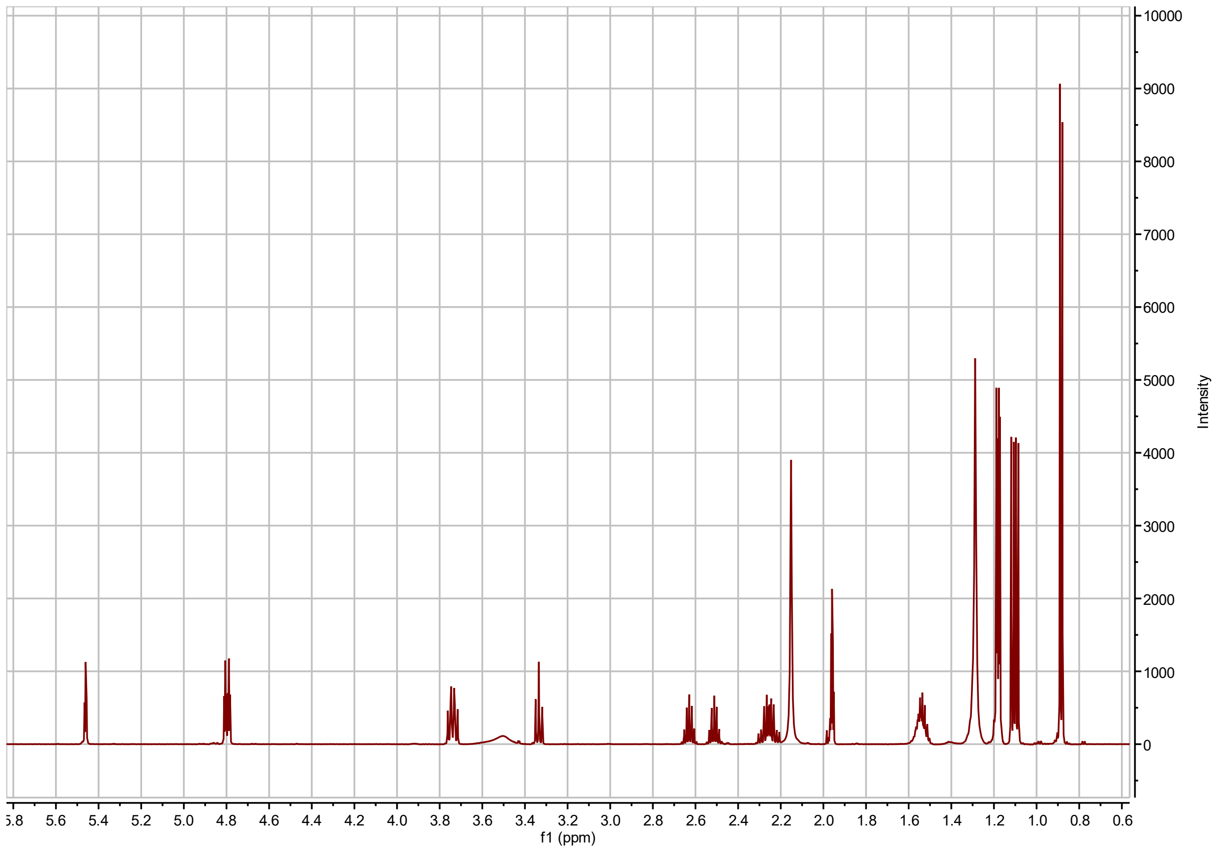


**I3:20(i4,i4,i12) $^{1}$H NMR**

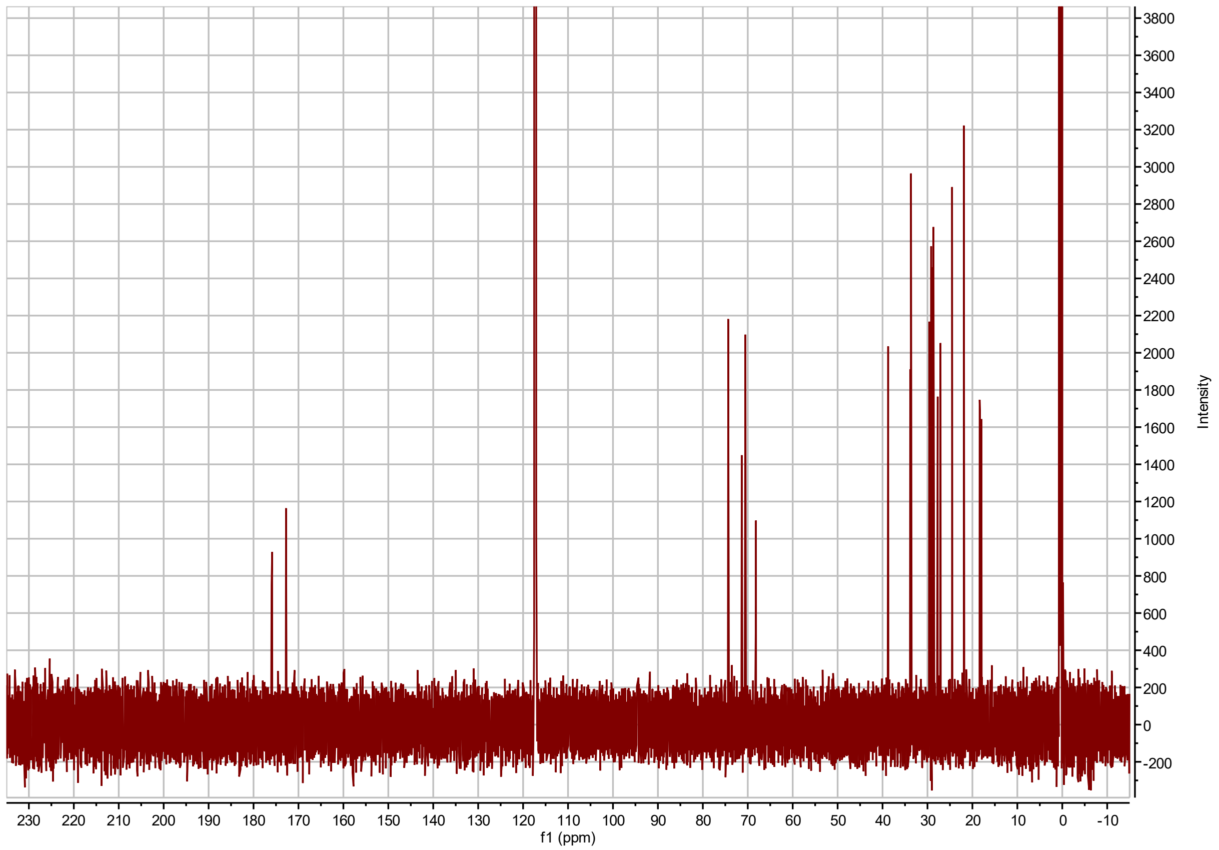


**I3:20(i4,i4,i12) $^{13}$C NMR**

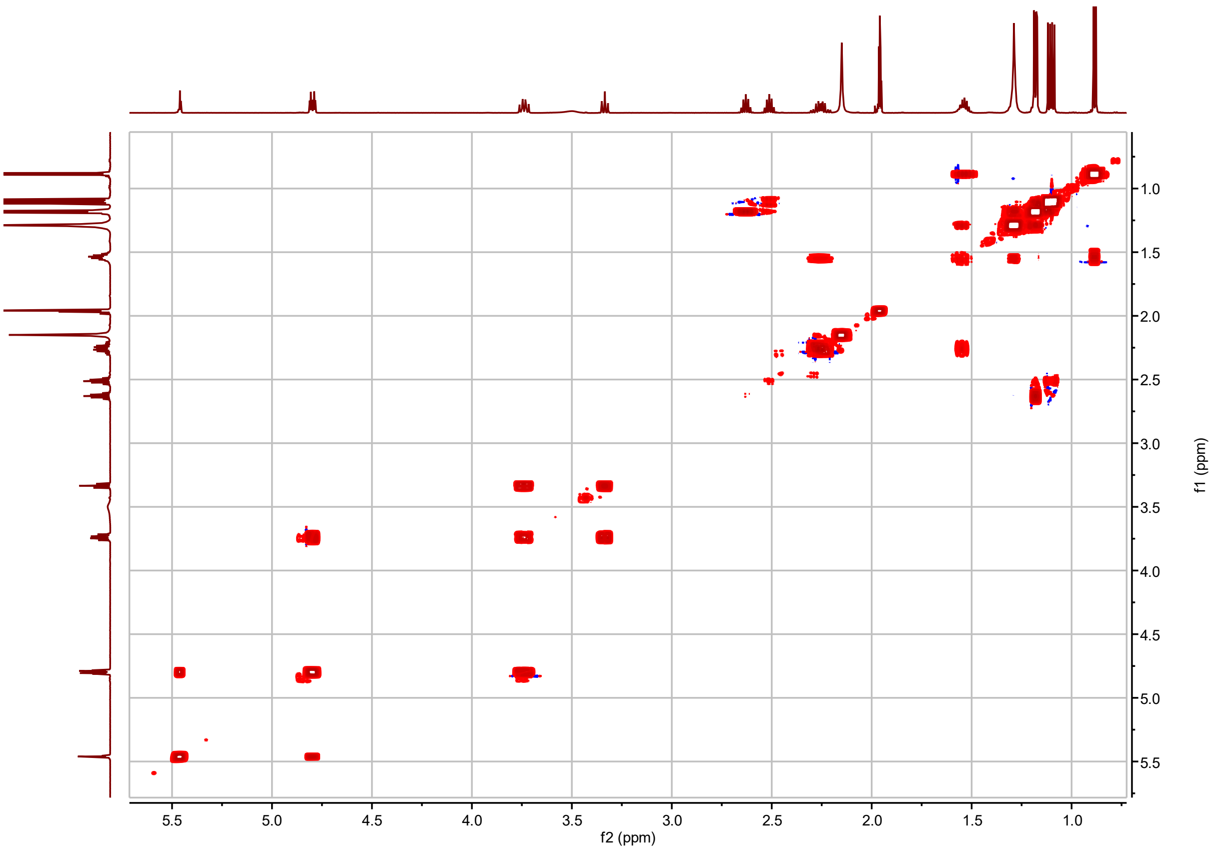


**I3:20(i4,i4,i12) $^{1}H$-$^{1}H$ COSY**

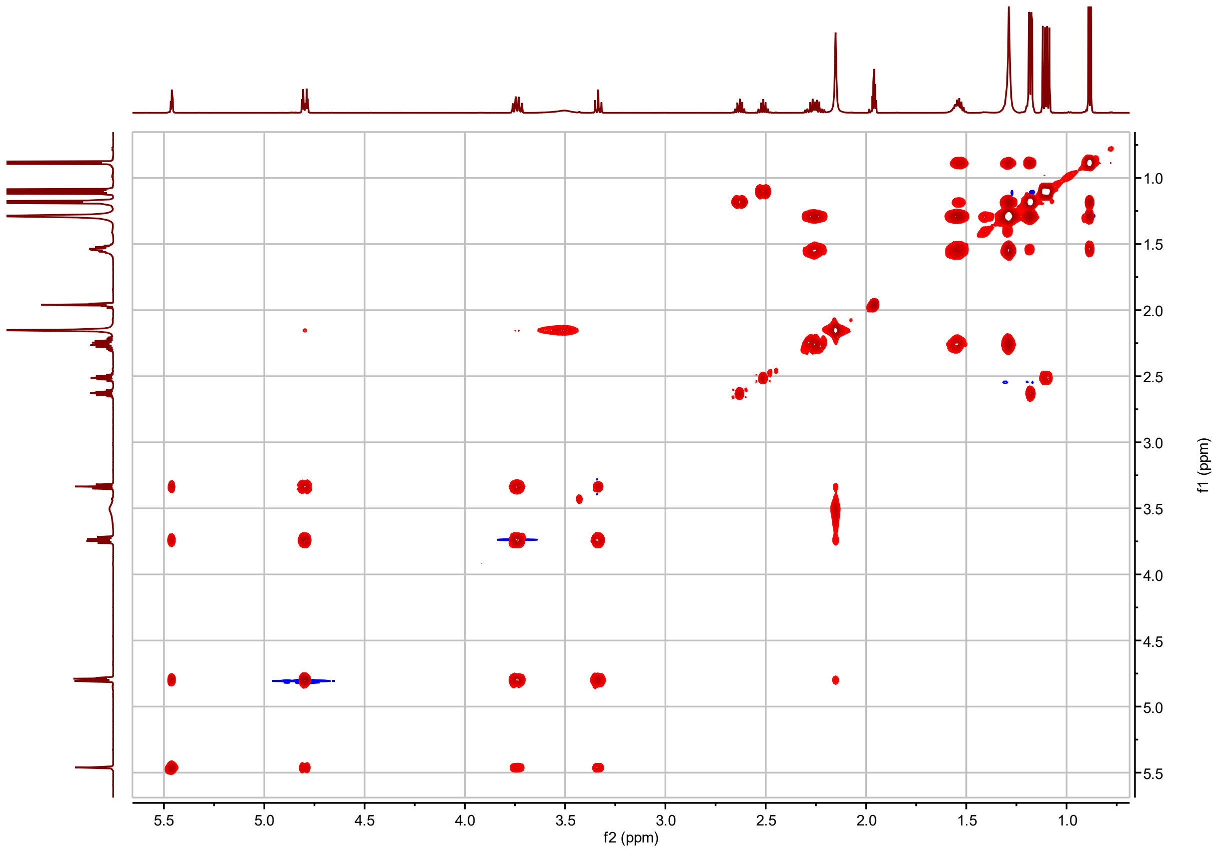


**I3:20(i4,i4,i12) $^{1}$H-$^{1}$H TOCSY**

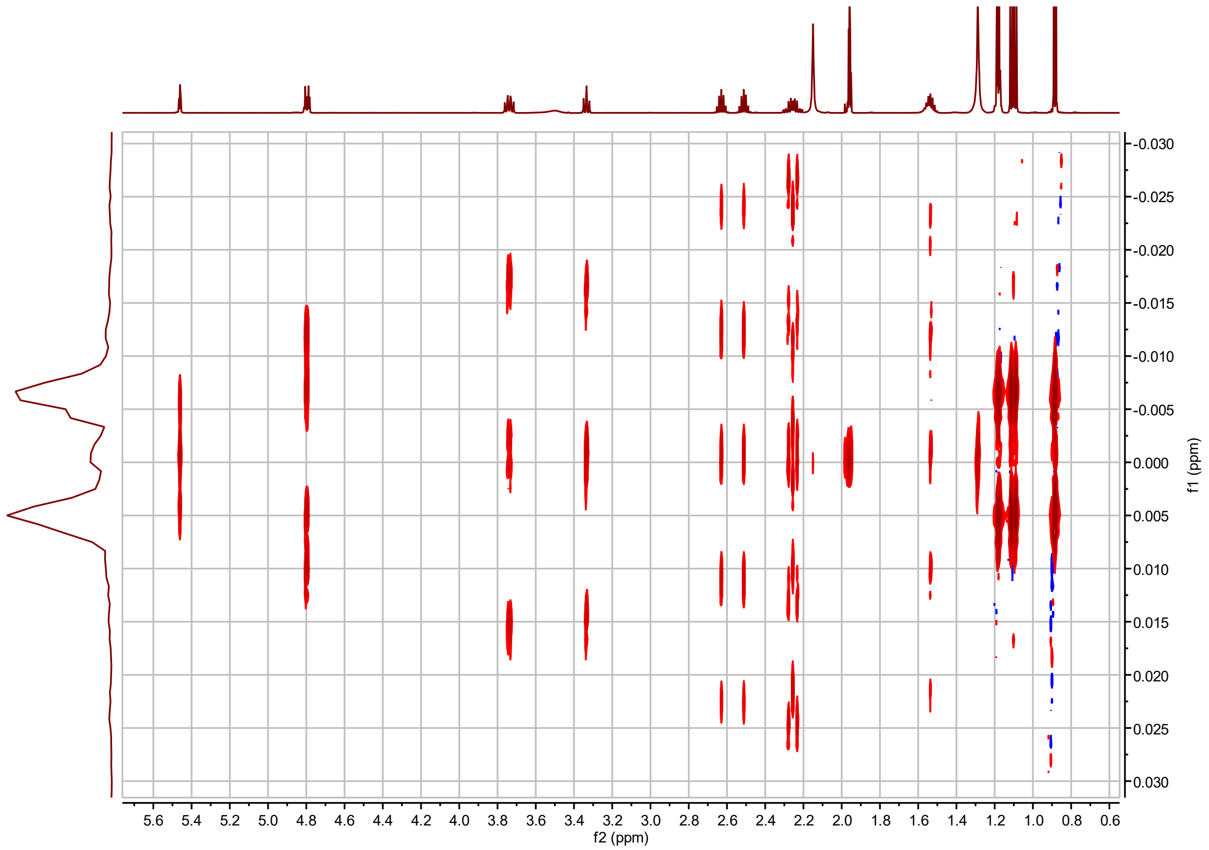


**I3:20(i4,i4,i12) *J*-resolved**

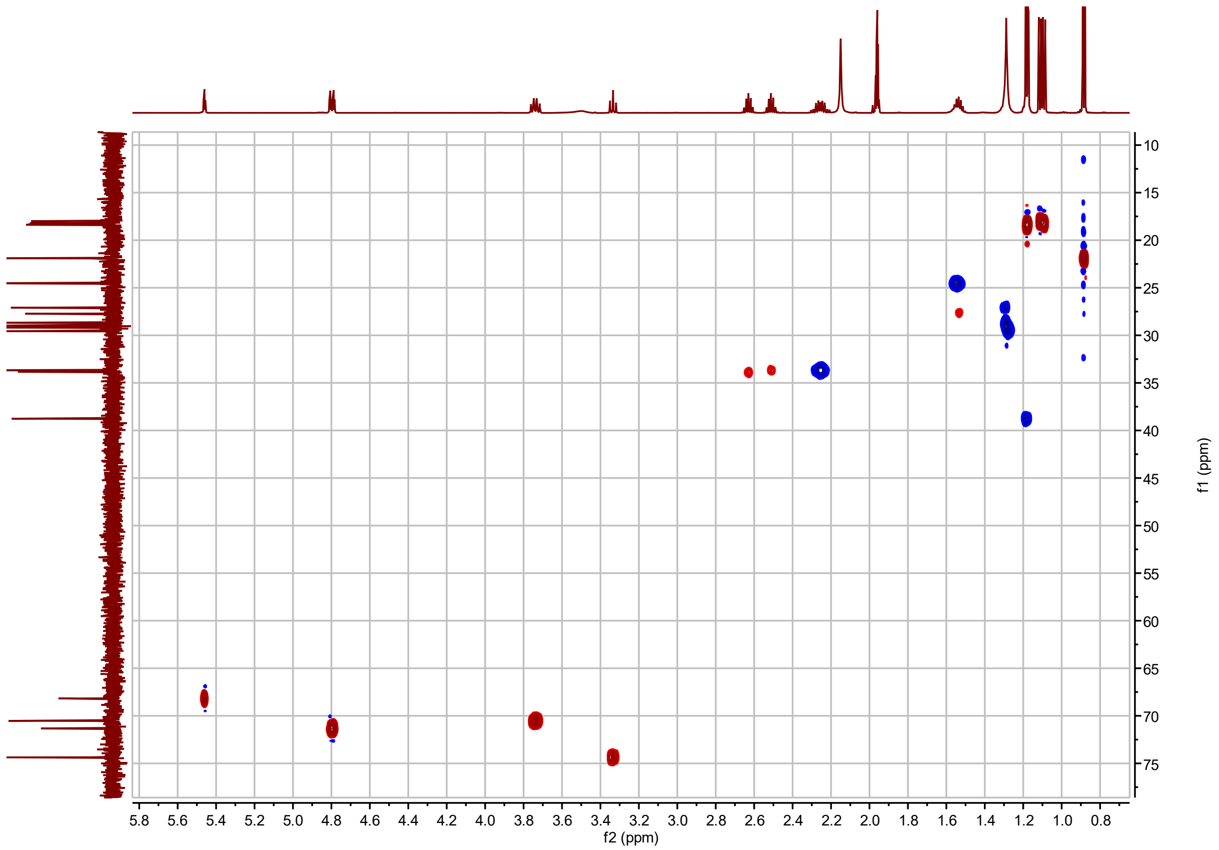


**I3:20(i4,i4,i12) $^{1}$H-$^{13}$C HSQC**

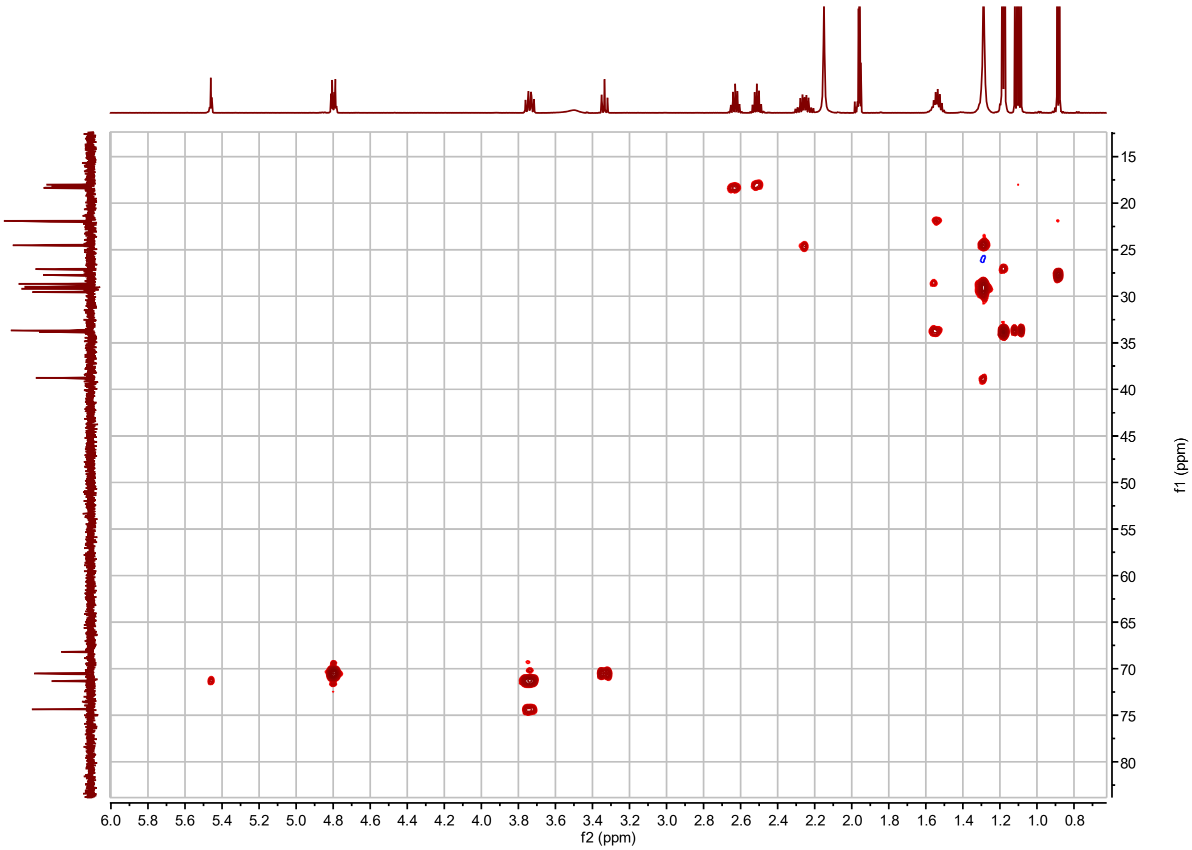


**I3:20(i4,i4,i12) $^{1}H$-$^{13}C$ H2BC**

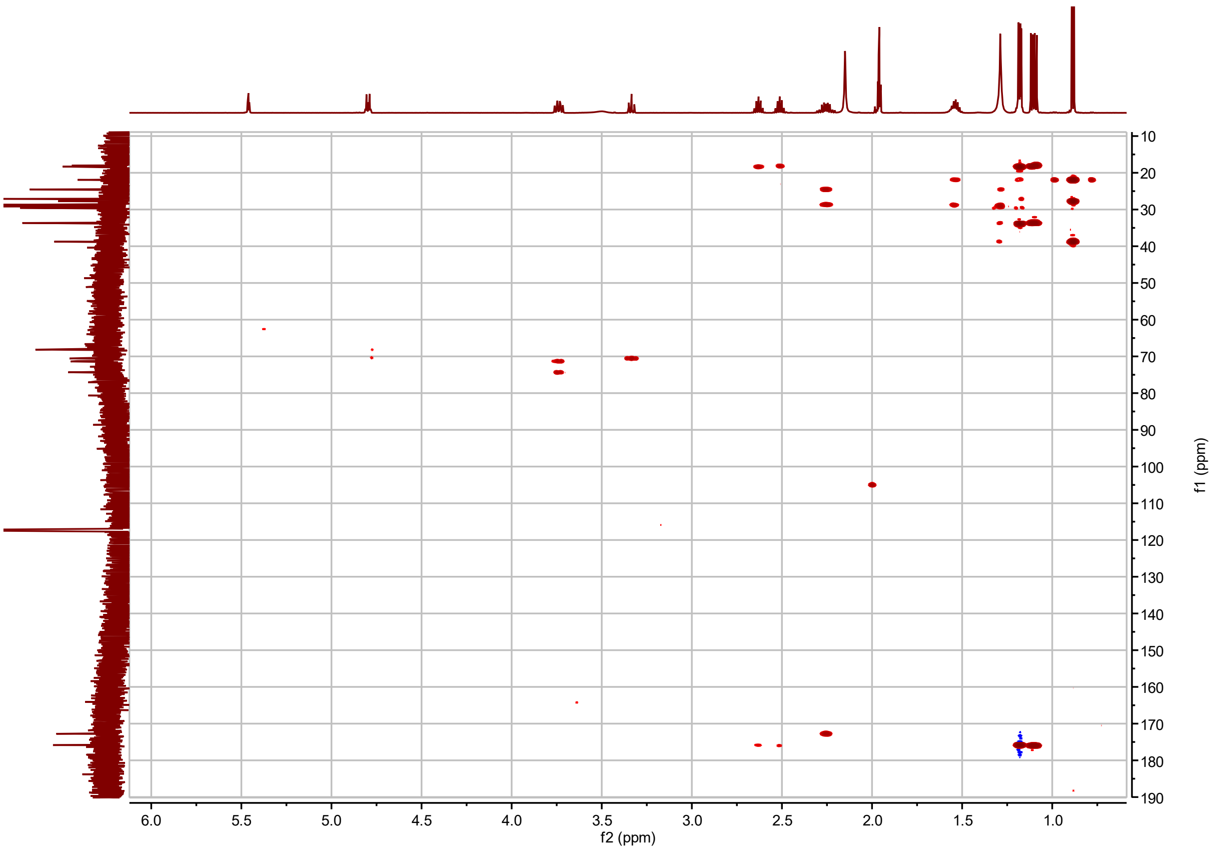


**I3:20(i4,i4,i12) $^{1}H$-$^{13}C$ HMBC with apodization optimized for correlations between acyl chain carbonyl carbon and C2 proton(s)**

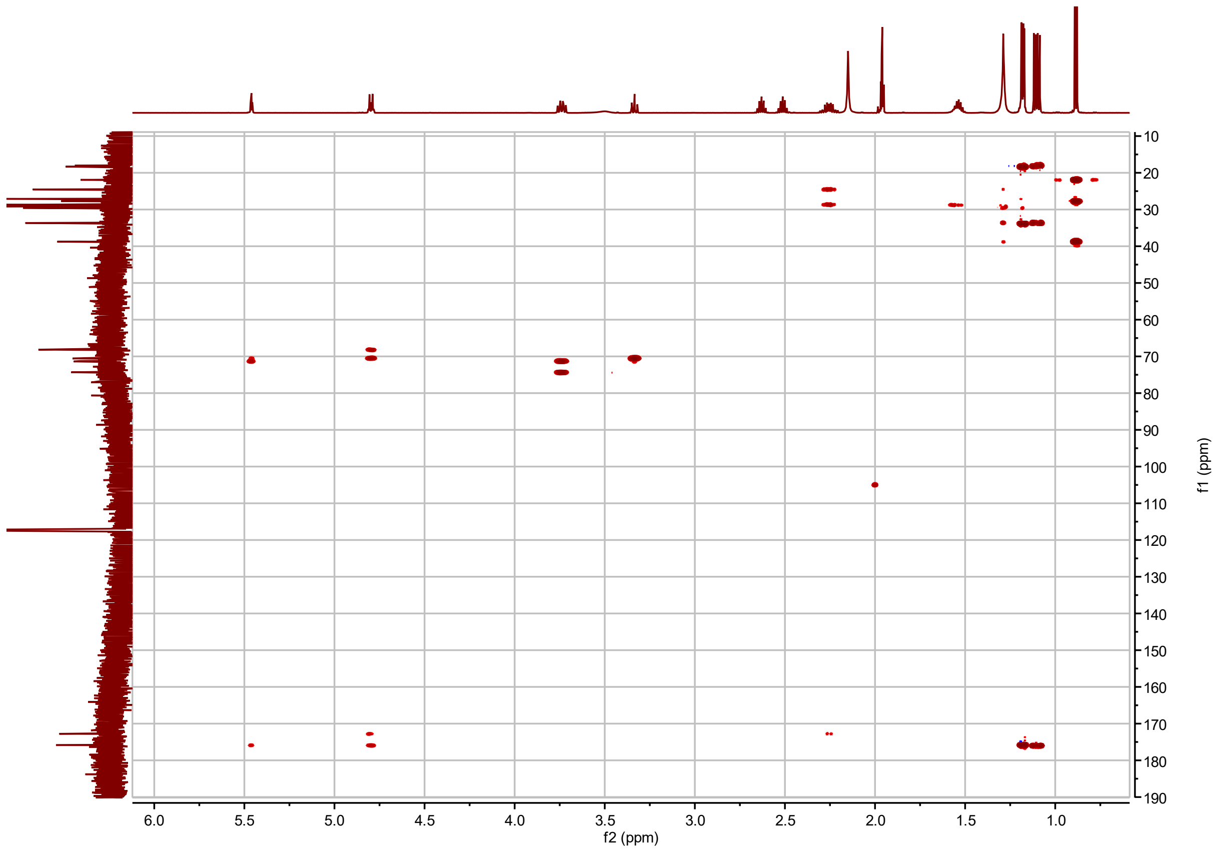


**I3:20(i4,i4,i12) $^{1}$H-$^{13}$C HMBC with apodization optimized for correlations between acyl chain carbonyl carbon and sugar ring protons**

**I3:22(i4,i4,i14) Chemical shifts and coupling constants.**

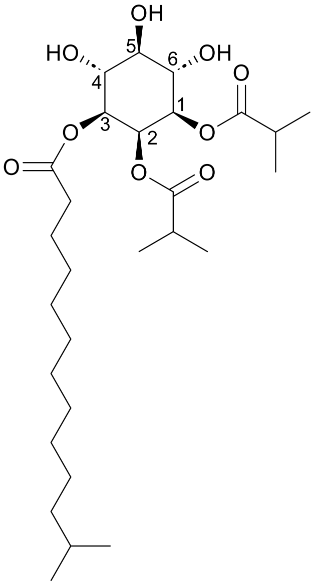


I3:22(i4,i4,i14)

Molecular Formula: $C_{28}H_{50}O_9$

Instrument: Varian 600 MHz Inova, Bruker Avance NEO 600 MHz NMR, Bruker Avance NEO 800 MHz

NMR solvent: $CD_3CN$

Fractions: 139-141

InChI Key: FPKCJTQXNJKTGE-RMLCQDLPSA-N

SMILES: CC(C)CCCCCCCCCCC(=O)O[C@H]1[C@H](O)[C@@H](O)[C@H](O)[C@@H](OC(=O)C(C)C)[C@H]1OC(=O)C(C)C

| Carbon # (Group) | $^{1}$H ($\delta$, ppm) | $^{13}$C ($\delta$, ppm) |
|---|---|---|
| 1(CH) | 4.80 (dt, *J* = 3.25, 10.23 Hz) | 71.34 or 71.31 |
| -1(CO) | | 176.03 or 175.89[a] |
| -2(CH) | 2.52 (hept, *J* = 6.96 Hz) | 33.85 |
| -3,4($CH_3$) | 1.11 (dd, *J* = 6.97, 13.05 Hz) | 18.40 – 18.00 |
| 2(CH) | 5.47 (t, *J* = 2.98 Hz) | 68.18 |
| -1(CO) | | 176.03 or 175.89[a] |
| -2(CH) | 2.64 (hept, *J* = 6.96 Hz) | 33.70 |
| -3,4($CH_3$) | 1.19 (dd, *J* = 2.59, 6.96 Hz) | 18.40 – 18.00 |
| 3(CH) | 4.80 (dt, *J* = 3.25, 10.23 Hz) | 71.34 or 71.31[a] |
| -1(CO) | | 172.81 |
| -2($CH_2$) | 2.27 (m) | 33.67 |
| -3($CH_2$) | 1.55 (m) | 24.53 |
| -4($CH_2$) | 1.29 (m) | 28.68 |
| -5-10($CH_2$) | 1.30 (m) | 29.63, 29.39, 29.36, 29.32, 29.16, 28.98 |
| -11($CH_2$) | 1.19 (m) | 38.78 |
| -12(CH) | 1.54 (m) | 27.74 |
| -13,14($CH_3$) | 0.89 (d, *J* = 6.64 Hz) | 21.92 |
| 4(CH) | 3.74 (td, *J* = 8.24, 9.85, 9.90 Hz) | 70.53 or 70.50[a] |
| 5(CH) | 3.34 (t, *J* = 9.31 Hz) | 74.36 |
| 6(CH) | 3.74 (td, *J* = 8.24, 9.85, 9.90 Hz) | 70.53 or 70.50[a] |

a – $^{13}$C signals not resolved in 2D spectra.

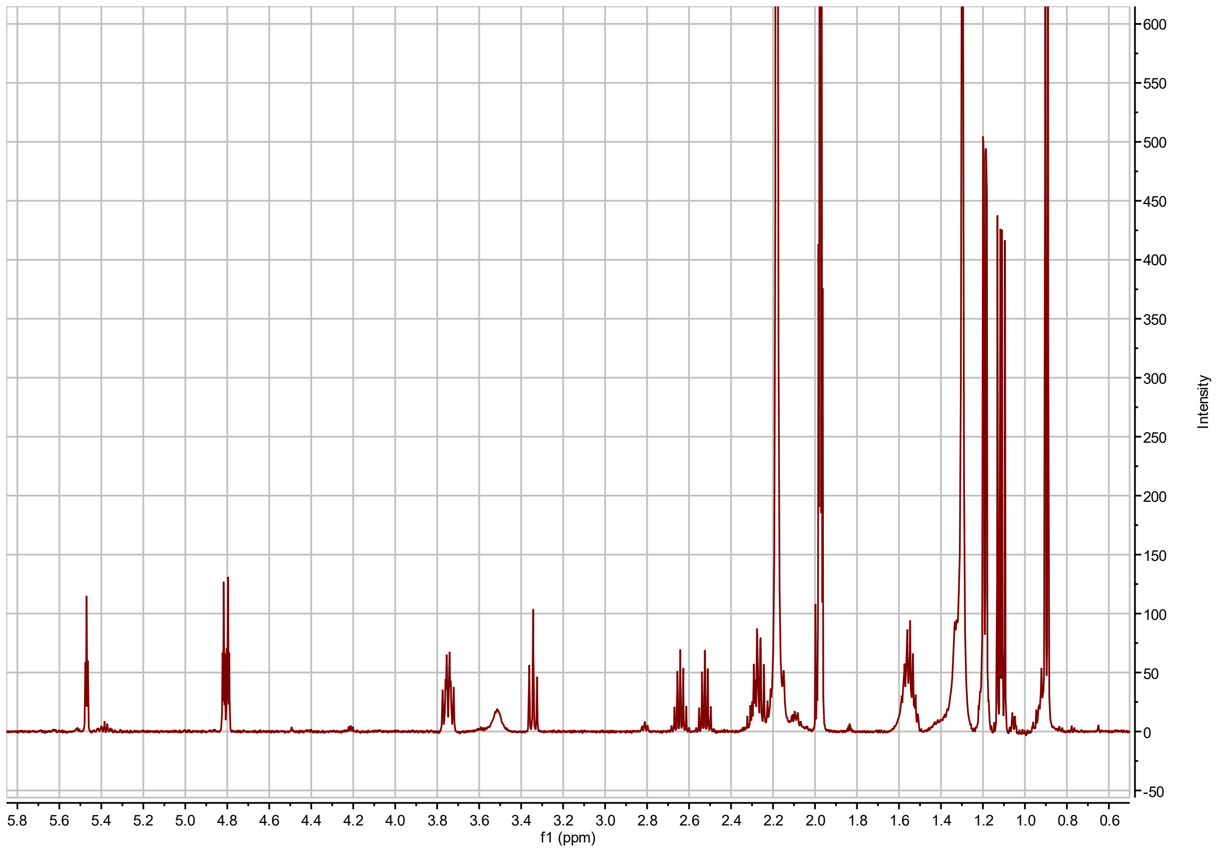


**I3:22(i4,i4,i14) $^1$H NMR**

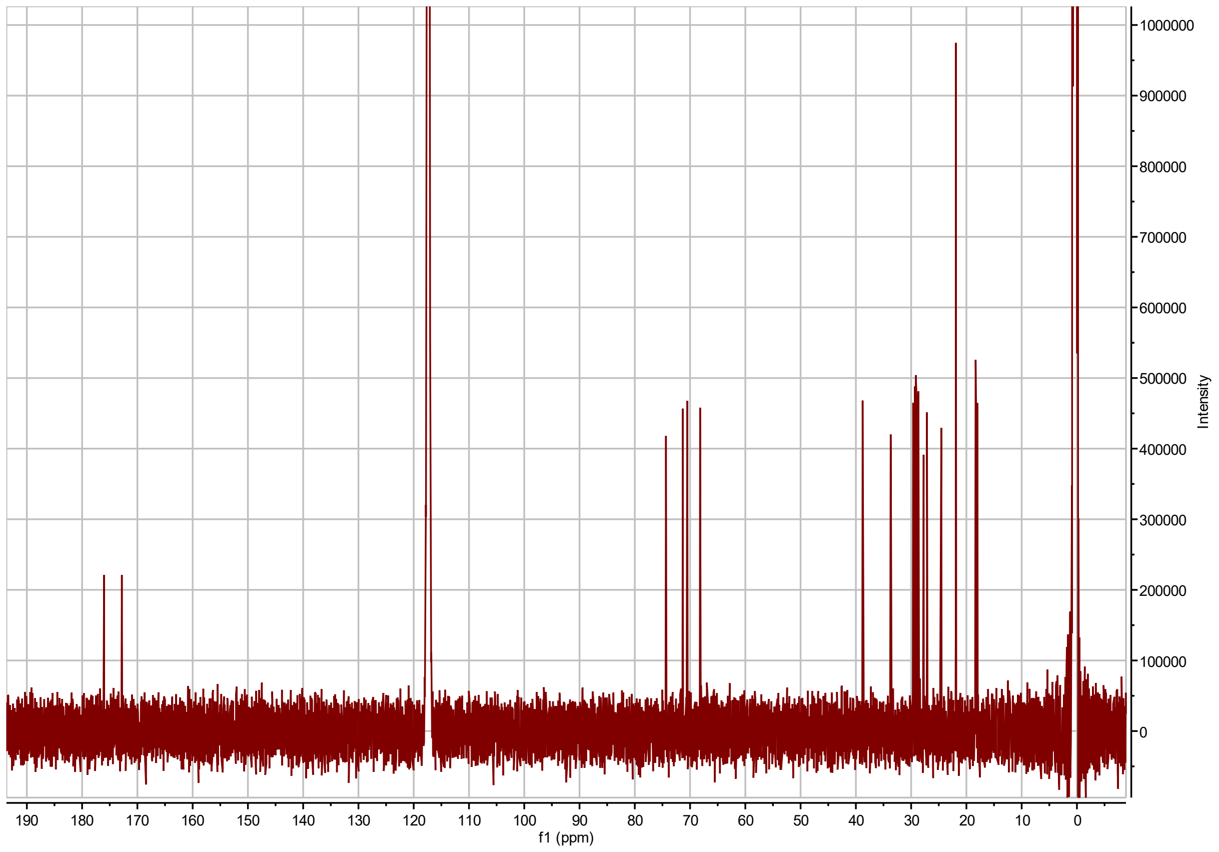


**I3:22(i4,i4,i14) $^{13}$C NMR**

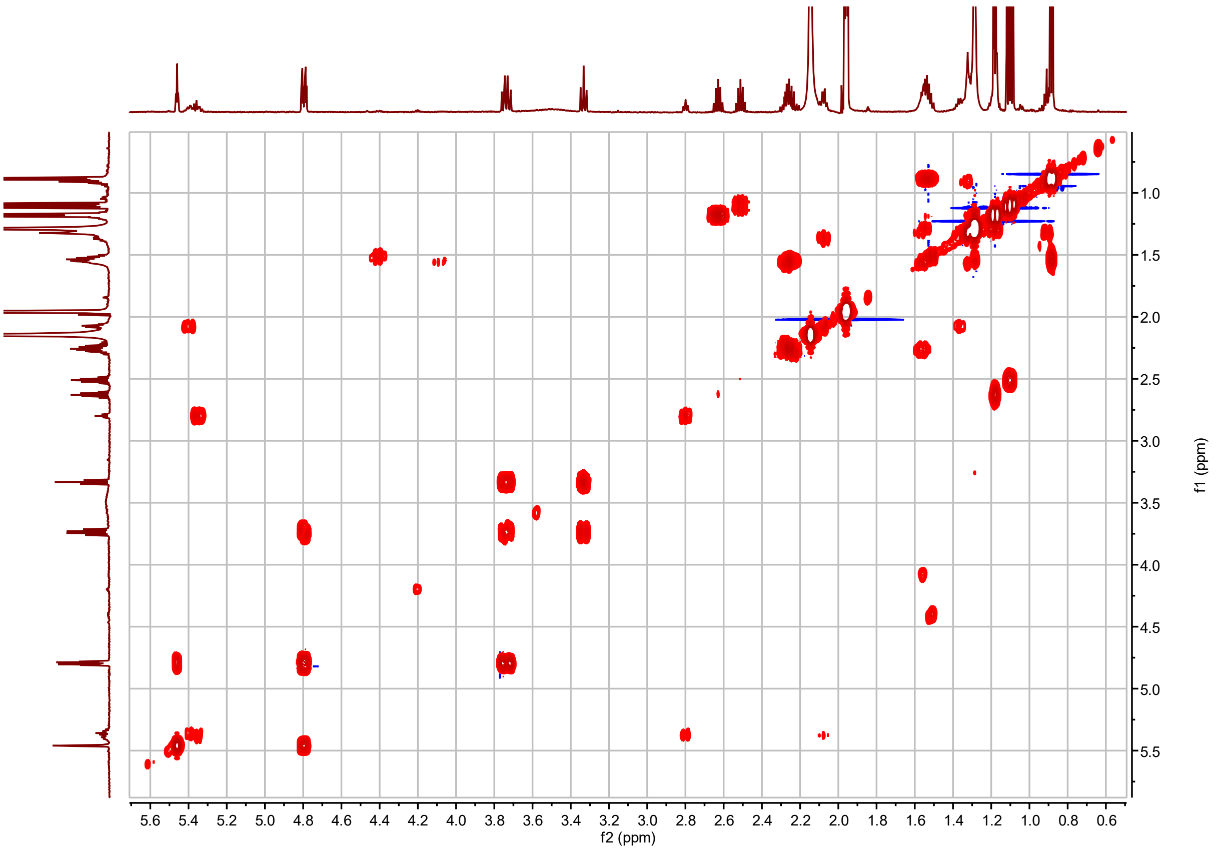


**I3:22(i4,i4,i14) $^1$H-$^1$H COSY**

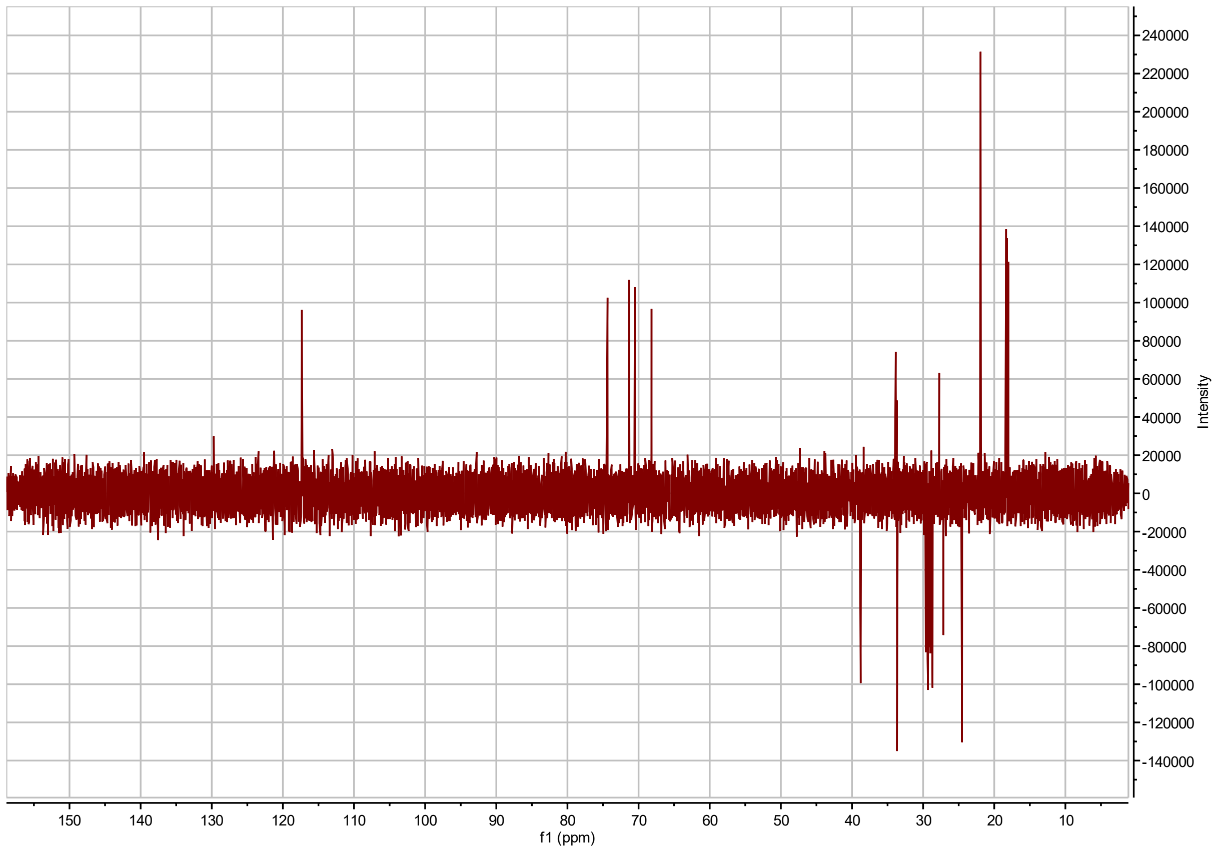


**I3:22(i4,i4,i14) $^{13}$C DEPT-Q**

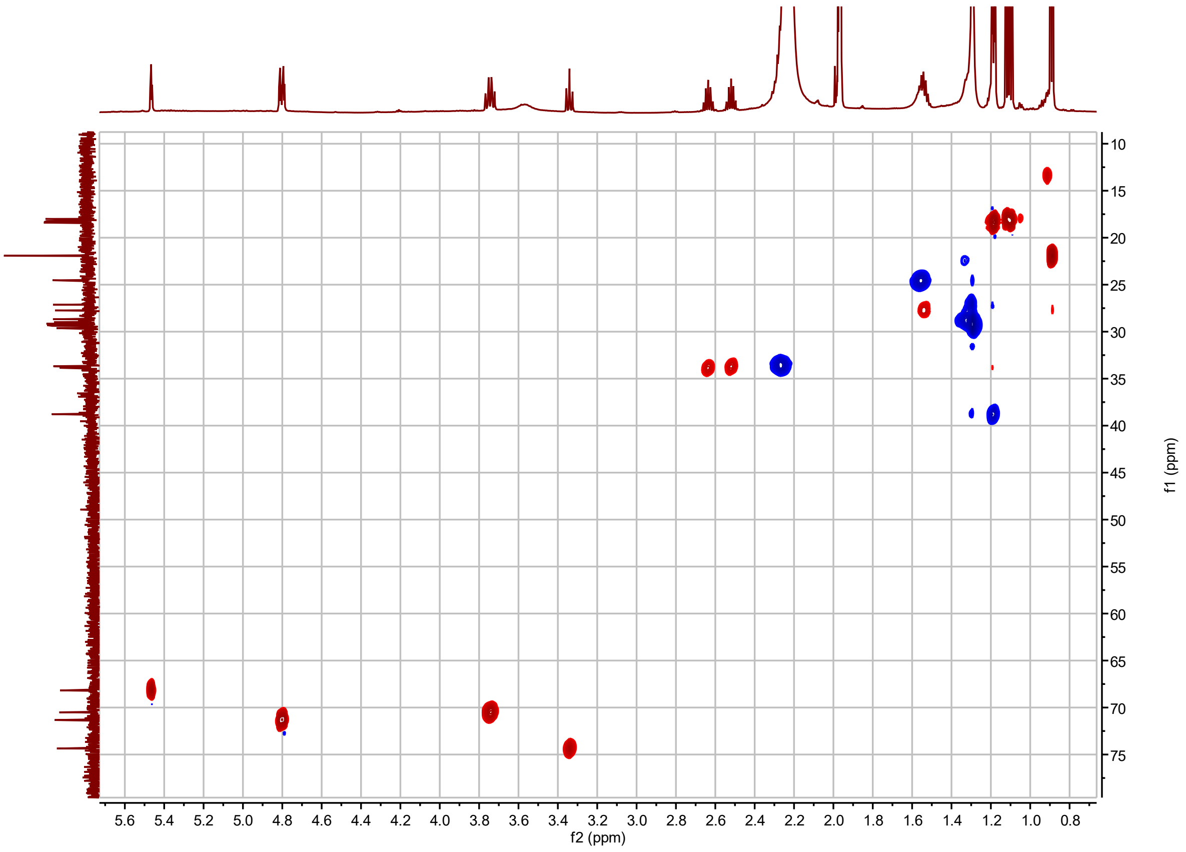


**I3:22(i4,i4,i14) $^{1}H$-$^{13}C$ HSQC**

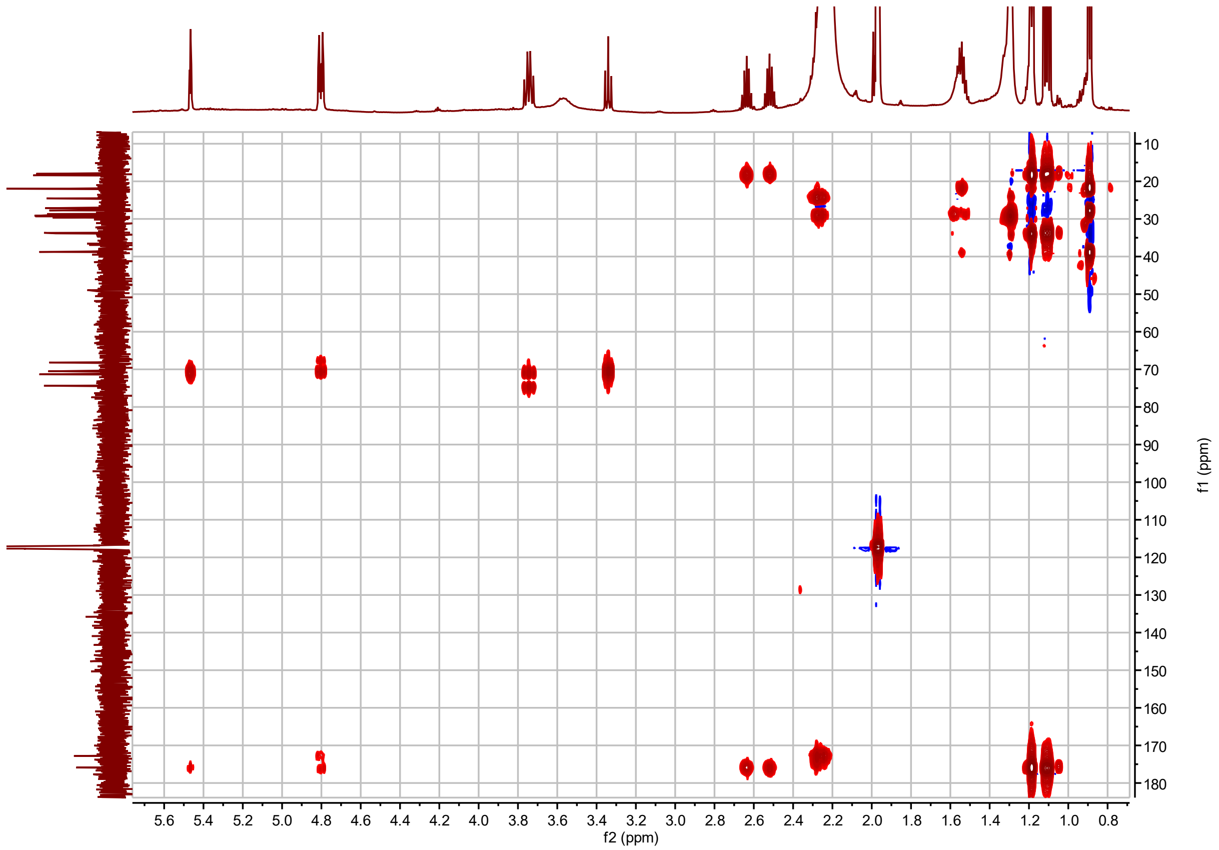


**I3:22(i4,i4,i14) $^{1}H$-$^{13}C$ HMBC**

**I3:22(i4,i4,n14) Chemical shifts and coupling constants.**

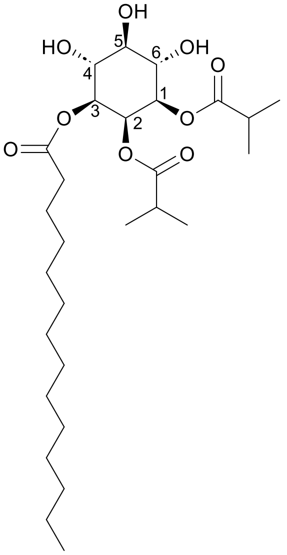


I3:22(i4,i4,n14)

Molecular Formula: $C_{28}H_{50}O_9$

Instrument: Bruker Avance NEO 800 MHz NMR

NMR solvent: $CD_3CN$

Fractions: 144-146

InChI Key: ZHGFFHULNGWPKL-RMLCQDLPSA-N

SMILES: CCCCCCCCCCCCCC(=O)O[C@H]1[C@H](O)[C@@H](O)[C@H](O)[C@@H](OC(=O)C(C)C)[C@H]1OC(=O)C(C)C

| Carbon # (Group) | $^{1}$H ($\delta$, ppm) | $^{13}$C ($\delta$, ppm) |
|---|---|---|
| 1(CH) | 4.80 (ddd, *J* = 2.98, 4.60, 10.20 Hz) | 71.89 or 71.87[a] |
| -1(CO) | | 176.56 |
| -2(CH) | 2.64 (hept, *J* = 6.97 Hz) | 34.41 |
| -3,4($CH_3$) | 1.18 (dd, *J* = 3.48, 6.98 Hz) | 18.96, 18.90 |
| 2(CH) | 5.46 (t, *J* = 2.97 Hz) | 68.72 |
| -1(CO) | | 176.43 |
| -2(CH) | 2.52 (hept, *J* = 6.97 Hz) | 34.26 or 34.23[a] |
| -3,4($CH_3$) | 1.11 (dd, *J* = 6.98, 17.92 Hz) | 18.79, 18.56 |
| 3(CH) | 4.80 (ddd, *J* = 2.98, 4.60, 10.20 Hz) | 71.89 or 71.87[a] |
| -1(CO) | | 173.34 |
| -2($CH_2$) | 2.26 (m) | 34.26 or 34.23[a] |
| -3($CH_2$) | 1.55 (m) | 25.10 |
| -4($CH_2$) | 1.29 (m) | 32.22 |
| -5-12($CH_2$) | 1.30 (m) | 29.95, 29.93, 29.89, 29.73, 29.65, 29.55, 29.24 |
| -13($CH_2$) | 1.33 (m) | 22.97 |
| -14($CH_3$) | 0.91 (m) | 13.96 |
| 4(CH) | 3.74 (q, *J* = 10.23 Hz) | 71.07 or 71.05[a] |
| 5(CH) | 3.34 (t, *J* = 9.33 Hz) | 74.90 |
| 6(CH) | 3.74 (q, *J* = 10.23 Hz) | 71.07 or 71.05[a] |

a – $^{13}$C signals not resolved in 2D spectra.

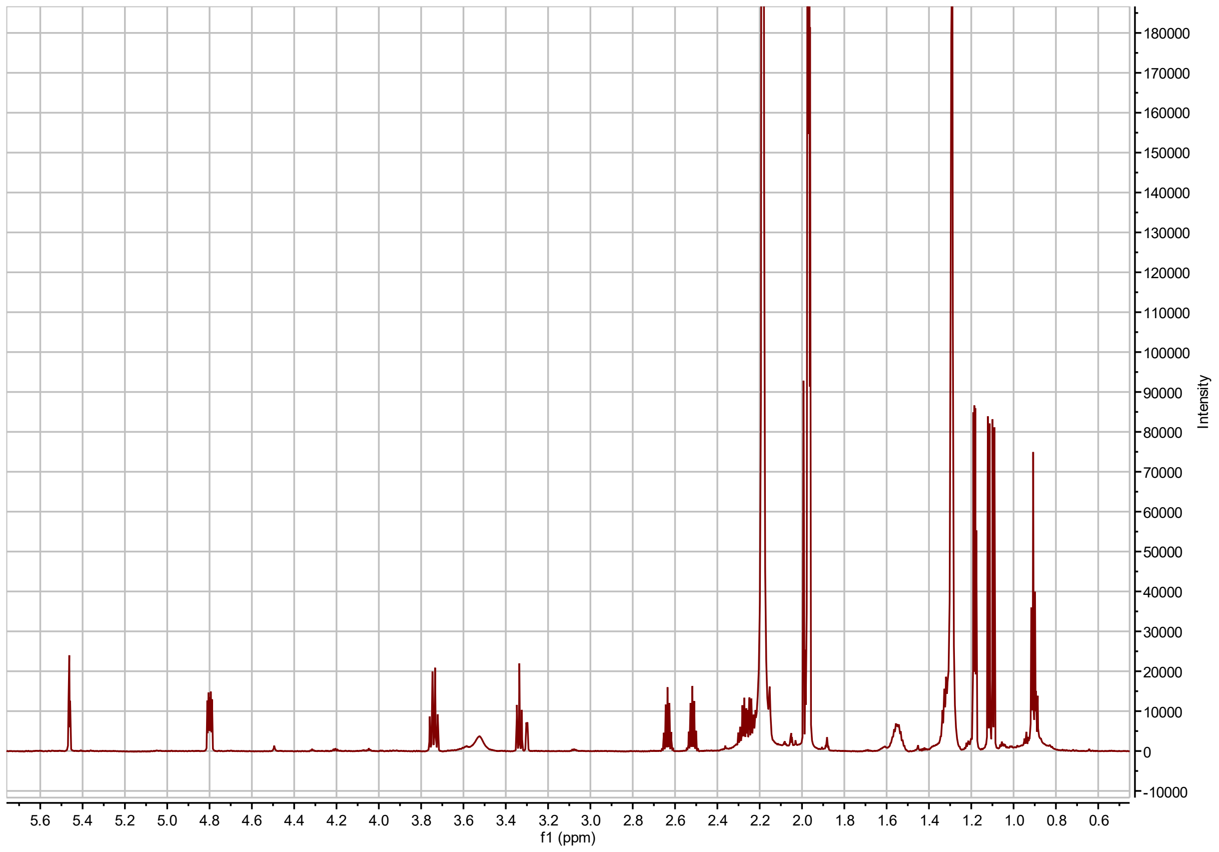


**I3:22(i4,i4,n14) $^{1}$H NMR**

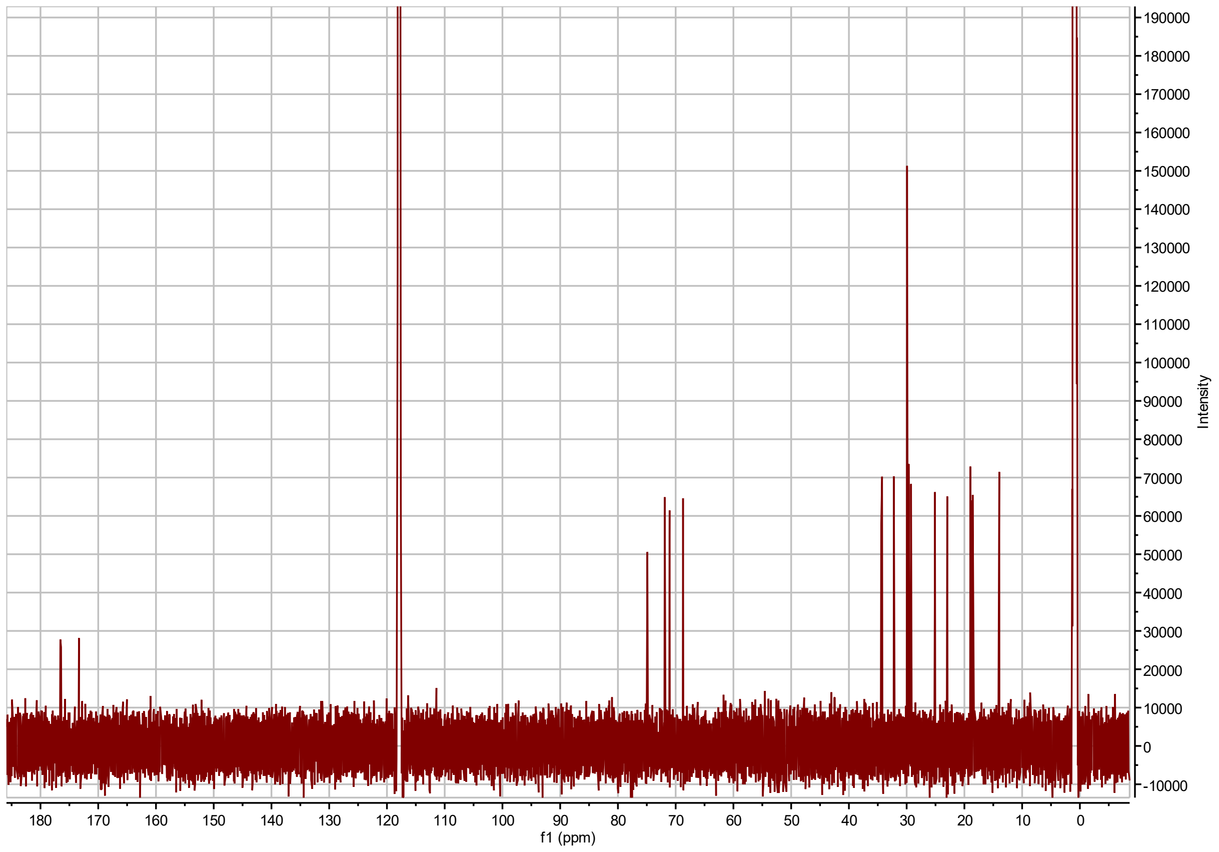

**I3:22(i4,i4,n14) $^{13}C$ NMR**

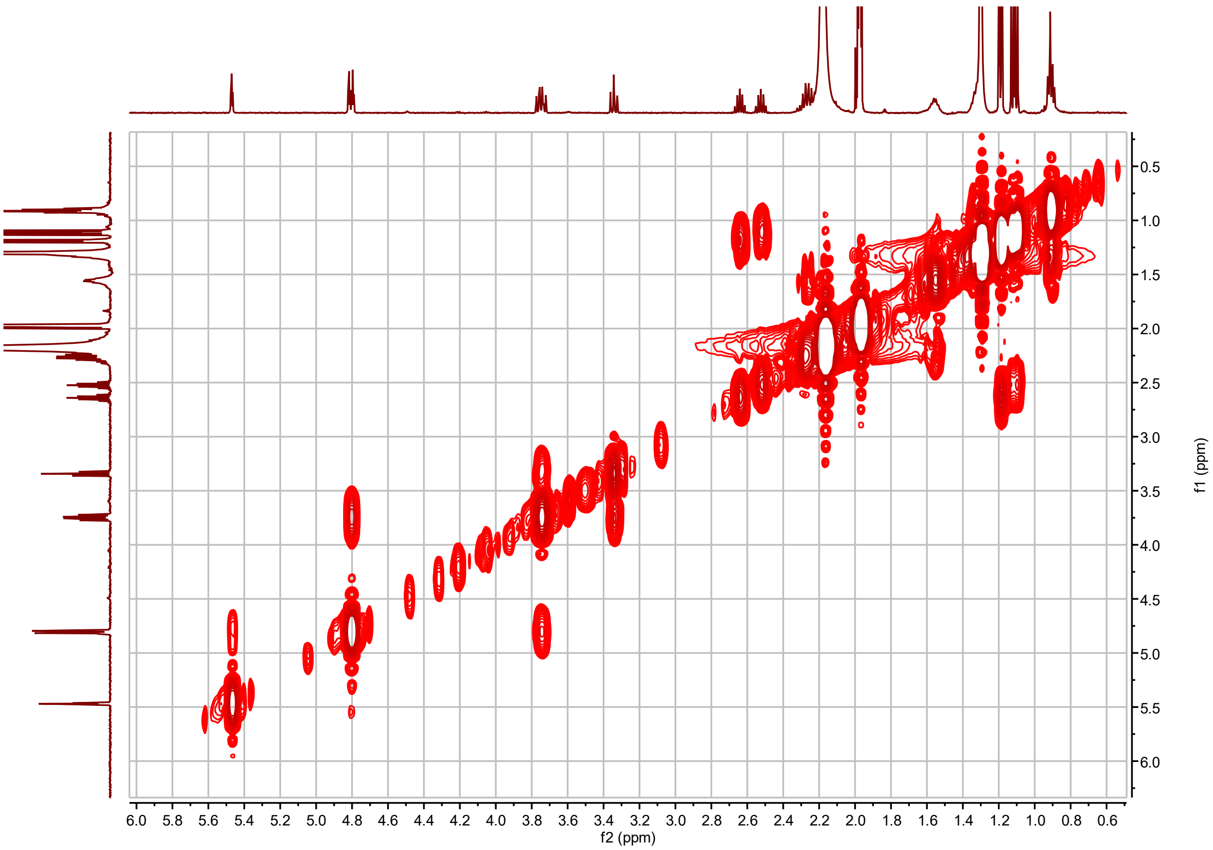


**I3:22(i4,i4,n14) $^1$H-$^1$H COSY**

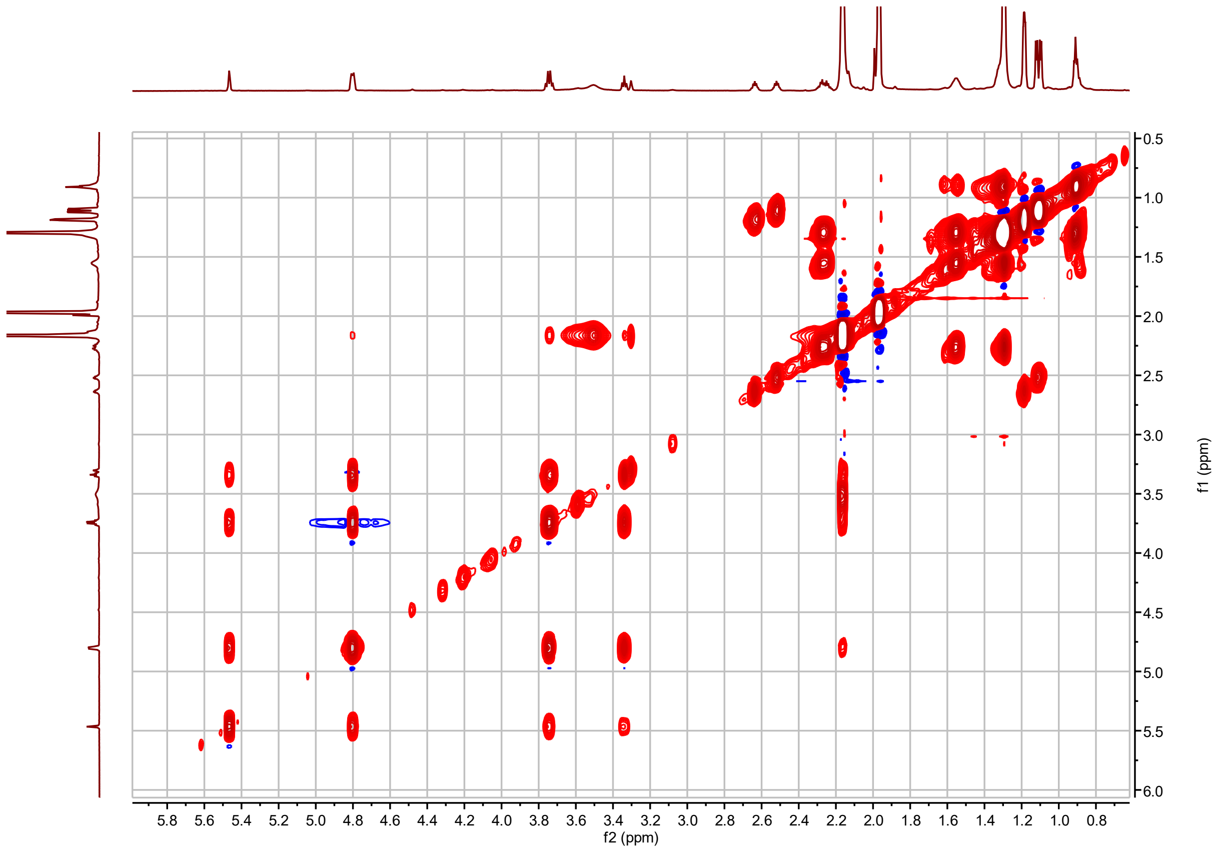


**I3:22(i4,i4,n14) $^{1}$H-$^{1}$H TOCSY**

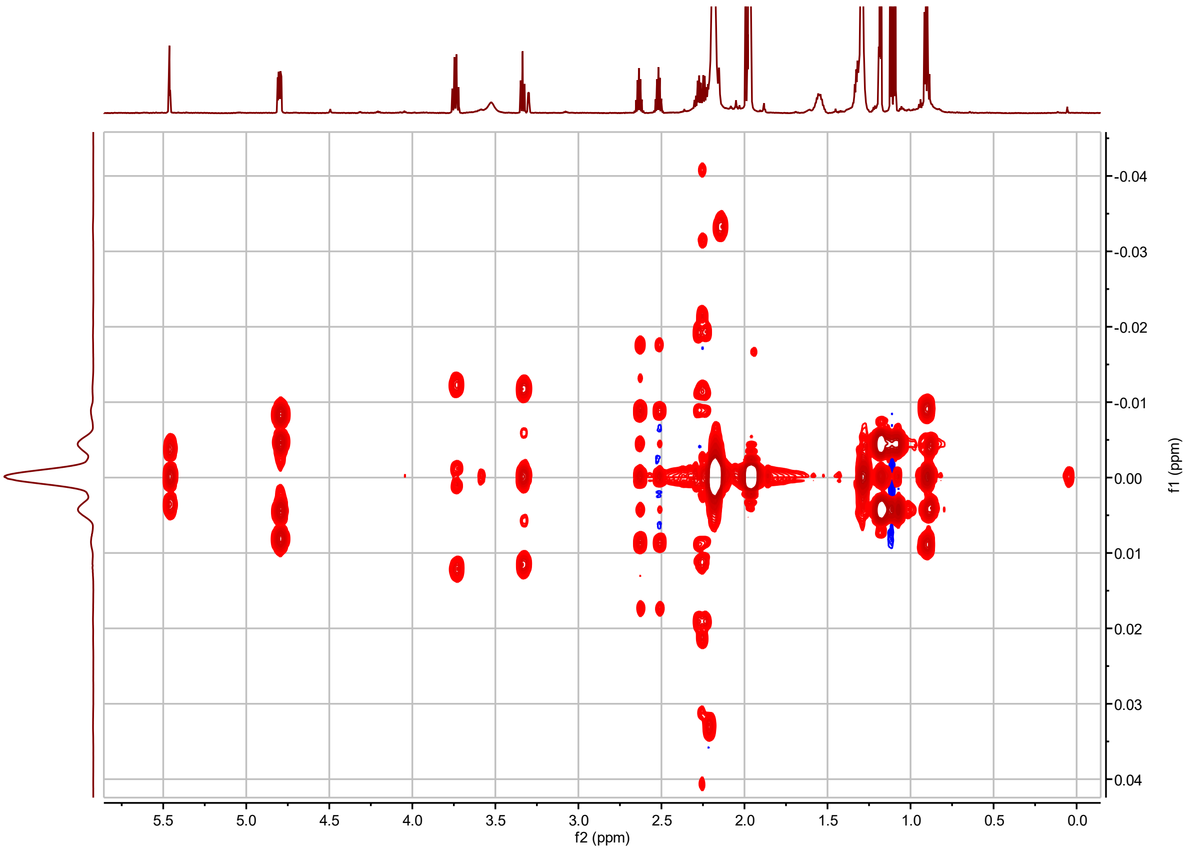


**I3:22(i4,i4,n14) *J*-resolved**

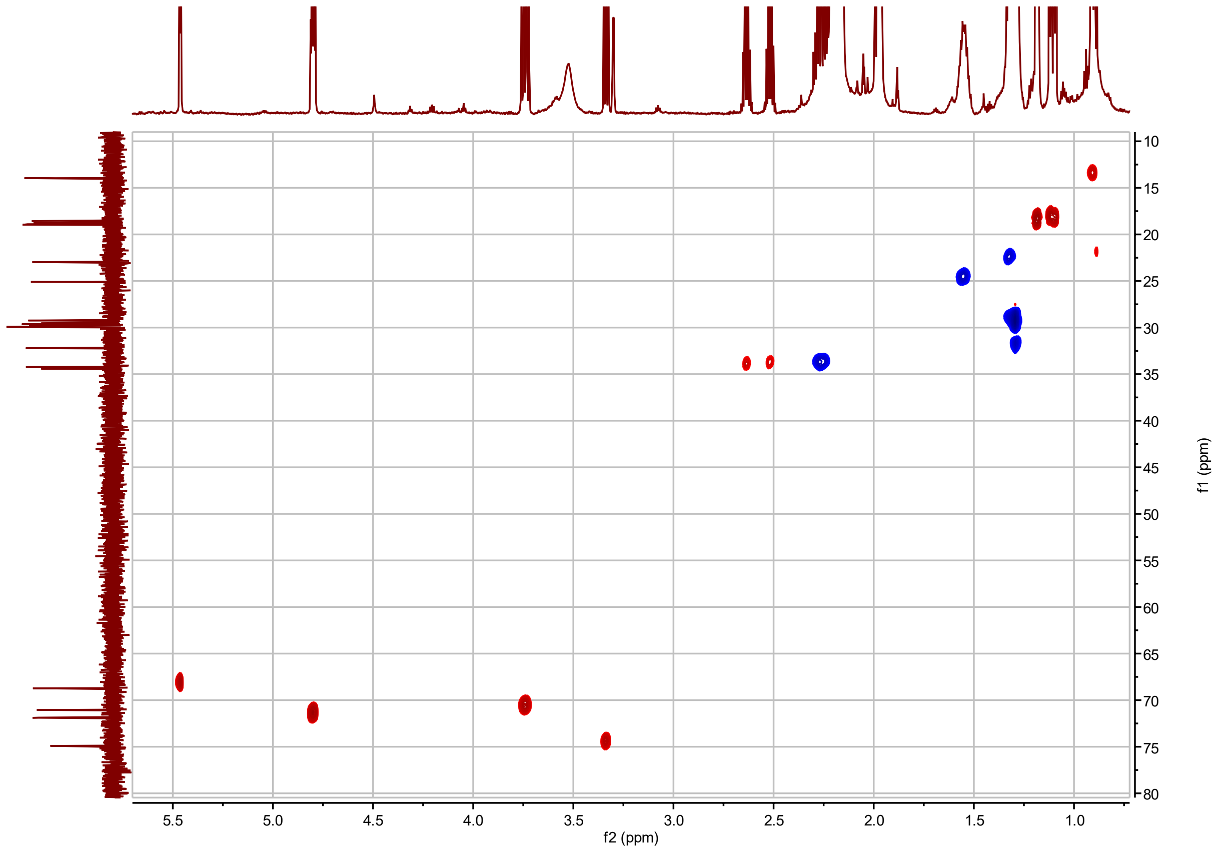


**I3:22(i4,i4,n14) $^{1}H$-$^{13}C$ HSQC**

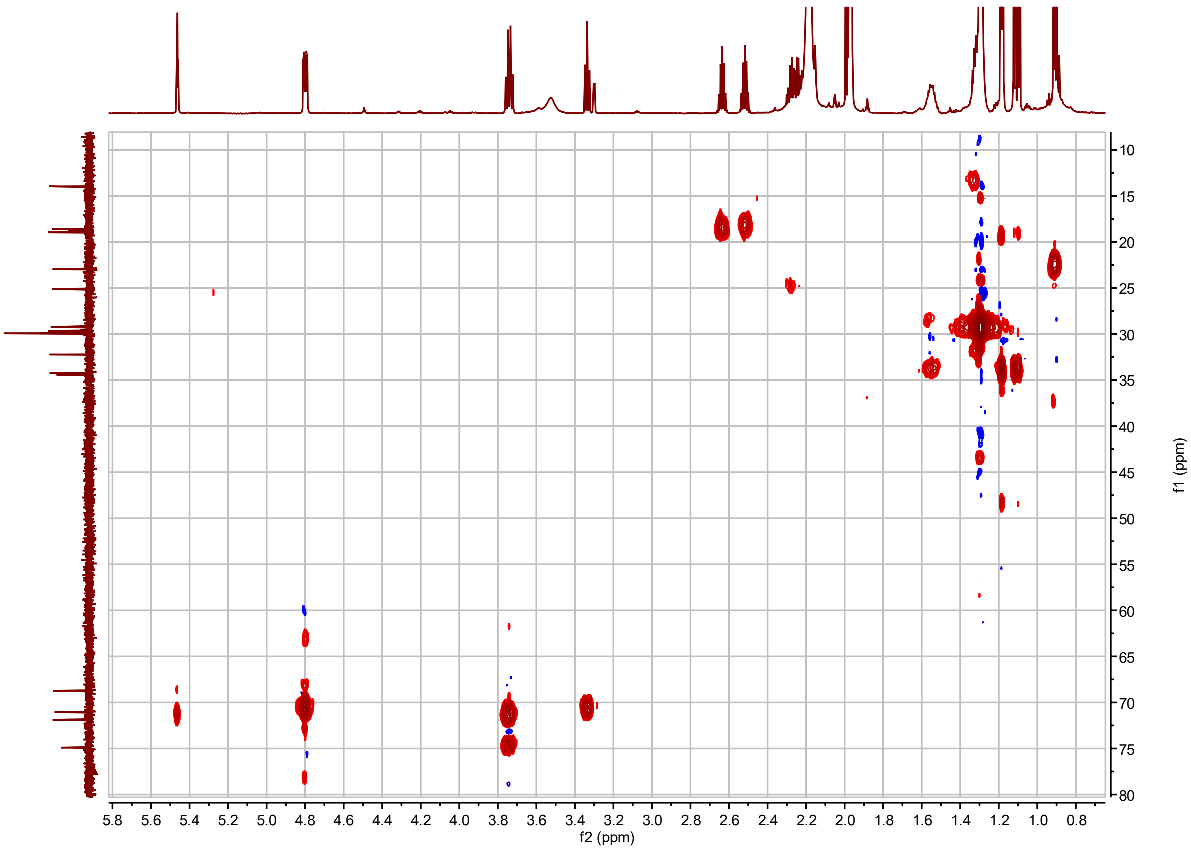


**I3:22(i4,i4,n14) $^{1}H$-$^{13}C$ H2BC**

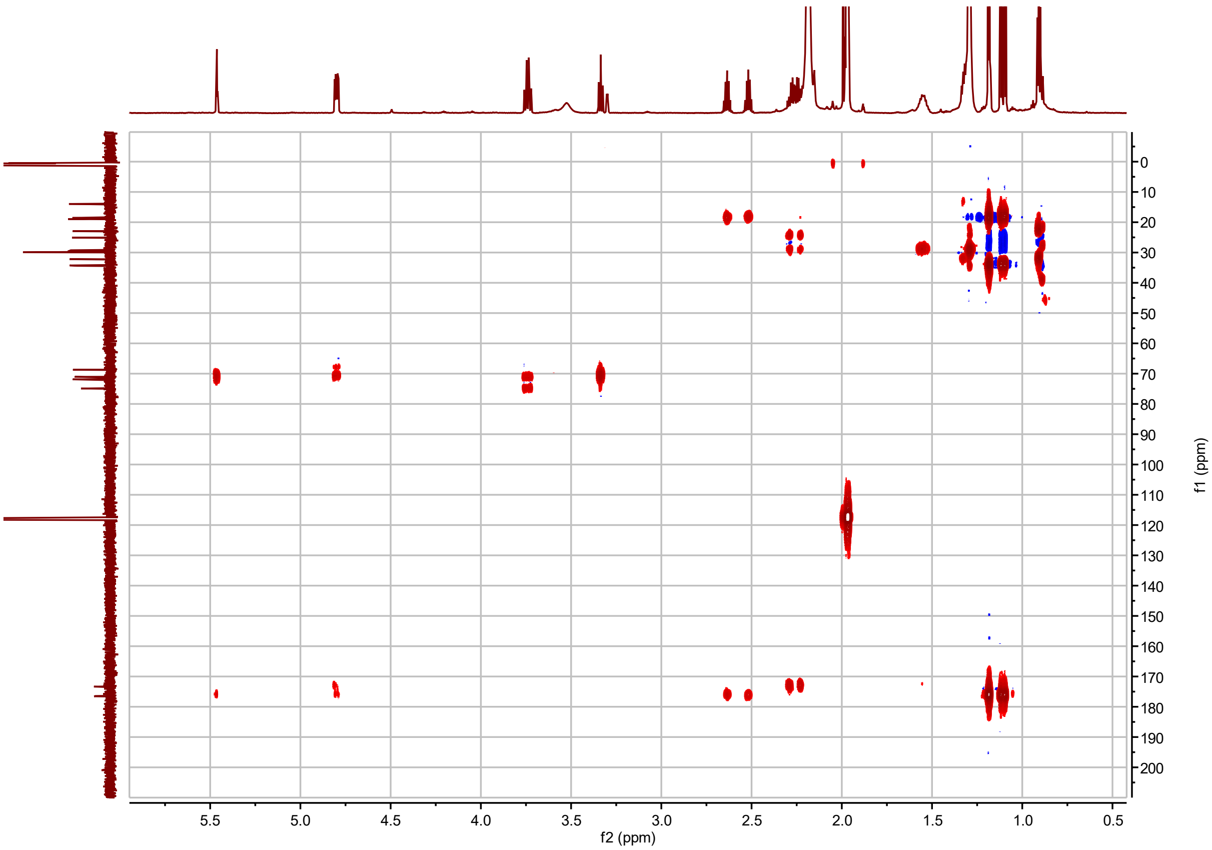


**I3:22(i4,i4,n14) $^{1}$H-$^{13}$C HMBC**

**I3:22(i4,i4,3-OH-i14) Chemical shifts and coupling constants.**

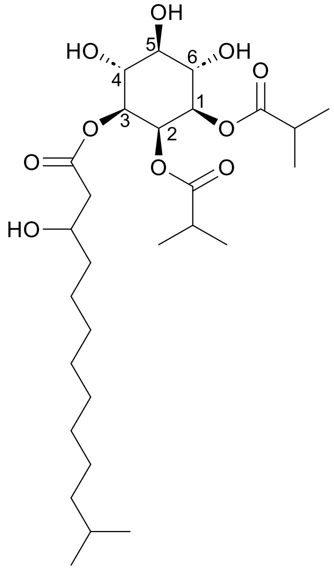

I3:22(i4,i4,3-OH-i14)

Molecular Formula: $C_{28}H_{50}O_{10}$

Instrument: Agilent 500 MHz DDR2

NMR solvent: $CD_3CN$

Fractions: 95-97

InChI Key: REKBIFNRGXIEET-JXDFUSHMSA-N

SMILES: CC(C)CCCCCCCCC(O)CC(=O)O[C@H]1[C@H](O)[C@@H](O)[C@H](O)[C@@H](OC(=O)C(C)C)[C@H]1OC(=O)C(C)C

| Carbon # (Group) | $^{1}$H ($\delta$, ppm) | $^{13}$C ($\delta$, ppm) |
|---|---|---|
| 1(CH) | 4.82 (dd, $J$ = 2.94, 10.21 Hz) | 71.34 |
| -1(CO) | | 175.98 |
| -2(CH) | 2.63 (hept, $J$ = 7.04, 7.04, 6.96, 6.96, 6.96 Hz) | 33.88 |
| -3,4($CH_3$) | 1.18 (dd, $J$ = 2.73, 6.97 Hz) | 18.33, 18.40 |
| 2(CH) | 5.48 (t, $J$ = 2.97 Hz) | 68.27 |
| -1(CO) | | 176.01 |
| -2(CH) | 2.51 (hept, $J$ = 7.00 Hz) | 33.70 |
| -3,4($CH_3$) | 1.11 (dd, $J$ = 7.00, 12.57 Hz) | 18.02, 18.22 |
| 3(CH) | 4.86 (dd, $J$ = 2.94, 10.18 Hz) | 71.50 |
| -1(CO) | | 171.16 |
| -2($CH_2$) | 2.46 (dd, $J$ = 4.25, 15.29 Hz)<br>2.30 (dd, $J$ = 8.44, 15.32 Hz) | 42.14 |
| -3(CHOH) | 3.92 (m) | 67.75 |
| -4($CH_2$) | 1.41 (m) | 36.53 |
| -5-9($CH_2$) | 1.29 (m) | 25.25, 29.26, 29.32, 29.38,29.65 |
| -10($CH_2$) | 1.29 (m) | 27.18 |
| -11($CH_2$) | 1.18 (m) | 38.7 |
| -12(CH) | 1.53 (hept, $J$ = 6.72 Hz) | 27.75 |
| -13,14($CH_3$) | 0.89 (d, $J$ = 6.64 Hz) | 21.89 |
| 4(CH) | 3.76 (td, $J$ = 5.22, 9.71, 9.76 Hz) | 70.66 or 70.54[a] |
| 5(CH) | 3.36 (t, $J$ = 9.31 Hz) | 74.15 |
| 6(CH) | 3.76 (td, $J$ = 5.22, 9.71, 9.76 Hz) | 70.66 or 70.54[a] |

a – $^{13}$C signals not resolved in 2D spectra.

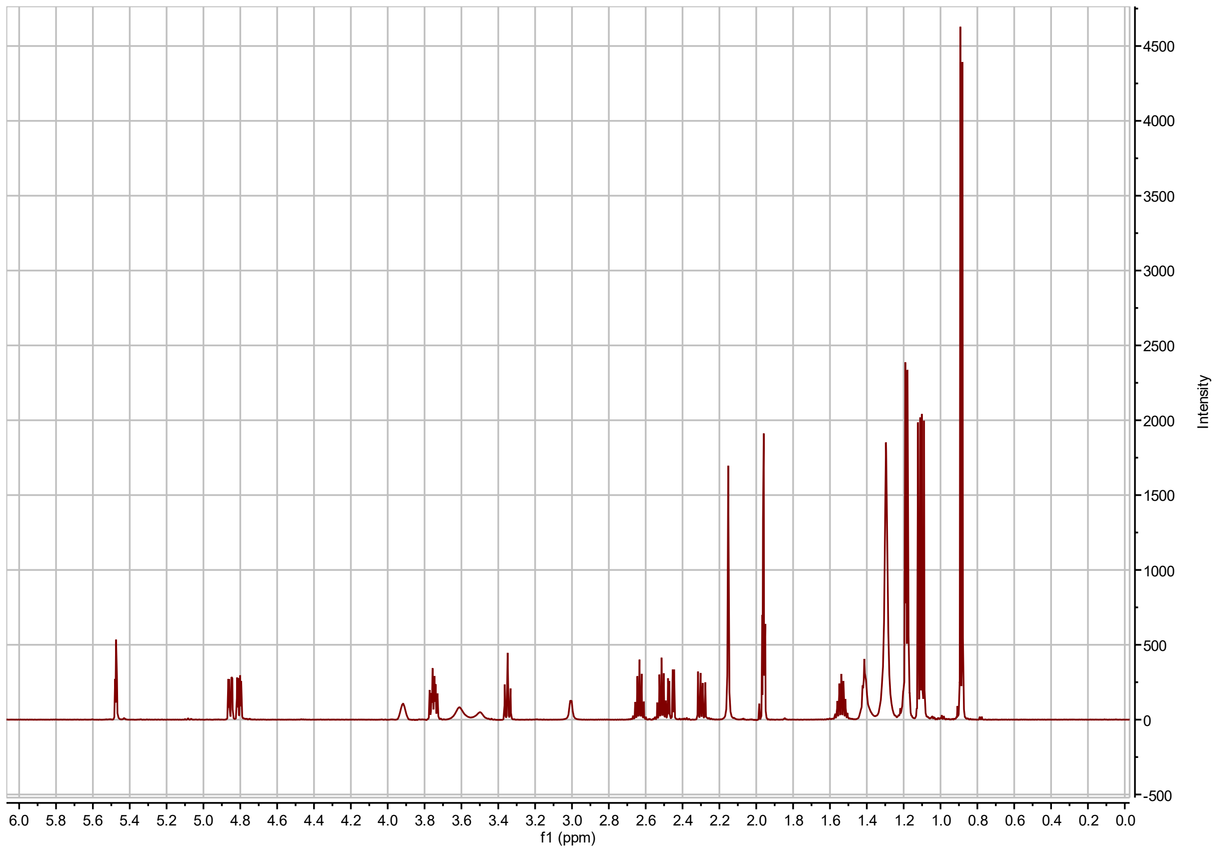


**I3:22(i4,i4,3-OH-i14) $^{1}$H NMR**

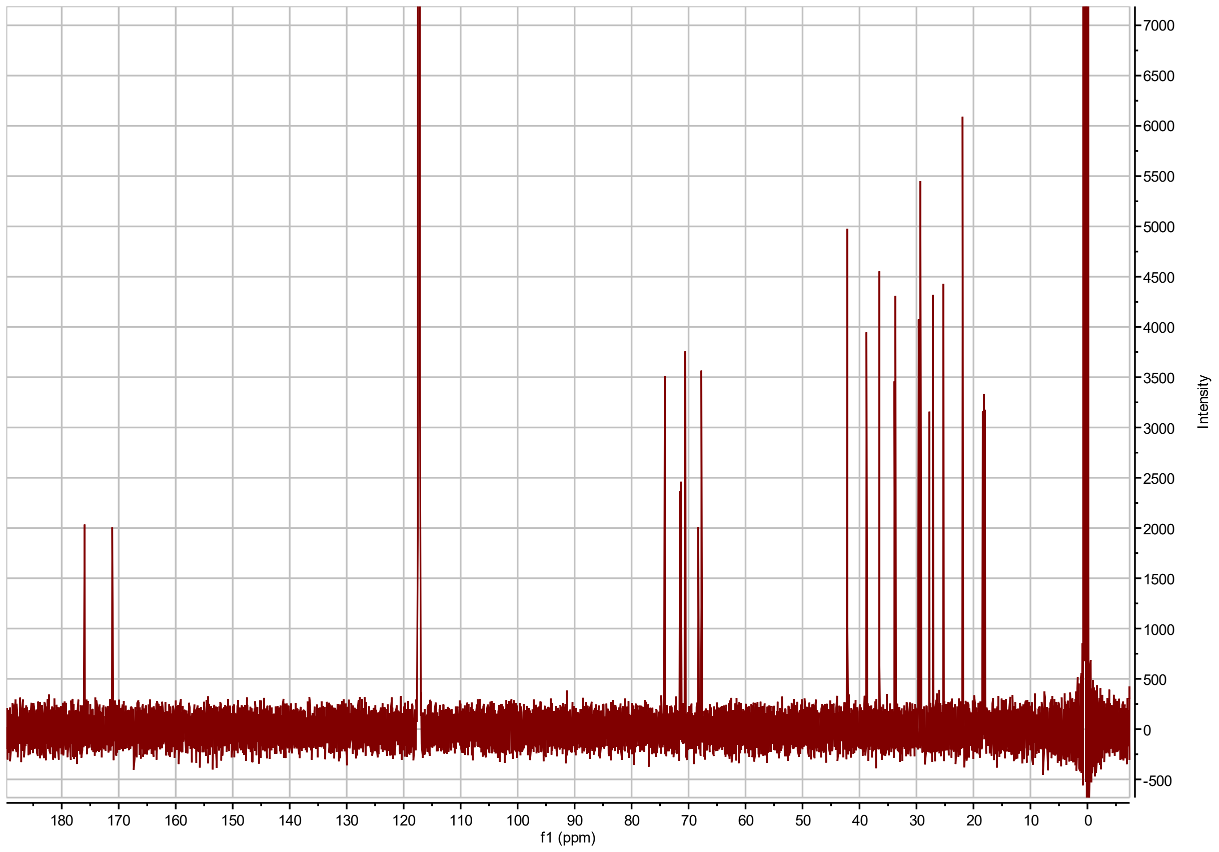


**I3:22(i4,i4,3-OH-i14) $^{13}$C NMR**

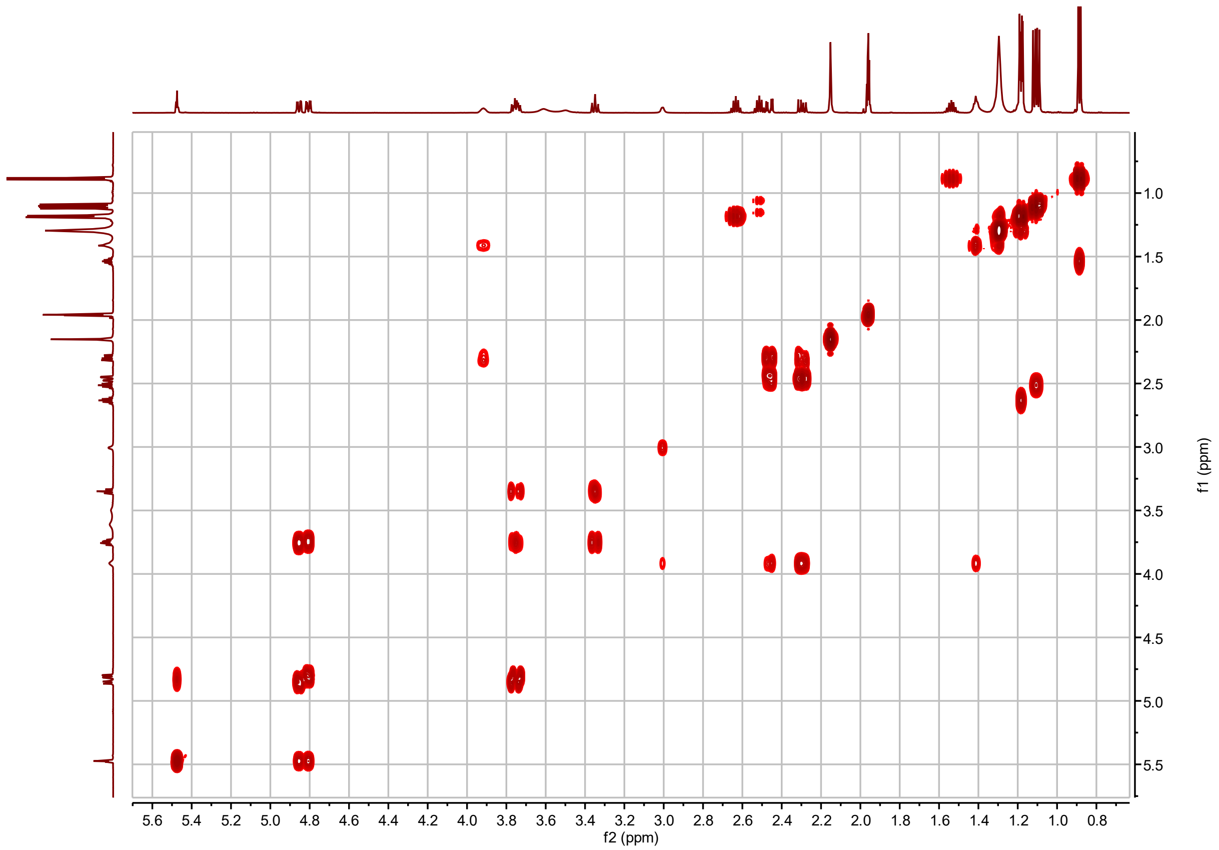


**I3:22(i4,i4,3-OH-i14) $^{1}$H-$^{1}$H COSY**

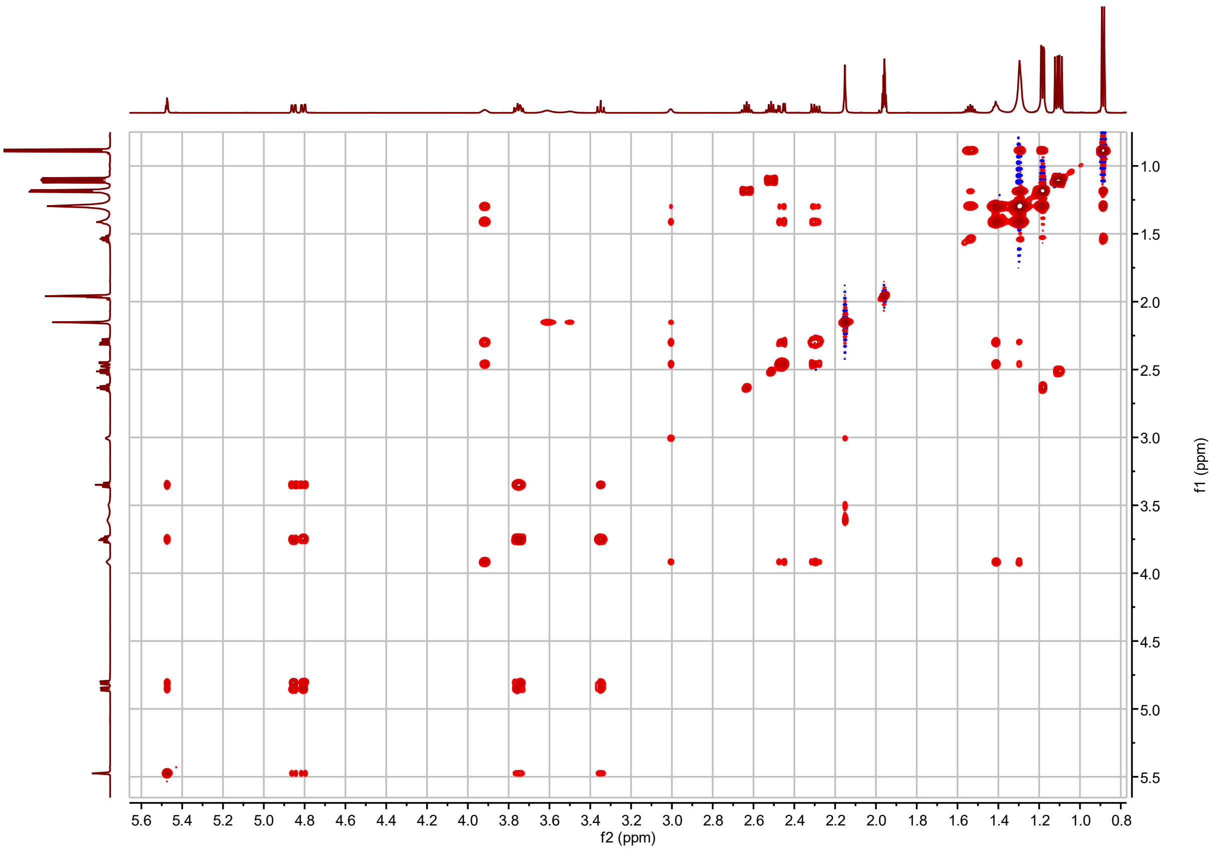


**I3:22(i4,i4,3-OH-i14) $^1$H-$^1$H TOCSY**

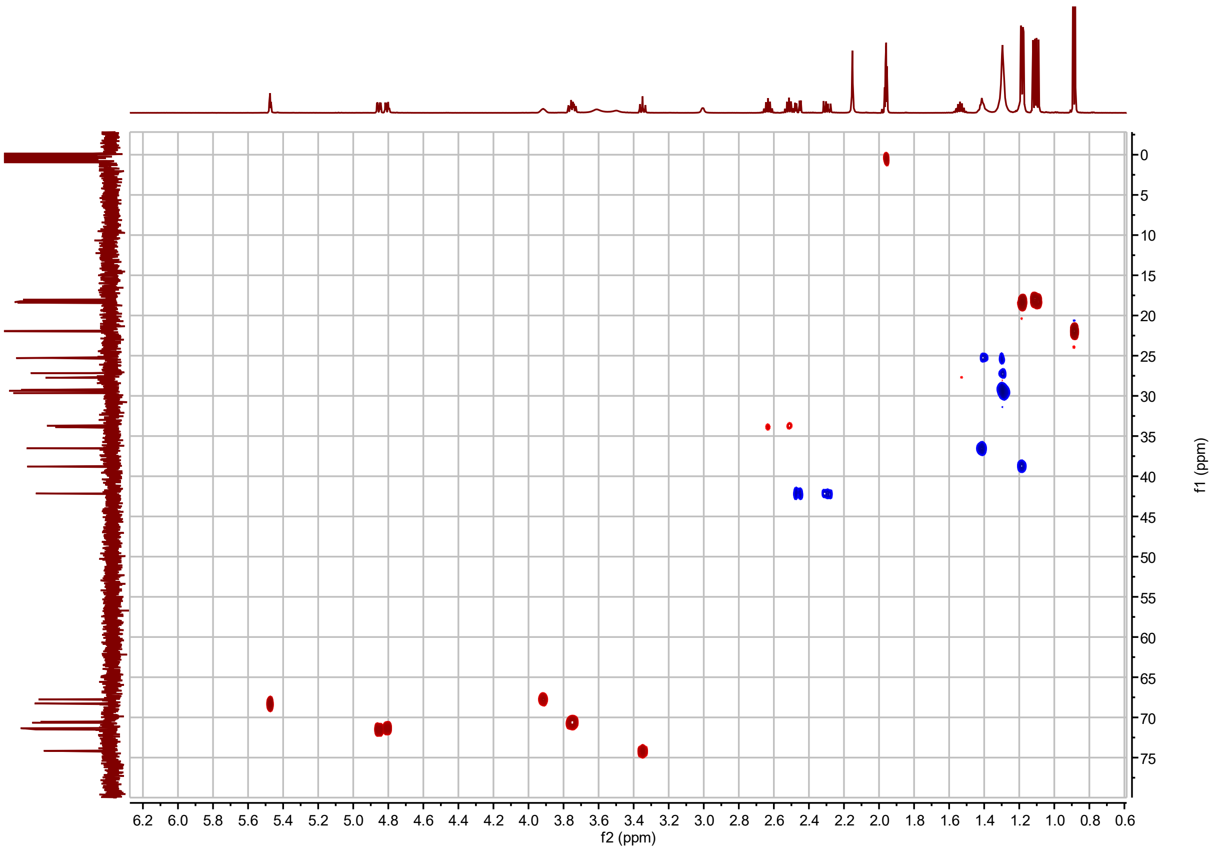


**I3:22(i4,i4,3-OH-i14) $^{1}H$-$^{13}C$ HSQC**

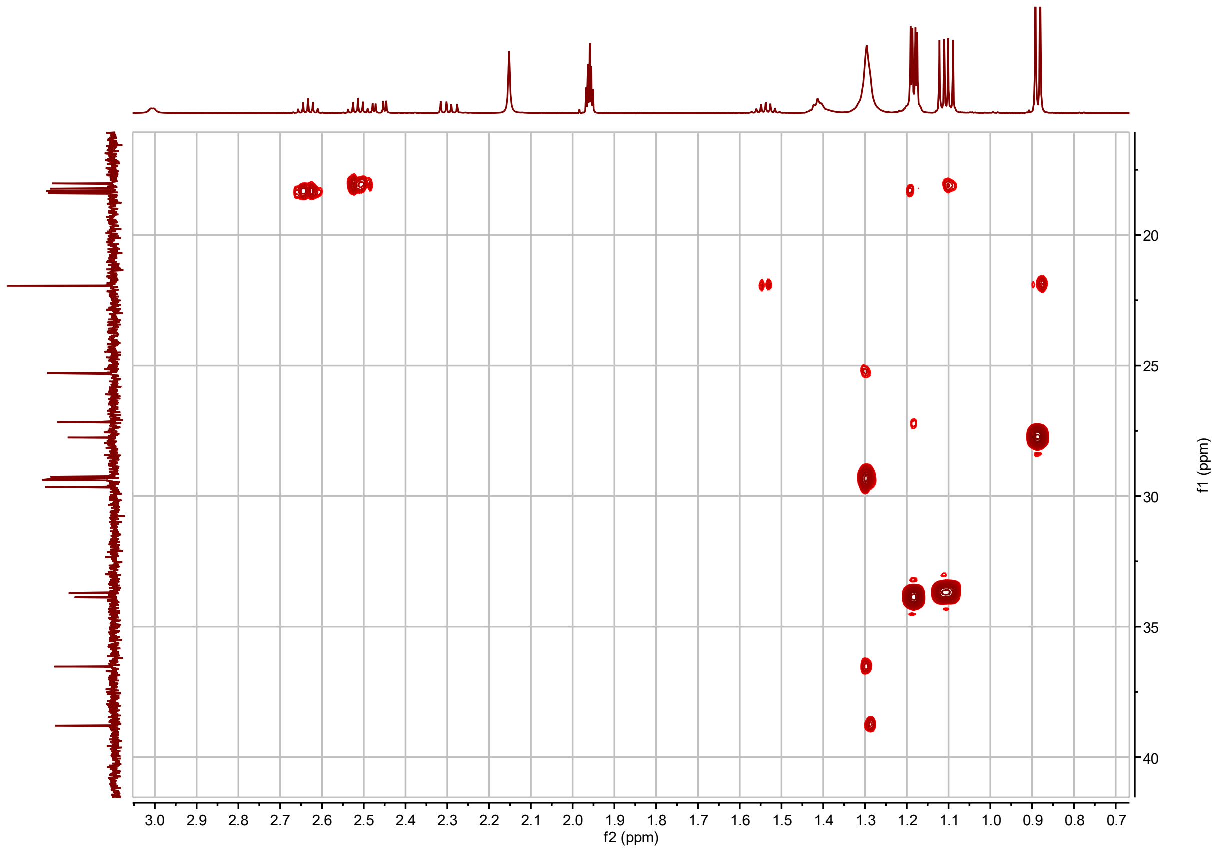


**I3:22(i4,i4,3-OH-i14) $^{1}$H-$^{13}$C H2BC**

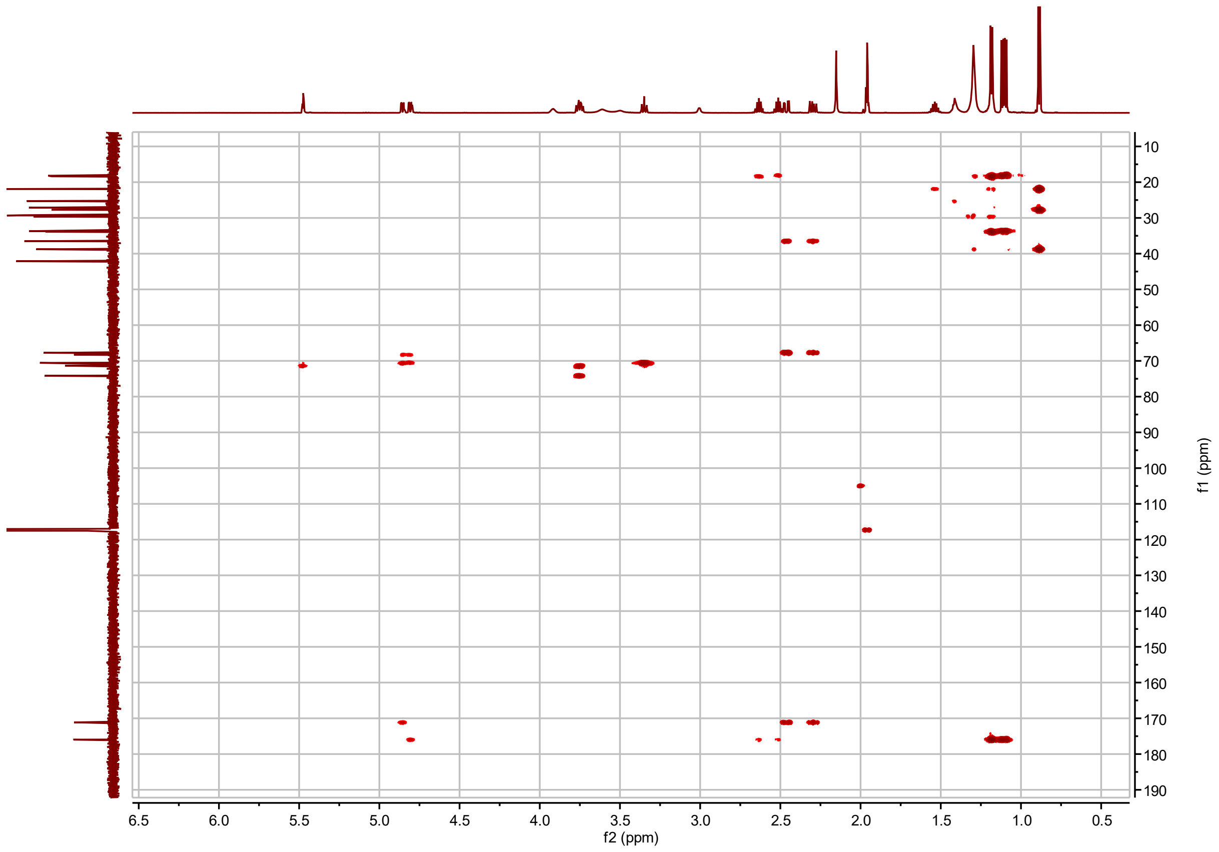


**I3:22(i4,i4,3-OH-i14) $^{1}H$-$^{13}C$ HMBC**

**I3:22(i4,i4,(3*R*)-OH-n14) Chemical shifts and coupling constants.**

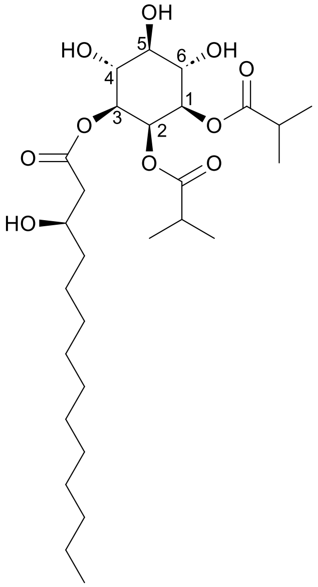


I3:22(i4,i4,(3*R*)-OH-n14)

Molecular Formula: $C_{28}H_{50}O_{10}$

Instrument: Agilent 500 MHz DDR2

NMR solvent: $CD_3CN$

Fractions: 101-103

InChI Key: KJJFWJBFXVGTFN-ARLPEISOSA-N

SMILES:
CCCCCCCCCCC[C@@H](O)CC(=O)O[C@H]1[C@H](O)[C@@H](O)[C@H](O)[C@@H](OC(=O)C(C)C)[C@H]1OC(=O)C(C)C

| Carbon # (Group) | $^1$H ($\delta$, ppm) | $^{13}$C ($\delta$, ppm) |
|---|---|---|
| 1(CH) | 4.82 (dd, *J* = 2.96, 10.21 Hz) | 71.34 |
| -1(CO) | | 175.98 |
| -2($C_H$) | 2.63 (hept, *J* = 7.04, 7.04, 6.96, 6.96, 6.96 Hz) | 33.88 |
| -3,4($CH_3$) | 1.18 (dd, *J* = 2.73, 6.97 Hz) | 18.33, 18.40 |
| 2(CH) | 5.48 (t, *J* = 2.97 Hz) | 68.27 |
| -1(CO) | | 176.01 |
| -2(CH) | 2.51 (hept, *J* = 7.00 Hz) | 33.71 |
| -3,4($CH_3$) | 1.11 (d, *J* = 7.00 Hz) | 18.02, 18.22 |
| 3(CH) | 4.86 (dd, *J* = 2.95, 10.17 Hz) | 71.50 |
| -1(CO) | | 171.16 |
| -2($CH_2$) | 2.30 (dd, *J* = 8.44, 15.32 Hz)<br>2.46 (dd, *J* = 4.25, 15.29 Hz) | 42.14 |
| -3(CHOH) | 3.92 (m) | 67.75 |
| -4($CH_2$) | 1.42 (m) | 36.53 |
| -5-12($CH_2$) | 1.41-1.30 (m) | 29.41, 29.38, 29.35, 29.25, 29.11, 25.30, 22.42 |
| -13($CH_2$) | 1.30 (m) | 31.67 |
| -14($CH_3$) | 0.91(m) | 13.42 |
| 4(CH) | 3.76 (m) | 70.54 or 70.66[a] |
| 5(CH) | 3.36 (t, *J* = 9.32 Hz) | 74.16 |
| 6(CH) | 3.76 (m) | 70.54 or 70.66[a] |

a – $^{13}$C signals not resolved in 2D spectra.

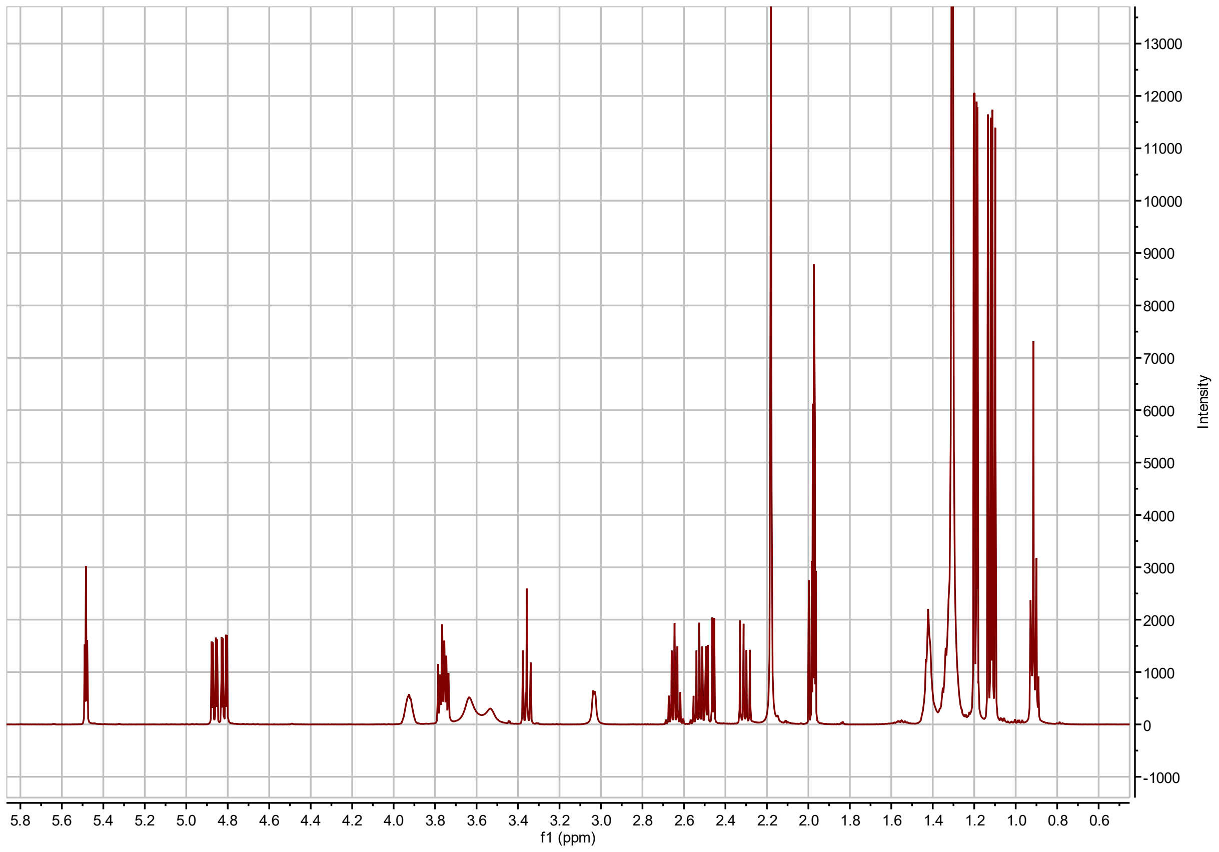


**I3:22(i4,i4,3-OH-n14) $^{1}$H NMR**

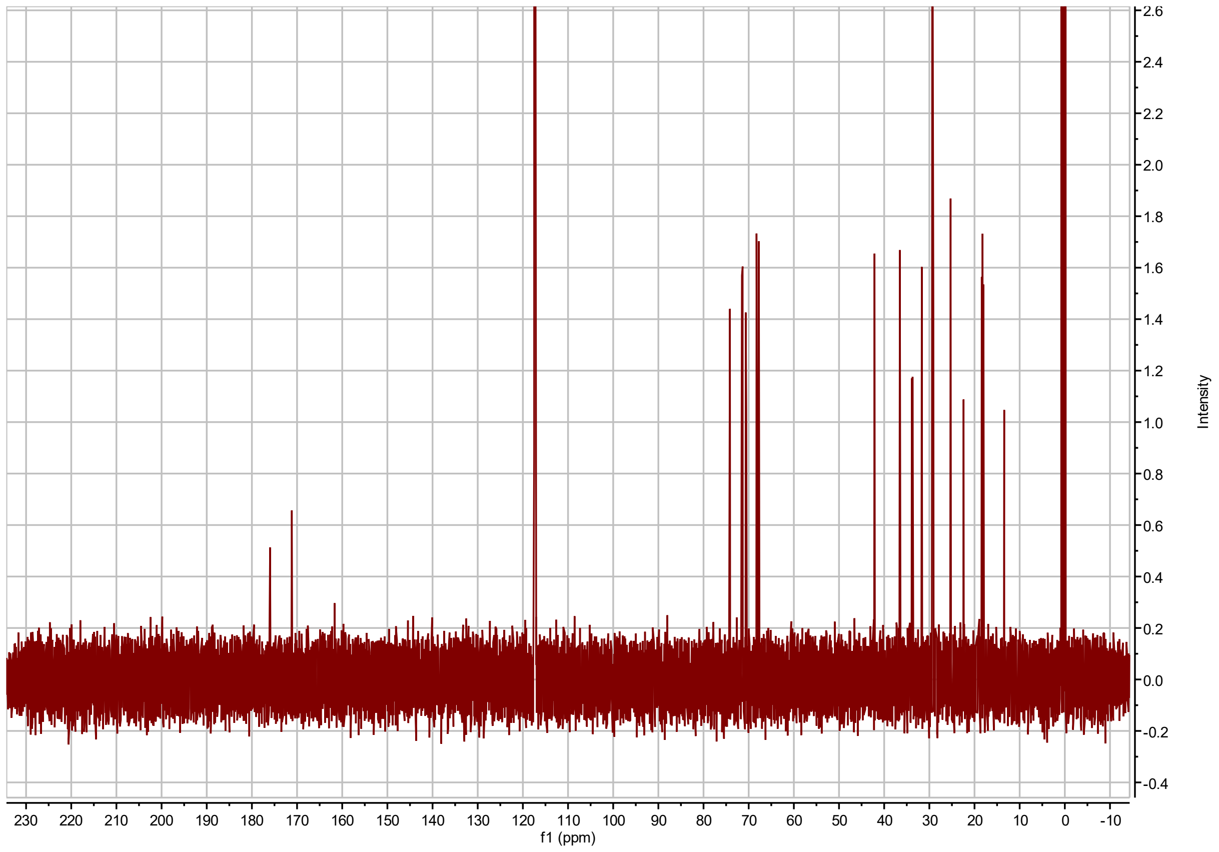


**I3:22(i4,i4,3-OH-n14) $^{13}$C NMR**

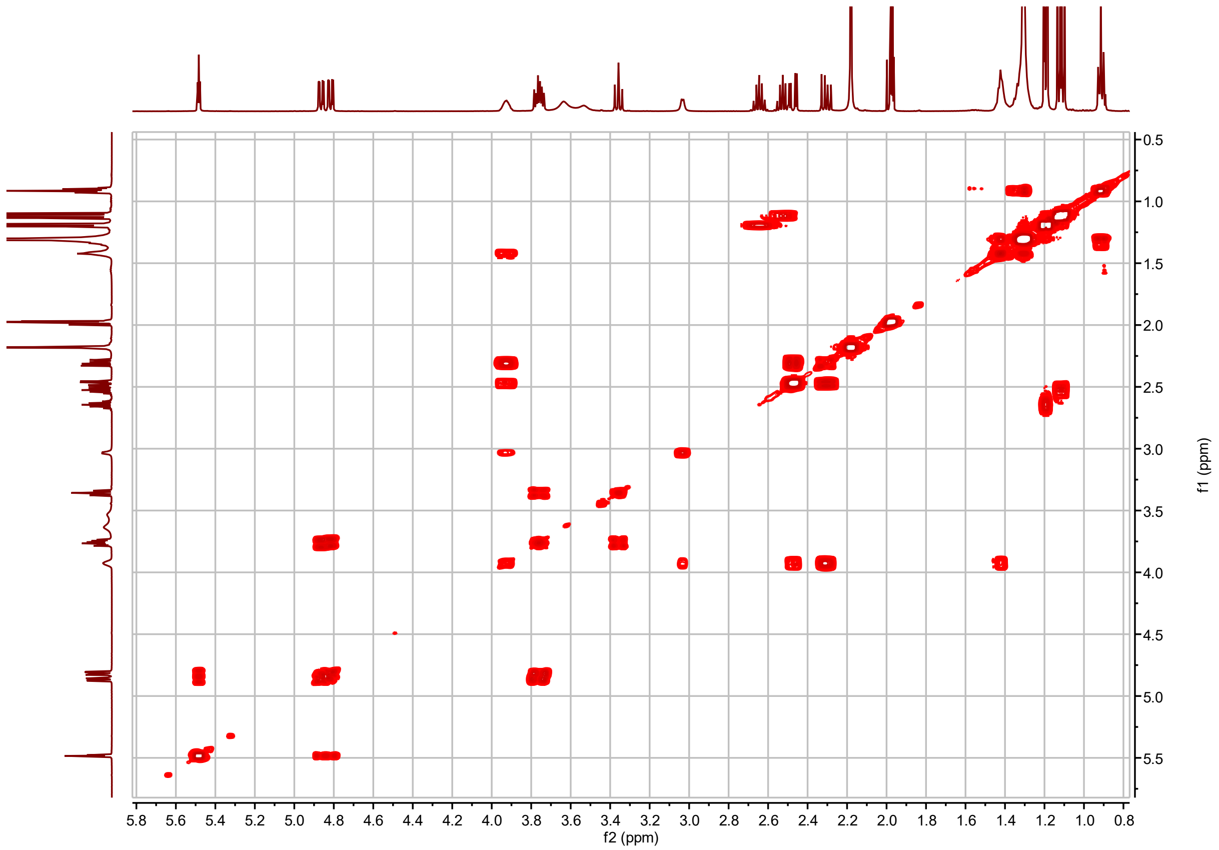


**I3:22(i4,i4,3-OH-n14) $^{1}$H-$^{1}$H COSY**

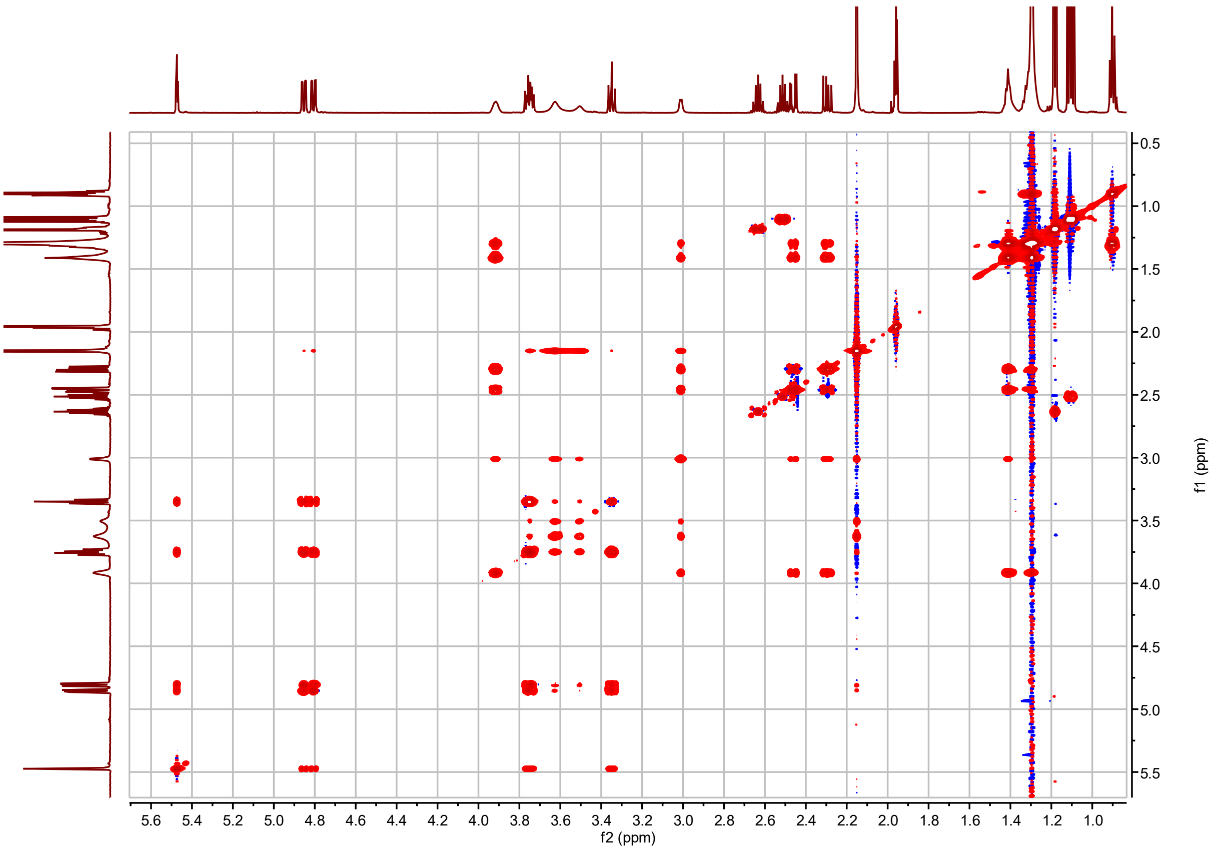


**I3:22(i4,i4,3-OH-n14) $^{1}$H-$^{1}$H TOCSY**

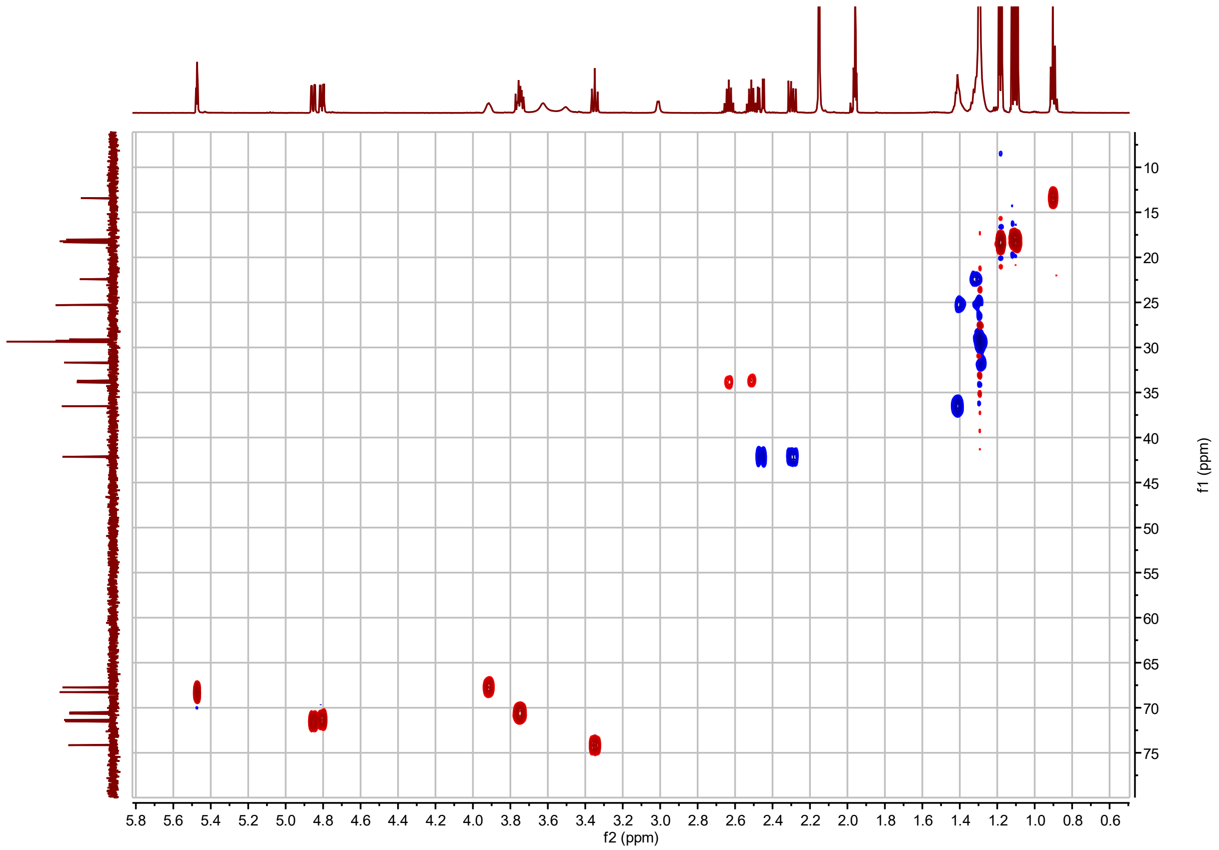


**I3:22(i4,i4,3-OH-n14) $^{1}H$-$^{13}C$ HSQC**

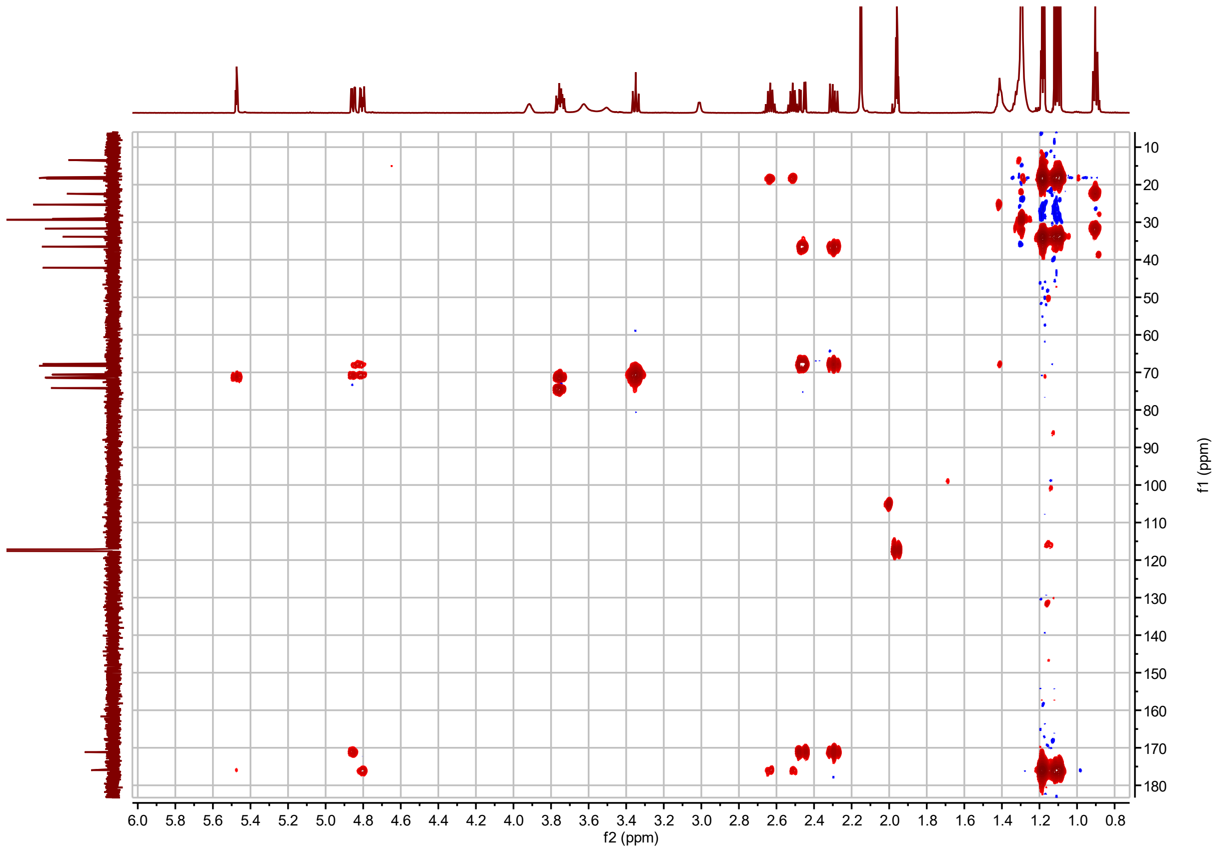


**I3:22(i4,i4,3-OH-n14) $^{1}$H-$^{13}$C HMBC**

**4-*O*-β-arabinosyl-*myo*-inositol Chemical shifts and coupling constants.**

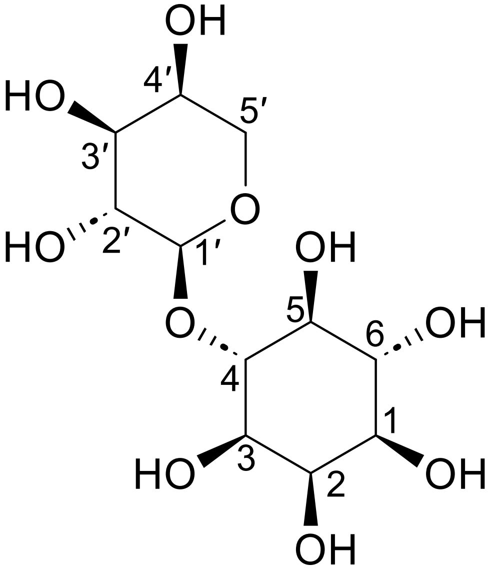


4-*O*-β-arabinosyl-*myo*-inositol derived from saponified AI4:18(2,4,4,8)

Molecular Formula: $C_{11}H_{20}O_{10}$

Instrument: Agilent 500 MHz DDR2

NMR solvent: $D_2O$

InChI Key: ZTUXXEBTGKCWOB-GZFMRODOSA-N

SMILES:
O[C@H]1CO[C@@H](O[C@@H]2[C@@H](O)[C@H](O)[C@@H](O)[C@@H](O)[C@H]2O)[C@H](O)[C@H]1O

| Carbon # (Group) | $^{1}$H ($\delta$, ppm) | $^{13}$C ($\delta$, ppm) |
|---|---|---|
| 1(CH) | 3.50 (dd, *J* = 3.3, 9.7 Hz) | 72.18 or 72.12[a] |
| 2(CH) | 3.86 (t, *J* = 2.95 Hz) | 71.96 |
| 3(CH) | 3.53 (dd, *J* = 2.9, 9.8 Hz) | 70.69 |
| 4(CH) | 3.60 (t, *J* = 10.1 Hz) | 81.93 |
| 5(CH) | 3.19 (t, *J* = 9.2 Hz) | 72.48 |
| 6(CH) | 3.44 (t, *J* = 10.4 Hz) | 72.18 or 72.12[a] |
| 1′(CH) | 4.37 (d, *J* = 7.61 Hz) | 103.72 |
| 2′(CH) | 3.43 (t, *J* = 9.8 Hz) | 71.18 |
| 3′(CH) | 3.48 (dd, *J* = 1.7, 13.5 Hz) | 72.18 or 72.12[a] |
| 4′(CH) | 3.75 (m) | 68.19 |
| 5′($CH_2$) | 3.77 (dd, *J* = 2.38, 13.4 Hz); 3.48 (dd, *J* = 1.7, 13.4 Hz) | 66.18 |

a – $^{13}$C signals not resolved in 2D spectra.

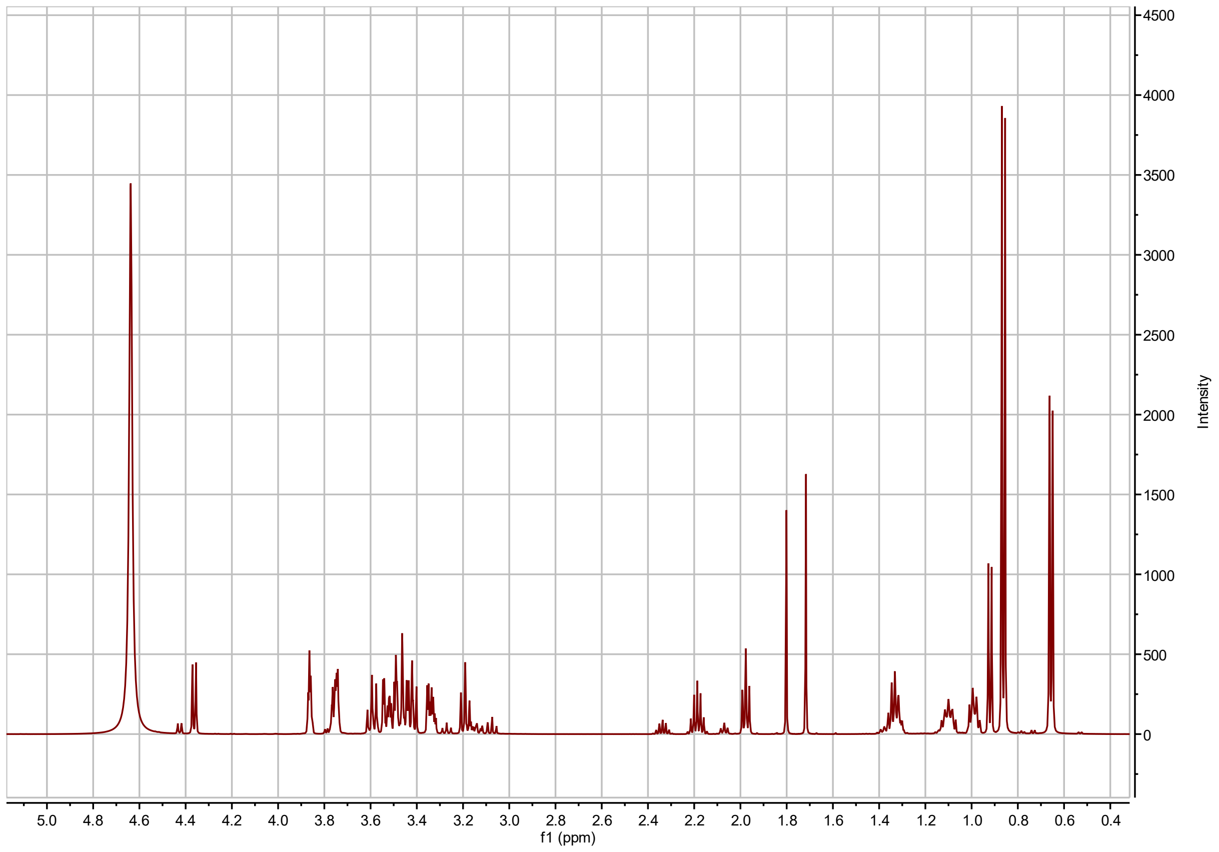


**4-*O*-arabinopyranosyl *myo*-inositol derived from saponified AI4:18(2,4,4,8) $^{1}$H NMR**

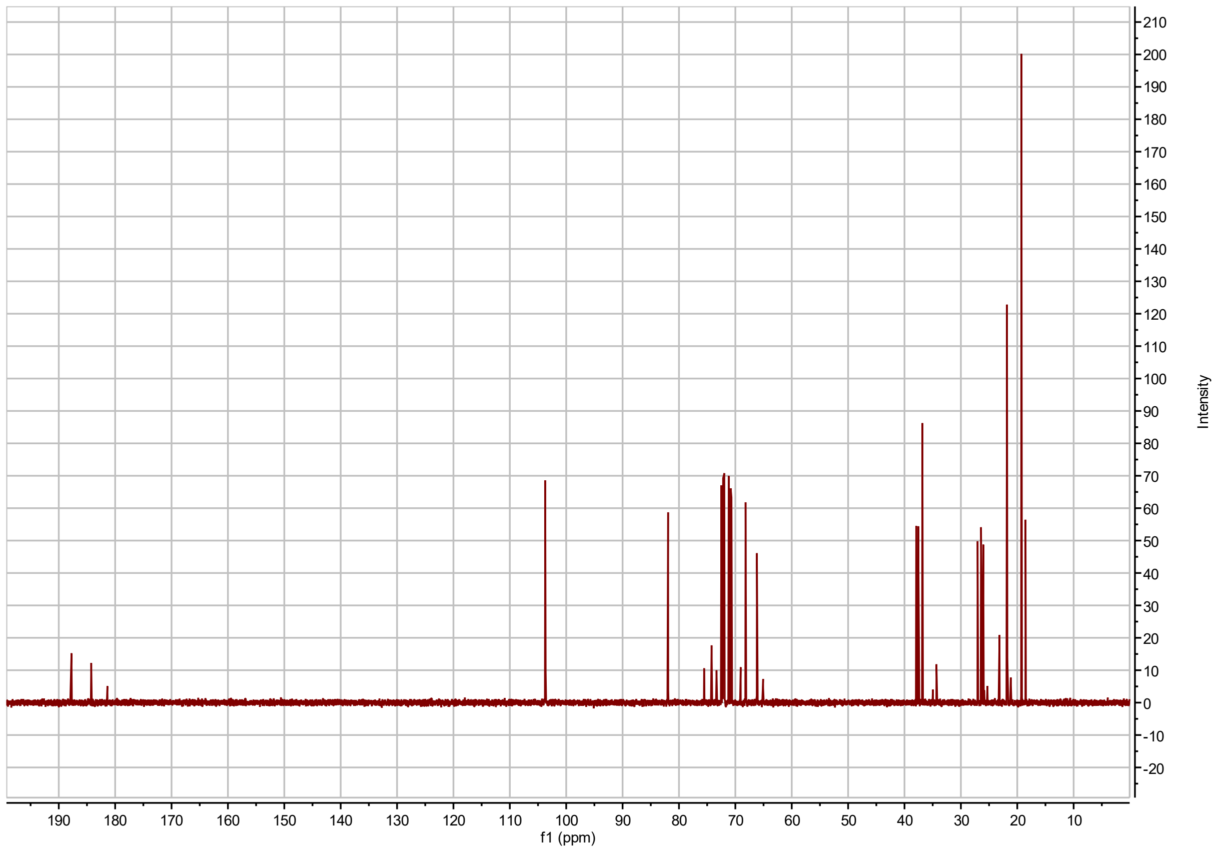


**4-*O*-arabinopyranosyl *myo*-inositol derived from saponified AI4:18(2,4,4,8) $^{13}C$ NMR**

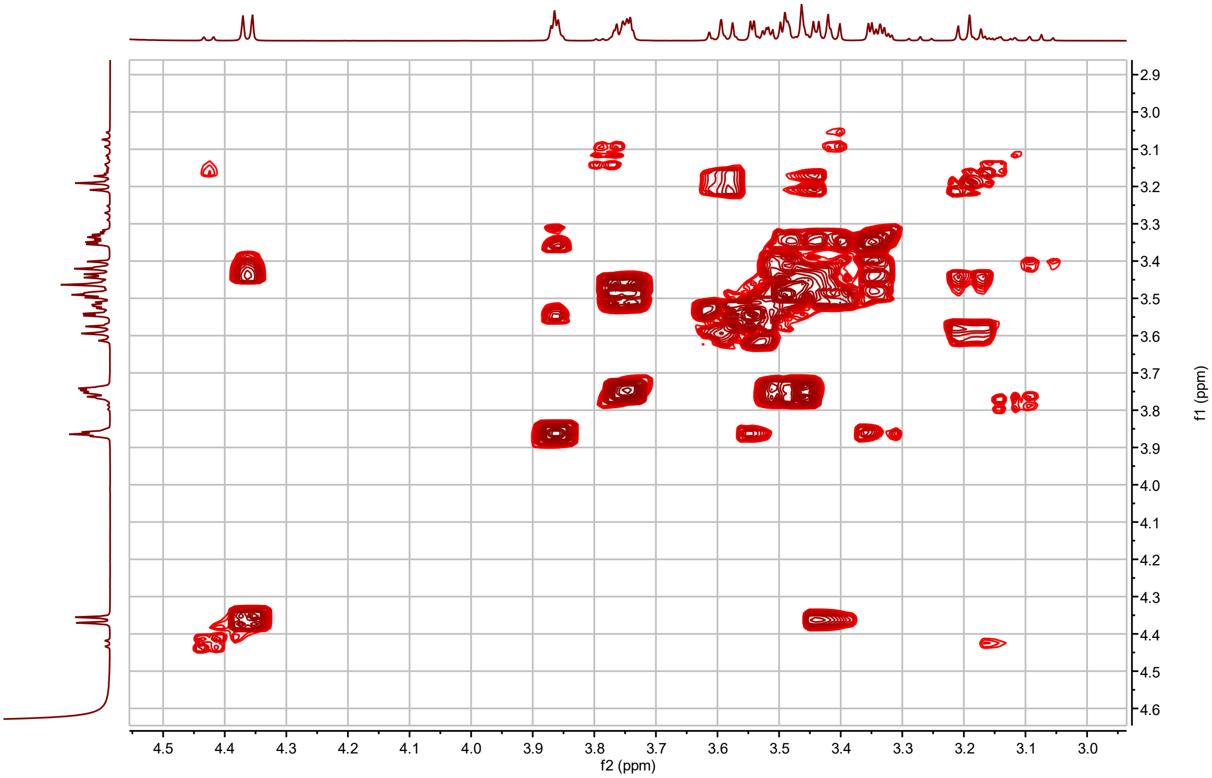


**4-*O*-arabinopyranosyl *myo*-inositol derived from saponified AI4:18(2,4,4,8) $^{1}$H-$^{1}$H COSY**

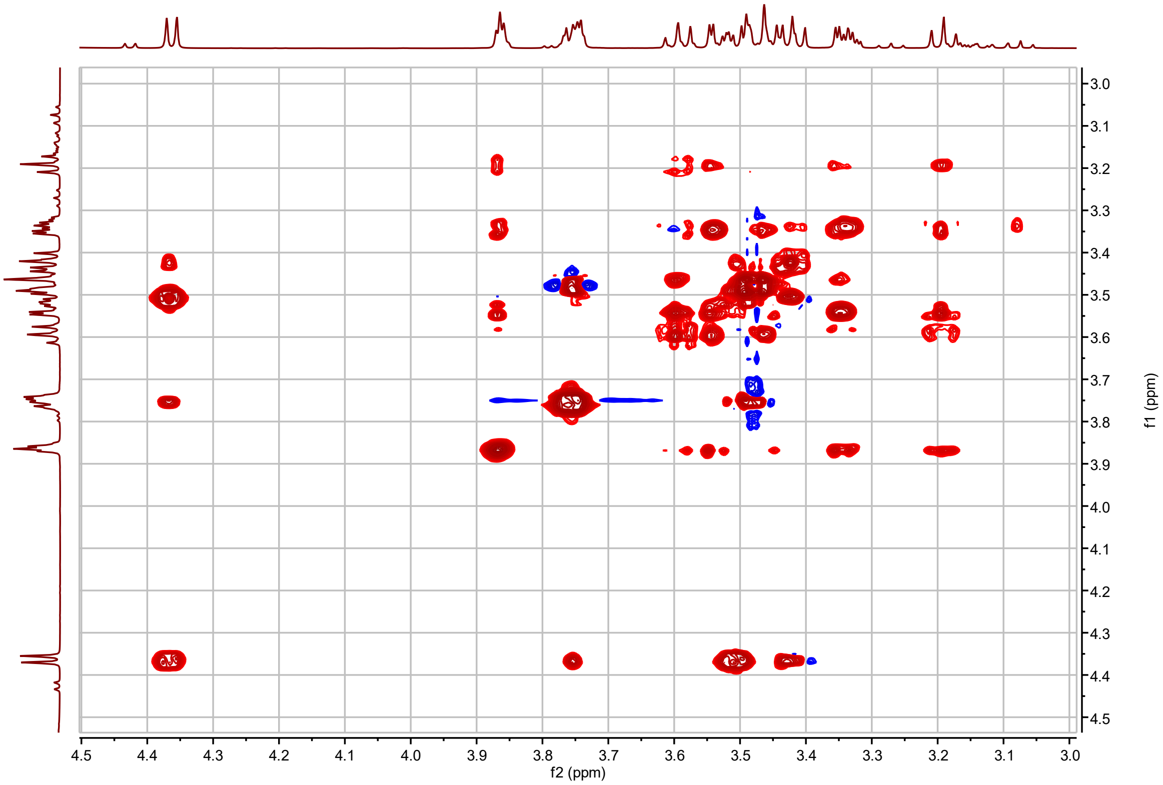


**4-*O*-arabinopyranosyl *myo*-inositol derived from saponified AI4:18(2,4,4,8) $^{1}$H-$^{1}$H TOSCY**

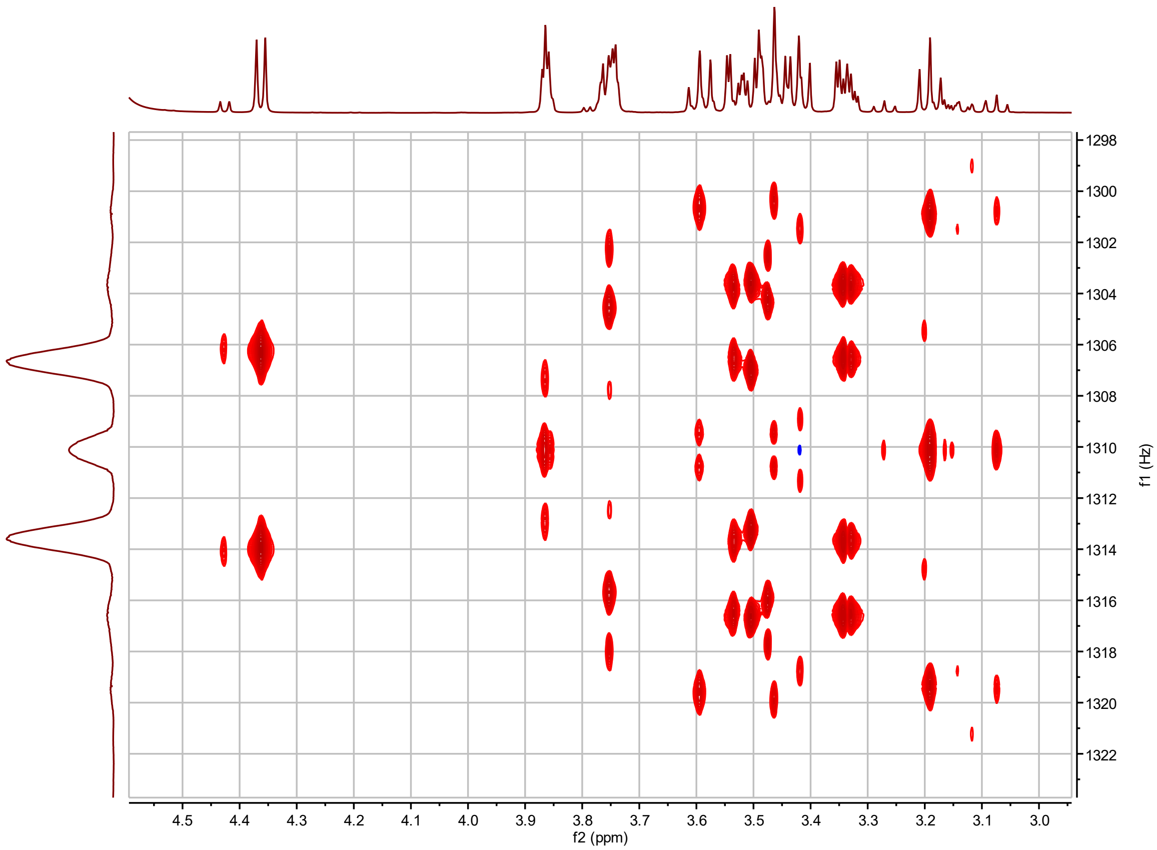


**4-*O*-arabinopyranosyl *myo*-inositol derived from saponified AI4:18(2,4,4,8) *J*-resolved**

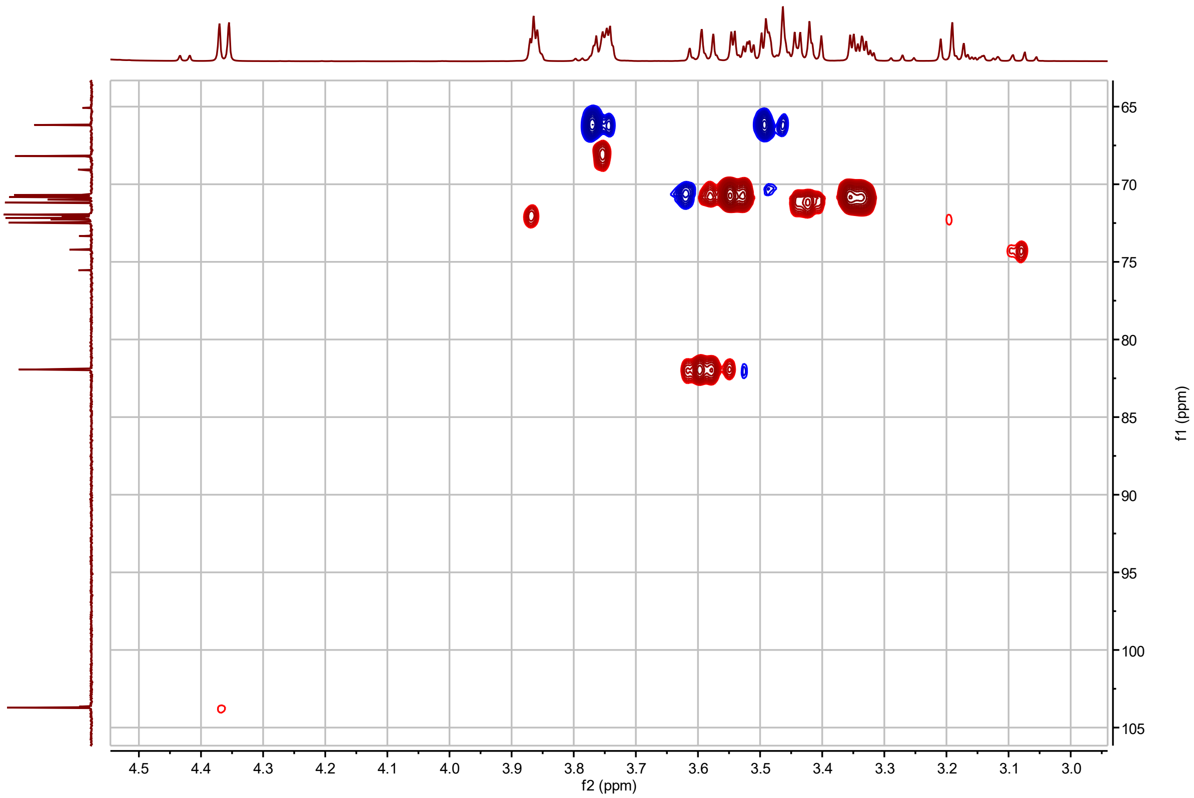


**4-*O*-arabinopyranosyl *myo*-inositol derived from saponified AI4:18(2,4,4,8) $^{1}H$-$^{13}C$ HSQC**

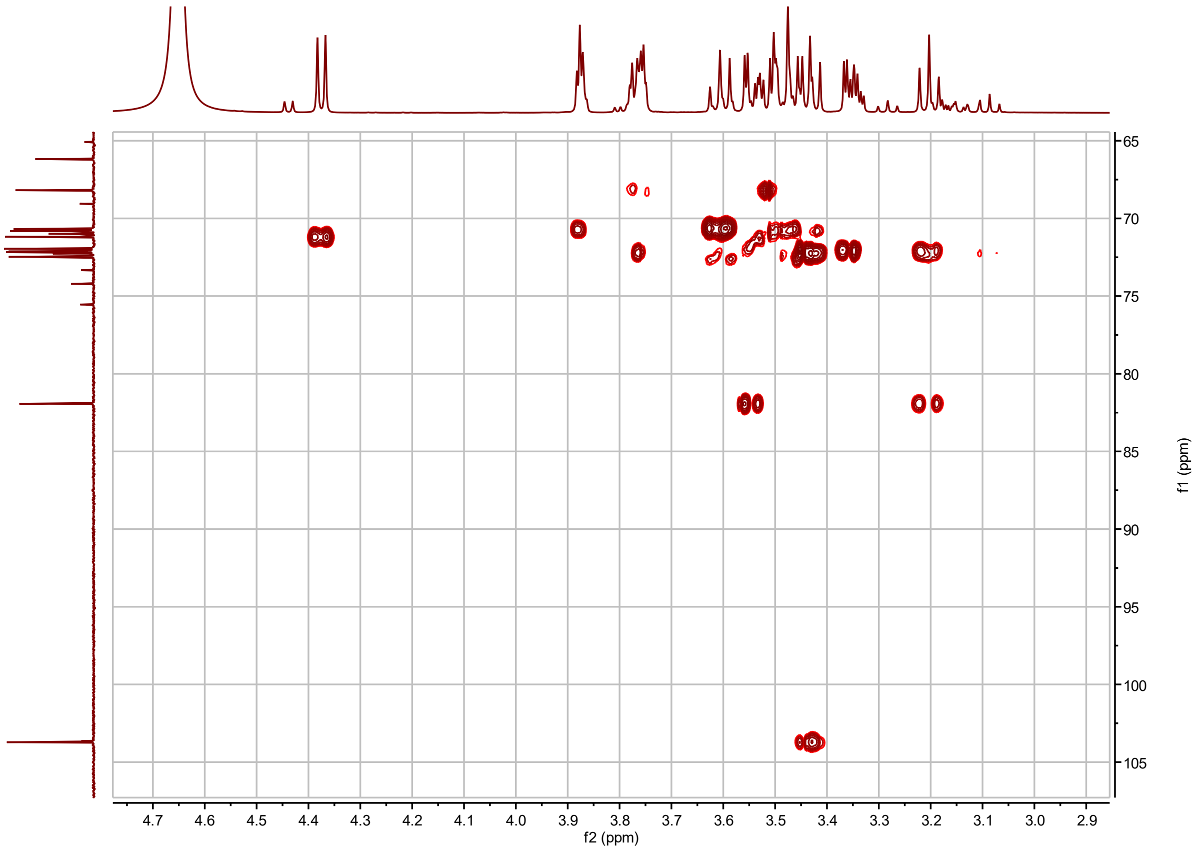


**4-*O*-arabinopyranosyl *myo*-inositol derived from saponified AI4:18(2,4,4,8) $^{1}H$-$^{13}C$ H2BC**

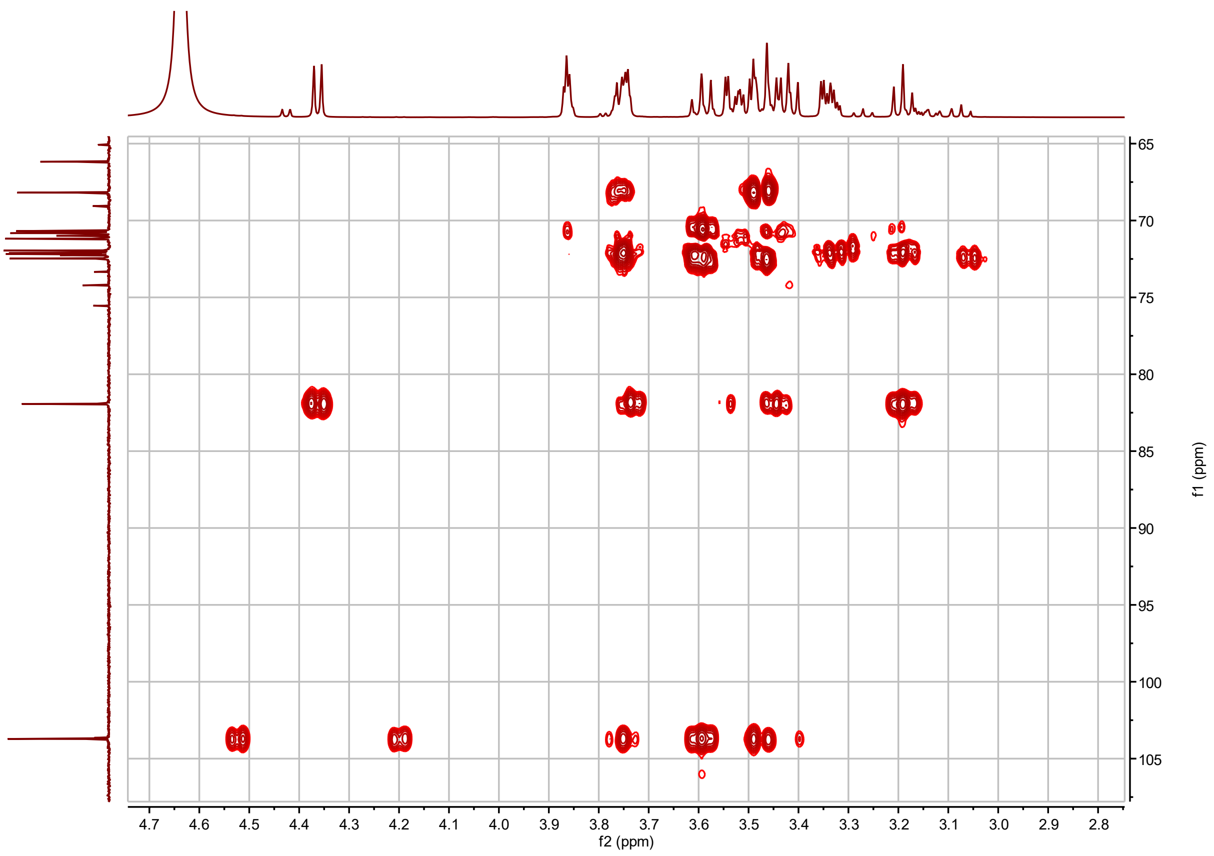


**4-*O*-arabinopyranosyl *myo*-inositol derived from saponified AI4:18(2,4,4,8) $^{1}H$-$^{13}C$ HMBC**

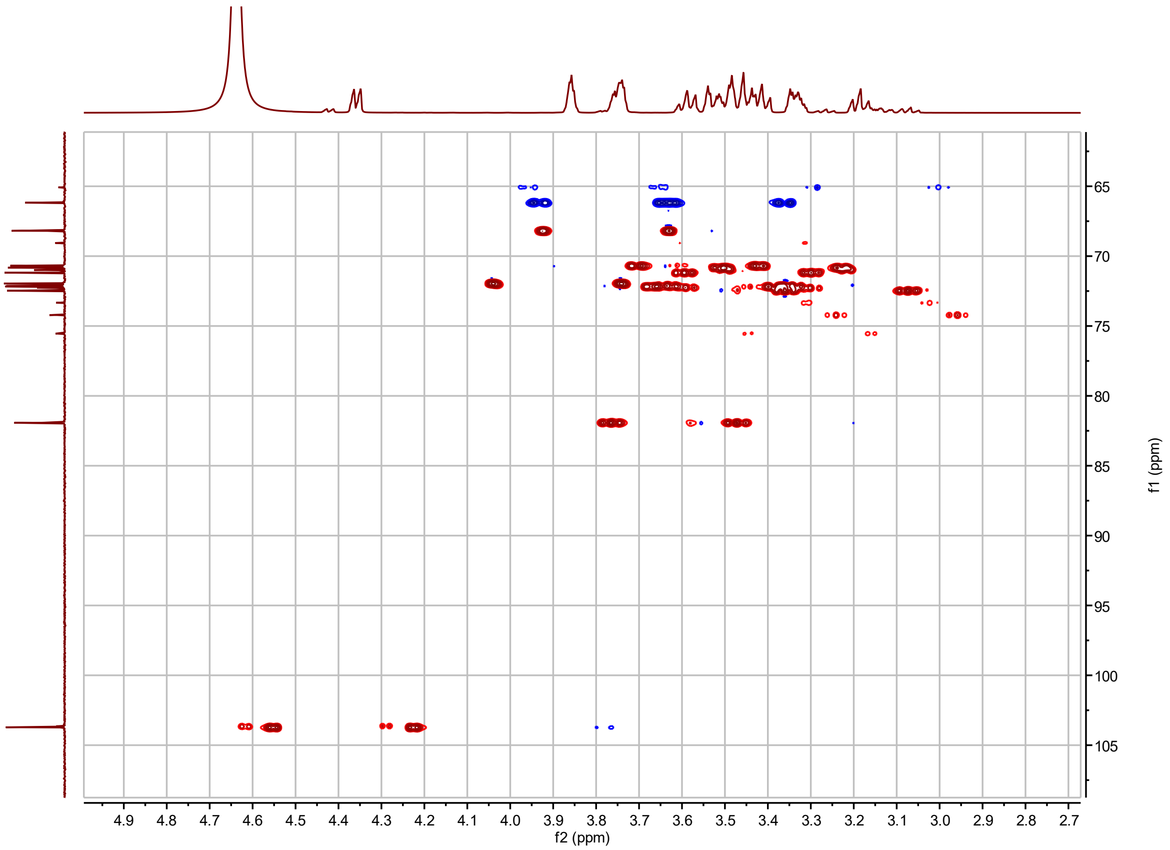


**4-*O*-arabinopyranosyl *myo*-inositol derived from saponified AI4:18(2,4,4,8) $^{1}H$-$^{13}C$ coupled HSQC**

**NMR metadata for the Agilent DDR2 500 MHz instruments.**

| | |
|---|---|
| Facility supervisor: | Dr. Daniel Holmes |
| Analyst | Paul D. Fiesel |
| Instrument location | MSU Max T. Rogers NMR Facility |
| Facility instrument title | Ahriman and Ormuzd |
| Manufacturer | Agilent |
| Field frequency lock | Acetonitrile-$d_3$; $D_2O$ |
| Additional solute | None |
| Solvent | $CD_3CN$: 600 μL; $D_2O$: 600 μL |
| Chemical shift standard | $CH_3CN$-$d_3$ ($\delta_H$ = 1.94 and $\delta_C$ = 118.70 ppm); $H_2O$-$d_2$ ($\delta_H$ = 4.36 ppm) |
| Concentration standard | None |
| Instrument | Agilent DDR2 500 MHz with 7600AS 96 autosamplers |
| Geographic location of instrument | 42.7288, -84.4745 |
| Magnet | 499.91 MHz |
| Probe | OneNMR Probe with Protune accessory for hands-off tuning |
| Acquisition software | VnmrJ 4.2A |
| Sample details | Kontes NMR tube, 8 in, Temperature @ 298K, no spinning |

**NMR metadata for the Varian Inova 600 MHz instrument.**

| | |
|---|---|
| Facility supervisor: | Dr. Daniel Holmes |
| Analyst | Paul D. Fiesel |
| Instrument location | MSU Max T. Rogers NMR Facility |
| Facility instrument title | Sobek |
| Manufacturer | Varian |
| Field frequency lock | Acetonitrile-$d_3$ |
| Additional solute | None |
| Solvent | $CD_3CN$: 600 μL |
| Chemical shift standard | $CH_3CN$-$d_3$ ($\delta_H$ = 1.94 and $\delta_C$ = 118.70 ppm) |
| Concentration standard | None |
| Instrument | Varian Inova 600 MHz |
| Geographic location of instrument | 42.7288, -84.4745 |
| Magnet | 599.77 MHz |
| Probe | Nalorac 5 mm PFG switchable probe pretuned for 1H, 13C |
| Acquisition software | VnmrJ 4.2A |
| Sample details | Kontes NMR tube, 8 in, Temperature @ 298K, no spinning |

**NMR metadata for the Bruker Avance NEO 600 MHz instrument.**

| | |
|---|---|
| Facility supervisor: | Dr. Daniel Holmes |
| Analyst | Paul D. Fiesel |
| Instrument location | MSU Max T. Rogers NMR Facility |
| Manufacturer | Bruker |
| Field frequency lock | Acetonitrile-$d_3$ |
| Additional solute | None |
| Solvent | $CD_3CN$: 600 μL |
| Chemical shift standard | $CH_3CN$-$d_3$ ($\delta_H$ = 1.94 and $\delta_C$ = 118.70 ppm) |
| Concentration standard | None |
| Instrument | Bruker Avance NEO 600 MHz NMR with shielded magnet |
| Geographic location of instrument | 42.7288, -84.4745 |
| Magnet | 600.32 MHz |
| Probe | 5 mm nitrogen cryogenic HCN Prodigy probe |
| Acquisition software | TopSpin 4.1.1 |
| Sample details | Kontes NMR tube, 8 in, Temperature @ 298K, no spinning |

**NMR metadata for the Bruker Avance NEO 800 MHz instrument.**

| | |
|---|---|
| Facility supervisor: | Dr. Daniel Holmes |
| Analyst | Paul D. Fiesel |
| Instrument location | MSU Max T. Rogers NMR Facility |
| Manufacturer | Bruker |
| Field frequency lock | Acetonitrile-$d_3$ |
| Additional solute | None |
| Solvent | $CD_3CN$: 600 µL |
| Chemical shift standard | $CH_3CN$-$d_3$ ($\delta_H$ = 1.94 and $\delta_C$ = 118.70 ppm) |
| Concentration standard | None |
| Instrument | Bruker Avance NEO 800 MHz NMR with shielded magnet |
| Geographic location of instrument | 42.7288, -84.4745 |
| Magnet | 800.33 MHz |
| Probe | 5 mm helium cryogenic HCN probe |
| Acquisition software | TopSpin 4.1.1 |
| Sample details | Kontes NMR tube, 8 in, Temperature @ 298K, no spinning |
